# Probing the KRas Switch II Groove by Fluorine NMR Spectroscopy

**DOI:** 10.1101/2022.07.15.500267

**Authors:** D. Matthew Peacock, Mark J. S. Kelly, Kevan M. Shokat

## Abstract

While there has been recent success in the development of KRas^G12C^ inhibitors, unmet needs for selective inhibitors and tool compounds targeting the remaining oncogenic KRas proteins remain. Here, we applied trifluoromethyl-containing ligands of KRas proteins as competitive probe ligands to assay the occupancy of the switch II pocket by ^19^F NMR spectroscopy. Structure-activity-relationship studies of probe ligands increased the sensitivity of the assay and identified structures that differentially detected each nucleotide state of KRas^G12D^. These differences in selectivity, combined with the high resolution of ^19^F NMR spectroscopy, enabled this method to be expanded to assay both nucleotide states of the protein simultaneously.

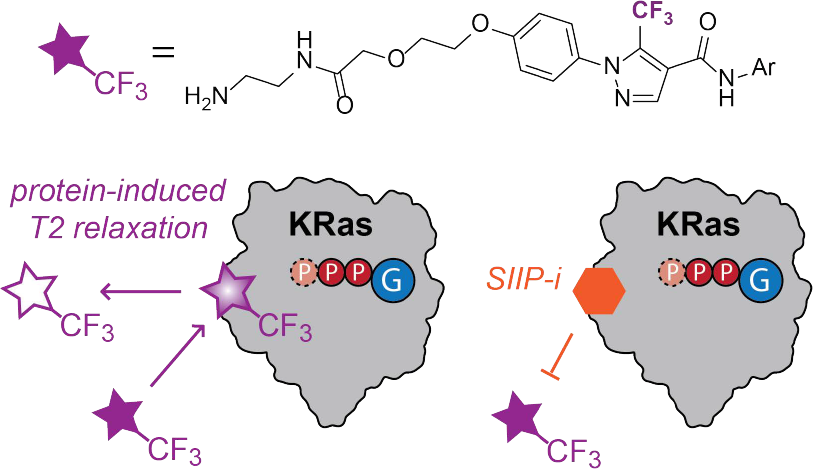

## INTRODUCTION

The proto-oncogene *KRAS* is among the most frequently mutated genes in human cancers, with mutations found in approximately 14% of patient samples.^1^ Its protein product (KRas) is a membrane-localized small GTPase responsible for cell growth and proliferation signaling through its effectors RAF and PI3K.^2^ Transforming mutations in *KRAS* (most commonly at G12, G13, or Q61) result in an increased cellular proportion of active GTP-bound KRas, typically by impairing the hydrolysis reaction of GTP.

Recent drug discovery campaigns relying on covalent chemistry have resulted in the first FDA-approved *KRAS* inhibitor sotorasib (AMG510), which selectively targets KRas proteins containing a G12C mutation.^3–12^ Sotorasib has weak reversible affinity to a cryptic pocket on the protein, termed the Switch II-pocket (SIIP), and relies on an irreversible covalent reaction with the mutant cysteine for its affinity and selectivity. In contrast, adagrasib (MRTX849) bears a weaker electrophile but possesses stronger reversible affinity to the same SIIP binding site (*K*_i_ = 4 μM).^13^ Further structure activity relationship (SAR) studies of the adagrasib scaffold led to MRTX1133, the first cell-active KRas^G12D^ SIIP inhibitor, in which the non-covalent interactions were optimized to achieve sub-picomolar binding affinity.^14^ Despite these recent successes, selective inhibitors for the protein products of many Ras-family oncogenes are still unknown, and tool compounds for the development of probes and assays are limited.

Protein-observed nuclear magnetic resonance (NMR) spectroscopy has proven to be a powerful tool in drug discovery generally and in the study of KRas protein-ligand interactions specifically.^15–17^ Furthermore, the two nucleotide-states of KRas can be resolved by protein-observed NMR spectroscopy, enabling nucleotide-cycling reactions and nucleotide-state-specific binding to be directly observed in mixed samples containing both GDP and GTP.^18–21^ While information-rich, protein-observed experiments are burdened by the requirements of high protein concentrations, long acquisition times, and isotopic labels. Ligand-observed NMR-spectroscopy alleviates these burdens; well-validated “probe” ligands can be applied to assay properties of the protein at lower concentrations, with faster acquisition times, and without the need for isotopic labelling.

In this study, we applied trifluoromethyl-containing ligands to the KRas switch II groove as probes to assay KRas proteins in both nucleotide-states by ^19^F NMR spectroscopy. One-dimensional ^19^F Carr-Purcell-Meiboom-Gill (CPMG1D) experiments were used to detect changes in the probes’ rates of transverse relaxation resulting from binding to the KRas protein. These probes were applied to assay competitive binding by SIIP-targeted inhibitors such as MRTX849, MRTX1133, and the cyclic peptide KD2.

## RESULTS AND DISCUSSION

### Reversible binding to the switch II groove is observed by NMR spectroscopy

We considered that a suitable probe structure to assay the switch II pocket of KRas proteins by ligand-observed NMR spectroscopy would have the following characteristics: (1) short residence time and weak affinity (*K*_D_ 10^−3^-10^−5^ M) to enable averaging of bound and unbound populations, (2) affinity for each of the two nucleotide states, and (3) an appropriate NMR handle to enable sensitive detection of the probe molecule under dilute conditions (≤100 μM). A previously reported disulfide-tethering screen against a KRas mutant containing an engineered cysteine (M72C) yielded a fragment (2C07, **1**) (Figure 1A) that we considered likely to serve as a starting point for ligand design to meet these requirements.^22^ This fragment occupied a site known as the switch II groove (SIIG), lying along a shallow lipophilic channel formed between α3 and SII. This disulfide fragment (**1**) and an acrylamide derived from it (**2**) reacted with KRas^M72C^ in both its inactive state (bound to GDP) and its active state (bound to GNP, a non-hydrolyzable GTP analog). Furthermore, the trifluoromethylpyrazole moiety in these compounds is an ideal structure for ligand-observed ^19^F NMR techniques; the three equivalent fluorines are observed as a strong singlet without any significant J-coupling due to the absence of nearby spin-active nuclei. The lack of biological fluorines greatly simplifies the analysis of ^19^F NMR spectra of protein-ligand mixtures when compared to ^1^H NMR spectra.^23^ However, the reversible affinities of these compounds to proteins lacking the M72C mutation were unconfirmed.

**Figure 1.**
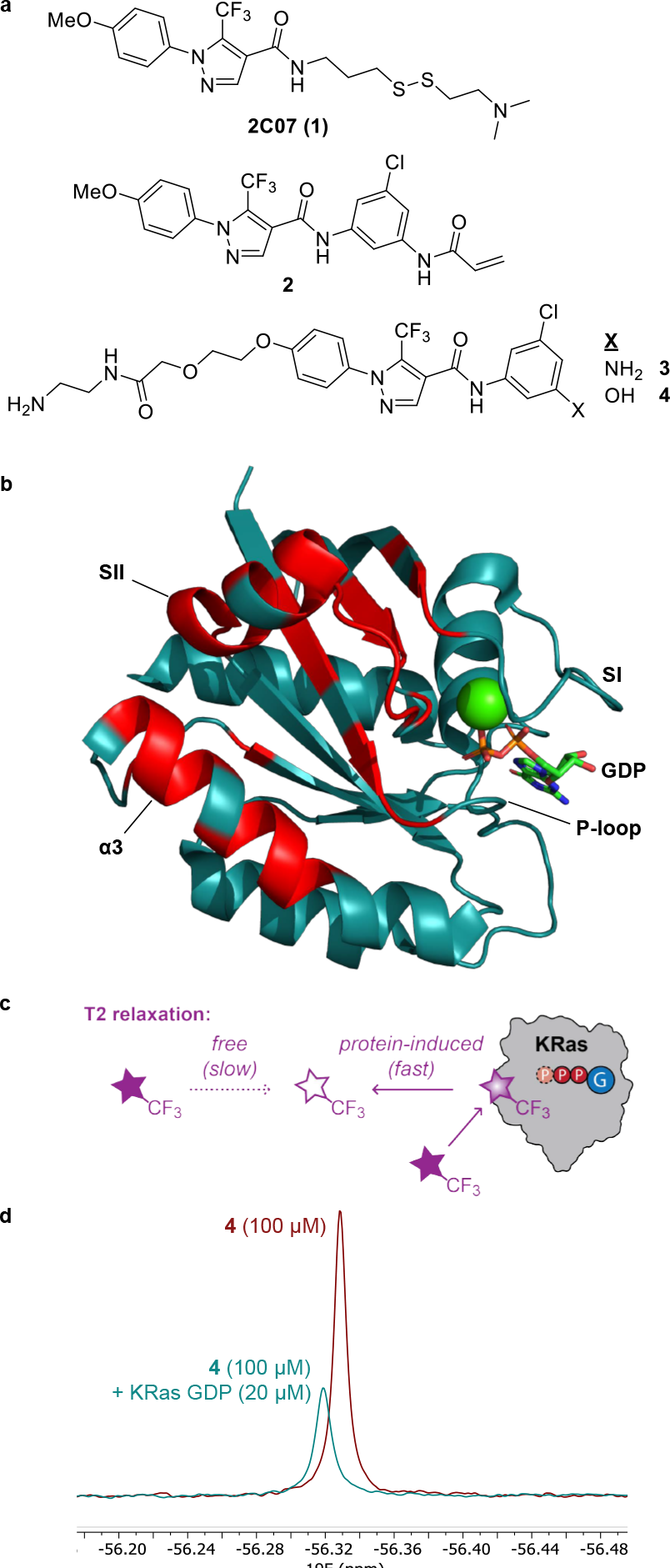
Solubilized derivatives of 2C07 bind the KRas SIIG. **a**, chemical structures of SIIG binders. **b**, top (>1σ) CSPs caused by 4 marked in red on KRas GDP structure 4LPK. **c**, cartoon depiction of KRas-induced T2 relaxation in a probe ligand. **d**, ^19^F CPMG NMR spectra (160 ms) of 4 in the absence (red) and presence (cyan) of KRas GDP.

Acrylamide **2** has very low aqueous solubility, and initial investigations of its reversible affinity were hindered by the formation of aggregates. To improve the solubility of **2**, the solvent-exposed methyl ether was extended into a polar solubilizing tail. The acrylamide was removed because oncogenic KRas proteins do not possess the engineered M72C mutation. These modifications resulted in compounds **3** and **4** (Fig 1A). Binding of **3** or **4** to uniformly ^15^N-labelled KRas^WT^ and KRas^G12D^ GDP 1-169 was observed by HSQC NMR spectroscopy, and perturbed peaks were mapped onto the previously determined structure of KRas GDP (Fig 1B, Supplementary Fig 1, PDB 4LPK). The largest of these perturbations are in the SII and α3 regions, which is consistent with expectations based on the structure of the KRas^M72C^**-1** GDP complex (PDB 5VBM).^22^ Smaller perturbations were observed in the dynamic P-loop and switch I (SI) regions, indicating that some conformational change is also induced by these binders. However, the reversible affinities of **3** and **4** were very weak; a 1 mM concentration of either ligand did not saturate the binding site (Supplementary Fig 1C).

### Ligand-observed CPMG NMR spectroscopy detects oncogenic KRas mutant proteins

We then sought to determine whether these SIIG-binders could serve as probe ligands to detect KRas proteins by ^19^F NMR spectroscopy. One-dimensional CPMG experiments [90-(τ-180-τ)_*n*_], in which a spin-echo with delay τ is looped *n* times, attenuate signal intensity as a function of the transverse relaxation rate (R_2_) and total spin echo time (2·τ·*n*). Prior studies have shown that reversible binding of proteins to fluorine-containing small molecules induces an increase in the observed ^19^F R_2_ at long values of τ, which maximize the exchange contribution to R_2_.^24–25^ Applying a 160 ms (τ = 20 ms, *n* = 4) CPMG filter to the ^19^F NMR spectrum of **4** (100 μM) in the presence of KRas GDP 1-169 resulted in a decrease in the measured integral (I) compared to the same spectrum acquired in the absence of protein (I_0_) (Fig 1C-D). However, these experiments required a relatively high concentration (20 μM) of protein to significantly reduce the integral of **4**. Further modifications to the probe structure were required to improve its binding affinity and the sensitivity of the ^19^F CPMG NMR assay.

To improve the sensitivity of this method for oncogenic KRas proteins and to explore the relationship between structure and nucleotide-state specificity, we synthesized a series of derivatives of **3** and **4** with varied structure at the aryl ring expected to be buried deepest within the pocket, and these derivatives were evaluated by CPMG NMR spectroscopy against both nucleotide states of KRas^G12D^ (6 μM) (Fig 2A-B). From this series, five compounds (**9, 12, 14, 15**, and **17**) showed increased sensitivity compared to **3** and **4**. Compound **9**, an isomer of **4** in which the hydroxy group is attached at C-4 of the aryl ring, specifically detected the GDP-state of KRas^G12D^. Compounds **12**, bearing a methyl group at C-2, and **17**, in which the phenyl ring is replaced by a naphthyl, detected both nucleotide states with similar sensitivity. Probe **9** relaxed more slowly in the absence of protein (R_2,free_ = 2.4 Hz) than did compounds containing the C-5 hydroxy group (R_2,free_ = 6.3 Hz for **4**, 4.4 Hz for **12**, and 5.0 Hz for **17**), increasing the sensitivity and dynamic range of integral measurements with this probe (Supplementary Fig 2). Furthermore, probe **9** was confirmed to be a tighter binder to the SIIG of U-^15^N KRas^G12D^ GDP by HSQC NMR compared to **4** (Supplementary Fig 3).

**Figure 2.**
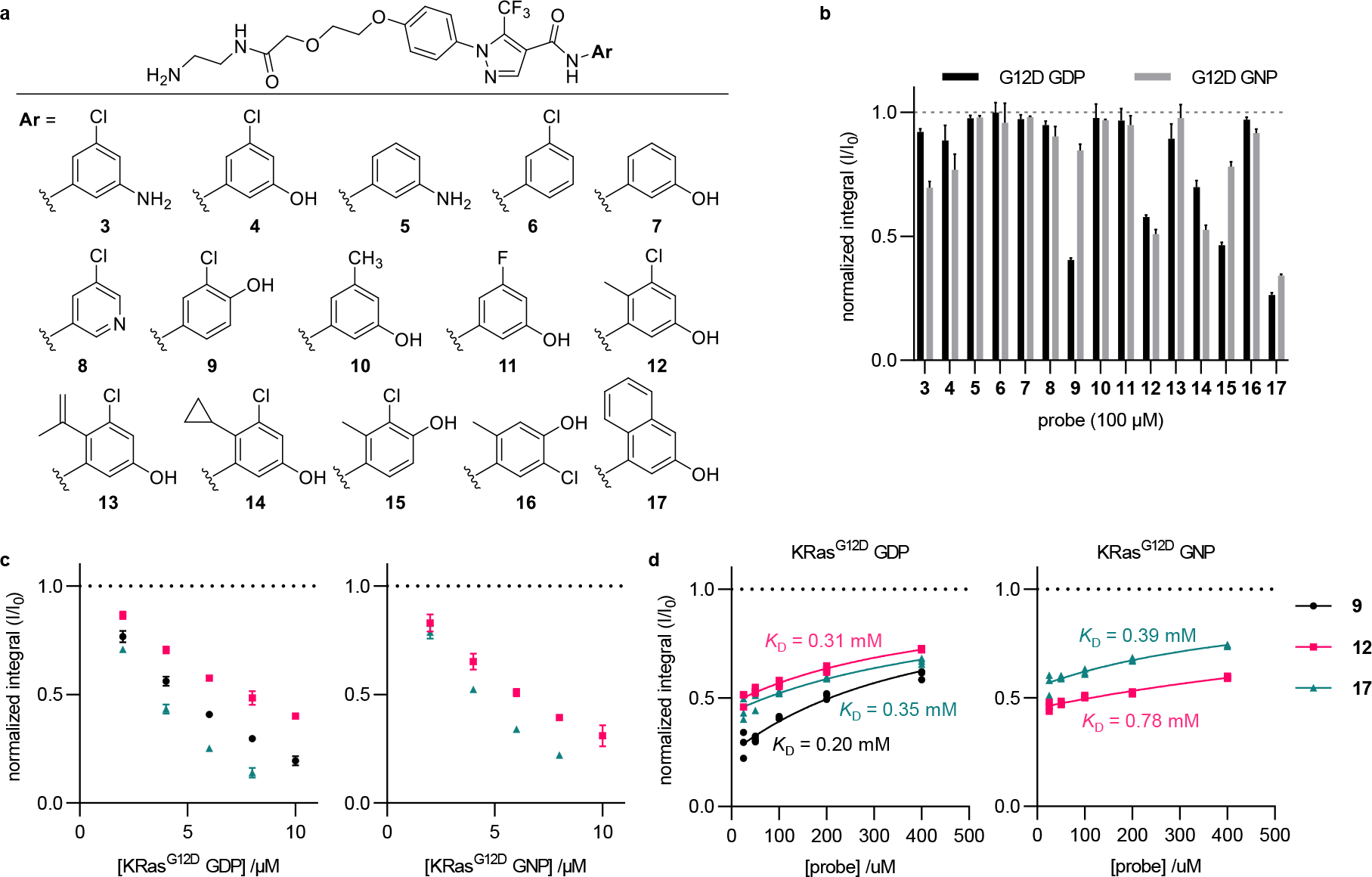
SAR of SIIG binders enables more sensitive detection of KRas^G12D^. **a**, chemical structures of SIIG binders. **b**, normalized integrals (I/I0) from^19^F CPMG NMR spectra (160 ms) of probes (100 μM) with KRas^G12D^ (6 μM); values are means ± errors propagated from SD (n=3). **c**, effect of varying protein concentration on normalized integrals with KRas^G12D^ (6 μM); values are means ± errors propagated from SD (n=3). **d**, effect of varying probe concentration on I/I0; [P]^0^ = 6 μM for probes **9** and **12**, 3 μM for probe **17**; individual points shown (n=3); data fit to equation 1.

The probe and protein concentrations were varied to determine the dependence on each binding partner. The normalized integrals of the probes decreased exponentially with increasing protein concentrations (Fig 2C, Supplementary Fig 2C-D). The normalized integrals (I/I_0_) increased with increasing probe concentrations, and *K*_D_ values for the probe-protein binding were extracted by fitting the data to equation 1 (Fig 2D). Probe **9** bound the GDP-state more strongly than did probes **12** or **17** (*K*_D_ 0.20 mM); however, we did not observe a clear correlation between these three probes’ binding affinities and sensitivities to detect KRas^G12D^ proteins.

Probes **9, 12**, and **17** were evaluated against both the inactive GDP and active GNP nucleotide-states of a panel of KRas proteins (three common mutations at G12 and three common mutations at Q61) to determine whether this method can be generalized across the protein products of the most frequent *KRAS* oncogenes (Fig 3). Probe **9** most sensitively detected the GDP-state of KRas^G12D^ and more weakly detected the remaining oncogene products excepting KRas^Q61R^ GNP (Fig 3A). Probes **12** and **17** detected both nucleotide states of all oncogene products tested, albeit with weaker sensitivity for KRas^Q61L^ GDP and KRas^Q61R^ GNP than for the others. The closely related protein HRas 1-166 contains only one residue difference within the SIIG binding site (Q95 in HRas vs H95 in KRas). HRas was only weakly detected by this method; however, the results with HRas^Q95H^ closely matched those obtained with KRas^WT^. Similarly, a recently identified mutation that confers resistance to adagrasib (Y96C) greatly weakened detection of KRas^G12D^.^26^

**Figure 3.**
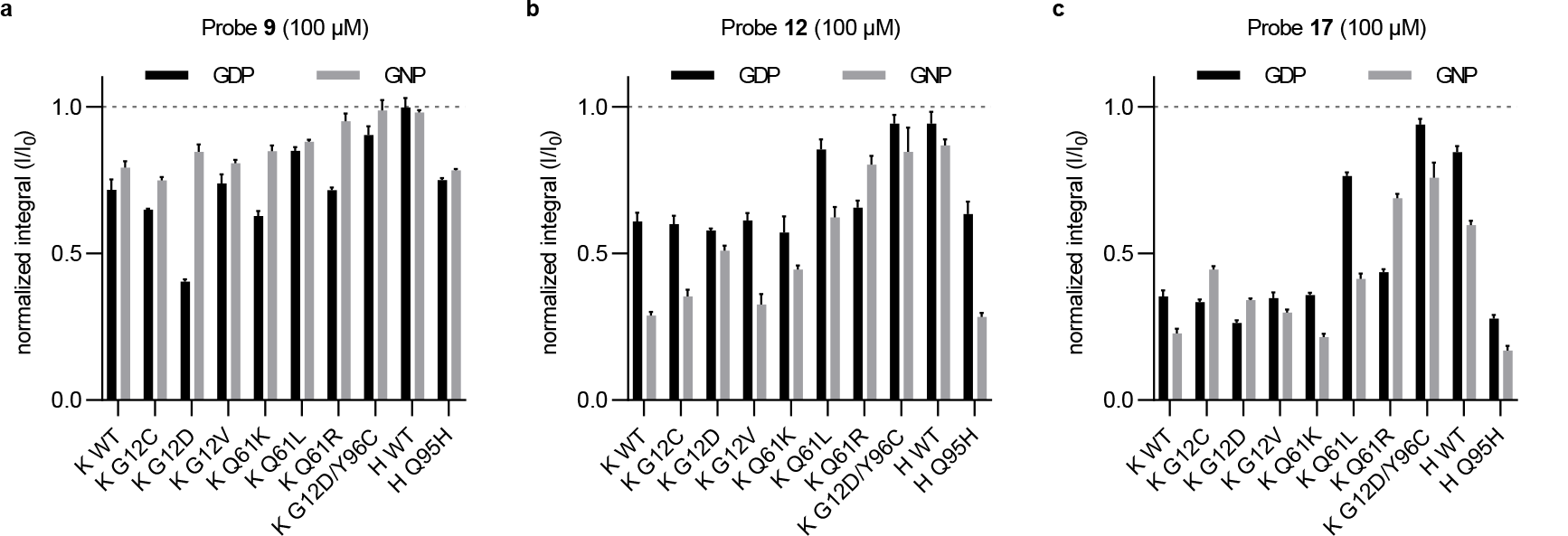
^19^F CPMG NMR spectroscopy detects a variety of oncogenic KRas proteins. Normalized integrals (I/I0) of 100 μM **9 (a), 12** (**b**), or **17** (**c**) in the presence of 6 μM KRas or HRas proteins from ^19^F CPMG NMR spectra (160 ms); values are means ± errors propagated from the standard deviations of I and I_0_ (n=3).

### SIIP-targeted inhibitors competitively displace SIIG-targeted ^19^F NMR probes

Having identified probe ligand structures that detected low micromolar concentrations of KRas proteins, we next sought to determine whether these probes could assay competitive reversible binding of ligands within or adjacent to the SIIG. Since the SIIG and SIIP are overlapping binding sites, we expected binding to either of them to be mutually exclusive (Supplementary Fig 4A). We measured the effect of MRTX849 on the normalized integral of probe **9** (100 μM) in the presence of KRas^G12D^ GDP (2 μM) (Fig 4A-B). MRTX849 displaced probe **9**, and the calculated fraction occupancy data were fit to equations 3 and 4 to extract the *K*_i_ of MRTX849 (2.9 μM). This result agrees well with the affinity expected based on kinetic data from its reaction with KRas^G12C^.^13^ In contrast, two G12C-targeted inhibitors based on different scaffolds (AMG510 and JDQ443) did not occupy the SIIP of KRas^G12D^ when tested at 60 μM (Supplementary Fig 4B-C).

**Fig 4.**
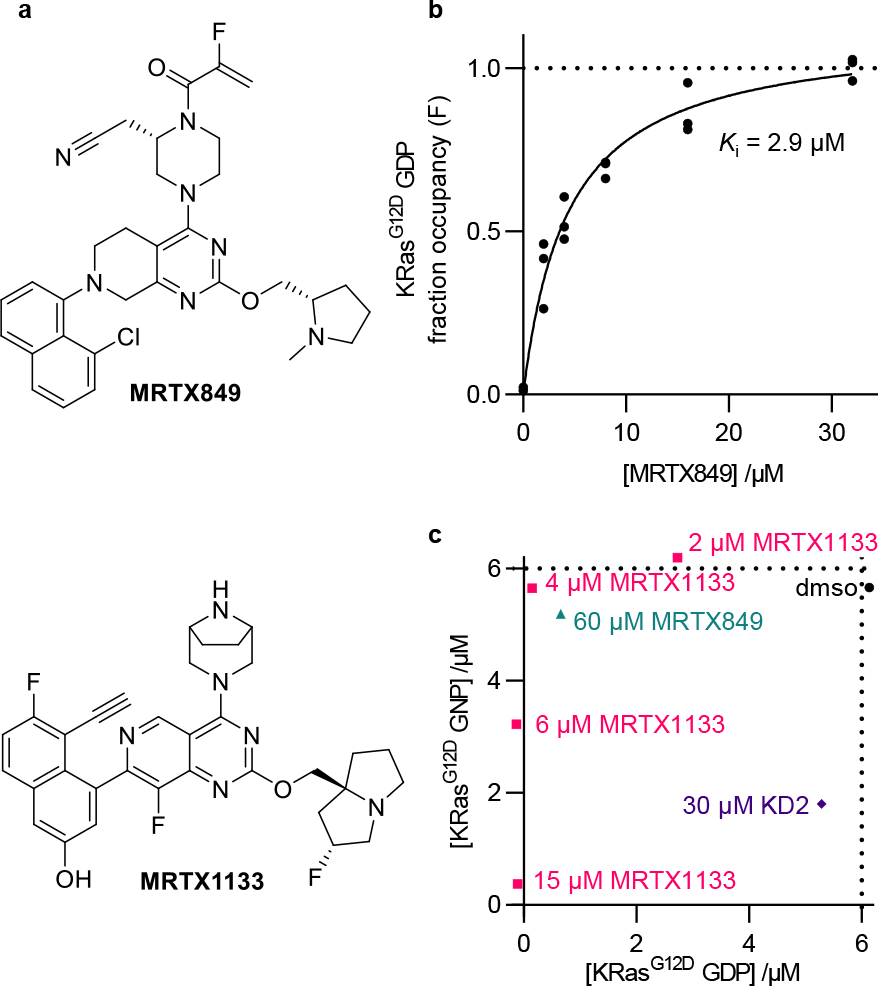
SIIP inhibitors compete with SIIG probes. **a**, chemical structures of MRTX849 and MRTX1133. **b**, fraction occupancy (F) of MRTX849 binding to KRas^G12D^ GDP (2 μM) calculated from ^19^F CPMG NMR spectra (320 ms) of **9** (100 μM) according to equations 2-4; individual points shown (n=3). **c**, simultaneous monitoring of both nucleotide states of KRas^G12D^ (6 μM each) with a mixture of **9** and **12** (50 μM each); concentrations were calculated with equations 5 and 6 from the means of R_2_ measurements (n=2-4).

Recognition of the two nucleotide-states of KRas proteins is a key property of inhibitors, relating both to an inhibitor’s mechanism of action and its ability to access constitutively active proteins. We sought to determine whether this ^19^F NMR method could be extended to assay the nucleotide-state of KRas^G12D^ and the nucleotide-state specificity of SIIP inhibitors. Since their resonances are resolved, and neither ligand saturates the binding site, the transverse relaxation rates of **9** and **12** could be measured simultaneously in a mixture. We tested whether this combination of probes could discriminate between the inactive GDP-state and the active and inactive conformations of the GTP state. The transverse relaxation rate of probe **9** exhibited a strong dependence of the nucleotide state and conformation of KRas^G12D^; R_2,**9**_ varied over 6 Hz between samples containing 6 μM of either KRas^G12D^ GDP, KRas^G12D^ GNP, KRas^G12D/T35S^ GNP (state 1 inactive conformation), or KRas^G12D^ GNP + RBD (state 2 active conformation) (Supplementary Fig 4D). Meanwhile, R_2,**12**_ varied relatively little (<2 Hz) among the same samples.

From this data, equations to calculate the individual concentrations of the GDP- and GNP-states in a mixture were determined (equations 5 and 6, Supplementary Fig 4D). Addition of the moderate-affinity binder MRTX849 (60 μM, GDP-state *K*_i_ = 2.9 μM) selectively occupied the GDP state of the protein (Fig 4C). The high-affinity binder MRTX1133 (GDP-state *K*_i_ < 1 pM) occupied both nucleotide states when added in excess (15 μM). However, sub-stoichiometric quantities (2, 4, and 6 μM) of MRTX1133 selectively occupied the GDP-state, consistent with the previously reported high GDP-state selectivity of a closely related structure.^21^ In contrast to the small molecule SIIP-binders, the cyclic peptide KD2 preferentially occupied the active GNP-state.^27^

## CONCLUSIONS

We have synthesized trifluoromethyl-containing ligands that bind to the SIIG of KRas proteins in both nucleotide-states, and we have shown that these compounds serve as ^19^F NMR spectroscopy probes for a variety of oncogenic KRas proteins. SAR studies identified modifications in the probe structure to improve the sensitivity and nucleotide-state selectivity of the assay. The probes were stoichiometrically competed by SIIP-targeted inhibitors, enabling their use to quantify the occupancy of the SIIP.

While many biochemical methods to assay competitive binding between two ligands are known, few allow simultaneous interrogation of two closely related proteins in a single sample, such as the two nucleotide states of KRas. Knowledge of the nucleotide-state selectivity of SIIP-inhibitors is likely important to understand their cellular engagement of constitutively activated KRas proteins, but current methods to directly query this selectivity are lacking. The method and probe ligands reported in this work enable the approximation of occupancy of both nucleotide-states from a single sample, allowing an inhibitor’s nucleotide-state selectivity to be assayed in a competitive manner.

The probe ligands described in this work can be used to detect KRas proteins at low μM concentrations with experiment times under 10 minutes. This sensitivity is sufficient to assay competitive binding, but the concentration of protein places a lower limit on quantifiable *K*_i_ values of competitors. The throughput and sensitivity of this method could both be further improved with continued efforts towards probe ligand design.

## Acknowledgments

We thank Ziyang Zhang (UCSF) for his advice and donation of samples used in this work (KRas^G12D/T35S^, TEV protease, RAF RBD, and KD2).

