## Supplementary Information for "Probing the KRas Switch II Groove by Fluorine NMR Spectroscopy"

*Supplementary Tables and Figures*

**Supplementary Table 1. Parameters from non-linear regressions to calculate $K_D$ and $K_i$.**

| figure | entry | equation | constrained constants | unconstrained | extracted value | 95% confidence interval |
|---|---|---|---|---|---|---|
| 2D | [**9**] dependence, $KRas^{G12D}$ GDP | 1 | $[P]_0$=6 µM, *t*=0.16 s | A=297 Hz | $K_D$=204 µM | 158-267 µM |
| 2D | [**12**] dependence, $KRas^{G12D}$ GDP | 1 | $[P]_0$=6 µM, *t*=0.16 s | A=240 Hz | $K_D$=307 µM | 256-374 µM |
| 2D | [**17**] dependence, $KRas^{G12D}$ GDP | 1 | $[P]_0$=3 µM, *t*=0.16 s | A=614 Hz | $K_D$=352 µM | 260-493 µM |
| 2D | [**12**] dependence, $KRas^{G12D}$ GNP | 1 | $[P]_0$=6 µM, *t*=0.16 s | A=644 Hz | $K_D$=778 µM | 655-941 µM |
| 2D | [**17**] dependence, $KRas^{G12D}$ GNP | 1 | $[P]_0$=3 µM, *t*=0.16 s | A=491 Hz | $K_D$=393 µM | 302-526 µM |
| 4B | [MRTX849] dependence, $KRas^{G12D}$ GDP | 3 | none | B=1.12 | $K_{i,obs}$=4.37 µM | 3.25-5.80 µM |

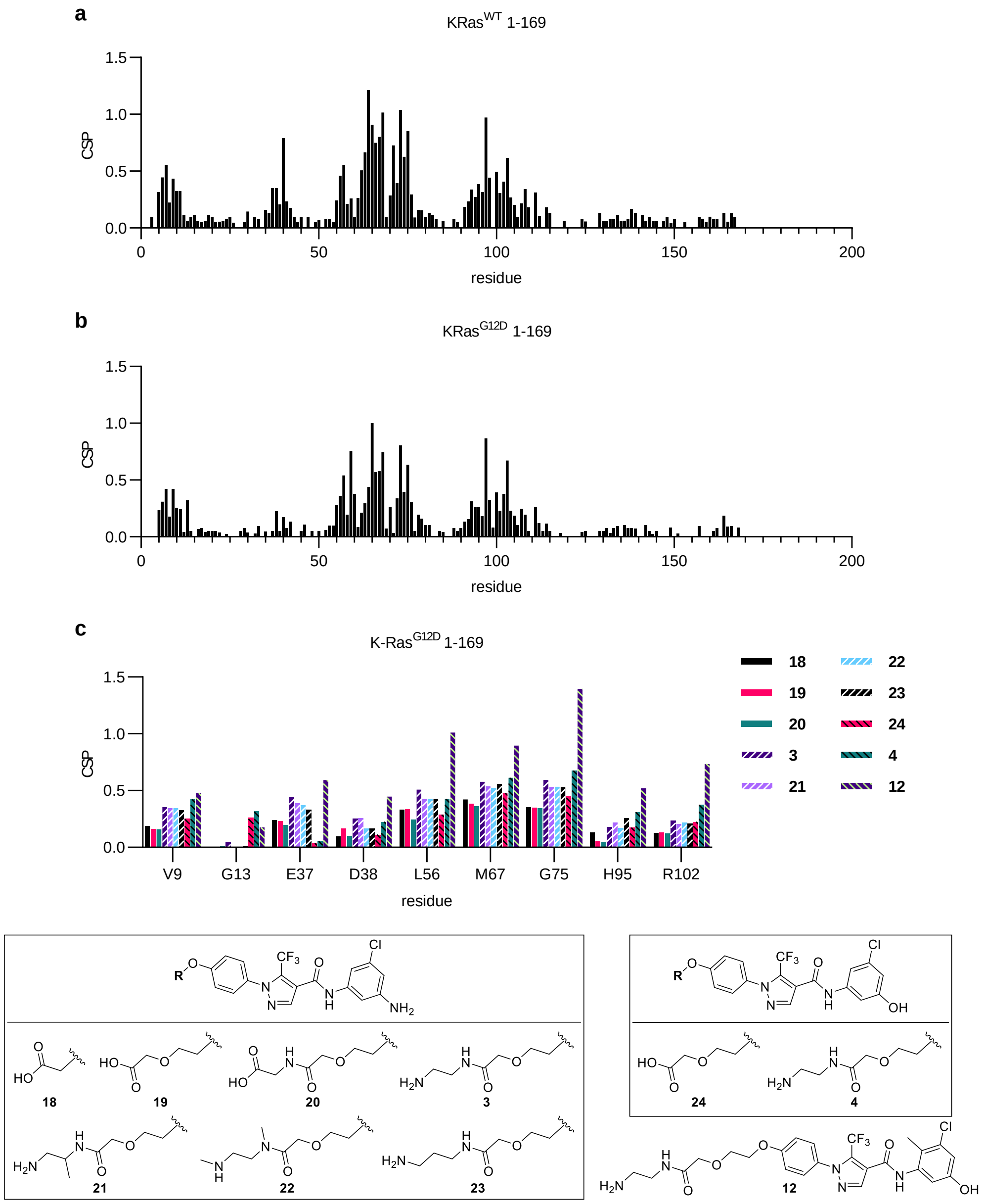

**Supplementary Fig 1. Effects of SIIG binders on the $^{1}$H-$^{15}$N HSQC NMR spectra of U-$^{15}$N KRas GDP 1-169.** Spectra recorded at pH 7.4 and 298 K with 10% dmso-$d_6$. $^{1}$H components of CSPs are scaled by 7 relative to $^{15}$N. **a-b**, CSPs observed in the spectra of WT (**a**) and G12D (**b**) proteins on the addition of **4** (1 mM). A threshold of 0.236 was set for inclusion in Fig 1b. **c**, effect of varying solubilizing group (1 mM compound) on the magnitude of CSPs, selected well-resolved and perturbed residues in P-loop (9, 13), SI (37, 38), interswitch (56), SII (67, 75), and α-3 (95, 102).

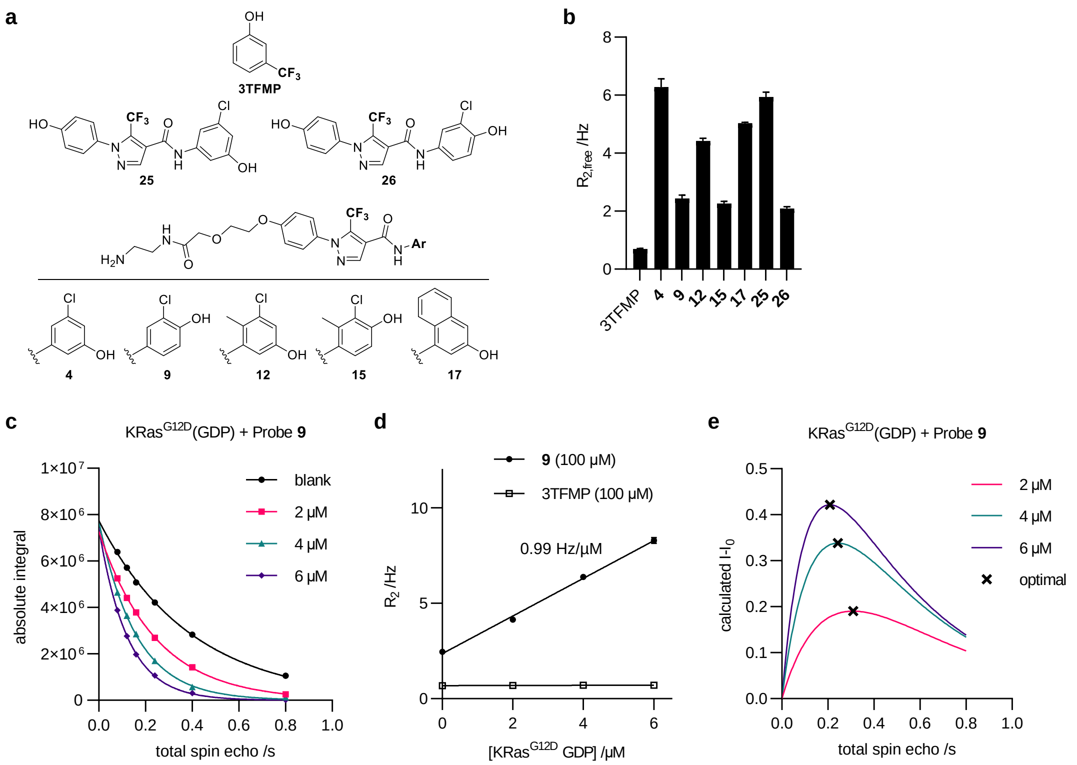

**Supplementary Fig 2. Rates of transverse relaxation measured by multi-point CPMG experiments. a**, chemical structures of solubilized probes (**4**, **9**, **12**, **15**, and **17**) and truncated derivatives (**25** and **26**) lacking the solubilizing group. **b**, effects of amide substituents and solubilizing group on the $^{19}$F NMR transverse relaxation rate ($R_{2,free}$) of trifluoromethylated probes (50 µM for **25** and **26**, 100 µM for others); $R_{2,free}$ measured by fitting absolute integrals to exponential decay functions of the total spin echo time (constant frequency); values are means ± SD with 6 timepoints per fit (n=3 for probes, n=15 for the internal standard 3TFMP). **c**, effect of varying [KRas$^{G12D}$ GDP] on the T2 relaxation curves of probe **9;** one example replicate for each concentration shown. **d**, effect of varying [KRas$^{G12D}$ GDP] on the observed transverse relaxation rate constants ($R_2$) of probe **9** and the standard 3-trifluoromethylphenol; values are means ± SD with 6 timepoints per fit (n=3). **e**, predicted differences in integrals ($I$-$I_0$) calculated from the rate constants shown in (**d**); the optimal spin echo periods that should be chosen to maximize signal-to-noise are marked with an **x**.

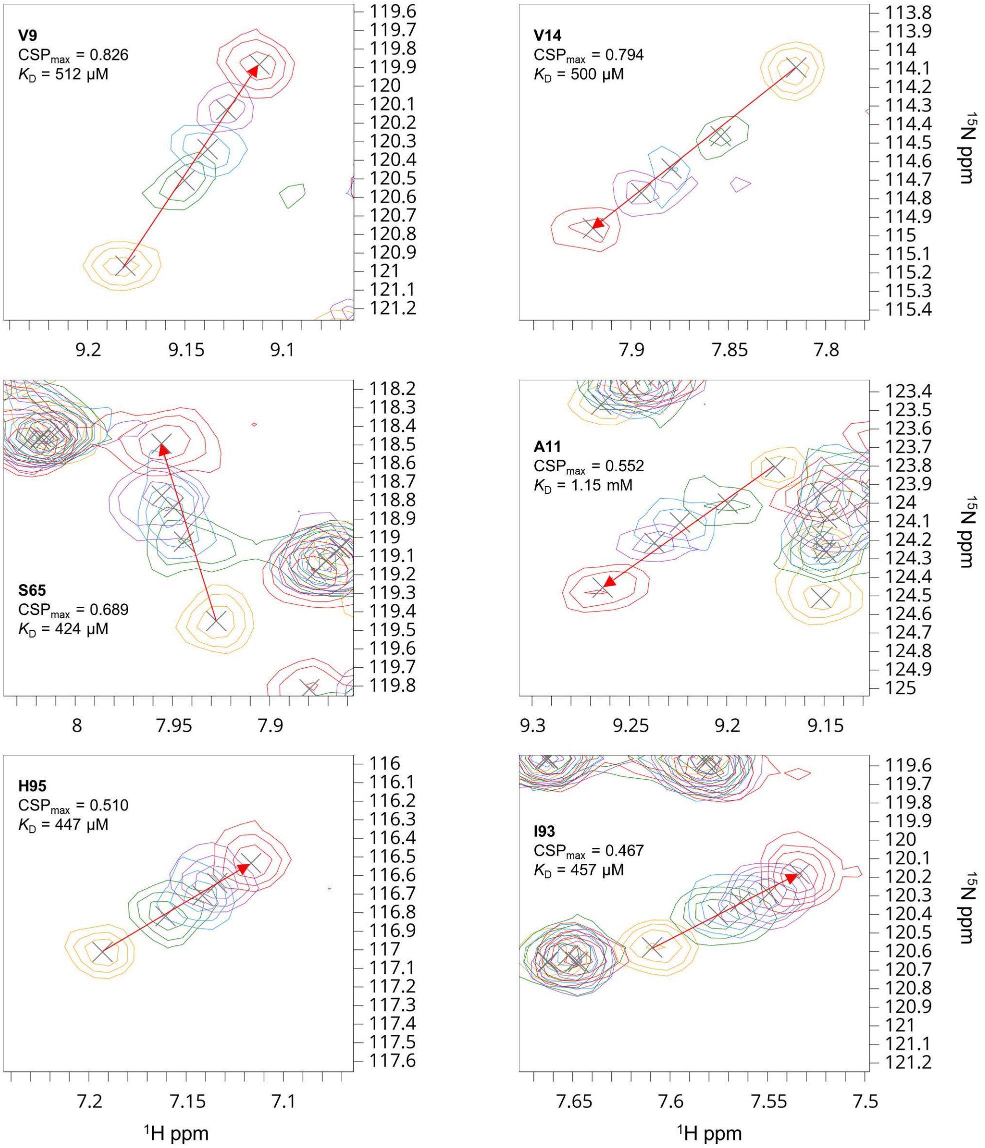

**Supplementary Fig 3. $K_D$ measurement of probe 9 binding to U-$^{15}$N-KRas$^{G12D}$ GDP 1-169 by HSQC NMR.** The six most-perturbed and well-resolved peaks were traced and fit to a single site binding curve with CCPNMR Analysis v3 to obtain an average $K_D$ of 0.58±0.28 mM; the most strongly perturbed peaks were too broadened to include in the analysis; $^{1}$H components of CSPs were weighted by 7 relative to $^{15}$N; spectra were acquired at pH 7.4 and 298 K with 10% dmso-$d_6$, 50 µM protein, and 0.20/0.33/0.50/1.0 mM **9**; each spectrum is a singlicate, the reported error is the SD from the six fit peaks.

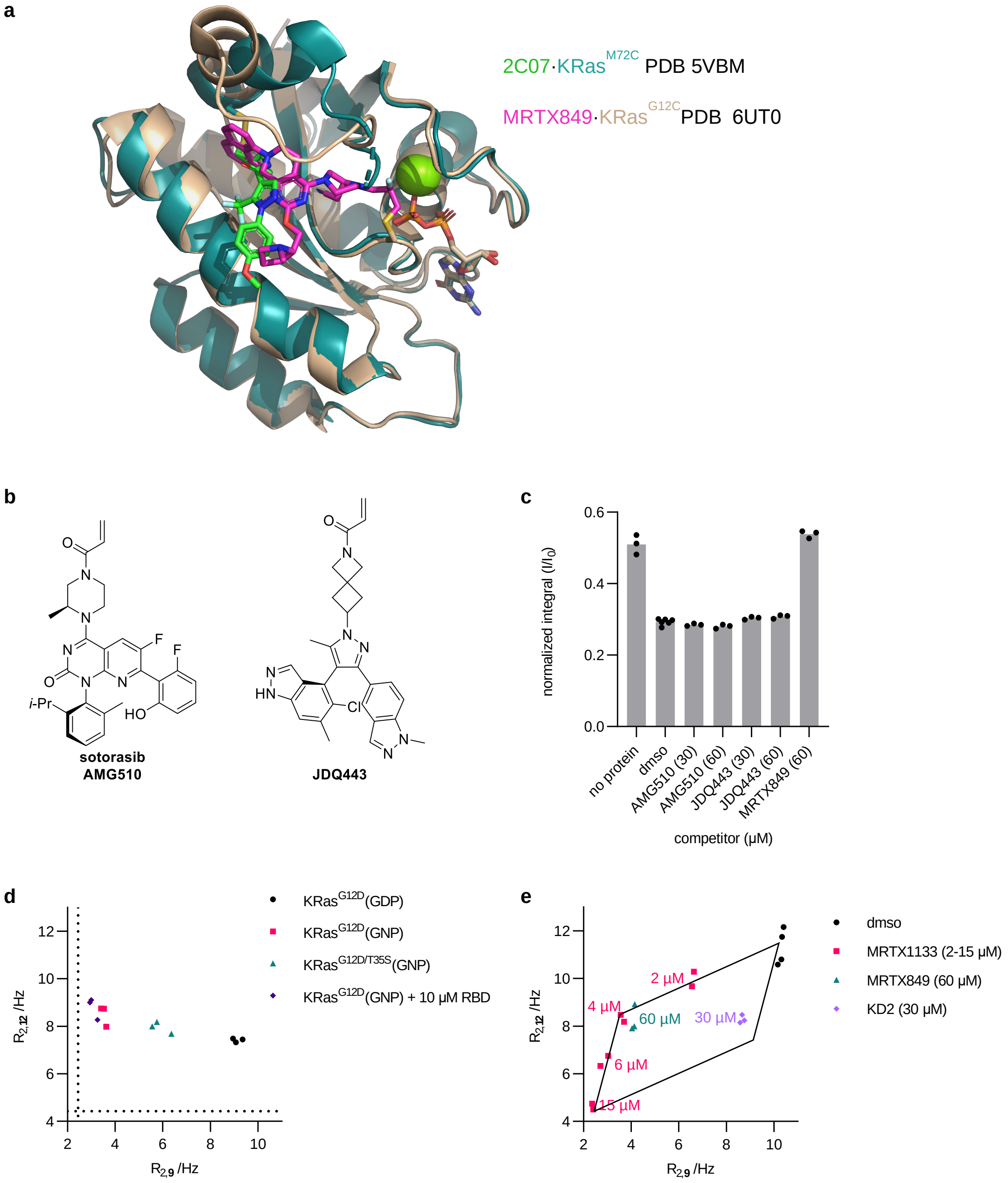

**Supplementary Fig 4. Design of a competitive assay to measure SIIP occupancy.** **a**, superimposed structures of covalently bound 2C07 (5VBM) and MRTX849 (PUT0), showing the expected overlap between the SIIG and SIIP binding sites. **b**, chemical structures of G12C-targeted SIIP inhibitors AMG510 and JDQ443. **c**, effects of SIIP-inhibitors on the integrals from $^{19}$F CPMG NMR spectra (320 ms) of probe **9** (100 µM) in the presence of KRas$^{G12D}$ (2 µM, GDP-bound); individual points shown (n=3 or 6). **d**, effect of the nucleotide state of KRas$^{G12D}$ (6 µM) on the transverse relaxation rates of probes **9** and **12** in mixture (50 µM each); $R_2$ measured by fitting absolute integrals to exponential decay functions of the total spin echo time (constant frequency) with five timepoints per fit; individual points shown (n=3); $R_{2,free}$ values previously measured (Supplementary Fig 2) marked as dotted lines. **e**, effect of SIIP inhibitors on the transverse relaxation rates of probes **9** and **12** in mixture (50 µM each) in the presence of both nucleotide states of KRas$^{G12D}$ (6 µM each, GDP- and GNP-bound); individual points shown (n=2); the lines defined by $R_{2,free}$ and constants C and D in equations 5 and 6 are plotted as solid lines.

### *Methods*

#### *Data analysis*

All non-linear regressions were performed in Graphpad Prism v9 with data points handled as individual values. Constrained constants are noted.

Integrals (I) from single-point CPMG experiments were calculated by dividing the probe's absolute integral by the standard's absolute integral (3-trifluoromethylphenol, 3TFMP) from the same sample. The resulting integral values (I) were averaged (n=3) and then normalized by dividing by the measurement obtained in the absence of protein ($I_0$, n=1 or 3). Errors in $I/I_0$ and $-\ln(I/I_0)$ were propagated from the standard deviations (SD) in I and $I_0$. In all cases, I and $I_0$ were measured from the same premixed aliquot of probe and 3-trifluoromethylphenol.

The $K_D$ values of probes were calculated by fitting normalized integrals to equation 1, where I and $I_0$ are the integrals in the presence and absence of protein, respectively, A is a constant in Hz, $t$ is the total spin-echo time ($2\cdot\tau\cdot n$), and $[P]_0$ and $[L]_0$ are the total concentrations of protein and probe, respectively (constraining $[P]_0$ and $t$). See the supplementary information for derivation.

$$I/I_0 = e^{-A\cdot t\cdot[P]_0/(K_D+[L]_0)} \quad (eq\ 1)$$

Transverse relaxation rate constants ($R_2$) were calculated from multi-point (constant frequency) CPMG experiments; the absolute integrals were fit to exponential decay functions (constraining plateau=0); the decay curve from each replicate sample was fit independently.

The fraction occupancy (F) of $KRas^{G12D}$ GDP was calculated by fitting normalized integrals to equation 2, where I is the integral relative to the internal standard (individual values n=3), $I_{0,0}$ is the measurement obtained in the absence of protein and competitor (mean n=3) and $I_{0,2}$ is the measurement obtained in the presence of protein (2 µM) and absence of competitor (mean n=3).

$$F = 1 - \ln\left(I/I_{0,0}\right) \div \ln\left(I_{0,2}/I_{0,0}\right) \quad (eq\ 2),$$

The $K_{i,obs}$ of MRTX849 was extracted by fitting the resulting fraction occupancy data to equation 3, where B is an unconstrained constant (expect 1 for complete competition), and corrected according to equation 4.

$$F = \frac{B\cdot[MRTX849]}{K_{i,obs}+[MRTX849]} \quad (eq\ 3)$$

$$K_i = \frac{K_{i,obs}}{1+[\mathbf{9}]/K_{D,\mathbf{9}}} \quad (eq\ 4).$$

The concentrations of $KRas^{G12D}$ GDP and $KRas^{G12D}$ GNP in mixed samples were calculated by solving the system of equations 5 and 6. $R_{2,free}$ values were previously measured from samples containing 100 µM of each probe individually (mean n=3). The constants C and D were calculated with $R_2$ measurements (mean n=3) from samples containing either nucleotide state (6 µM) and the mixture of two probes (50 µM each).

$$R_{2,\mathbf{9}} = C_{\mathbf{9}}[GDP] + D_{\mathbf{9}}[GNP] + R_{2,\mathbf{9},free} \quad (eq\ 5)$$

$$R_{2,\mathbf{12}} = C_{\mathbf{12}}[GDP] + D_{\mathbf{12}}[GNP] + R_{2,\mathbf{12},free} \quad (eq\ 6)$$

*Preparation of Ras proteins.*

The plasmids for bacterial expression of HRas 1-166 (His-TEV-N; pProEx; ampicillin resistance) and KRas 1-169 (His-TEV-N; pJ411; kanamycin resistance) have been previously published.[1] Mutants were obtained by site-directed mutagenesis (Q5, NEB; or Genscript Biotech). BL21(DE3) competent cells were transformed with 1-2 ng of plasmid. Cultures for the preparation of natural abundance isotope proteins were grown at 37 °C (A600 0.4-0.6) in Terrific Broth (Sigma Aldrich), induced (1 mM IPTG), and allowed to express protein overnight at 18 °C. Cultures for the preparation of U-$^{15}$N proteins were grown and induced in M9 minimal media supplemented with 1 g/L $^{15}NH_4Cl$ (CIL). Cells were collected by centrifugation, lysed by sonication (Tris buffer pH 8.0, supplemented with 1 mM PMSF or 1 tablet per 50 ml EDTA-free cOmplete, Roche), and cleared by centrifugation. His-tagged Ras proteins were isolated with Co TALON resin (Takara Bio), the nucleotide was exchanged to GDP by dialysis (0.5 mM EDTA), the His tag was cleaved with TEV protease, and the protein was purified by anion exchange (Tris buffer pH 8.0, 50-500 NaCl gradient, HiTrap Q HP, Cytiva) and gel filtration (HEPES storage buffer pH 7.5, Superdex 75, GE or Cytiva). Nucleotide exchange from GDP to GPPNHP (Jena Biosciences) was performed with samples of protein reserved prior to the final gel filtration purification and according to a published procedure comprising EDTA-mediated exchange, desalting, and cleavage of residual GDP with an alkaline phosphatase (CIP or Quick CIP, NEB)[1-4], and then purified by gel filtration. Proteins were concentrated (10k Amicon Ultra, Millipore) to 0.5-1 mM, concentrations were determined by uv (280 nm, $\varepsilon$ = 13410 $M^{-1}cm^{-1}$ for HRas 1-166 and 11920 $M^{-1}cm^{-1}$ for KRas 1-169), and aliquots were flash frozen in liquid $N_2$ and stored at -80 °C.

Storage buffer: 20 mM HEPES, 150 mM NaCl, 1 mM $MgCl_2$. Titrated to pH 7.5 with NaOH. Storage buffer for U-$^{15}$N proteins: 40 mM HEPES, 150 mM NaCl, 4 mM $MgCl_2$, 5% glycerol, 7% $D_2O$. Titrated to pH 7.4 with NaOH.

*$^{1}$H-$^{15}$N HSQC NMR sample preparation and acquisition.*

General procedure: a 0.030 µmol aliquot of U-$^{15}$N protein in storage buffer was diluted to 255 µl with HSQC NMR sample buffer on ice. DSS (15 µl, 20 mM in buffer) and the small molecule ligand were added (30 µl, 10 mM in dmso-*d*$_6$), the sample was mixed by vortex, and the resulting solution was transferred to a 5 mm $D_2O$-matched Shigemi NMR tube (BMS-3). The final concentrations of protein and ligand were 0.10 and 1.0 mM, respectively. To measure the $K_D$ of probe **9**, a 600 µl sample with 50 µM KRas$^{G12D}$(GDP) and a 300 µl sample with 50 µM KRas$^{G12D}$(GDP) and 1.0 mM **9** were prepared; intermediate concentrations of **9** were obtained by mixing these two samples.

1D $^{1}$H (ZGESGP) and 2D $^{1}$H-$^{15}$N fast HSQC (FHSQCCF3GPPH, NS = 8, TDF1 = 256, GARP decoupling) spectra were recorded at 298 K on an 800 MHz Bruker Avance NEO spectrometer equipped with an actively shielded Z-gradient 5 mm TCI cryoprobe (H&F/C/N) using programs from the pulse program library (TopSpin 4.0.8). Spectra were processed and analyzed with Bruker Topspin 4.0 and CCPNMR Analysis v3.[5] $^{1}$H chemical shifts were referenced to 1 mM DSS at 0 ppm; $^{15}$N chemical shifts were referenced indirectly with $\Xi$ = 0.101329118. Backbone assignments were imported from the BMRB (entry 27720 for WT and entry 27719 for G12D).[6]

HSQC NMR buffer: 40 mM HEPES, 150 mM NaCl, 4 mM $MgCl_2$, 7% $D_2O$. Titrated to pH 7.4 with NaOH.

*$^{19}$F CPMG NMR sample preparation and acquisition.*

An aliquot of protein in storage buffer was diluted to 480 µl with sample buffer, then allowed to warm to ambient temperature. x µl of a dmso-*d*$_6$ solution containing the competitor molecule, y µl of a dmso-*d*$_6$ solution of the probe and 3-trifluoromethylphenol (premixed, 10 mM each), and 20-x-y µl of dmso-*d*$_6$ were added (4% dmso-*d*$_6$ final). The sample was mixed and transferred to a 5 mm NMR tube. For the dual-probe measurements, a dmso-*d*$_6$ stock containing both probes and 3-trifluoromethylphenol (premixed, 5.0 mM each) was used. The number of replicates (n) reflects independently prepared protein-containing samples. Blank samples (no protein) were prepared in triplicate for probes **9**, **12**, and **17** and in singlicate for the remaining probes.

$^{19}$F CPMG (CPMG1D) spectra were recorded at 298 K on a 600 MHz Bruker Avance NEO spectrometer equipped with an actively shielded Z-gradient 5 mm TCI cryoprobe (H&F/C/N) using programs from the pulse program library (TopSpin 4.0.6). Spectra were processed and visualized with MestReNova v14. Single-point CPMG experiments were performed with the following key parameters: DS = 8 or 16, NS = 32 or 64, D1 = 4 s, acquisition time = 1.18 s (fidres=0.85 Hz) D20 ($\tau$) = 20 ms, O1P = -59 ppm, and variable CPMG loop counter L4 ($n$). Spectra were processed with TD points and 2 Hz exponential apodization. Multi-point CPMG experiments to determine $R_2$ were performed as a series of five or six CPMG experiments with parameters described above except with D20 ($\tau$) = 10 ms and L4 ($n$) an even number varied from 2 to 40.

Sample buffer: 20 mM HEPES, 150 mM NaCl, 1 mM $MgCl_2$, 7% $D_2O$. Titrated to pH 7.5 with NaOH.

*Derivation of equation 1.*

The dependency of the observed $R_2$ on protein and probe concentrations has been previously published:

$$R_2 = \frac{[PL]}{[L]_0} R_{2,bound} + \left(1 - \frac{[PL]}{[L]_0}\right) R_{2,free} + \frac{[PL]}{[L]_0}\left(1 - \frac{[PL]}{[L]_0}\right)^2 \frac{4\pi^2(\delta_{free} - \delta_{bound})^2}{k_{off}} \qquad (\textit{eq 7}),$$

where $[L]_0$, $[P]_0$, and [PL] are the concentrations of total protein, total probe, and bound probe, respectively; $R_{2,bound}$ and $R_{2,free}$ are the transverse relaxation rate constants of bound and free probe, respectively; $\delta_{free}$-$\delta_{bound}$ is the difference in chemical shift; and $k_{off}$ is the probe dissociation rate constant.[7] When the occupancy of the ligand is low (i.e. $[L]_0 >> [P]_0$ and $[L] \approx [L]_0$), the dependency of $R_2$ on the bound protein can be approximated as:

$$R_2 \approx R_{2,free} + A\, {}^{[PL]}\!/_{[L]_0} \qquad (\textit{eq 8}),$$

where A is a constant in Hz representing the contributions of the bound ligand and exchange process to $R_2$. Therefore, the probe's measured integral

$$I = C \cdot e^{-R_2 \cdot t} = C \cdot e^{-(R_{2,free} + A[PL]/[L]_0)t} \qquad (\textit{eq 9}),$$

where C is a unitless constant, and $t$ is the total spin echo time. $I_0 = I$ when $[PL] = 0$; therefore,

$$I = I_0 \cdot e^{-A \cdot t \cdot [PL]/[L]_0} \qquad (\textit{eq 10}),$$

and the normalized integral

$$I/I_0 = e^{-A \cdot t \cdot [PL]/[L]_0} \qquad (\textit{eq 11}).$$

The dissociation constant

$$K_D = \frac{[P][L]}{[PL]} \approx \frac{([P]_0 - [PL])[L]_0}{[PL]} = \frac{[P]_0[L]_0}{[PL]} - [L]_0 \quad (\textit{eq 12});$$

therefore,

$$I/I_0 = e^{-A \cdot t \cdot [P]_0/(K_D + [L]_0)} \quad (\textit{eq 1}).$$

### *Chemical synthesis*

*General experimental details.* Solvents referred to as "dry" were either purchased (Acros extra dry) or dried with a solvent system containing activated molecular sieves. Other solvents were purchased from Fisher Scientific or Sigma Aldrich and used as received (ACS reagent or HPLC grade). Thin layer chromatography (TLC) was performed with indicator-containing glass-backed silica plates (F254, EMD) and visualized under a 254 nm uv lamp. Liquid chromatography mass spectroscopy (LC/MS) was performed with a Waters Acquity UPLC and Xevo QTOF. Normal-phase flash column chromatography was performed on a Teledyne Isco Combiflash Rf+ system. Preparative reverse-phase high pressure liquid chromatography (HPLC) was performed on either a Waters Autopurification or Teledyne Isco EZPREP system equipped with a 20x300 C18 column and uv/vis detector. NMR spectra were acquired on a 400 MHz Bruker Avance III HD or 600 MHz Avance NEO spectrometer. $^{1}$H and $^{13}$C chemical shifts were referenced to the solvent peak ($CHCl_3$ $^{1}$H δ 7.26 ppm, $CDCl_3$ $^{13}$C δ 77.16, DMSO-$d_5$ $^{1}$H δ 2.50 ppm, DMSO-$d_6$ $^{13}$C δ 39.52). $^{19}$F NMR chemical shifts were referenced indirectly with Ξ = 0.94094011.

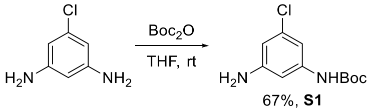

***tert*-Butyl (3-amino-5-chlorophenyl)carbamate (S1).** 5-Chloro-1,3-benzenediamine (714 mg, 5.01 mmol) was dissolved in dry THF (8 ml) in a 50 ml round bottom flask under Ar. Di-*tert*-butyl decarbonate (1.09 g, 5.01, 1.0 equiv mmol) was dissolved in dry THF (4.5 ml), and this solution was added portionwise to the reaction mixture with stirring over 20 min. The reaction mixture was stirred at ambient temperature overnight. Volatiles were removed at reduced pressure, and the crude product was purified by flash column chromatography (hexanes/ethyl acetate gradient) to give the title compound as a white solid (815 mg, 3.36 mmol, 67% yield).
$^{1}$H NMR (400 MHz, DMSO) δ 9.20 (s, 1H), 6.69 (t, *J* = 1.9 Hz, 1H), 6.64 (t, *J* = 1.9 Hz, 1H), 6.19 (t, *J* = 2.0 Hz, 1H), 5.33 (s, 2H), 1.45 (s, 9H).

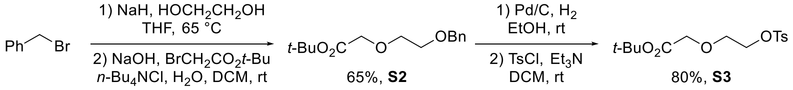

***tert*-Butyl 2-(2-(benzyloxy)ethoxy)acetate (S2).**
Step 1. Sodium hydride (60% w/w oil dispersion, 3.3 g, 83 mmol, 1.1 equiv) was suspended in dry THF (150 ml) in an oven-dried 250 ml round bottom flask under Ar. Ethylene glycol (25 ml, 0.45 mol, 5.9 equiv) was added slowly with stirring while venting the evolved gas through a needle with a gentle flow of Ar, and the mixture was stirred at ambient temperature for 1 h. Benzyl bromide (9.0 ml, 76 mmol) was then added with stirring, a reflux condenser was attached, and the reaction mixture was heated at reflux overnight. The reaction mixture was cooled in an ice bath and quenched by slowly adding saturated $NH_4Cl_{(aq)}$. The mixture was transferred to a separatory funnel and washed twice with sat. $NH_4Cl_{(aq)}$ (ca. 75 ml) and once with brine (ca. 75 ml). The aqueous layers were extracted further with EA in the same sequence (ca. 100 ml). The organic layers were dried over $MgSO_4$ and filtered. Volatiles were removed at reduced pressure to give crude 2-benzyloxyethanol as a yellow oil, which was carried on to the next step without purification.

Step 2. Crude 2-benzyloxyethanol was dissolved in DCM (150 ml) in a 500 ml round bottom flask. The solution was cooled in an ice bath and *tert*-butyl bromoacetate (17 ml, 0.12 mol, 1.5 equiv), *n*-$Bu_4NCl$ (20.8 g, 74.8 mmol, 1.0 equiv), and $NaOH_{(aq)}$ (150 ml, 37% w/w) were added in sequence with stirring. The resulting biphasic mixture was stirred (maximum speed) at 0 °C. The reaction was monitored by TLC from the organic layer (top). When TLC indicated high conversion of the starting material (4 h), the mixture was transferred to a separatory funnel, and the layers were separated. The organic layer was washed with brine (ca. 100 ml). The aqueous layers were extracted further with DCM (ca. 100 ml) in the same sequence. The organic layers were dried over $Na_2SO_4$ and filtered, and volatiles were removed at reduced pressure. The crude product was purified by column chromatography (800 ml silica, isocratic 10% EA/Hex, Rf 0.21) to give the title compound as a clear and colorless oil (12.9 g, 48.4 mmol, 65% yield for 2 steps).

$^{1}H$ NMR (400 MHz, $CDCl_3$) δ 7.38 – 7.26 (m, 5H), 4.58 (s, 2H), 4.04 (s, 2H), 3.78 – 3.71 (m, 2H), 3.71 – 3.63 (m, 2H), 1.47 (s, 9H).

***tert*-Butyl 2-(2-(tosyloxy)ethoxy)acetate (S3).**

Step 1. *Tert*-butyl 2-(2-(benzyloxy)ethoxy)acetate (2.66 g, 9.99 mmol) was dissolved in absolute EtOH (40 ml) in a 100 ml round bottom flask under Ar, and the mixture was cooled to 0 °C in an ice bath. Palladium on carbon (532 mg, 10% w/w, 0.500 mmol, 5.0 mol%) was added to the flask. A balloon of hydrogen was attached, and the atmosphere was exchanged by purging. The cooling bath was removed, and the reaction mixture was stirred at ambient temperature overnight, at which point TLC indicated high conversion of the starting material (product Rf 0.68 in EA, $KMnO_4$ stain). The hydrogen atmosphere was purged with Ar, and the mixture was filtered through a pad of Celite, which was rinsed twice with absolute EtOH (20 ml). Volatiles were removed at reduced pressure to give crude *tert*-butyl 2-(2-hydroxyethoxy)acetate as a light brown oil, which was carried on to the next step without purification.

Step 2. Crude *tert*-butyl 2-(2-hydroxyethoxy)acetate was redissolved in dry DCM (50 ml) in an oven-dried 100 ml round bottom flask under Ar. Triethylamine (2.1 ml, 15 mmol, 1.5 equiv) was added with stirring, and the mixture was cooled to 0 °C in an ice bath. Freshly recrystallized tosyl chloride (2.10 g, 11.0 mmol, 1.1 equiv) was added with stirring, the cooling bath was removed, and the reaction mixture was stirred at ambient temperature overnight. The mixture was transferred to a separatory funnel and washed with 1 M $HCl_{(aq)}$ (60 ml). The acidic aqueous layer was extracted further with 50 ml DCM. The organic layers were dried over $Na_2SO_4$ and filtered, and volatiles were removed at reduced pressure. The crude product was purified by flash column chromatography (hexanes/ethyl acetate gradient, Rf 0.28 in 20% EA/Hex) to give the title compound as a clear and colorless oil (2.63 g, 7.96 mmol, 80% yield for 2 steps).

$^{1}H$ NMR (400 MHz, $CDCl_3$) δ 7.81 (d, J = 8.4 Hz, 2H), 7.34 (d, J = 7.8 Hz, 2H), 4.24 – 4.16 (m, 2H), 3.94 (s, 2H), 3.80 – 3.72 (m, 2H), 2.45 (s, 3H), 1.46 (s, 9H).

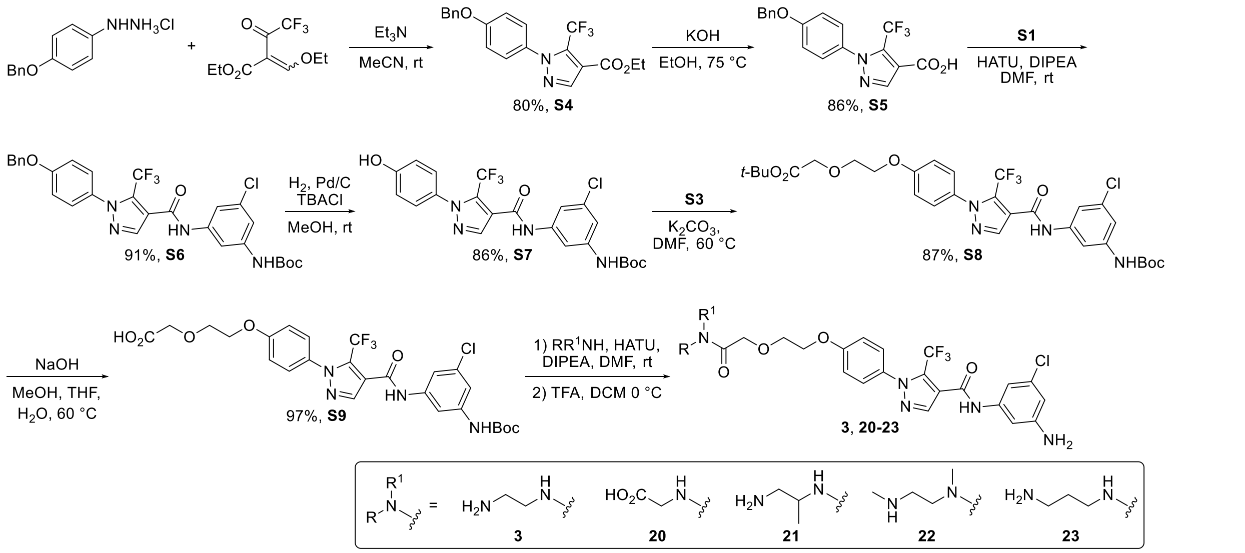

**Ethyl 1-(4-(benzyloxy)phenyl)-5-(trifluoromethyl)-1*H*-pyrazole-4-carboxylate (S4).**

4-(Benzyloxy)phenylhydrazine hydrochloride (2.50 g, 9.97 mmol) was suspended in MeCN (25 ml) in a 250 ml round bottom flask under Ar. Triethylamine (1.4 ml, 1.0 g, 10 mmol, 1.0 equiv) was added with stirring at ambient temperature. The mixture was stirred for ca. 15 minutes, then ethyl-2-(ethoxymethylene)-4,4,4-trifluoro-3-oxo-butanoate (2.0 ml, 2.5 g, 10 mmol, 1.0 equiv) and another portion of triethylamine (1.4 ml, 1.0 g, 10 mmol, 1.0 equiv) were added in sequence. The mixture was stirred at ambient temperature for 1 h. Volatiles were removed at reduced pressure, and the residue was resuspended in DCM (ca. 40 ml). The mixture was washed with $HCl_{(aq)}$ (0.5 M, ca. 40 ml), which was then extracted further with DCM (ca. 40 ml). The organic layers were dried over $Na_2SO_4$ and filtered, and volatiles were removed at reduced pressure. The crude product was purified by flash column

chromatography (hexanes/ethyl acetate gradient, Rf 0.43 in 20% EA/Hex) to give the title compound as an orange solid (3.13 g, 8.01 mmol, 80% yield).
$^{1}$H NMR (400 MHz, $CDCl_3$) δ 8.09 (s, 1H), 7.47 – 7.35 (m, 5H), 7.33 (d, *J* = 8.9 Hz, 2H), 7.06 (d, *J* = 9.0 Hz, 2H), 5.12 (s, 2H), 4.37 (q, *J* = 7.1 Hz, 2H), 1.38 (t, *J* = 7.1 Hz, 3H). $^{19}$F NMR (376 MHz, $CDCl_3$) δ -55.5. ESI$^+$ (LC/MS) calc'd 391.1264, found 391.1296.

**1-(4-(Benzyloxy)phenyl)-5-(trifluoromethyl)-1*H*-pyrazole-4-carboxylic acid (S5).** Ethyl 1-(4-(benzyloxy)yphenyl)-5-(trifluoromethyl)-1*H*-pyrazole-4-carboxylate (1.24 g, 3.18 mmol) was suspended in absolute ethanol (16 ml) in a 20 ml scintillation vial. An aqueous solution of potassium hydroxide (0.48 ml, 40% w/w, 0.67 g, 4.8 mmol, 1.5 equiv) was added with stirring, and the mixture was heated at 75 °C with stirring for 4 h. The mixture was transferred to a separatory funnel and diluted with water and diethyl ether (ca. 60 ml each). The layers were mixed and separated. The aqueous layer was acidified to pH <1 by adding concentrated $HCl_{(aq)}$ and extracted twice with ethyl acetate (ca. 60 ml each). The ethyl acetate layers were dried over $Na_2SO_4$ and filtered. Volatiles were removed at reduced pressure, and the crude product was recrystallized by dissolved in boiling toluene (ca. 15 ml) and then cooling to 0 °C. The solid was collected by filtration and rinsed thrice with cold hexanes (ca. 4 ml each). The filtrate was concentrated, and a second crop was obtained by the same procedure with lower volumes. The two crops were combined to give the title compound as off-white crystals (991 mg, 2.74 mmol, 86% yield).
$^{1}$H NMR (400 MHz, $CDCl_3$) δ 8.17 (s, 1H), 7.48 – 7.36 (m, 5H), 7.35 (d, *J* = 8.7 Hz, 2H), 7.07 (d, *J* = 9.0 Hz, 2H), 5.13 (s, 2H). $^{19}$F NMR (376 MHz, $CDCl_3$) δ -55.6. *Note: the -$CO_2\underline{H}$ resonance was not located in the $^{1}$H NMR spectrum.* ESI$^+$ (LC/MS) calc'd 363.0951, found 363.0989.

***tert*-Butyl (3-(1-(4-(benzyloxy)phenyl)-5-(trifluoromethyl)-1*H*-pyrazole-4-carboxamido)-5-chlorophenyl)carbamate (S6).** 1-(4-(Benzyloxy)phenyl)-5-(trifluoromethyl)-1*H*-pyrazole-4-carboxylic acid (543 mg, 1.50 mmol), *tert*-butyl (3-amino-5-chlorophenyl)carbamate (400 mg, 1.65 mmol, 1.1 equiv), and HATU (855 mg, 2.25 mmol, 1.5 equiv) were added to an oven-dried 1 dram vial under Ar. The mixture was cooled to 0 °C in an ice bath and dissolved in dry DMF (0.8 ml). Di-(*iso*-propyl)ethylamine (0.80 ml, 4.6 mmol, 3.1 equiv) was added slowly with stirring. The reaction mixture was stirred at 0 °C for approximately 10 min, then the cooling bath was removed, and the mixture was stirred at ambient temperature overnight. The mixture was diluted with ethyl acetate and sat. $NH_4C_{(aq)}$ (ca. 40 ml each) and transferred to a separatory funnel. The layers were mixed and separated, and the aqueous layer was extracted further with ethyl acetate (ca. 40 ml). The organic layers were dried over $Na_2SO_4$ and filtered, and volatiles were removed at reduced pressure. The crude product was purified by flash column chromatography (hexanes/ethyl acetate gradient, Rf 0.80 in 50% EA/Hex) to give the title compound as a yellow-brown solid (802 mg, 1.37 mmol, 91% yield).
$^{1}$H NMR (400 MHz, DMSO) δ 10.64 (s, 1H), 9.64 (s, 1H), 8.26 (s, 1H), 7.85 (s, 1H), 7.54 – 7.33 (m, 8H), 7.25 (t, *J* = 1.9 Hz, 1H), 7.20 (d, *J* = 9.0 Hz, 2H), 5.20 (s, 2H), 1.48 (s, 9H). $^{19}$F NMR (376 MHz, DMSO) δ -55.0. ESI$^+$ (LC/MS) calc'd 609.1487, found 609.1503 ($MNa^+$).

***tert*-Butyl (3-chloro-5-(1-(4-hydroxyphenyl)-5-(trifluoromethyl)-1*H*-pyrazole-4-carboxamido)phenyl)carbamate (S7).** *tert*-Butyl (3-(1-(4-(benzyloxy)phenyl)-5-(trifluoromethyl)-1*H*-pyrazole-4-carboxamido)-5-chlorophenyl)carbamate (772 mg, 1.32 mmol) and tetrabutylammonium chloride (402 mg, 1.45 mmol, 1.1 equiv) were suspended in MeOH (13 ml) in a 20 ml scintillation vial. The mixture was cooled to 0 °C in an ice bath, and palladium (10% w/w on carbon, 70 mg, 0.066 mmol, 5.0 mol%) was added. The atmosphere was exchanged to hydrogen by purging with a balloon, and the cooling bath was removed. The mixture was stirred at ambient temperature under an atmosphere of hydrogen (balloon, ca. 1 atm) for 2 h. The atmosphere was purged with Ar, and the mixture was filtered through a plug of Celite, which was rinsed with methanol (ca. 10 ml). Volatiles were removed at reduced pressure, and the crude product was purified by flash column chromatography (hexanes/ethyl acetate gradient, Rf 0.51 in 50% EA/Hex) to give the title compound as a white solid (562 mg, 1.13 mmol, 86% yield).
$^{1}$H NMR (400 MHz, DMSO) δ 10.63 (s, 1H), 10.09 (s, 1H), 9.64 (s, 1H), 8.23 (s, 1H), 7.85 (t, *J* = 1.9 Hz, 1H), 7.51 (t, *J* = 1.9 Hz, 1H), 7.31 (d, *J* = 8.7 Hz, 2H), 7.25 (t, *J* = 1.9 Hz, 1H), 6.91 (d, *J* = 8.8 Hz, 2H), 1.48 (s, 9H). $^{19}$F NMR (376 MHz, DMSO) δ -55.1. ESI$^+$ (LC/MS) calc'd 519.1017, found 519.1024 ($MNa^+$).

***tert*-Butyl 2-(2-(4-(4-((3-((tert-butoxycarbonyl)amino)-5-chlorophenyl)carbamoyl)-5-(trifluoromethyl)-1*H*-pyrazol-1-yl)phenoxy)ethoxy)acetate (S8).** *tert*-Butyl (3-chloro-5-(1-(4-hydroxyphenyl)-5-(trifluoromethyl)-1*H*-pyrazole-4-carboxamido)phenyl)carbamate (497 mg, 1.00 mmol), potassium carbonate (225 mg, 1.63 mmol, 1.6 equiv), and dry DMF (4 ml) were added to an oven-dried 20 ml scintillation vial under Ar. The mixture was heated at 60 °C with stirring for 15 min. Then, *tert*-butyl 2-(2-(tosyloxy)ethoxy)acetate (606 mg, 1.83 mmol, 1.8 equiv) was added with stirring, and the reaction was heated at 60 °C with stirring overnight. The mixture was cooled to ambient temperature, diluted with ethyl acetate and sat. $NH_4Cl_{(aq)}$ (ca. 15 ml each), and transferred to a separatory funnel. The layers were mixed and separated, and the organic layer was washed again with brine (ca. 15 ml). The aqueous layers were extracted further in the same sequence with ethyl acetate (ca. 15 ml). The organic layers were dried over $Na_2SO_4$ and filtered, and volatiles were removed at reduced pressure. The crude product was purified by flash column chromatography (hexanes/ethyl acetate gradient) to give the title compound as a white solid (573 mg, 0.875 mmol, 87% yield).
$^1$H NMR (400 MHz, DMSO) δ 10.64 (s, 1H), 9.64 (s, 1H), 8.26 (s, 1H), 7.85 (t, *J* = 1.9 Hz, 1H), 7.52 (t, *J* = 1.9 Hz, 1H), 7.44 (d, *J* = 8.9 Hz, 2H), 7.25 (t, *J* = 1.9 Hz, 1H), 7.12 (d, *J* = 9.0 Hz, 2H), 4.26 – 4.17 (m, 2H), 4.08 (s, 2H), 3.89 – 3.79 (m, 2H), 1.48 (s, 9H), 1.42 (s, 9H). $^{19}$F NMR (376 MHz, DMSO) δ -55.0. ESI$^+$ (LC/MS) calc'd 677.1960, found 677.1991 (MNa$^+$).

**2-(2-(4-(4-((3-((*tert*-Butoxycarbonyl)amino)-5-chlorophenyl)carbamoyl)-5-(trifluoromethyl)-1*H*-pyrazol-1-yl)phenoxy)ethoxy)acetic acid (S9).** *tert*-Butyl 2-(2-(4-(4-((3-((tert-butoxycarbonyl)amino)-5-chlorophenyl)carbamoyl)-5-(trifluoromethyl)-1*H*-pyrazol-1-yl)phenoxy)ethoxy)acetate (564 mg, 0.861 mmol) was dissolved in a 1:1:1 mixture of THF, methanol, and water (8 ml). An aqueous solution of sodium hydroxide (15% w/w, 0.40 ml, 0.47 g, 1.8 mmol, 2.0 equiv) was added with stirring, and the mixture was heated at 60 °C with stirring for 30 min. The mixture was cooled to ambient temperature and concentrated at reduced pressure. The residue was diluted with water, acidified by adding $HCl_{(aq)}$, and extracted with ethyl acetate. The organic layer was filtered through $Na_2SO_4$, and volatiles were removed at reduced pressure to give the title compound as a white powder (502 mg, 0.838 mmol, 97% yield).
$^1$H NMR (400 MHz, DMSO) δ 12.63 (s, 1H), 10.64 (s, 1H), 9.64 (s, 1H), 8.26 (s, 1H), 7.85 (t, *J* = 1.8 Hz, 1H), 7.52 (t, *J* = 1.9 Hz, 1H), 7.44 (d, *J* = 8.9 Hz, 2H), 7.25 (t, *J* = 2.0 Hz, 1H), 7.13 (d, *J* = 9.0 Hz, 2H), 4.25 – 4.17 (m, 2H), 4.12 (s, 2H), 3.89 – 3.81 (m, 2H), 1.48 (s, 9H). $^{19}$F NMR (376 MHz, DMSO) δ -55.0. ESI$^+$ (LC/MS) calc'd 621.1334, found 621.1324 (MNa$^+$).

**General procedure for the preparation of 3 and 20-23.**
Step 1. 2-(2-(4-(4-((3-((*tert*-Butoxycarbonyl)amino)-5-chlorophenyl)carbamoyl)-5-(trifluoromethyl)-1*H*-pyrazol-1-yl)phenoxy)ethoxy)acetic acid and HATU (1.0 equiv) were added to an oven-dried 1 dram vial under Ar. The mixture was cooled to 0 °C in an ice bath, and DMF (0.10 M), amine nucleophile (2.0 equiv), and DIPEA (1.0 equiv) were added in sequence. After 15 min, the cooling bath was removed, and the mixture was stirred at ambient temperature overnight. The mixture was diluted with ethyl acetate and saturated $NH_4Cl_{(aq)}$ (ca. 2 ml each). The layers were mixed and separated, and the aqueous layer was extracted twice more with ethyl acetate (ca. 2 ml each). The organic layers were filtered through $Na_2SO_4$, and volatiles were removed at reduced pressure. The crude product was purified by flash column chromatography (hexanes/ethyl acetate gradient).

Step 2. The bis-Boc-protected product from step 1 was dissolved (or suspended) in DCM (1.5 ml) in a 20 ml scintillation vial, and the mixture was cooled to 0 °C in an ice bath. TFA (0.5 ml) was added with stirring, and the mixture was stirred at 0 °C until LC/MS analysis indicated complete cleavage of the Boc protecting groups (typically 1 h). Volatiles were evaporated under a stream of Ar, and the crude product was purified by high pressure liquid chromatography (C18, water/acetonitrile gradient with 0.1% formic acid). Acetonitrile was removed at reduced pressure, and the resulting aqueous mixture was frozen and lyophilized to afford the product as a 1:1 formic acid salt.

***N*-(3-amino-5-chlorophenyl)-1-(4-(2-(2-((2-aminoethyl)amino)-2-oxoethoxy)ethoxy)phenyl)-5-(trifluoromethyl)-1*H*-pyrazole-4-carboxamide (3).** Prepared as described above from 2-(2-(4-(4-((3-((*tert*-butoxycarbonyl)amino)-5-chlorophenyl)carbamoyl)-5-(trifluoromethyl)-1*H*-pyrazol-1-yl)phenoxy)ethoxy)acetic acid (30.0 mg, 0.0500 mmol), *tert*-butyl (2-aminoethyl)carbamate (16 µl, 0.10 mmol, 2.0 equiv), HATU (19 mg, 0.050 mmol, 1.0 equiv), and DIPEA (8 µl, 0.05 mmol, 1 equiv) to give the title compound in the form of a 1:1 formic acid salt (10 mg, 0.017 mmol, 34% yield).

$^{1}H$ NMR (400 MHz, DMSO) δ 10.34 (s, 1H), 8.33 (br, 1H), 8.23 (s, 1H), 8.0-6.0 (v. br), 7.90 (br, 1H), 7.45 (d, *J* = 8.9 Hz, 2H), 7.14 (d, *J* = 8.9 Hz, 2H), 6.95 (s, 1H), 6.90 (s, 1H), 6.35 (t, *J* = 1.9 Hz, 1H), 5.50 (br, 2H), 4.30 – 4.22 (m, 2H), 3.90 – 3.81 (m, 2H), 3.26 – 3.19 (m, 2H), 2.78 – 2.69 (m, 2H). $^{19}F$ NMR (376 MHz, DMSO) δ -55.0. ESI$^{+}$ (LC/MS) calc'd 541.1573, found 541.1598. *Note: the -N$\underline{H}_3^{+}$ resonance was not located in the $^{1}H$ NMR spectrum, and the -C$\underline{H}_2$N- resonance at 3.2 ppm is partially obscured by the broad $H_2O$ resonance.*

**(2-(2-(4-(4-((3-Amino-5-chlorophenyl)carbamoyl)-5-(trifluoromethyl)-1*H*-pyrazol-1-yl)phenoxy)ethoxy)acetyl)glycine (20).** Prepared as described above from 2-(2-(4-(4-((3-((*tert*-butoxycarbonyl)amino)-5-chlorophenyl)carbamoyl)-5-(trifluoromethyl)-1*H*-pyrazol-1-yl)phenoxy)ethoxy)acetic acid (30.0 mg, 0.0500 mmol), glycine *tert*-butyl ester hydrochloride (10 mg, 0.060 mmol, 1.2 equiv), HATU (19 mg, 0.050 mmol, 1.0 equiv), and DIPEA (26 µl, 0.15 mmol, 3.0 equiv) to give the title compound (10.4 mg, 0.0190 mmol, 37% yield).
$^{1}H$ NMR (400 MHz, DMSO) δ 12.56 (br, 1H), 10.33 (s, 1H), 8.22 (s, 1H), 8.02 (t, *J* = 5.9 Hz, 1H), 7.44 (d, *J* = 8.9 Hz, 2H), 7.15 (d, *J* = 9.0 Hz, 2H), 6.95 (s, 1H), 6.90 (s, 1H), 6.35 (t, *J* = 2.0 Hz, 1H), 5.50 (br, 2H), 4.32 – 4.19 (m, 2H), 4.02 (s, 2H), 3.92 – 3.83 (m, 2H), 3.79 (d, *J* = 6.0 Hz, 2H). $^{19}F$ NMR (376 MHz, DMSO) δ -55.0. ESI$^{+}$ (LC/MS) calc'd 556.1205, found 556.1196.

**(*rac*)-*N*-(3-Amino-5-chlorophenyl)-1-(4-(2-(2-((1-aminopropan-2-yl)amino)-2-oxoethoxy)ethoxy)phenyl)-5-(trifluoromethyl)-1*H*-pyrazole-4-carboxamide (21).** Prepared as described above from 2-(2-(4-(4-((3-((*tert*-butoxycarbonyl)amino)-5-chlorophenyl)carbamoyl)-5-(trifluoromethyl)-1*H*-pyrazol-1-yl)phenoxy)ethoxy)acetic acid (30.0 mg, 0.0500 mmol), *tert*-butyl *N*-(2-aminopropyl)carbamate (17.5 mg, 0.100 mmol, 2.0 equiv), HATU (19 mg, 0.050 mmol, 1.0 equiv), and DIPEA (9 µl, 0.05 mmol, 1 equiv) to give the title compound in the form of a 1:1 formic acid salt (25 mg, 0.042 mmol, 83% yield). $^{1}H$ NMR (400 MHz, DMSO) δ 10.33 (s, 1H), 8.31 (s, 1H), 8.23 (s, 1H), 7.56 (d, *J* = 8.5 Hz, 1H), 7.45 (d, *J* = 8.9 Hz, 2H), 7.14 (d, *J* = 9.0 Hz, 2H), 6.95 (s, 1H), 6.90 (s, 1H), 6.35 (t, *J* = 1.9 Hz, 1H), 5.50 (s, 2H), 4.25 (t, *J* = 4.4 Hz, 2H), 3.97 (d, *J* = 2.9 Hz, 2H), 3.89 – 3.83 (m, 2H), 2.72 – 2.60 (m, 2H), 1.05 (d, *J* = 6.7 Hz, 3H). $^{19}F$ NMR (376 MHz, DMSO) δ -55.0. ESI$^{+}$ (LC/MS) calc'd 555.1729, found 555.1738.

***N*-(3-amino-5-chlorophenyl)-1-(4-(2-(2-(methyl(2-(methylamino)ethyl)amino)-2-oxoethoxy)ethoxy)phenyl)-5-(trifluoromethyl)-1*H*-pyrazole-4-carboxamide (22).** Prepared as described above from 2-(2-(4-(4-((3-((*tert*-butoxycarbonyl)amino)-5-chlorophenyl)carbamoyl)-5-(trifluoromethyl)-1*H*-pyrazol-1-yl)phenoxy)ethoxy)acetic acid (30.0 mg, 0.0500 mmol), *tert*-butyl methyl(2-(methylamino)ethyl)carbamate hydrochloride (13.5 mg, 0.060 mmol, 1.2 equiv), HATU (19 mg, 0.050 mmol, 1.0 equiv), and DIPEA (26 µl, 0.15 mmol, 3.0 equiv) to give the title compound in the form of a 1:1 formic acid salt (13.5 mg, 0.022 mmol, 44% yield).
$^{1}H$ NMR (400 MHz, DMSO) δ 10.33 (s, 1H), 8.25 (br, 1H), 8.23 (s, 1H), 7.44 (d, *J* = 8.9 Hz, 2H), 7.13 (d, *J* = 9.0 Hz, 2H), 6.95 (s, 1H), 6.90 (s, 1H), 6.35 (t, *J* = 2.0 Hz, 1H), 5.50 (br, 2H), 4.30 – 4.17 (m, 4H), 3.87 – 3.78 (m, 2H), 3.41 (t, *J* = 6.0 Hz, *major* 2H), 3.36 – 3.30 (m, *minor* 2H), 2.92 (s, *major* 3H), 2.80 (s, *minor* 3H), 2.76 (t, *J* = 6.3 Hz, *major* 2H), 2.72 – 2.64 (m, *minor* 1H), 2.38 (s, *major* 3H), 2.31 (s, *minor* 3H). $^{19}F$ NMR (376 MHz, DMSO) δ -55.0. ESI$^{+}$ (LC/MS) calc'd 569.1886, found 569.1863.

***N*-(3-amino-5-chlorophenyl)-1-(4-(2-(2-((3-aminopropyl)amino)-2-oxoethoxy)ethoxy)phenyl)-5-(trifluoromethyl)-1*H*-pyrazole-4-carboxamide (23).** Prepared as described above from 2-(2-(4-(4-((3-((*tert*-butoxycarbonyl)amino)-5-chlorophenyl)carbamoyl)-5-(trifluoromethyl)-1*H*-pyrazol-1-yl)phenoxy)ethoxy)acetic acid (30.0 mg, 0.0500 mmol), *tert*-butyl (3-aminopropyl)carbamate (17.5 mg, 0.10 mmol, 2.0 equiv), HATU (19 mg, 0.050 mmol, 1.0 equiv), and DIPEA (9 µl, 0.05 mmol, 1 equiv) to give the title compound in the form of a 1:1 formic acid salt (25.4 mg, 0.046 mmol, 91% yield). $^{1}H$ NMR (400 MHz, DMSO) δ 10.34 (s, 1H), 8.38 (s, 1H), 8.23 (s, 1H), 7.91 (t, *J* = 5.9 Hz, 1H), 7.45 (d, *J* = 8.9 Hz, 2H), 7.14 (d, *J* = 9.0 Hz, 2H), 6.95 (s, 1H), 6.90 (s, 1H), 6.35 (t, *J* = 2.0 Hz, 1H), 5.50 (br, 2H), 4.29 – 4.21 (m, 2H), 3.97 (s, 2H), 3.89 – 3.80 (m, 2H), 3.17 (q, *J* = 6.5 Hz, 2H), 2.68 (t, *J* = 7.1 Hz, 2H), 1.61 (p, *J* = 6.8 Hz, 2H). $^{19}F$ NMR (376 MHz, DMSO) δ -55.0. ESI$^{+}$ (LC/MS) calc'd 555.1729, found 555.1738.

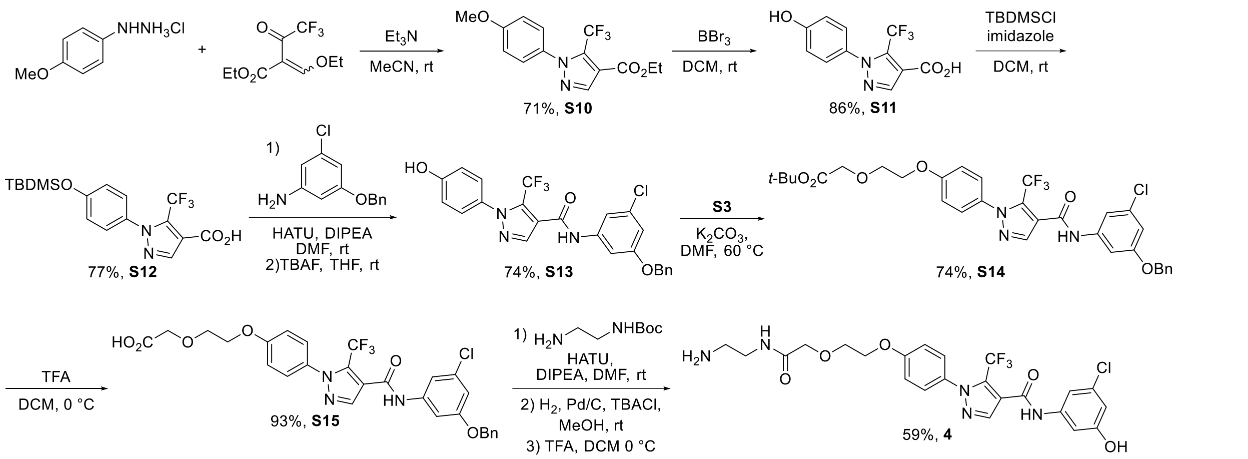

**Ethyl 1-(4-methoxyphenyl)-5-(trifluoromethyl)-1*H*-pyrazole-4-carboxylate (S10)** was prepared as reported in the literature.[8]

**1-(4-Hydroxyphenyl)-5-(trifluoromethyl)-1*H*-pyrazole-4-carboxylic acid (S11).** Ethyl 1-(4-methoxyphenyl)-5-(trifluoromethyl)-1*H*-pyrazole-4-carboxylate (716 mg, 2.50 mmol) was dissolved in dry DCM (50 ml) in an oven-dried 250 ml round bottom flask under Ar, and the solution was cooled to 0 °C in an ice bath. Boron tribromide (2.4 ml. 25 mmol, 10 equiv) was added to the reaction flask slowly with stirring at 0 °C. The reaction mixture was stirred overnight, allowing the ice bath to slowly thaw and warm to ambient temperature. The reaction mixture was again cooled to 0 °C and slowly transferred via cannula into a flask containing water (25 ml) at 0 °C. The resulting white suspension was stirred at 0 °C for 30 min, then the bath was removed, and the mixture was stirred at ambient temperature for 1 h. DCM was removed at reduced pressure, and the aqueous mixture was diluted to ca. 60 ml with water and extracted thrice with ethyl acetate (60/25/25 ml). The organic layers were washed with brine (25 ml) in the same sequence and dried over $Na_2SO_4$. Volatiles were removed at reduced pressure, and the crude product was purified by flash column chromatography (hexanes/ethyl acetate gradient) to give the title compound as an off-white solid (587 mg, 2.16, 86% yield).
$^1$H NMR (400 MHz, DMSO) δ 13.26 (s, 1H), 10.06 (s, 1H), 8.17 (s, 1H), 7.30 (d, $J$ = 8.8 Hz, 2H), 6.88 (d, $J$ = 8.8 Hz, 2H). $^{19}$F NMR (376 MHz, DMSO) δ -54.6. ESI$^-$ (LC/MS) calc'd 271.0336, found 271.0360 (M-H$^+$).

**1-(4-((*tert*-Butyldimethylsilyl)oxy)phenyl)-5-(trifluoromethyl)-1*H*-pyrazole-4-carboxylic acid (S12).** 1-(4-Hydroxyphenyl)-5-(trifluoromethyl)-1*H*-pyrazole-4-carboxylic acid (413 mg, 1.52 mmol) was dissolved in dry DCM (10.5 ml) in a scintillation vial under Ar. Imidazole (310 mg, 4.55 mmol, 3.0 equiv) and *tert*-butyldimethylchlorosilane (343 mg, 2.28 mmol, 1.5 equiv) were added in that order with stirring. The reaction mixture was stirred at ambient temperature for 1 h, then additional portions of imidazole (155 mg, 2.28 mmol, 1.5 equiv) and *tert*-butyldimethylchlorosilane (343 mg, 2.28 mmol, 1.5 equiv) were added, and then mixture was stirred at ambient temperature overnight (ca. 20 h). The solution was washed with aqueous citric acid (5% w/w, 10 ml), and the aqueous layer was extracted further with DCM (ca. 10 ml). The organic layers were dried over $Na_2SO_4$, and volatiles were removed at reduced pressure to give the doubly protected crude product *tert*-butyldimethylsilyl 1-(4-((tert-butyldimethylsilyl)oxy)phenyl)-5-(trifluoromethyl)-1*H*-pyrazole-4-carboxylate as a white solid. This sample was dissolved in an acetic acid/methanol/DCM mixture (0.2:2:98 v/v, 4 ml) and stirred with silica (2 ml) at ambient temperature until TLC indicated high conversion to a lower Rf spot (Rf irreproducible, streaks). This sample was then purified by flash column chromatography (hexanes/ethyl acetate gradient); fractions containing the doubly-protected intermediate were combined and resubjected to the same cleavage procedure. Fractions containing the singly protected product were combined, and volatiles were removed at reduced pressure to give the title compound as a white solid (452 mg, 1.17 mmol, 77% yield).
$^1$H NMR (400 MHz, $CDCl_3$) δ 8.17 (s, 1H), 7.29 (d, $J$ = 8.8 Hz, 2H), 6.93 (d, $J$ = 8.8 Hz, 2H), 1.00 (s, 9H), 0.24 (s, 6H). $^{19}$F NMR (376 MHz, $CDCl_3$) δ -55.6. ESI$^+$ (LC/MS) calc'd 387.1347, found 387.1334. *Note: The -COOH resonance was not located in the $^1$H NMR spectrum, likely due to fast exchange.*

***N*-(3-(Benzyloxy)-5-chlorophenyl)-1-(4-hydroxyphenyl)-5-(trifluoromethyl)-1*H*-pyrazole-4-carboxamide (S13).**

Step1. An oven-dried 1-dram vial under Ar was charged with 1-(4-((*tert*-butyldimethylsilyl)oxy)phenyl)-5-(trifluoromethyl)-1*H*-pyrazole-4-carboxylic acid (77.2 mg, 0.200 mmol), 3-benzyloxy-5-chloro-aniline (51.3 mg, 0.220 mmol, 1.1 equiv), and HATU (114 mg, 0.300 mmol, 1.5 equiv). The mixture was cooled to 0 °C in an ice bath, and DMF (0.80 ml) and DIPEA (0.11 ml, 0.63 mmol, 3.2 equiv) were added in that order with stirring. The reaction mixture was stirred at 0 °C for approximately 10 min, then the cooling bath was removed, and the mixture was stirred at ambient temperature overnight. The reaction mixture was diluted with $HCl_{(aq)}$ (1 M, 6 ml) and extracted thrice with ethyl acetate (4 ml each). The organic layers were filtered through $Na_2SO_4$, and volatiles were removed at reduced pressure. The crude product was carried on to the next step without purification.

Step 2. The reaction residue from step 1 was redissolved in dry THF (1 ml), and the solution was cooled to 0 °C in an ice bath. TBAF (1 M in THF, 0.30 ml, 1.5 equiv) was added with stirring, and the reaction mixture was stirred at 0 °C for 30 min, at which point TLC indicated high conversion of the TDMS ether. Saturated $NH_4Cl_{(aq)}$ (5 ml) was added, and the mixture was extracted thrice with ethyl acetate (4 ml each). The organic layers were filtered through $Na_2SO_4$, and volatiles were removed at reduced pressure. The crude product was purified by flash column chromatography (hexanes/ethyl acetate gradient, Rf 0.44 in 50% EA/Hex) to give the title compound (72.0 mg, 0.148 mmol, 74% yield).

$^1$H NMR (400 MHz, DMSO) δ 10.62 (s, 1H), 10.09 (s, 1H), 8.23 (s, 1H), 7.49 – 7.27 (m, 9H), 6.95 – 6.87 (m, 3H), 5.13 (s, 2H). $^{19}$F NMR (376 MHz, DMSO) δ -55.1. ESI$^+$ (LC/MS) calc'd 488.0984, found 488.0973.

***tert*-Butyl 2-(2-(4-(4-((3-(benzyloxy)-5-chlorophenyl)carbamoyl)-5-(trifluoromethyl)-1*H*-pyrazol-1-yl)phenoxy)ethoxy)acetate (S14).** A 20 ml scintillation vial under Ar was charged with *N*-(3-(benzyloxy)-5-chlorophenyl)-1-(4-hydroxyphenyl)-5-(trifluoromethyl)-1*H*-pyrazole-4-carboxamide (71.0 mg, 0.146 mmol), *tert*-Butyl 2-(2-(tosyloxy)ethoxy)acetate (92 mg, 0.28 mmol, 1.9 equiv), potassium carbonate (ca. 33 mg, 0.24 mmol, 1.6 equiv), and DMF (0.58 ml). The reaction mixture was heated at 60 °C with stirring overnight. The reaction mixture was diluted with saturated $NH_4Cl_{(aq)}$ (4 ml) and extracted five times with ethyl acetate (4/2/2/2/2 ml). The organic layers were filtered through $Na_2SO_4$, and volatiles were removed at reduced pressure. The crude product was purified by flash column chromatography (hexanes/ethyl acetate gradient, Rf 0.61 in 50% EA/Hex) to give the title compound as a viscous yellow oil (69.7 mg, 0.108 mmol, 74% yield).

$^1$H NMR (400 MHz, DMSO) δ 10.63 (s, 1H), 8.26 (s, 1H), 7.50 – 7.32 (m, 9H), 7.12 (d, *J* = 9.0 Hz, 2H), 6.90 (t, *J* = 2.1 Hz, 1H), 5.13 (s, 2H), 4.24 – 4.18 (m, 2H), 4.08 (s, 2H), 3.87 – 3.81 (m, 2H), 1.42 (s, 9H). $^{19}$F NMR (376 MHz, DMSO) δ -55.0. ESI$^+$ (LC/MS) calc'd 646.1926, found 646.1959.

**2-(2-(4-(4-((3-(Benzyloxy)-5-chlorophenyl)carbamoyl)-5-(trifluoromethyl)-1*H*-pyrazol-1-yl)phenoxy)ethoxy)acetic acid (S15).** *tert*-Butyl 2-(2-(4-(4-((3-(benzyloxy)-5-chlorophenyl)carbamoyl)-5-(trifluoromethyl)-1*H*-pyrazol-1-yl)phenoxy)ethoxy)acetate (68.7 mg, 0.106 mmol) was dissolved in DCM (1.5 ml) in a 20 ml scintillation vial under Ar, and the solution was cooled to 0 °C in an ice bath. TFA (0.5 ml) was added slowly with stirring, and the reaction mixture was stirred at 0 °C for 2 h. Volatiles were evaporated under a stream of Ar, and the crude product was purified by flash column chromatography (0 to 10% MeOH gradient in DCM with 0.2% acetic acid, Rf 0.28 in 0.2/10/90 AcOH/MeOH/DCM) to give the title compound as an off-white solid (58.4 mg, 0.0990 mmol, 93% yield).

$^1$H NMR (400 MHz, DMSO) δ 12.61 (br, 1H), 10.63 (s, 1H), 8.26 (s, 1H), 7.50 – 7.32 (m, 9H), 7.13 (d, *J* = 9.0 Hz, 2H), 6.90 (t, *J* = 2.1 Hz, 1H), 5.13 (s, 2H), 4.24 – 4.18 (m, 2H), 4.12 (s, 2H), 3.90 – 3.81 (m, 2H). $^{19}$F NMR (376 MHz, DMSO) δ -55.0. ESI$^+$ (LC/MS) calc'd 590.1300, found 590.1321.

***N*-(3-Chloro-5-hydroxyphenyl)-1-(4-(2-(2-((2-aminoethyl)amino)-2-oxoethoxy)ethoxy)phenyl)-5-(trifluoromethyl)-1*H*-pyrazole-4-carboxamide (4).**

Step 1. An oven-dried 1-dram vial was charged with 2-(2-(4-(4-((3-(benzyloxy)-5-chlorophenyl)carbamoyl)-5-(trifluoromethyl)-1*H*-pyrazol-1-yl)phenoxy)ethoxy)acetic acid (29.5 mg, 0.0500 mmol) and HATU (28.5 mg, 0.0750, 1.5 equiv) and flushed with Ar. The mixture was cooled in an ice bath and DMF (0.5 ml), *tert*-butyl (2-aminoethyl)carbamate (16 µl, 0.10 mmol, 2.0 equiv), and DIPEA (17 µl, 0.10 mmol, 2.0 equiv) were added in that order with stirring. The reaction mixture was stirred for 15 min, then the cooling bath was removed, and the mixture was stirred at ambient temperature overnight. The mixture was diluted with EA (2 ml) and washed with 0.5 M

$HCl_{(aq)}$ (2 ml) and brine (2 ml). The aqueous layers were extracted twice more with EA (2 ml each) in the same sequence, and the organic layers were filtered through $Na_2SO_4$. Volatiles were removed at reduced pressure, and the crude product was purified by flash column chromatography (2 to 10% methanol gradient in DCM, Rf 0.47 in 10% MeOH/DCM).

Step 2. The sample obtained from step 1 was dissolved in MeOH (0.52 ml) in a scintillation vial. Tetrabutylammonium chloride (16 mg, 0.058 mmol, 1.1 equiv) was added, the vial was flushed with Ar, and the mixture was cooled to 0 °C in an ice bath. Palladium on carbon (10% w/w, 3 mg, 0.003 mmol, 5 mol%) was added with stirring. A balloon of hydrogen was attached, and the atmosphere was exchanged by purging. The reaction mixture was stirred at 0 °C for 15 min, then the cooling bath was removed, and the mixture was stirred at ambient temperature until LC/MS analysis indicated complete cleavage of the benzyl ether (5 h). The atmosphere was purged with Ar, and the mixture was filtered through a plug of Celite, which was rinsed thrice with MeOH (1 ml each). Volatiles were removed at reduced pressure.

Step 3. The sample obtained from step 2 was dissolved in DCM (1.5 ml) in a scintillation vial, and the solution was cooled to 0 °C in an ice bath. TFA (0.5 ml) was added slowly with stirring, and the mixture was stirred at 0 °C for 1 h. Volatiles were evaporated under a stream of Ar. The crude product was purified by preparative reverse-phase HPLC (C18, water/acetonitrile gradient + 0.1% formic acid). Pure fractions were combined, and MeCN was removed at reduced pressure. The resulting aqueous mixture was frozen and lyophilized to give the title compound and a flocculent white solid (17.4 mg, 0.0296 mmol, 56% yield).
$^1$H NMR (400 MHz, DMSO) δ 10.53 (s, 1H), 8.32 (s, 1H), 8.26 (s, 1H), 7.86 (t, $J$ = 5.8 Hz, 1H), 7.45 (d, $J$ = 8.9 Hz, 2H), 7.26 (t, $J$ = 1.9 Hz, 1H), 7.19 (t, $J$ = 2.0 Hz, 1H), 7.14 (d, $J$ = 9.0 Hz, 2H), 6.56 (t, $J$ = 2.1 Hz, 1H), 4.29 – 4.22 (m, 2H), 3.98 (s, 2H), 3.90 – 3.81 (m, 2H), 3.21 (q, $J$ = 6.2 Hz, 2H), 2.72 (t, $J$ = 6.4 Hz, 2H). $^{19}$F NMR (376 MHz, DMSO) δ -55.0. ESI$^+$ (LC/MS) calc'd 542.1413, found 542.1412. *Note: the -N$\underline{H}_3^+$ and -O$\underline{H}$ resonances were not located in the $^1$H NMR spectrum, and the -C$\underline{H}_2$N- resonance at 3.2 ppm is partially obscured by the broad $H_2O$ resonance.*

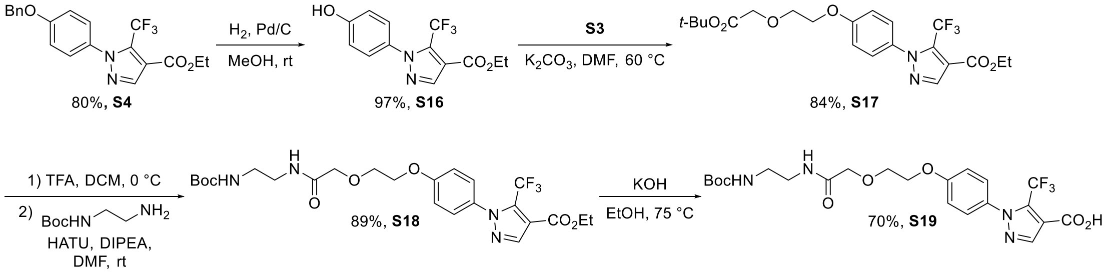

**Ethyl 1-(4-hydroxyphenyl)-5-(trifluoromethyl)-1*H*-pyrazole-4-carboxylate (S16).** A 50 ml round bottom flask was charged with ethyl 1-(4-(benzyloxy)yphenyl)-5-(trifluoromethyl)-1*H*-pyrazole-4-carboxylate (1.56 g, 4.00 mmol). The atmosphere was evacuated and refilled with Ar, MeOH (16 ml) was added to the flask, and the mixture was cooled to 0 °C in an ice bath. Palladium (213 mg, 10% w/w on carbon, 0.200 mmol, 5.00 mol%) was added. The atmosphere was purged with Ar and then with hydrogen. The ice bath was removed, and the mixture was stirred at ambient temperature under an atmosphere of hydrogen (balloon, ca. 1 atm) for 18 h. The balloon was removed, and the atmosphere was purged with Ar. The mixture was filtered through a pad of Celite, which was rinsed thrice with MeOH (2 ml each). Volatiles were removed from the filtrate at reduced pressure, and the crude product was purified by flash column chromatography (hexanes/ethyl acetate gradient) to give the title compound as a light yellow solid (1.16 g, 3.87 mmol, 97% yield).
$^1$H NMR (400 MHz, $CDCl_3$) δ 8.09 (s, 1H), 7.29 (d, J = 8.8 Hz, 2H), 6.92 (d, J = 8.8 Hz, 2H), 4.37 (q, J = 7.2 Hz, 2H), 1.38 (t, J = 7.1 Hz, 3H). $^{19}$F NMR (376 MHz, $CDCl_3$) δ -55.5. ESI$^+$ (LC/MS) calc'd 301.0795, found 301.0792.

**Ethyl 1-(4-(2-(2-(*tert*-butoxy)-2-oxoethoxy)ethoxy)phenyl)-5-(trifluoromethyl)-1*H*-pyrazole-4-carboxylate (S17).** A oven-dried scintillation vial was charged with ethyl 1-(4-hydroxyphenyl)-5-(trifluoromethyl)-1*H*-pyrazole-4-carboxylate (601 mg, 2.00 mmol) and $K_2CO_3$ (332 mg, 2.40 mmol, 1.20 equiv) and flushed with Ar. Dry DMF (3 ml) was added, and the mixture was heated at 60 °C with stirring for ca. 15 min. *Tert*-butyl 2-(2-(tosyloxy)ethoxy)acetate (727 mg, 2.20 mmol, 1.10 equiv) was dissolved in DMF (1 ml), and the solution was added

slowly to the reaction flask with stirring at 60 °C. The reaction mixture was heated at 60 °C with stirring overnight, at which point LC/MS indicated a high conversion of the starting material. The mixture was diluted with ethyl acetate (ca. 15 ml), transferred to a separatory funnel, and washed with sat. $NH_4Cl_{(aq)}$ and brine (ca. 15 ml each). The aqueous layers were extracted further with ethyl acetate (ca. 15 ml) in the same sequence, and the organic layers were dried over $Na_2SO_4$ and filtered. Volatiles were removed at reduced pressure, and the crude product was purified by flash column chromatography (hexanes/ethyl acetate gradient, Rf 0.26 in 20% EA/Hex) to give the title compound as a viscous yellow oil (767 mg, 1.67 mmol, 84% yield).
$^1$H NMR (400 MHz, $CDCl_3$) δ 8.08 (s, 1H), 7.32 (d, J = 8.9 Hz, 2H), 7.01 (d, J = 9.0 Hz, 2H), 4.37 (q, J = 7.2 Hz, 2H), 4.26 – 4.21 (m, 2H), 4.10 (s, 2H), 3.98 – 3.93 (m, 2H), 1.49 (s, 9H), 1.38 (t, J = 7.1 Hz, 3H). $^{19}$F NMR (376 MHz, $CDCl_3$) δ -55.6. ESI$^+$ (LC/MS) calc'd 481.1557, found 481.1557 ($MNa^+$). *Note: This sample contained an impurity (ca. 25 mol%) derived from the alkyl tosylate, which was carried through the next 2 steps.*

**Ethyl 1-(4-((2,2-dimethyl-4,9-dioxo-3,11-dioxa-5,8-diazatridecan-13-yl)oxy)phenyl)-5-(trifluoromethyl)-1*H*-pyrazole-4-carboxylate (S18).**
Step 1. Ethyl 1-(4-(2-(2-(*tert*-butoxy)-2-oxoethoxy)ethoxy)phenyl)-5-(trifluoromethyl)-1*H*-pyrazole-4-carboxylate (767 mg, 1.67 mmol) was dissolved in DCM (8 ml) in a scintillation vial under Ar. The mixture was cooled to 0 °C in an ice bath, and TFA (2 ml) was added slowly with stirring. The bath was removed, and the reaction mixture with stirred at ambient temperature for 2 h. Volatiles were evaporated under a stream of Ar. The residue was redissolved in DCM (4 ml), volatiles were again evaporated under a stream of Ar, and the same sequence repeated once more. The crude 2-(2-(4-(4-(ethoxycarbonyl)-5-(trifluoromethyl)-1*H*-pyrazol-1-yl)phenoxy)ethoxy)acetic acid was carried on to the following step without purification.
$^1$H NMR (400 MHz, $CDCl_3$) δ 8.09 (s, 1H), 7.35 (d, J = 8.9 Hz, 2H), 7.01 (d, J = 9.0 Hz, 2H), 4.37 (q, J = 7.2 Hz, 2H), 4.28 (s, 2H), 4.26 – 4.22 (m, 2H), 4.04 – 4.00 (m, 2H), 1.38 (t, J = 7.1 Hz, 3H). $^{19}$F NMR (376 MHz, $CDCl_3$) δ -55.5. ESI$^+$ (LC/MS) calc'd 403.1111, found 403.1122. *Note: The -COOH resonance was not located in the $^1$H NMR spectrum, likely due to fast exchange.*

Step 2. *N*-Boc-ethylenediamine (0.53 ml, 0.54 g, 3.4 mmol, 2.0 equiv) and HATU (763 mg, 2.01 mmol, 1.20 equiv) were added to the 20 ml scintillation vial containing crude 2-(2-(4-(4-(ethoxycarbonyl)-5-(trifluoromethyl)-1*H*-pyrazol-1-yl)phenoxy)ethoxy)acetic acid from the previous step. The atmosphere was flushed with Ar, the mixture was cooled to 0 °C in an ice bath, and DMF (8.5 ml) and DIPEA (0.58 ml, 3.3 mmol, 2.0 equiv) were added in that order. The cooling bath was removed, and the reaction mixture was stirred at ambient temperature overnight. The mixture was diluted with ethyl acetate (ca. 20 ml) and transferred to a separatory funnel. The mixture was washed with $HCl_{(aq)}$ (0.5 M, ca. 20 ml, and brine (ca. 20 ml). The aqueous layers were extracted further with EA (ca. 20 ml) in the same sequence. The organics layers were dried over $Na_2SO_4$ and filtered. Volatiles were removed at reduced pressure, and the crude product was purified by flash column chromatography (dichloromethane/methanol gradient, then hexanes/ethyl acetate gradient, Rf 0.52 in 10% MeOH/DCM) to give the title compound as a viscous clear and colorless oil (810 mg, 1.49 mmol, 89% yield).
$^1$H NMR (400 MHz, DMSO-$d_6$) δ 8.26 (s, 1H), 7.84 – 7.72 (m, 1H), 7.46 (d, J = 8.9 Hz, 2H), 7.12 (d, J = 9.0 Hz, 2H), 6.85 (t, J = 5.9 Hz, 1H), 4.31 (q, J = 7.1 Hz, 2H), 4.26 – 4.21 (m, 2H), 3.95 (s, 2H), 3.86 – 3.80 (m, 2H), 3.14 (q, J = 6.3 Hz, 2H), 3.00 (q, J = 6.2 Hz, 2H), 1.36 (s, 9H), 1.30 (t, J = 7.1 Hz, 3H). $^{19}$F NMR (376 MHz, DMSO-$d_6$) δ -54.8. ESI$^+$ (LC/MS) calc'd 567.2037, found 567.1982 ($MNa^+$).

**1-(4-((2,2-Dimethyl-4,9-dioxo-3,11-dioxa-5,8-diazatridecan-13-yl)oxy)phenyl)-5-(trifluoromethyl)-1*H*-pyrazole-4-carboxylic acid (S19).** Ethyl 1-(4-((2,2-dimethyl-4,9-dioxo-3,11-dioxa-5,8-diazatridecan-13-yl)oxy)phenyl)-5-(trifluoromethyl)-1*H*-pyrazole-4-carboxylate (809 mg, 1.49 mmol) was suspended in absolute EtOH (5 ml) in a scintillation vial under Ar. The mixture was heated at 75°C with stirring, and $KOH_{(aq)}$ (300 µl, 40% w/w, 420 mg, 2.99 mmol, 2.00 equiv) was added with stirring. The reaction mixture was stirred at 75 °C for 1 h. The mixture was cooled to 30 °C, and volatiles were evaporated under a stream of Ar. The crude conjugate base was dissolved in water (ca. 20 ml), transferred to a separatory funnel, and washed with ether (15 ml). The organic layer was extracted further with water (10 ml). The combined aqueous layers were acidified to pH <1 by adding $HCl_{(aq)}$ (2 ml, 3 M). The mixture was extracted twice with ethyl acetate (20 ml). The organic extracts were dried over $Na_2SO_4$ and filtered. Volatiles were removed at reduced pressure to give a white solid. The crude product was recrystallized by dissolving in boiling toluene (ca. 15 ml), filtering through cotton, and then cooling to 0 °C. The white solid was

collected by filtration, rinsed with cold hexanes (2x4 ml), and dried under vacuum to give the title compound as a white powder (536 mg, 1.04 mmol, 70% yield).
$^1$H NMR (400 MHz, DMSO-$d_6$) δ 13.28 (s, 1H), 8.19 (s, 1H), 7.79 (t, J = 5.9 Hz, 1H), 7.45 (d, J = 8.9 Hz, 2H), 7.11 (d, J = 9.0 Hz, 2H), 6.85 (t, J = 5.7 Hz, 1H), 4.29 – 4.21 (m, 2H), 3.95 (s, 2H), 3.87 – 3.79 (m, 2H), 3.14 (q, J = 6.3 Hz, 2H), 3.00 (q, J = 6.2 Hz, 2H), 1.36 (s, 9H). $^{19}$F NMR (376 MHz, DMSO-$d_6$) δ -54.6. ESI$^+$ (LC/MS) calc'd 539.1724, found 539.1724 (MNa$^+$).

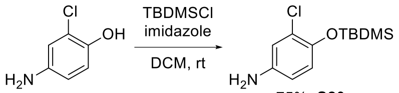

75%, **S20**

**4-((*tert*-Butyldimethylsilyl)oxy)-3-chloroaniline (S20).** 4-Amino-2-chlorophenol (574 mg, 4.00 mmol) was dissolved in dry DCM (8 ml) in a 20 ml scintillation vial under Ar. Imidazole (327 mg, 4.80 mmol, 2.40 equiv) was added with stirring, then the mixture was cooled to 0 °C in an ice bath. TBDMSCl (633 mg, 4.20 mmol, 1.05 equiv) was dissolved in dry DCM (2 ml), and this solution was added dropwise to the reaction vial with stirring at 0 °C. After 15 min, the ice bath was removed, and the reaction mixture was stirred at ambient temperature overnight. The mixture was transferred to a separatory funnel, diluted to ca. 20 ml with DCM, and washed with water (ca. 20 ml). A few drops of sat $NaHCO_{3(aq)}$ were added to improve the separation of layers. The aqueous layer was extracted further with DCM (ca. 20 ml). The organic layers were dried over $Na_2SO_4$ and filtered. Volatiles were removed at reduced pressure, and the crude product was purified by flash column chromatography (hexanes/ethyl acetate gradient, Rf 0.31 in 20% EA/Hex) to give the title compound as a red-brown oil (769 mg, 2.98 mmol, 75% yield).
$^1$H NMR (400 MHz, DMSO) δ 6.67 (d, *J* = 8.6 Hz, 1H), 6.60 (d, *J* = 2.7 Hz, 1H), 6.42 (dd, *J* = 8.6, 2.7 Hz, 1H), 4.90 (br, 2H), 0.97 (s, 9H), 0.14 (s, 6H). ESI$^+$ (LC/MS) calc'd 258.1076, found 258.1076.

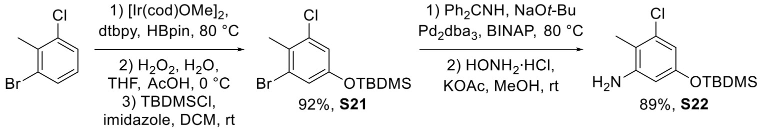

92%, **S21** 89%, **S22**

**5-((*tert*-Butyldimethylsilyl)oxy)-1-bromo-3-chloro-2-methylbenzene (S21).**
Step 1. (1,5-Cyclooctadiene)(methoxy)iridium(I) dimer (13.3 mg, 0.0200 mmol, 1.00 mol%) and 4,4-di-*tert*-butyl-2,2-bipyridyl (10.7 mg, 0.0400 mmol, 2.00 mol%) were placed in a 50 ml flame-dried Schlenk flask under Ar. The atmosphere was evacuated and refilled with Ar, then the flask was cooled to 0 °C in an ice bath. Pinacolborane (0.44 ml, 0.38 g, 3.0 mmol, 1.5 equiv) and 1-bromo-3-chloro-2-methylbenzene (0.26 ml, 0.41 g, 2.0 mmol) were added to the flask by syringe through a septum with stirring. The atmosphere was again evacuated and refilled with Ar, then the ice bath was removed. The reaction mixture was heated at 80 °C for 2 h. The mixture was cooled to ambient temperature, dissolved in DCM (ca. 2 ml), and filtered through a plug of silica, which was rinsed thrice with DCM (2 ml each). Volatiles were removed at reduced pressure. The crude 3-bromo-5-chloro-4-methylphenylboronic acid pinacol ester was carried on the next step without further purification.
$^1$H NMR (400 MHz, $CDCl_3$) δ 7.86 (s, 1H), 7.71 (s, 1H), 2.53 (s, 3H), 1.33 (s, 12H).

Step 2. The crude product from step 1 was suspended in a mixture of THF (5 ml), water (2.5 ml), and acetic acid (2.5 ml) in a 20 ml scintillation vial, and the mixture was cooled to 0 °C in an ice bath. $H_2O_{2(aq)}$ (30% w/w, 4.5 ml, 20 equiv) was slowly added to the reaction vial with stirring. The cooling bath was removed, and the mixture was stirred at ambient temperature for 4 h. The mixture was transferred to a separatory funnel and diluted with ethyl acetate and water until each layer was approximately 20 ml. Sat. $NaHCO_{3(aq)}$ was added dropwise to improve the separation of layers. The layers were separated, and the organic layer washed again with brine (ca. 20 ml). The aqueous layers were extracted further with ethyl acetate (ca. 20 ml) in the same sequence. The organic layers were dried over $Na_2SO_4$ and filtered. Volatiles were removed at reduced pressure, and the crude product was purified by flash column chromatography (hexanes/ethyl acetate gradient, Rf 0.47 in 20% EA/Hex) to give 3-bromo-5-chloro-4-methylphenol as a white solid. *Note: the measured mass (540 mg) is an overestimation due to the presence of residual solvent, which was not removed under vacuum to avoid inadvertently evaporating the desired product.*
$^1$H NMR (400 MHz, $CDCl_3$) δ 7.47 (s, 1H), 6.92 (s, 1H), 5.38 (s, 1H), 2.33 (s, 3H).

Step 3. Imidazole (163 mg, 2.40 mmol, 1.20 equiv) and dry DCM (9 ml) were added to the 20 ml scintillation vial containing the product from step 2 under Ar, and the mixture was cooled to 0 °C in an ice bath. *tert*-Butyldimethylchlorosilane (332 mg, 2.20 mmol, 1.10 equiv) was dissolved in dry DCM (1 ml), and this solution was added dropwise to the reaction vial with stirring. The cooling bath was removed, and the mixture was stirred at ambient temperature overnight. The solution was washed with water (ca. 10 ml), and the aqueous layers was extracted further with DCM (ca. 10 ml). The organic layers were filtered through $Na_2SO_4$, and volatiles were removed at reduced pressure. The crude product was purified by flash column chromatography (hexanes, Rf >0.8) to give the title compound as a clear and colorless oil (616 mg, 1.84 mmol, 92% yield over 3 steps).
$^1$H NMR (400 MHz, $CDCl_3$) δ 6.98 (d, *J* = 2.4 Hz, 1H), 6.83 (d, *J* = 2.5 Hz, 1H), 2.42 (s, 3H), 0.97 (s, 10H), 0.20 (s, 6H).

**5-((*tert*-Butyldimethylsilyl)oxy)-3-chloro-2-methylaniline (S22).**
Step 1. An oven-dried 1 dram vial was charged with 5-((*tert*-butyldimethylsilyl)oxy)-1-bromo-3-chloro-2-methylbenzene ( 134 mg, 0.399 mmol), BINAP (10 mg, 0.016 mmol, 4.0 mol%), $Pd_2dba_3$ (7.3 mg, 0.0080 mmol, 2.0 mol%), and benzophenone imine (74 µl, 0.44 mmol, 1.1 equiv) and flushed with Ar. In a separate oven-dried 10 ml flask, NaO*t*-Bu was dissolved in dry toluene to make a 0.24 M solution, which was then sparged with Ar. A portion of this NaO*t*-Bu solution (2.0 ml, 0.48 mmol, 1.2 equiv) was added to the reaction vial through a septum. The reaction mixture was heated at 80 °C with stirring for 16 h. After cooling to ambient temperature, the mixture was directly loaded onto a plug of silica and purified by flash column chromatography (hexanes/ethyl acetate gradient, Rf 0.76 in 20% EA/Hex) to give *N*-(5-((*tert*-butyldimethylsilyl)oxy)-3-chloro-2-methylphenyl)-1,1-diphenylmethanimine as a yellow oil (164 mg, 0.376 mmol, 94% yield).

Step 2. The product from step 1 was dissolved in methanol (2 ml) in a 20 ml scintillation vial. Sodium acetate (78 mg, 0.95 mmol, 2.4 equiv) and hydroxylamine hydrochloride (56 mg, 0.81 mmol, 2.0 equiv) were added in that order with stirring. The reaction mixture was stirred at ambient temperature for 90 min. Volatiles were removed at reduced pressure, and the residue was redissolved in ethyl acetate (ca. 2 ml) and washed with water (ca. 2 ml). The aqueous layer was extracted further with ethyl acetate (ca. 2 ml), the organic layers were filtered through $Na_2SO_4$, and volatiles were removed at reduced pressure. The crude product was purified by flash column chromatography (hexanes/ethyl acetate gradient, Rf 0.50 in 20% EA/Hex) to give the title compound as a clear and colorless oil (96.7 mg, 0.356 mmol, 89% yield over 2 steps).
$^1$H NMR (400 MHz, DMSO) δ 6.15 (d, *J* = 2.5 Hz, 1H), 6.07 (d, *J* = 2.4 Hz, 1H), 5.17 (br, 2H), 2.01 (s, 3H), 0.93 (s, 9H), 0.15 (s, 6H). ESI$^+$ (LC/MS) calc'd 272.1232, found 272.1230.

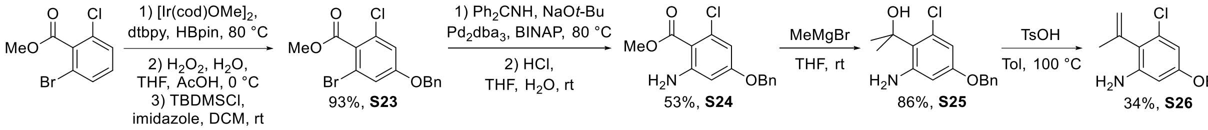

**Methyl 4-(benzyloxy)-2-bromo-6-chlorobenzoate (S23).**
Step 1. Pinacolborane (0.44 ml, 3.0 mmol, 1.5 equiv), 4,4-di-*tert*-butyl-2,2-bipyridyl (10.7 mg, 0.0399 mmol, 2.0 mol%), and (1,5-cyclooctadiene)(methoxy)iridium(I) dimer (13.3 mg, 0.0201 mmol, 1.0 mol%) were placed in a flame-dried 100 ml round bottom flask under Ar. The mixture was frozen in an acetone/dry ice bath, and the atmosphere was evacuated and refilled with Ar thrice. The mixture was allowed to warm to ambient temperature, and methyl 2-bromo-6-chlorobenzoate (499 mg, 2.00 mmol) was added with stirring. The atmosphere was evacuated and refilled with Ar. The reaction mixture was heated at 80 °C with stirring for 2 h, then cooled to ambient temperature. The atmosphere was evacuated and refilled with Ar. At this point, $^1$H NMR spectroscopy indicated high conversion of the arene. The mixture was dissolved in DCM (ca. 2 ml) and filtered through a pad of silica (ca. 2 ml slurry with DCM), which was rinsed with thrice with DCM (total filtrate volume ca. 8 ml). Volatiles were removed at reduced pressure. The crude (3-bromo-5-chloro-4-(methoxycarbonyl)phenyl)boronic acid pinacol ester was carried on the next step without further purification. $^1$H NMR (400 MHz, $CDCl_3$) δ 7.88 (d, *J* = 0.9 Hz, 1H), 7.75 (d, *J* = 0.9 Hz, 1H), 3.98 (s, 3H), 1.34 (s, 12H).

Step 2. The crude product from step 1 was suspended in a mixture of THF (5 ml), water (2.5 ml), and acetic acid (2.5 ml) in a 20 ml scintillation vial, and the mixture was cooled to 0 °C in an ice bath. $H_2O_{2(aq)}$ (30% w/w, 4.5 ml, 20 equiv) was slowly added to the reaction vial with stirring, and the mixture was stirred at 0 °C for 2 h. The mixture was transferred to a separatory funnel and diluted with ethyl acetate and water until each layer was approximately 20 ml. Sat. $NaHCO_{3(aq)}$ was added dropwise to improve the separation of layers (aqueous layer pH ca. 3). The layers were separated, and the organic layer washed again with brine (20 ml). The aqueous layers were extracted further with ethyl acetate (20 ml) in the same sequence. The organic layers were dried over $Na_2SO_4$ and filtered. Volatiles were removed at reduced pressure, and the crude product was purified by flash column chromatography (hexanes/ethyl acetate gradient, Rf 0.26 in 20% EA/Hex) to give methyl 2-bromo-6-chloro-4-hydroxybenzoate as a white solid (531 mg, 2.00 mmol, quant. yield).

Step 3. The product from step 2 was dissolved in dry acetone (10 ml) in a 100 ml round bottom flask under Ar. Potassium carbonate (0.55 g, 4.0 mmol, 2.0 equiv) and benzyl bromide (261 µl, 2.20 mmol, 1.1 equiv) were added with stirring in that order, and the reaction mixture was stirred at ambient temperature overnight. At this point, TLC indicated incomplete conversion. An addition portion of benzyl bromide (43.5 µl, 0.366 mmol, 0.18 equiv) was added, and the mixture was heated at 50 °C with stirring for 2 h. At this point, TLC indicated high conversion. The mixture was allowed to cool to ambient temperature and filtered through cotton, which was rinsed thrice with acetone (5 ml each). Volatiles were removed at reduced pressure, and the crude product was purified by flash column chromatography (hexanes/ethyl acetate gradient, Rf 0.53 in 20% EA/Hex) to give the title compound as a clear and colorless oil (660 mg, 1.86 mmol, 93% yield over 3 steps). $^1H$ NMR (400 MHz, $CDCl_3$) δ 7.44 – 7.33 (m, 5H), 7.11 (d, $J$ = 2.3 Hz, 1H), 6.97 (d, $J$ = 2.3 Hz, 1H), 5.05 (s, 2H), 3.95 (s, 3H).

**Methyl 2-amino-4-(benzyloxy)-6-chlorobenzoate (S24).**
Step 1. An oven-dried 1 dram vial was charged with methyl 4-(benzyloxy)-2-bromo-6-chlorobenzoate (356 mg, 1.00 mmol), BINAP (25 mg, 0.040 mmol, 4.0 mol%), and benzophenone imine (185 µl, 1.10 mmol, 1.1 equiv) and flushed with Ar. In a separate oven-dried 10 ml flask, NaO*t*-Bu was dissolved in dry toluene to make a 0.30 M solution. A portion of this NaO*t*-Bu solution (5.0 ml, 1.5 mmol, 1.5 equiv) was added to the reaction vial through a septum, and the mixture was sparged with Ar. $Pd_2dba_3$ (18 mg, 0.020 mol, 2.0 mol%) was added, and the vial was sealed with a PTFE-lined cap. The reaction mixture was heated at 80 °C with stirring overnight. After cooling to ambient temperature, the liquid was filtered through a plug of Celite. The sticky residue remaining in the vial was suspended in water (ca. 2 ml) and extracted twice with ethyl acetate (ca. 2 ml each), and the plug of Celite was rinsed with these organic extracts. Volatiles were removed at reduced pressure. The crude product was purified by flash column chromatography (hexanes/ethyl acetate gradient) to give methyl 4-(benzyloxy)-2-chloro-6-((diphenylmethylene)amino)benzoate as a yellow oil (291 mg).

Step 2. The product from step 1 was dissolved in THF (2 ml) and water (1 ml) in a 20 ml scintillation vial. $HCl_{(aq)}$ (3.0 M, 10 drops) was added with stirring, and the reaction mixture was stirred at ambient temperature overnight. The mixture was basified by adding $K_2CO_{3(aq)}$ (1.0 M, ca. 5 ml) and extracted twice with ethyl acetate (ca. 5 ml each). The organic layers were filtered through $Na_2SO_4$, and volatiles were removed at reduced pressure. The crude product was purified by flash column chromatography (hexanes/ethyl acetate gradient, Rf 0. 11 in 10% EA/Hex) to give the title compound as a light yellow oil (154 mg, 0.528 mmol, 53% yield).
$^1H$ NMR (400 MHz, DMSO) δ 7.45 – 7.31 (m, 5H), 6.34 (d, $J$ = 2.4 Hz, 1H), 6.33 (d, $J$ = 2.4 Hz, 1H), 6.03 (br, 2H), 5.06 (s, 2H), 3.78 (s, 3H).

**2-(2-Amino-4-(benzyloxy)-6-chlorophenyl)propan-2-ol (S25).** Methyl 2-amino-4-(benzyloxy)-6-chlorobenzoate (152 mg, 0.521 mmol) was dissolved in dry THF (1.5 ml) in a 20 ml scintillation vial under Ar. The solution was cooled to 0 C in an ice bath, and methylmagnesium bromide (3.0 M in $Et_2O$, 0.52 ml, 1.6 mmol, 3.0 equiv) was added with stirring. The cooling bath was removed, and the mixture was stirred at ambient temperature for 1 h, at which point TLC (50% EA/Hex) indicated incomplete conversion to a product with lower Rf. An additional portion of methylmagnesium bromide (0.10 ml, 0.30 mmol, 0.6 equiv) was added with stirring, and the reaction mixture was stirred at ambient temperature for an additional 1 h, at which point TLC indicated high conversion of the starting material. The reaction mixture was cooled to 0 °C in an ice bath, and the reaction was quenched by slowing adding a few drops of saturated $NHC_{4(aq)}$. The mixture was portioned between water and ethyl acetate (ca. 5 ml each). The layers were mixed and separated, and the organic layer was washed further with brine (5 ml). The aqueous layers were extracted twice further with ethyl acetate (5 ml each) in the same sequence. The organic layers were filtered through $Na_2SO_4$, and volatiles were removed at reduced pressure. The crude product was purified by flash column chromatography (hexanes/ethyl acetate gradient, Rf 0.71 in 50% EA/Hex) to give the title compound (131 mg, 0.449 mmol, 86% yield).

$^1$H NMR (400 MHz, DMSO) δ 7.43 – 7.29 (m, 5H), 6.22 (d, *J* = 2.8 Hz, 1H), 6.18 (d, *J* = 2.7 Hz, 1H), 6.15 (br, 2H), 5.60 (s, 1H), 4.98 (s, 2H), 1.64 (s, 6H).

**5-(Benzyloxy)-3-chloro-2-(prop-1-en-2-yl)aniline (S26).** 2-(2-Amino-4-(benzyloxy)-6-chlorophenyl)propan-2-ol (130 mg, 0.446 mmol) was suspended in dry toluene (2 ml) in a 20 ml scintillation vial under Ar. Tosic acid monohydrate (9.0 mg, 0.047 mmol, 11 mol%) and powdered molecular sieves (4 Å, one small scoop ca. 0.05 $cm^3$) were added, and the reaction mixture was heated at 100 °C with stirring for 3 h, and which point TLC and LC/MS analysis indicated complete conversion of the starting material to form 3 products (m/z = 234, 274, and 314). The mixture was purified by flash column chromatography (hexanes/ethyl acetate gradient, Rf 0.47 in 50% EA/Hex; followed by hexanes/DCM gradient, Rf 0.37 in 50% hexanes/DCM) to give the title compound as a viscous colorless oil (41.5 mg, 0.152 mmol, 34% yield).
$^1$H NMR (400 MHz, DMSO) δ 7.43 – 7.35 (m, 4H), 7.35 – 7.30 (m, 1H), 6.29 (d, *J* = 2.4 Hz, 1H), 6.28 (d, *J* = 2.4 Hz, 1H), 5.34 (p, *J* = 1.9 Hz, 1H), 5.00 (s, 2H), 4.99 (br, 2H), 4.84 – 4.82 (m, 1H), 1.90 – 1.88 (m, 3H). ESI$^+$ (LC/MS) calc'd 274.0993, found 274.0988.

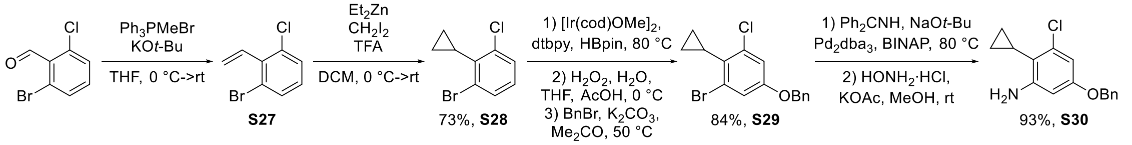

**1-Bromo-3-chloro-2-vinylbenzene (S27).** Methyltriphenylphosphonium bromide (1.72 g, 4.82 g, 1.20 equiv) was suspended in dry THF (4 ml) in a flame-dried 50 ml round bottom flask under Ar, and the suspension was cooled to 0 °C in an ice bath. A solution of potassium *tert*-butoxide in THF (1.0 M, 5.5 ml, 5.5 mmol, 1.4 equiv) was added slowly to the reaction flask with stirring, and the mixture was stirred for an additional 20 min. The cooling bath was removed, and the mixture was allowed to warm to ambient temperature over 10 min. 2-Bromo-6-chloro-benzaldehyde (878 mg, 4.00 mmol) was dissolved in dry THF (2 ml), and this solution was slowly added through a septum to the reaction flask with stirring. The reaction mixture was stirred at ambient temperature for 3 h. The reaction mixture was quenched by adding sat. $NH_4Cl_{(aq)}$ (2 ml) with stirring, transferred to a separatory funnel, diluted with ethyl acetate (ca. 15 ml), and washed with sat. $NH_4Cl_{(aq)}$ (ca. 20 ml). The organic layer was then washed with brine (ca. 20 ml), and the aqueous layers were extracted further with ethyl acetate (ca. 20 ml) in the same sequence. The organic layers were dried over $Na_2SO_4$ and filtered. Volatiles were removed at reduced pressure, and the crude product was purified by flash column chromatography (hexanes/ethyl acetate gradient, Rf 0.74 in 10% EA/Hex) to give the title compound as a clear and colorless oil. *Note: the measured mass (949 mg) is an overestimation due to the presence of residual solvent, which was not removed under vacuum to avoid inadvertently evaporating the desired product.*
$^1$H NMR (400 MHz, $CDCl_3$) δ 7.51 (dd, *J* = 8.0, 1.2 Hz, 1H), 7.36 (dd, *J* = 8.0, 1.2 Hz, 1H), 7.02 (t, *J* = 8.0 Hz, 1H), 6.66 (dd, *J* = 17.8, 11.6 Hz, 1H), 5.72 (dd, *J* = 8.9, 1.3 Hz, 1H), 5.68 (q, *J* = 1.3 Hz, 1H).

**1-Bromo-3-chloro-2-cyclopropylbenzene (S28).** Diethylzinc (1.0 M in hexanes, 12 ml, 12 mmol, 3.0 equiv) was added to a 50 ml flame-dried Schlenk flask under Ar, and the solution was cooled to 0 °C in an ice bath. TFA (0.93 ml, 1.4 g, 12 mmol, 3.0 equiv) was dissolved in dry, Ar-sparged DCM (4 ml) in a separate flame-dried 25 ml round bottom flask under Ar, and this solution was slowly transferred to the reaction flask containing diethylzinc with stirring by cannula. The resulting white suspension was stirred at 0 °C for an additional 20 min. In the flask that previously contained the TFA solution, diiodomethane (0.97 ml, 3.2 g, 12 mmol, 3.0 equiv) was dissolved in dry, Ar-sparged DCM (4 ml), and this solution was slowly transferred to the reaction flask with stirring by cannula. The reaction mixture was stirred at 0 °C for an additional 20 min. The sample of 1-bromo-3-chloro-2-vinylbenzene prepared in the previous step was dissolved in dry, Ar-sparged DCM (4 ml) in a 100 ml round bottom flask. This solution was slowly transferred to the reaction flask with stirring by cannula, and the flask and cannula were rinsed twice with DCM (4 ml each). The cooling bath was removed, and the reaction mixture was stirred at ambient temperature for 6 h. The mixture was cooled to 0 °C in an ice bath, and the reaction was quenched by slowly adding $HCl_{(aq)}$ (1.0 M, 25 ml). The mixture was transferred to a separatory funnel, the layers were separated, and the organic layer was washed further with sat. $NaHCO_{3(aq)}$ (ca. 20 ml). The aqueous layers were extracted further with DCM (ca. 20 ml) in the same sequence, and the organic layers were dried over $Na_2SO_4$ and filtered. Volatiles were removed at reduced pressure, and the crude product was purified by flash column chromatography (hexanes/ethyl

acetate gradient, Rf 0.74 in hexanes) to give the title compound as a clear and colorless oil (675 mg, 2.92 mmol, 73% yield). *Note: residual $CH_2I_2$ in the initially obtained sample of this compound was found to poison the subsequent Ir-catalyzed borylation reaction; repeated purification by column chromatography was required to reduce this impurity to ca. 2 mol% and obtain a suitable sample. The 675 mg yield noted here reflects the sample recovered and repurified from the unsuccessful initial attempts.*
$^1$H NMR (400 MHz, $CDCl_3$) δ 7.46 (dd, *J* = 8.0, 1.3 Hz, 1H), 7.30 (dd, *J* = 8.0, 1.3 Hz, 1H), 6.99 (td, *J* = 8.0, 0.9 Hz, 1H), 1.77 (tt, *J* = 8.5, 5.7 Hz, 1H), 1.21 – 1.15 (m, 2H), 0.81 – 0.73 (m, 2H).

**5-(Benzyloxy)-1-bromo-3-chloro-2-cyclopropylbenzene (S29).**
Step 1. 1-Bromo-3-chloro-2-cyclopropylbenzene (675 mg, 2.92 mmol), pinacolborane (0.63 ml, 4.3 mmol, 1.5 equiv), and 4,4-di-*tert*-butyl-2,2-bipyridyl (16 mg, 0.060 mmol, 2.0 mol%) were placed in a flame-dried 100 ml round bottom flask under Ar. The mixture was frozen in an acetone/dry ice bath, and the atmosphere was evacuated and refilled with Ar thrice. The mixture was allowed to warm to ambient temperature, and (1,5-cyclooctadiene)(methoxy)iridium(I) dimer (20 mg, 0.030 mmol, 1.0 mol%) was added with stirring to mix. The atmosphere was exchanged with the same freeze and cycle procedure described above. The reaction mixture was heated at 80 °C for 2.5 h, then cooled to ambient temperature. At this point, $^1$H NMR spectroscopy indicated ca. 8% conversion of the arene. Additional portions of (1,5-cyclooctadiene)(methoxy)iridium(I) dimer (20 mg, 0.030 mmol, 1.0 mol%), 4,4-di-*tert*-butyl-2,2-bipyridyl (16 mg, 0.060 mmol, 2.0 mol%), and pinacolborane (0.63 ml, 4.3 mmol, 1.5 equiv) were added, the atmosphere was exchanged by the same procedure described above, and the mixture was heated at 80 °C for an additional 2 h, then cooled to ambient temperature. At this point, $^1$H NMR spectroscopy indicated ca. 55% conversion of the arene. Additional portions of (1,5-cyclooctadiene)(methoxy)iridium(I) dimer (20 mg, 0.030 mmol, 1.0 mol%), 4,4-di-*tert*-butyl-2,2-bipyridyl (16 mg, 0.060 mmol, 2.0 mol%), and pinacolborane (0.63 ml, 4.3 mmol, 1.5 equiv) were added, the atmosphere was exchanged by the same procedure described above, and the mixture was heated at 80 °C for an additional 2 h, then cooled to ambient temperature. At this point, $^1$H NMR spectroscopy indicated high conversion of the arene. The mixture was dissolved in DCM (ca. 4 ml) and filtered through a pad of silica (ca. 10 ml slurry with DCM), which was rinsed with DCM (total filtrate volume ca. 30 ml). Volatiles were removed at reduced pressure. The crude 3-bromo-5-chloro-4-cyclopropylphenylboronic acid pinacol ester was carried on the next step without further purification.
$^1$H NMR (400 MHz, $CDCl_3$) δ 7.86 (d, *J* = 1.2 Hz, 1H), 7.70 (d, *J* = 1.2 Hz, 1H), 1.78 (tt, *J* = 8.6, 5.8 Hz, 1H), 1.33 (s, 12H), 1.21 – 1.15 (m, 2H), 0.80 – 0.74 (m, 2H).

Step 2. The crude product from step 1 was suspended in a mixture of THF (7 ml), water (3.5 ml), and acetic acid (3.5 ml) in a 100 ml round bottom flask, and the mixture was cooled to 0 °C in an ice bath. $H_2O_{2(aq)}$ (30% w/w, 6.6 ml, 20 equiv) was slowly added to the reaction with stirring, and the mixture was stirred at 0 °C for 2 h. The mixture was transferred to a separatory funnel and diluted with ethyl acetate and water until each layer was approximately 75 ml. Saturated $NaHCO_{3(aq)}$ was added dropwise to improve the separation of layers. The layers were separated, and the organic layer washed again with brine (ca. 100 ml). The aqueous layers were extracted further with ethyl acetate (ca. 75 ml) in the same sequence. The organic layers were dried over $Na_2SO_4$ and filtered. Volatiles were removed at reduced pressure, and the crude product was purified by flash column chromatography (hexanes/ethyl acetate gradient, Rf 0.42 in 20% EA/Hex) to give 3-bromo-5-chloro-4-cyclopropylphenol as an orange-brown solid. *Note: the measured mass (765 mg) is an overestimation due to the presence of residual solvent, which was not removed under vacuum to avoid inadvertently evaporating the desired product.*
$^1$H NMR (400 MHz, $CDCl_3$) δ 7.00 (d, *J* = 2.6 Hz, 1H), 6.84 (d, *J* = 2.6 Hz, 1H), 4.84 (br, 1H), 1.67 (tt, *J* = 8.4, 5.6 Hz, 1H), 1.17 – 1.08 (m, 2H), 0.76 – 0.64 (m, 2H).

Step 3. The product from step 2 was dissolved in dry acetone (15 ml) in a 50 ml round bottom flask under Ar. Potassium carbonate (0.85 g, 6.2 mmol, 2.0 equiv) was added, and the mixture was heated at 50 °C with stirring for 30 min. Then, benzyl bromide (0.40 ml, 0.58 g, 1.1 equiv) was added with stirring, and the reaction mixture was heated at 50 °C with stirring overnight. The mixture was allowed to cool to ambient temperature and filtered through cotton, which was rinsed thrice with acetone (5 ml each). Volatiles were removed at reduced pressure, and the crude product was purified by flash column chromatography (hexanes/ethyl acetate gradient, Rf 0.74 in 20% EA/Hex) to give the title compound as a clear and colorless oil (827 mg, 2.45 mmol, 84% yield over 3 steps).

$^{1}$H NMR (400 MHz, $CDCl_3$) δ 7.43 – 7.38 (m, 4H), 7.38 – 7.31 (m, 1H), 7.13 (d, *J* = 2.6 Hz, 1H), 6.96 (d, *J* = 2.6 Hz, 1H), 5.00 (s, 2H), 1.69 (tt, *J* = 8.5, 5.7 Hz, 1H), 1.16 – 1.08 (m, 2H), 0.75 – 0.68 (m, 2H).

**5-(Benzyloxy)-3-chloro-2-cyclopropylaniline (S30).**
Step 1. An oven-dried 1 dram vial was charged with 5-((*tert*-butyldimethylsilyl)oxy)-1-bromo-3-chloro-2-cyclopropylbenzene ( 135 mg, 0.400 mmol), BINAP (10 mg, 0.016 mmol, 4.0 mol%), and benzophenone imine (74 µl, 0.44 mmol, 1.1 equiv) and flushed with Ar. In a separate oven-dried 10 ml flask, NaO*t*-Bu was dissolved in dry toluene to make a 0.30 M solution. A portion of this NaO*t*-Bu solution (2.0 ml, 0.60 mmol, 1.5 equiv) was added to the reaction vial through a septum, and the mixture was sparged with Ar. $Pd_2dba_3$ (7.3 mg, 0.0080 mmol, 2.0 mol%) was added, and the vial was sealed with a PTFE-lined cap. The reaction mixture was heated at 80 °C for 15 h with stirring. After cooling to ambient temperature, the mixture was directly loaded onto a plug of silica and purified by flash column chromatography (hexanes/ethyl acetate gradient, Rf 0.65 in 20% EA/Hex) to give *N*-(5-(benzyloxy)-3-chloro-2-cyclopropylphenyl)-1,1-diphenylmethanimine as a yellow oil. *Note: the measured mass (201 mg) is an overestimation due to impurities.*

Step 2. The product from step 1 was dissolved in methanol (4 ml) in a 20 ml scintillation vial. Sodium acetate (78 mg, 0.95 mmol, 2.4 equiv) and hydroxylamine hydrochloride (56 mg, 0.81 mmol, 2.0 equiv) were added in that order with stirring. The reaction mixture was stirred at ambient temperature for 2 h. Volatiles were under a stream of Ar, and the residue was redissolved in DCM (ca. 4 ml) and water (ca. 4 ml). The aqueous layer was basified by adding a few drops of 1 M $NaOH_{(aq)}$, then the layers were mixed and separated. The aqueous layer was extracted twice more with DCM (ca. 4 ml each), the organic layers were filtered through $Na_2SO_4$, and volatiles were evaporated under a stream of Ar. The crude product was purified by flash column chromatography (hexanes/ethyl acetate gradient, then isocratic DCM, Rf 0.29 in 20% EA/Hex) to give the title compound as a clear and colorless oil (102 mg, 0.371 mmol, 93% yield over 2 steps).
$^{1}$H NMR (400 MHz, DMSO) δ 7.43 – 7.35 (m, 4H), 7.35 – 7.29 (m, 1H), 6.24 (s, 2H), 5.24 (br, 2H), 4.98 (s, 2H), 1.36 (tt, *J* = 8.2, 5.5 Hz, 1H), 1.04 – 0.93 (m, 2H), 0.48 – 0.39 (m, 2H). ESI$^{+}$ (LC/MS) calc'd 274.0993, found 274.0988.

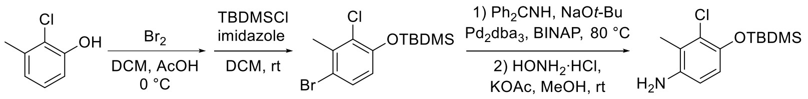

73%, **S31**
89%, **S32**

**4-((*tert*-Butyldimethylsilyl)oxy)-1-bromo-3-chloro-2-methylbenzene (S31).**
Step 1. 2-Chloro-3-methyl-phenol (715 mg, 5.02 mmol) was dissolved in DCM (7.5ml) and acetic acid (2.5 ml) in a 20 ml scintillation vial. The solution was cooled to 0 °C in an ice bath, and bromine (0.26 ml, 0.81 g, 5.1 mmol, 1.0 equiv) was added dropwise with stirring, allowing the red color to dissipate after each drop. The red color persisted after adding the last drop, and TLC indicated complete consumption of the starting material. The reaction was quenched by adding a saturated aqueous solution of sodium thiosulfate (ca. 2 ml) at 0 °C, then allowing the mixture to warm to ambient temperature. The mixture was transferred to a separatory funnel, and DCM and water were added until each layer was approximately 20 ml. The pH of the aqueous layer was adjusted to approximately 6 by adding sat. $NaHCO_{3(aq)}$. The layers were mixed and separated, and the aqueous layer was extracted further with DCM (ca. 20 ml). The organic layers were dried over $Na_2SO_4$ and filtered. Volatiles were removed at reduced pressure, and the crude product was purified by flash column chromatography (hexanes/ethyl acetate gradient, Rf 0.45 in 20% EA/Hex) to give 4-bromo-2-chloro-3-methyl-phenol.
$^{1}$H NMR (400 MHz, $CDCl_3$) δ 7.36 (d, *J* = 8.7 Hz, 1H), 6.79 (d, *J* = 8.8 Hz, 1H), 5.56 (s, 1H), 2.50 (s, 3H).

Step 2. Imidazole (409 mg, 6.00 mmol, 1.20 equiv) and dry DCM (20 ml) were added to the 250 ml round bottom flask containing the product from step 1, and the mixture was cooled to 0 °C in an ice bath. TBDMSCl (754 mg, 5.00 mmol, 1.00 equiv) was dissolved in dry DCM (5 ml), and this solution was added to the reaction flask slowly with stirring at 0 °C. The cooling bath was removed and the reaction mixture was stirred at ambient temperature overnight. The mixture was transferred to a separatory funnel and washed with water (ca. 25 ml). The aqueous layer was extracted further with DCM (ca. 25 ml). The organic layers were dried over $Na_2SO_4$ and filtered. Volatiles were removed at reduced pressure. The crude product was purified by flash column chromatography (hexanes/ethyl

acetate gradient, Rf 0.87 in 20% EA/hex) to give the title compound as a clear and colorless oil that turned to a white solid on standing (1.22 g, 3.65 mmol, 73% yield).
$^1$H NMR (400 MHz, $CDCl_3$) δ 7.30 (d, *J* = 8.7 Hz, 1H), 6.63 (d, *J* = 8.7 Hz, 1H), 2.50 (s, 3H), 1.03 (s, 9H), 0.22 (s, 6H).

**4-((*tert*-Butyldimethylsilyl)oxy)-3-chloro-2-methylaniline (S32).**
Step 1. An oven-dried 1 dram vial was charged with 4-((*tert*-butyldimethylsilyl)oxy)-1-bromo-3-chloro-2-methylbenzene ( 134 mg, 0.399 mmol), BINAP (10 mg, 0.016 mmol, 4.0 mol%), and benzophenone imine (74 µl, 0.44 mmol, 1.1 equiv) and flushed with Ar. In a separate oven-dried 25 ml flask, NaO*t*-Bu was dissolved in dry toluene to make a 0.24 M solution, which was then sparged with Ar. A portion of this NaO*t*-Bu solution (2.0 ml, 0.48 mmol, 1.2 equiv) was added to the reaction vial through a septum. $Pd_2dba_3$ (7.3 mg, 0.0080 mmol, 2.0 mol%) was added, and the vial was sealed with a PTFE-lined cap. The reaction mixture was heated at 80 °C with stirring for 16 h. After cooling to ambient temperature, the mixture was directly loaded onto a plug of silica and purified by flash column chromatography (hexanes/ethyl acetate gradient, Rf 0.75 in 20% EA/Hex) to give *N*-(4-((*tert*-butyldimethylsilyl)oxy)-3-chloro-2-methylphenyl)-1,1-diphenylmethanimine as a viscous yellow oil.

Step 2. The product from step 1 was dissolved in methanol (2 ml) in a 20 ml scintillation vial. Sodium acetate (78 mg, 0.95 mmol, 2.4 equiv) and hydroxylamine hydrochloride (56 mg, 0.81 mmol, 2.0 equiv) were added in that order with stirring. The reaction mixture was stirred at ambient temperature for 2 h. The mixture was filtered through a plug of celite, and volatiles were removed at reduced pressure. The crude product was purified by flash column chromatography (1 to 5% MeOH in DCM, Rf 0.78 in 10% MeOH/DCM) to give the title compound as a clear and colorless oil (96.6 mg, 0.355 mmol, 89% yield over 2 steps).
$^1$H NMR (400 MHz, DMSO) δ 6.59 (d, *J* = 8.6 Hz, 1H), 6.50 (d, *J* = 8.6 Hz, 1H), 4.68 (br, 2H), 2.10 (s, 3H), 0.97 (s, 9H), 0.14 (s, 6H). $ESI^+$ (LC/MS) calc'd 272.1232, found 272.1230.

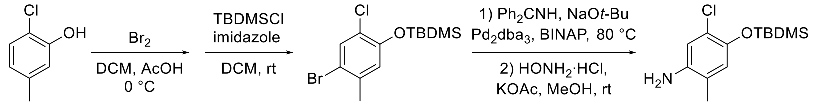

92%, **S33**
78%, **S34**

**4-((*tert*-Butyldimethylsilyl)oxy)-1-bromo-5-chloro-2-methylbenzene (S33).**
Step 1. 2-Chloro-5-methyl-phenol (715 mg, 5.02 mmol) was dissolved in DCM (7.5ml) and acetic acid (2.5 ml) in a 20 ml scintillation vial. The solution was cooled to 0 °C in an ice bath, and bromine (0.26 ml, 0.81 g, 5.1 mmol, 1.0 equiv) was added dropwise with stirring, allowing the red color to dissipate after each drop. The red color persisted after adding the last drop, and TLC indicated complete consumption of the starting material. The reaction was quenched by adding a saturated aqueous solution of sodium thiosulfate (ca. 2 ml) at 0 °C, then allowing the mixture to warm to ambient temperature. The mixture was transferred to a separatory funnel, and DCM and water were added until each layer was approximately 20 ml. The pH of the aqueous layer was adjusted to approximately 6 by adding sat. $NaHCO_{3(aq)}$. The layers were mixed and separated, and the aqueous layer was extracted further with DCM (ca. 20 ml). The organic layers were dried over $Na_2SO_4$ and filtered. Volatiles were removed at reduced pressure, and the crude product was purified by flash column chromatography (hexanes/ethyl acetate gradient, Rf 0.45 in 20% EA/Hex) to give 4-bromo-2-chloro-5-methyl-phenol.
$^1$H NMR (400 MHz, $CDCl_3$) δ 7.47 (s, 1H), 6.92 (s, 1H), 5.38 (s, 1H), 2.33 (s, 3H).

Step 2. Imidazole (409 mg, 6.00 mmol, 1.20 equiv) and dry DCM (20 ml) were added to the 250 ml round bottom flask containing the product from step 1, and the mixture was cooled to 0 °C in an ice bath. TBDMSCl (754 mg, 5.00 mmol, 1.00 equiv) was dissolved in dry DCM (5 ml), and this solution was added to the reaction flask slowly with stirring at 0 °C. The cooling bath was removed, and the reaction mixture was stirred at ambient temperature overnight. The mixture was transferred to a separatory funnel and washed with water (ca. 25 ml). The aqueous layer was extracted further with DCM (ca. 25 ml). The organic layers were dried over $Na_2SO_4$ and filtered. Volatiles were removed at reduced pressure. The crude product was purified by flash column chromatography (hexanes/ethyl acetate gradient, Rf 0.87 in 20% EA/hex) to give the title compound as a clear and colorless oil (1.55 g, 4.62 mmol, 92% yield).
$^1$H NMR (400 MHz, $CDCl_3$) δ 7.49 (s, 1H), 6.75 (s, 1H), 2.31 (s, 3H), 1.02 (s, 9H), 0.22 (s, 6H).

**4-((*tert*-Butyldimethylsilyl)oxy)-5-chloro-2-methylaniline (S34).**
Step 1. An oven-dried 1 dram vial was charged with 4-((*tert*-butyldimethylsilyl)oxy)-1-bromo-5-chloro-2-methylbenzene ( 134 mg, 0.399 mmol), BINAP (10 mg, 0.016 mmol, 4.0 mol%), and benzophenone imine (74 µl, 0.44 mmol, 1.1 equiv) and flushed with Ar. In a separate oven-dried 25 ml flask, NaO*t*-Bu was dissolved in dry toluene to make a 0.24 M solution, which was then sparged with Ar. A portion of this NaO*t*-Bu solution (2.0 ml, 0.48 mmol, 1.2 equiv) was added to the reaction vial through a septum. $Pd_2dba_3$ (7.3 mg, 0.0080 mmol, 2.0 mol%) was added, and the vial was sealed with a PTFE-lined cap. The reaction mixture was heated at 80 °C for 16 h with stirring. After cooling to ambient temperature, the mixture was directly loaded onto a plug of silica and purified by flash column chromatography (hexanes/ethyl acetate gradient, Rf 0.75 in 20% EA/Hex) to give *N*-(4-((*tert*-butyldimethylsilyl)oxy)-5-chloro-2-methylphenyl)-1,1-diphenylmethanimine as a viscous yellow oil.

Step 2. The product from step 1 was dissolved in methanol (2 ml) in a 20 ml scintillation vial. Sodium acetate (78 mg, 0.95 mmol, 2.4 equiv) and hydroxylamine hydrochloride (56 mg, 0.81 mmol, 2.0 equiv) were added in that order with stirring. The reaction mixture was stirred at ambient temperature for 2 h. The mixture was filtered through a plug of Celite, and volatiles were removed at reduced pressure. The crude product was purified by flash column chromatography (1 to 5% MeOH gradient in DCM, Rf 0.81 in 10% MeOH/DCM) to give the title compound as a clear and colorless oil (85.0 mg, 0.313 mmol, 78% yield over 2 steps).
$^1$H NMR (400 MHz, DMSO) δ 6.63 (s, 1H), 6.59 (s, 1H), 4.65 (br, 2H), 1.99 (s, 3H), 0.97 (s, 9H), 0.14 (s, 6H). ESI$^+$ (LC/MS) calc'd 272.1232, found 272.1230.

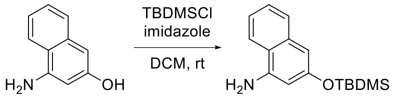

90%, **S35**

**3-((*tert*-Butyldimethylsilyl)oxy)naphthalen-1-amine (S35).** 4-Aminonaphthalen-2-ol (50.0 mg, 0.314 mmol) and imidazole (27.0 mg, 0.397 mmol, 1.26 equiv) were suspended in DCM (ca. 0.75 ml) in an oven-dried 1-dram vial under Ar. In a separate glass tube, TBDMSCl (48 mg, 0.32 mmol, 1.0 equiv) was dissolved in DCM (ca. 0.75 ml), and the resulting solution was slowly transferred to the reaction vial with stirring. The reactions mixture was stirred at ambient temperature for 30 min. The mixture was loaded directly onto silica and purified by flash column chromatography (hexanes/ethyl acetate gradient, Rf 0.25 in 20% EA/Hex) to give the title compound as a light orange oil (77.6 mg, 0.284 mmol, 90% yield).
$^1$H NMR (400 MHz, DMSO) δ 7.93 (d, *J* = 8.4 Hz, 1H), 7.57 (d, *J* = 8.1 Hz, 1H), 7.31 (ddd, *J* = 8.0, 6.7, 1.2 Hz, 1H), 7.17 (ddd, *J* = 8.2, 6.8, 1.3 Hz, 1H), 6.47 (d, *J* = 2.2 Hz, 1H), 6.31 (d, *J* = 2.2 Hz, 1H), 5.73 (s, 2H), 0.97 (s, 9H), 0.21 (s, 6H). ESI$^+$ (LC/MS) calc'd 274.1622, found 274.1631.

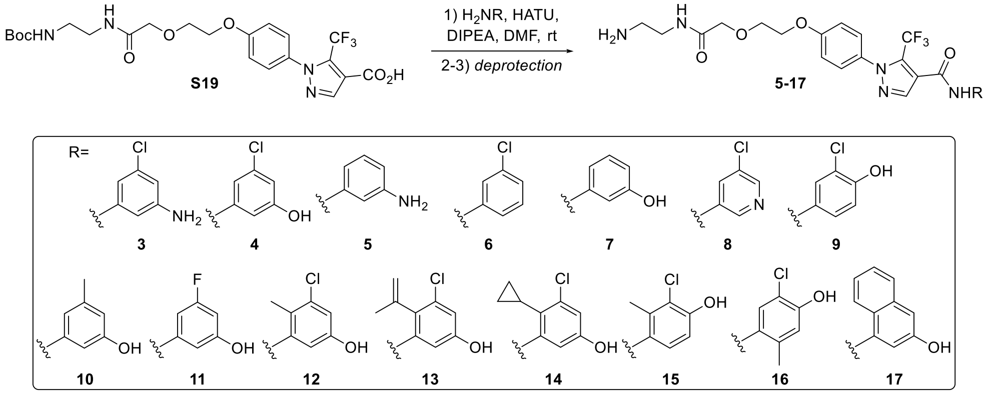

**General procedure for the preparation of 5-17.**
Step 1 (amide coupling). An oven-dried 1-dram vial was charged with 1-(4-((2,2-dimethyl-4,9-dioxo-3,11-dioxa-5,8-diazatridecan-13-yl)oxy)phenyl)-5-(trifluoromethyl)-1*H*-pyrazole-4-carboxylic acid, the amine coupling partner, and HATU and flushed with Ar. The mixture was cooled in an ice bath and DMF (0.2 M) and DIPEA were added in that order with stirring. The reaction mixture was stirred for approximately 10 min, then the cooling bath was removed, and the mixture was stirred at ambient temperature overnight. The mixture was diluted with EA (ca. 10x reaction volume) and washed with 0.5 M $HCl_{(aq)}$ (ca. 10x reaction volume). The aqueous layer was extracted twice

further with EA (ca. 10x reaction volume), and the organic layers were dried over $Na_2SO_4$ and filtered. Volatiles were removed at reduced pressure, and the crude product was purified by flash column chromatography.

Step 2-1 (benzyl ether hydrogenolysis). The sample obtained from step 1 was dissolved in MeOH in a scintillation vial. Tetrabutylammonium chloride was added, the vial was flushed with Ar, and the mixture was cooled to 0 °C in an ice bath. Palladium on carbon (10% w/w) was added with stirring. A balloon of hydrogen was attached, and the atmosphere was exchanged by purging. The cooling bath was removed, and the mixture was stirred at ambient temperature until LC/MS and/or TLC analysis indicated complete cleavage of the benzyl ether (typically overnight). The atmosphere was purged with Ar, and the mixture was filtered through a plug of Celite, which was rinsed thrice with MeOH. Volatiles were removed at reduced pressure, and the crude product was purified by flash column chromatography.

Step 2-2 (TBDMS ether cleavage). The sample obtained from step 1 was dissolved in THF in a scintillation vial. TBAF (1M in THF) was added in portions, and the mixture was stirred at ambient temperature until TLC analysis indicated high conversion of the TBDMS ether. Volatiles were evaporated under a stream of Ar, and the crude product was purified by flash column chromatography.

Step 2-3 or 3 (Boc cleavage). The sample obtained from step 1 or 2 was dissolved in DCM in a scintillation vial, and the solution was cooled to 0 °C in an ice bath. TFA was added slowly with stirring, and the mixture was stirred at 0 °C until LC/MS analysis indicated complete cleavage of the carbamate (30 min to 2 h). Volatiles were evaporated under a stream of Ar. The crude product was purified by preparative reverse-phase HPLC (C18, water/acetonitrile gradient + 0.1% formic acid or TFA). Pure fractions were combined, and MeCN was removed at reduced pressure. The resulting aqueous mixture was frozen and lyophilized to give the title compound.

***N*-(3-Aminophenyl)-1-(4-(2-(2-((2-aminoethyl)amino)-2-oxoethoxy)ethoxy)phenyl)-5-(trifluoromethyl)-1*H*-pyrazole-4-carboxamide (5).**

Step 1. The amide coupling was performed as described above with 1-(4-((2,2-dimethyl-4,9-dioxo-3,11-dioxa-5,8-diazatridecan-13-yl)oxy)phenyl)-5-(trifluoromethyl)-1*H*-pyrazole-4-carboxylic acid (25.8 mg, 0.0500 mmol), HATU (28.5, 0.0750 mmol, 1.5 equiv), *N*-Boc-*m*-phenylenediamine (11.5 mg, 0.0552 mmol, 1.1 equiv), DIPEA (27 µl, 0.15 mmol, 3.0 equiv), and DMF (0.25 ml). The crude product was purified by flash column chromatography (2 to 10% MeOH gradient in DCM, Rf 0.57 in 10% MeOH/DCM).

Step 2. The Boc cleavage was performed as described above with TFA (0.5 ml) in DCM (1.5 ml) at 0 °C for 2 h. The crude product was purified by preparative reverse-phase HPLC (C18, water/acetonitrile gradient + 0.1% formic acid) to give the title compound in the form of a 1:1 formic acid salt (6.5 mg, 0.012 mmol, 24% yield). $^{1}$H NMR (400 MHz, DMSO) δ 10.18 (s, 1H), 8.30 (s, 1H), 8.21 (s, 1H), 7.86 (br, 1H), 7.44 (d, *J* = 8.8 Hz, 2H), 7.14 (d, *J* = 8.9 Hz, 2H), 7.06 (s, 1H), 6.95 (t, *J* = 8.0 Hz, 1H), 6.76 (d, *J* = 7.9 Hz, 1H), 6.34 – 6.29 (m, 1H), 5.12 (br, 2H), 4.28 – 4.21 (m, 2H), 3.98 (s, 2H), 3.89 – 3.82 (m, 2H), 3.22 (q, *J* = 6.2 Hz, 2H), 2.72 (t, *J* = 6.3 Hz, 2H). $^{19}$F NMR (376 MHz, DMSO) δ -54.9. ESI$^+$ (LC/MS) calc'd 507.1962, found 507.1942. *Note: the -N*$\underline{H}_3^+$ *resonance was not located in the* $^{1}$*H NMR spectrum, and the -C*$\underline{H}_2$*N- resonance at 3.2 ppm is partially obscured by the broad* $H_2O$ *resonance.*

***N*-(3-Chlorophenyl)-1-(4-(2-(2-((2-aminoethyl)amino)-2-oxoethoxy)ethoxy)phenyl)-5-(trifluoromethyl)-1*H*-pyrazole-4-carboxamide (6).**

Step 1. The amide coupling was performed as described above with 1-(4-((2,2-dimethyl-4,9-dioxo-3,11-dioxa-5,8-diazatridecan-13-yl)oxy)phenyl)-5-(trifluoromethyl)-1*H*-pyrazole-4-carboxylic acid (25.8 mg, 0.0500 mmol), HATU (28.5, 0.0750 mmol, 1.5 equiv), 3-chloroaniline (7.6 mg, 0.060 mmol, 1.2 equiv), DIPEA (27 µl, 0.15 mmol, 3.0 equiv), and DMF (0.25 ml). The crude product was purified by flash column chromatography (2 to 10% MeOH gradient in DCM, Rf 0.51 in 10% MeOH/DCM).

Step 2. The Boc cleavage was performed as described above with TFA (0.5 ml) in DCM (1.5 ml) at 0 °C for 2 h. The crude product was purified by preparative reverse-phase HPLC (C18, water/acetonitrile gradient + 0.1% formic acid) to give the title compound in the form of a 1:1 formic acid salt (10.1 mg, 0.0177, 35% yield). $^{1}$H NMR (400 MHz, DMSO) δ 10.69 (br, 1H), 8.31 (br, 1H), 8.28 (s, 1H), 7.89 (t, *J* = 2.1 Hz, 1H), 7.85 (br, 1H), 7.60 (d, *J* = 8.7 Hz, 1H), 7.46 (d, *J* = 8.9 Hz, 2H), 7.40 (t, *J* = 8.1 Hz, 1H), 7.19 (dd, *J* = 8.1, 2.1 Hz, 1H), 7.14 (d, *J* = 9.0 Hz, 2H), 4.28 – 4.23 (m, 2H), 3.98 (s, 2H), 3.88 – 3.83 (m, 2H), 3.21 (q, *J* = 6.3 Hz, 2H), 2.71 (t, *J* = 6.4 Hz, 2H). $^{19}$F NMR

(376 MHz, DMSO) δ -55.0.ESI$^+$ (LC/MS) calc'd 526.1464, found 526.1456. *Note: the -N*$\underline{H}_3^+$ *resonance was not located in the* $^1$*H NMR spectrum, and the -C*$\underline{H}_2$*N- resonance at 3.2 ppm is partially obscured by the broad* $H_2O$ *resonance.*

**N-(3-Hydroxyphenyl)-1-(4-(2-(2-((2-aminoethyl)amino)-2-oxoethoxy)ethoxy)phenyl)-5-(trifluoromethyl)-1*H*-pyrazole-4-carboxamide (7).**

Step 1. The amide coupling was performed as described above with 1-(4-((2,2-dimethyl-4,9-dioxo-3,11-dioxa-5,8-diazatridecan-13-yl)oxy)phenyl)-5-(trifluoromethyl)-1*H*-pyrazole-4-carboxylic acid (25.8 mg, 0.0500 mmol), HATU (28.5, 0.0750 mmol, 1.5 equiv), 3-aminophenol (10.9 mg, 0.100 mmol, 2.0 equiv), DIPEA (35 µl, 0.20 mmol, 4.0 equiv), and DMF (0.25 ml). The crude product was purified by flash column chromatography (2 to 10% MeOH gradient in DCM, Rf 0.56 in 10% MeOH/DCM).

Step 2. The Boc cleavage was performed as described above with TFA (0.5 ml) in DCM (1.5 ml) at 0 °C for 2 h. The crude product was purified by preparative reverse-phase HPLC (C18, water/acetonitrile gradient + 0.1% formic acid) to give the title compound in the form of a 1:1 formic acid salt (10.2 mg, 0.0184, 37% yield). $^1$H NMR (400 MHz, DMSO) δ 10.37 (s, 1H), 9.46 (br, 1H), 8.31 (s, 1H), 8.23 (s, 1H), 7.86 (br, 1H), 7.45 (d, *J* = 8.9 Hz, 2H), 7.29 (s, 1H), 7.18 – 7.09 (m, 3H), 7.06 (d, *J* = 7.9 Hz, 1H), 6.51 (d, *J* = 7.0 Hz, 1H), 4.29 – 4.21 (m, 2H), 3.98 (s, 2H), 3.89 – 3.82 (m, 2H), 3.22 (q, *J* = 6.2 Hz, 2H), 2.72 (t, *J* = 6.3 Hz, 2H). $^{19}$F NMR (376 MHz, DMSO) δ -55.0. ESI$^+$ (LC/MS) calc'd 508.1803, found 508.1812. *Note: the -N*$\underline{H}_3^+$ *resonance was not located in the* $^1$*H NMR spectrum, and the -C*$\underline{H}_2$*N- resonance at 3.2 ppm is partially obscured by the broad* $H_2O$ *resonance.*

**N-(5-Chloropyridin-3-yl)-1-(4-(2-(2-((2-aminoethyl)amino)-2-oxoethoxy)ethoxy)phenyl)-5-(trifluoromethyl)-1*H*-pyrazole-4-carboxamide (8).**

Step 1. The amide coupling was performed as described above with 1-(4-((2,2-dimethyl-4,9-dioxo-3,11-dioxa-5,8-diazatridecan-13-yl)oxy)phenyl)-5-(trifluoromethyl)-1*H*-pyrazole-4-carboxylic acid (25.8 mg, 0.0500 mmol), HATU (28.5, 0.0750 mmol, 1.5 equiv), 5-chloropyridin-3-amine (7.7 mg, 0.060 mmol, 1.2 equiv), DIPEA (35 µl, 0.20 mmol, 4.0 equiv), and DMF (0.25 ml). The crude product was purified by flash column chromatography (2 to 10% MeOH gradient in DCM, Rf 0.49 in 10% MeOH/DCM).

Step 2. The Boc cleavage was performed as described above with TFA (0.25 ml) in DCM (0.75 ml) at 0 °C for 1 h. The crude product was purified by preparative reverse-phase HPLC (C18, water/acetonitrile gradient + 0.1% formic acid) to give the title compound in the form of a 1:1 formic acid salt (11.8 mg, 0.0206, 41% yield). $^1$H NMR (400 MHz, DMSO) δ 10.96 (br, 1H), 8.78 (d, *J* = 2.2 Hz, 1H), 8.40 (d, *J* = 2.3 Hz, 1H), 8.35 – 8.28 (m, 3H), 7.86 (br, 1H), 7.47 (d, *J* = 8.9 Hz, 2H), 7.15 (d, *J* = 9.0 Hz, 2H), 4.28 – 4.22 (m, 2H), 3.98 (s, 2H), 3.88 – 3.82 (m, 2H), 3.22 (q, *J* = 6.4 Hz, 2H), 2.72 (t, *J* = 6.3 Hz, 2H). $^{19}$F NMR (376 MHz, DMSO) δ -55.0. ESI$^+$ (LC/MS) calc'd 527.1416, found 527.1415. *Note: the -N*$\underline{H}_3^+$ *resonance was not located in the* $^1$*H NMR spectrum, and the -C*$\underline{H}_2$*N- resonance at 3.2 ppm is partially obscured by the broad* $H_2O$ *resonance.*

**N-(3-Chloro-4-hydroxyphenyl)-1-(4-(2-(2-((2-aminoethyl)amino)-2-oxoethoxy)ethoxy)phenyl)-5-(trifluoromethyl)-1*H*-pyrazole-4-carboxamide (9).**

Step 1. The amide coupling was performed as described above with 1-(4-((2,2-dimethyl-4,9-dioxo-3,11-dioxa-5,8-diazatridecan-13-yl)oxy)phenyl)-5-(trifluoromethyl)-1*H*-pyrazole-4-carboxylic acid (51.6 mg, 0.100 mmol), HATU (57 mg, 0.15 mmol, 1.5 equiv), 4-((*tert*-butyldimethylsilyl)oxy)-3-chloroaniline (31 mg, 0.12 mmol, 1.2 equiv), DIPEA (35 µl, 0.20 mmol, 2.0 equiv), and DMF (2 ml). *Modification: the amine coupling partner was added after stirring the carboxylic acid, HATU, DMF, and DIPEA for 1 h at ambient temperature.* The crude product from step 1 was carried on to step 2 without purification.

Step 2. The TBDMS cleavage was performed as described above with two portions of TBAF (0.10 ml each, 1.0 M in THF, 1.0 equiv each) added 1 h apart in THF (1 ml) at ambient temperature. Full cleavage was observed 1 h after adding the second portion. The crude product was purified by flash column chromatography (2 to 15% MeOH gradient in DCM, Rf 0.29 in 10% MeOH/DCM).

Step 3. The Boc cleavage was performed as described above with TFA (0.5 ml) in DCM (1.5 ml) at 0 °C for 1 h. The crude product was purified by preparative reverse-phase HPLC (C18, water/acetonitrile gradient + 0.1% TFA) to give the title compound in the form of a 1:1 TFA salt (61.0 mg, 0.0930 mmol, 93% yield). $^1$H NMR (600 MHz,

DMSO) δ 10.40 (d, $J$ = 4.5 Hz, 1H), 10.06 (br, 1H), 8.24 (s, 1H), 7.99 (q, $J$ = 5.4 Hz, 1H), 7.77 (d, $J$ = 2.5 Hz, 1H), 7.73 (br, 3H), 7.45 (d, $J$ = 8.8 Hz, 2H), 7.40 (dd, $J$ = 8.8, 2.6 Hz, 1H), 7.14 (d, $J$ = 8.9 Hz, 2H), 6.96 (d, $J$ = 8.7 Hz, 1H), 4.29 – 4.22 (m, 2H), 4.01 (s, 2H), 3.90 – 3.83 (m, 2H), 3.35 (q, $J$ = 6.2 Hz, 2H), 2.89 (h, $J$ = 6.0 Hz, 2H). $^{13}$C NMR (151 MHz, DMSO) δ 170.1, 159.3, 158.6, 149.5, 139.5, 131.7, 131.1, 129.8 (q, $J$ = 38.9 Hz), 127.5, 121.4, 121.1, 119.9, 119.4 (q, $J$ = 270.2 Hz), 119.1, 116.5, 114.9, 70.1, 69.4, 67.4, 38.8, 36.1. $^{19}$F NMR (376 MHz, DMSO) δ -54.9.ESI$^+$ (LC/MS) calc'd 542.1413, found 542.1427. *Note: the $^{13}$C and $^{19}$F NMR resonances of the $F_3CCO_2^-$ anion are not included in this list.*

**$N$-(3-Hydroxy-5-methylphenyl)-1-(4-(2-(2-((2-aminoethyl)amino)-2-oxoethoxy)ethoxy)phenyl)-5-(trifluoromethyl)-1$H$-pyrazole-4-carboxamide (10).**
Step 1. The amide coupling was performed as described above with 1-(4-((2,2-dimethyl-4,9-dioxo-3,11-dioxa-5,8-diazatridecan-13-yl)oxy)phenyl)-5-(trifluoromethyl)-1*H*-pyrazole-4-carboxylic acid (25.8 mg, 0.0500 mmol), HATU (28.5, 0.0750 mmol, 1.5 equiv), 3-amino-5-methylphenol (9.2 mg, 0.075 mmol, 1.5 equiv), and DIPEA (0.60 M in DMF, 0.25 ml, 0.15 mmol, 3.0 equiv). The crude product was purified by flash column chromatography (2 to 10% MeOH gradient in DCM, Rf 0.41 in 10% MeOH/DCM).

Step 2. The Boc cleavage was performed as described above with TFA (0.25 ml) in DCM (0.75 ml) at 0 °C for 1 h. The crude product was purified by preparative reverse-phase HPLC (C18, water/acetonitrile gradient + 0.1% formic acid) to give the title compound in the form of a 1:1 formic acid salt (11.8 mg, 0.0208, 42% yield). $^1$H NMR (400 MHz, DMSO) δ 10.28 (s, 1H), 9.34 (br, 1H), 8.32 (s, 1H), 8.22 (s, 1H), 7.86 (br, 1H), 7.44 (d, $J$ = 8.8 Hz, 2H), 7.14 (d, $J$ = 9.0 Hz, 2H), 7.07 (s, 1H), 6.93 (s, 1H), 6.34 (s, 1H), 4.28 – 4.23 (m, 2H), 3.98 (s, 2H), 3.88 – 3.83 (m, 2H), 3.24 – 3.17 (m, 2H), 2.75 – 2.69 (m, 2H), 2.20 (s, 3H). $^{19}$F NMR (376 MHz, DMSO) δ -55.0. ESI$^+$ (LC/MS) calc'd 522.1959, found 522.1951. *Note: the -N$\underline{H}_3^+$ resonance was not located in the $^1$H NMR spectrum, and the -C$\underline{H}_2$N- resonance at 3.2 ppm is partially obscured by the broad $H_2O$ resonance.*

**$N$-(3-Fluoro-5-hydroxyphenyl)-1-(4-(2-(2-((2-aminoethyl)amino)-2-oxoethoxy)ethoxy)phenyl)-5-(trifluoromethyl)-1$H$-pyrazole-4-carboxamide (11).**
Step 1. The amide coupling was performed as described above with 1-(4-((2,2-dimethyl-4,9-dioxo-3,11-dioxa-5,8-diazatridecan-13-yl)oxy)phenyl)-5-(trifluoromethyl)-1*H*-pyrazole-4-carboxylic acid (25.8 mg, 0.0500 mmol), HATU (28.6, 0.0752 mmol, 1.5 equiv), 3-benzyloxy-5-fluoroaniline (16.3 mg, 0.0750 mmol, 1.5 equiv), and DIPEA/DMF solution (0.60 M, 0.25 ml, 0.15 mmol, 3.0 equiv). The crude product was purified by flash column chromatography (2 to 10% MeOH gradient in DCM, Rf 0.43 in 10% MeOH/DCM).

Step 2. The benzyl ether hydrogenolysis was performed as described above with Pd/C (3 mg, 0.003 mmol, 5 mol%) and MeOH (1 ml). The crude product was purified by flash column chromatography (2 to 15% MeOH gradient in DCM, Rf 0.39 in 10% MeOH/DCM).

Step 3. The Boc cleavage was performed as described above with TFA (0.25 ml) in DCM (0.75 ml) at 0 °C for 1 h. The crude product was purified by preparative reverse-phase HPLC (C18, water/acetonitrile gradient + 0.1% formic acid) to give the title compound in the form of a 1:1 formic acid salt (11.8 mg, 0.0206, 41% yield). $^1$H NMR (400 MHz, DMSO) δ 10.55 (s, 1H), 8.34 (s, 1H), 8.25 (s, 1H), 7.87 (t, $J$ = 5.5 Hz, 1H), 7.45 (d, $J$ = 8.9 Hz, 2H), 7.14 (d, $J$ = 8.9 Hz, 2H), 7.08 – 6.99 (m, 2H), 6.32 (dt, $J$ = 10.8, 2.3 Hz, 1H), 4.28 – 4.22 (m, 2H), 3.98 (s, 2H), 3.89 – 3.82 (m, 2H), 3.21 (q, $J$ = 6.2 Hz, 2H), 2.71 (t, $J$ = 6.4 Hz, 2H). $^{19}$F NMR (376 MHz, DMSO) δ -55.0, -112.0 (t, $J$ = 10.8 Hz). ESI$^+$ (LC/MS) calc'd 526.1708, found 526.1697. *Note: the -N$\underline{H}_3^+$ and -O$\underline{H}$ resonances were not located in the $^1$H NMR spectrum, and the -C$\underline{H}_2$N- resonance at 3.2 ppm is partially obscured by the broad $H_2O$ resonance.*

**$N$-(3-Chloro-5-hydroxy-2-methylphenyl)-1-(4-(2-(2-((2-aminoethyl)amino)-2-oxoethoxy)ethoxy)phenyl)-5-(trifluoromethyl)-1$H$-pyrazole-4-carboxamide (12).**
Step 1. The amide coupling was performed as described above with 1-(4-((2,2-dimethyl-4,9-dioxo-3,11-dioxa-5,8-diazatridecan-13-yl)oxy)phenyl)-5-(trifluoromethyl)-1*H*-pyrazole-4-carboxylic acid (51.6 mg, 0.100 mmol), HATU (57 mg, 0.15 mmol, 1.5 equiv), 5-((*tert*-butyldimethylsilyl)oxy)-3-chloro-2-methylaniline (32 mg, 0.12 mmol, 1.2 equiv), DIPEA (35 µl, 0.20 mmol, 2.0 equiv), and DMF (2 ml). *Modification: the amine coupling partner was added after stirring the carboxylic acid, HATU, DMF, and DIPEA for 1 h at ambient temperature.* The crude product from step 1 was carried on to step 2 without purification.

Step 2. The TBDMS cleavage was performed as described above with two portions of TBAF (0.10 ml each, 1.0 M in THF, 2.0 equiv) added 1 h apart in THF (1 ml) at ambient temperature. Full cleavage was determined 1 h after adding the second portion. The crude product was purified by flash column chromatography (2 to 15% MeOH gradient in DCM, Rf 0.44 in 10% MeOH/DCM).

Step 3. The Boc cleavage was performed as described above with TFA (0.5 ml) in DCM (1.5 ml) at 0 °C for 1 h. The crude product was purified by preparative reverse-phase HPLC (C18, water/acetonitrile gradient + 0.1% TFA) to give the title compound in the form of a 1:1 TFA salt (49.8 mg, 0.0743 mmol, 74% yield). $^{1}$H NMR (600 MHz, DMSO) δ 10.13 (s, 1H), 9.80 (br, 1H), 8.28 (s, 1H), 7.99 (q, *J* = 5.5 Hz, 1H), 7.74 (br, 3H), 7.46 (d, *J* = 8.8 Hz, 2H), 7.14 (d, *J* = 8.9 Hz, 2H), 6.84 (s, 1H), 6.77 (d, *J* = 2.5 Hz, 1H), 4.28 – 4.23 (m, 2H), 4.01 (s, 2H), 3.89 – 3.85 (m, 2H), 3.35 (q, *J* = 6.2 Hz, 2H), 2.89 (h, *J* = 5.9 Hz, 2H), 2.14 (s, 3H). $^{13}$C NMR (151 MHz, DMSO) δ 170.1, 159.3, 155.5, 139.5, 137.5, 133.8, 131.7, 129.9 (q, *J* = 38.7 Hz), 127.5, 121.3, 120.8, 119.4 (q, 270.5 Hz), 114.9, 113.9, 112.4, 70.1, 69.4, 67.4, 38.8, 36.1, 14.2. $^{19}$F NMR (376 MHz, DMSO) δ -54.8. ESI$^+$ (LC/MS) calc'd 556.1569, found 556.1605. *Note: the $^{13}$C and $^{19}$F NMR resonances of the $F_3CCO_2^-$ anion are not included in this list.*

**N-(3-Chloro-5-hydroxy-2-(prop-1-en-2-yl)phenyl)-1-(4-(2-(2-((2-aminoethyl)amino)-2-oxoethoxy)ethoxy)phenyl)-5-(trifluoromethyl)-1H-pyrazole-4-carboxamide (13).**

Step 1. The amide coupling was performed as described above with 1-(4-((2,2-dimethyl-4,9-dioxo-3,11-dioxa-5,8-diazatridecan-13-yl)oxy)phenyl)-5-(trifluoromethyl)-1*H*-pyrazole-4-carboxylic acid (25.8 mg, 0.0500 mmol), HATU (28.5, 0.0750 mmol, 1.5 equiv), 5-(benzyloxy)-3-chloro-2-(prop-1-en-2-yl)aniline (14 mg, 0.051 mmol, 1.0 equiv), DIPEA (17 µl, 0.10 mmol, 2.0 equiv), and DMF (0.25 ml). After 3 d, LC/MS analysis indicated low conversion of the carboxylic acid. An additional portion of HATU (14 mg, 0.037 mmol, 0.75 equiv) was added, the mixture was stirred for another 1 d, a second additional portion of HATU was added (14 ml, 0.037 mmol, 0.75 equv), and the mixture was stirred for another 1 d. The crude product was purified by flash column chromatography (2 to 15% MeOH gradient in DCM, Rf 0.43 in 10% MeOH/DCM).

Step 2. The benzyl ether hydrogenolysis was performed as described above with Pd/C (6 mg, 0.006 mmol, 0.1 equiv), TBACl (14 mg, 0050 mmol, 1.0 equiv), and MeOH (1 ml). The crude product was purified by flash column chromatography (2 to 15% MeOH gradient in DCM, Rf 0.39 in 10% MeOH/DCM).

Step 3. The Boc cleavage was performed as described above with TFA (0.5 ml) in DCM (1.5 ml) at 0 °C for 30 min. The crude product was purified by preparative reverse-phase HPLC (C18, water/acetonitrile gradient + 0.1% formic acid) to give the title compound in the form of a 1:1 formic acid salt (10.1 mg, 0.0161, 32% yield). $^{1}$H NMR (400 MHz, DMSO) δ 9.59 (s, 1H), 8.29 (s, 1H), 8.11 (s, 1H), 7.82 (br, 1H), 7.45 (d, *J* = 8.9 Hz, 2H), 7.13 (d, *J* = 9.0 Hz, 2H), 7.03 (d, *J* = 2.5 Hz, 1H), 6.77 (d, *J* = 2.4 Hz, 1H), 5.34 – 5.31 (m, 1H), 4.86 (s, 1H), 4.28 – 4.22 (m, 2H), 3.98 (s, 2H), 3.88 – 3.83 (m, 2H), 3.23 – 3.16 (m, 2H), 2.74 – 2.68 (m, 2H), 1.96 (s, 3H). $^{19}$F NMR (376 MHz, DMSO) δ -54.9. ESI$^+$ (LC/MS) calc'd 582.1726, found 582.1712. *Note: the -N$\underline{H}_3^+$ and -O$\underline{H}$ resonances were not located in the $^{1}$H NMR spectrum, and the -C$\underline{H}_2$N- resonance at 3.2 ppm is partially obscured by the broad $H_2O$ resonance.*

**N-(3-Chloro-2-cyclopropyl-5-hydroxyphenyl)-1-(4-(2-(2-((2-aminoethyl)amino)-2-oxoethoxy)ethoxy)phenyl)-5-(trifluoromethyl)-1H-pyrazole-4-carboxamide (14).**

Step 1. The amide coupling was performed as described above with 1-(4-((2,2-dimethyl-4,9-dioxo-3,11-dioxa-5,8-diazatridecan-13-yl)oxy)phenyl)-5-(trifluoromethyl)-1*H*-pyrazole-4-carboxylic acid (25.8 mg, 0.0500 mmol), HATU (28.6, 0.0752 mmol, 1.5 equiv), 5-(benzyloxy)-3-chloro-2-cyclopropylaniline (14 mg, 0.0750 mmol, 1.5 equiv), and DIPEA (26 µl, 0.15 mmol, 3.0 equiv), and DMF (1 ml). After 3 d, LC/MS analysis indicated low conversion of the carboxylic acid. The reaction mixture was heated at 50 °C with stirring for an additional 1 d. An additional portion of HATU (14 mg, 0.037 mmol, 0.75 equiv) was added, and the mixture was stirred for another 1 d. The crude product was purified by flash column chromatography (2 to 15% MeOH gradient in DCM).

Step 2. The benzyl ether hydrogenolysis was performed as described above with Pd/C (3 mg, 0.003 mmol, 5 mol%), TBACl (14 mg, 0050 mmol, 1.0 equiv), and MeOH (1 ml). The crude product was purified by flash column chromatography (2 to 15% MeOH gradient in DCM).

Step 3. The Boc cleavage was performed as described above with TFA (0.5 ml) in DCM (1.5 ml) at 0 °C for 1 h. The crude product was purified by preparative reverse-phase HPLC (C18, water/acetonitrile gradient + 0.1% formic acid) to give the title compound in the form of a 1:1 formic acid salt (7.2 mg, 0.011, 23% yield). $^1$H NMR (400 MHz, DMSO) δ 10.04 (s, 1H), 8.35 – 8.18 (m, 2H), 7.88 (br, 1H), 7.47 (d, $J$ = 8.5 Hz, 2H), 7.14 (d, $J$ = 8.9 Hz, 2H), 6.93 (d, $J$ = 2.5 Hz, 1H), 6.72 (d, $J$ = 2.5 Hz, 1H), 4.30 – 4.21 (m, 2H), 3.99 (s, 2H), 3.90 – 3.83 (m, 2H), 3.28 – 3.21 (m, 2H), 2.84 – 2.71 (m, 2H), 1.66 – 1.55 (m, 1H), 1.00 – 0.91 (m, 2H), 0.50 – 0.43 (m, 2H). $^{19}$F NMR (376 MHz, DMSO) δ -54.8. ESI$^+$ (LC/MS) calc'd 582.1726, found 582.1712. *Note: the -N$\underline{H}_3{}^+$ and -O$\underline{H}$ resonances were not located in the $^1$H NMR spectrum, and the -C$\underline{H}_2$N- resonance at 3.2 ppm is partially obscured by the broad $H_2O$ resonance.*

**_N_-(3-Chloro-4-hydroxy-2-methylphenyl)-1-(4-(2-(2-((2-aminoethyl)amino)-2-oxoethoxy)ethoxy)phenyl)-5-(trifluoromethyl)-1_H_-pyrazole-4-carboxamide (15).**

Step 1. The amide coupling was performed as described above with 1-(4-((2,2-dimethyl-4,9-dioxo-3,11-dioxa-5,8-diazatridecan-13-yl)oxy)phenyl)-5-(trifluoromethyl)-1*H*-pyrazole-4-carboxylic acid (25.8 mg, 0.0500 mmol), HATU (22.8 mg, 0.0600 mmol, 1.2 equiv), 4-((*tert*-butyldimethylsilyl)oxy)-3-chloro-2-methylaniline (20.4 mg, 0.0750 mmol, 1.5 equiv), DIPEA (17 µl, 0.10 mmol, 2.0 equiv), and DMF (0.25 ml). *Modification: the amine coupling partner was added after stirring the carboxylic acid, HATU, DMF, and DIPEA for 1 h at ambient temperature.* The crude product was purified by flash column chromatography (2 to 15% MeOH gradient in DCM). Two resolved peaks with similar uv absorbance and similar retention on TLC (Rf 0.38 in 10% MeOH/DCM) were collected; the second of these peaks resulted from inadvertent cleavage of the TBDMS ether.

Step 2. The TBDMS cleavage was performed as described above with TBAF (0.10 ml, 1.0 M in THF, 2.0 equiv) in THF (1 ml) at ambient temperature for 1 h. The crude product was purified by flash column chromatography (2 to 15% MeOH gradient in DCM, Rf 0.38 in 10% MeOH/DCM), and these fractions were combined with those of the second peak from step 1.

Step 3. The Boc cleavage was performed as described above with TFA (0.25 ml) in DCM (1 ml) at 0 °C for 30 min. The crude product was purified by preparative reverse-phase HPLC (C18, water/acetonitrile gradient + 0.1% formic acid) to give the title compound in the form of a 1:1 formic acid salt (17.2 mg, 0.0286 mmol, 57% yield). $^1$H NMR (400 MHz, DMSO) δ 10.02 (s, 1H), 8.31 (s, 1H), 8.26 (s, 1H), 7.91 – 7.83 (m, 1H), 7.45 (d, $J$ = 8.9 Hz, 2H), 7.14 (d, $J$ = 9.0 Hz, 2H), 7.07 (d, $J$ = 8.7 Hz, 1H), 6.90 – 6.83 (m, 1H), 4.29 – 4.22 (m, 2H), 3.98 (s, 2H), 3.89 – 3.82 (m, 2H), 3.22 (q, $J$ = 6.3 Hz, 2H), 2.73 (t, $J$ = 6.3 Hz, 2H), 2.22 (s, 3H). $^{19}$F NMR (376 MHz, DMSO) δ -54.8. ESI$^+$ (LC/MS) calc'd 556.1569, found 556.1605. *Note: the -N$\underline{H}_3{}^+$ and -O$\underline{H}$ resonances were not located in the $^1$H NMR spectrum, and the -C$\underline{H}_2$N- resonance at 3.2 ppm is partially obscured by the broad $H_2O$ resonance.*

**_N_-(5-Chloro-4-hydroxy-2-methylphenyl)-1-(4-(2-(2-((2-aminoethyl)amino)-2-oxoethoxy)ethoxy)phenyl)-5-(trifluoromethyl)-1_H_-pyrazole-4-carboxamide (16).**

Step 1. The amide coupling was performed as described above with 1-(4-((2,2-dimethyl-4,9-dioxo-3,11-dioxa-5,8-diazatridecan-13-yl)oxy)phenyl)-5-(trifluoromethyl)-1*H*-pyrazole-4-carboxylic acid (25.8 mg, 0.0500 mmol), HATU (22.8 mg, 0.0600 mmol, 1.2 equiv), 4-((*tert*-butyldimethylsilyl)oxy)-5-chloro-2-methylaniline (20.4 mg, 0.0750 mmol, 1.5 equiv), DIPEA (17 µl, 0.10 mmol, 2.0 equiv), and DMF (0.25 ml). *Modification: the amine coupling partner was added after stirring the carboxylic acid, HATU, DMF, and DIPEA for 1 h at ambient temperature.* The crude product was purified by flash column chromatography (2 to 15% MeOH gradient in DCM). Two resolved peaks with similar uv absorbance and similar retention on TLC (Rf 0.38 in 10% MeOH/DCM) were collected; the second of these peaks resulted from inadvertent cleavage of the TBDMS ether.

Step 2. The TBDMS cleavage was performed as described above with TBAF (0.10 ml, 1.0 M in THF, 2.0 equiv) in THF (1 ml) at ambient temperature for 1 h. The crude product was purified by flash column chromatography (2 to 15% MeOH gradient in DCM, Rf 0.38 in 10% MeOH/DCM), and these fractions were combined with those of the second peak from step 1.

Step 3. The Boc cleavage was performed as described above with TFA (0.25 ml) in DCM (1 ml) at 0 °C for 30 min. The crude product was purified by preparative reverse-phase HPLC (C18, water/acetonitrile gradient + 0.1% formic acid) to give the title compound in the form of a 1:1 formic acid salt (17.4 mg, 0.0289 mmol, 58% yield). $^1$H NMR (400 MHz, DMSO) δ 9.88 (s, 1H), 8.29 (br, 1H), 8.24 (s, 1H), 7.84 (br, 1H), 7.45 (d, $J$ = 8.8 Hz, 2H), 7.30 (s, 1H), 7.14 (d, $J$ = 8.9 Hz, 2H), 6.86 (s, 1H), 4.29 – 4.22 (m, 2H), 3.98 (s, 2H), 3.89 – 3.82 (m, 2H), 3.24 – 3.19 (m, 2H), 2.77 – 2.69 (m, 2H), 2.14 (s, 3H). $^{19}$F NMR (376 MHz, DMSO) δ -54.8. ESI$^+$ (LC/MS) calc'd 556.1569, found

556.1605. *Note: the -NH$_3^+$ and -OH resonances were not located in the $^1$H NMR spectrum, and the -CH$_2$N- resonance at 3.2 ppm is partially obscured by the broad H$_2$O resonance.*

***N*-(3-Hydroxynapthalen-1-yl)-1-(4-(2-(2-((2-aminoethyl)amino)-2-oxoethoxy)ethoxy)phenyl)-5-(trifluoromethyl)-1*H*-pyrazole-4-carboxamide (17).**

Step 1. The amide coupling was performed as described above with 1-(4-((2,2-dimethyl-4,9-dioxo-3,11-dioxa-5,8-diazatridecan-13-yl)oxy)phenyl)-5-(trifluoromethyl)-1*H*-pyrazole-4-carboxylic acid (12.0 mg, 0.0232 mmol), HATU (10.6 mg, 0.0279 mmol, 1.2 equiv), 3-((*tert*-butyldimethylsilyl)oxy)naphthalen-1-amine (10.0 mg, 0.0366 mmol, 1.6 equiv), DIPEA (8.0 µl, 0.046 mmol, 2.0 equiv), and DMF (0.12 ml). *Modification: the amine coupling partner was added after stirring the carboxylic acid, HATU, DMF, and DIPEA for 1 h at ambient temperature.* The crude product was carried on to the next step without purification.

Step 2. The TBDMS cleavage was performed as described above with TBAF (0.050 ml, 1.0 M in THF, 2.2 equiv) in THF (0.5 ml) at ambient temperature for 1 h. The crude product was purified by flash column chromatography (2 to 15% MeOH gradient in DCM).

Step 3. The Boc cleavage was performed as described above with TFA (0.25 ml) in DCM (0.75 ml) at 0 °C for 1 h. The crude product was purified by preparative reverse-phase HPLC (C18, water/acetonitrile gradient + 0.1% TFA) to give the title compound in the form of a 1:1 TFA salt (12.3 mg, 0.0183mmol, 79% yield). $^1$H NMR (600 MHz, DMSO) δ 10.53 (s, 1H), 9.86 (br, 1H), 8.41 (br, 1H), 8.01 (t, *J* = 6.0 Hz, 1H), 7.95 (d, *J* = 8.4 Hz, 1H), 7.77 (br, 3H), 7.73 (d, *J* = 8.3 Hz, 1H), 7.49 (d, *J* = 8.4 Hz, 2H), 7.43 (ddd, *J* = 8.1, 6.7, 1.2 Hz, 1H), 7.39 (s, 1H), 7.31 (ddd, *J* = 8.2, 6.7, 1.3 Hz, 1H), 7.16 (d, *J* = 8.9 Hz, 2H), 7.07 (d, *J* = 2.4 Hz, 1H), 4.30 – 4.24 (m, 2H), 4.01 (s, 2H), 3.91 – 3.85 (m, 2H), 3.36 (q, *J* = 6.2 Hz, 2H), 2.90 (h, *J* = 6.0 Hz, 2H). $^{13}$C NMR (151 MHz, DMSO) δ 170.1, 159.9, 159.3, 154.7, 139.7, 135.0, 134.1, 131.8, 129.9 (q, *J* = 38.8 Hz), 127.5, 126.5, 126.4, 122.9, 122.8, 122.7, 121.1, 119.5 (q, *J* = 270.4 Hz), 115.1, 114.9, 107.3, 70.1, 69.4, 67.4, 38.8, 36.1. $^{19}$F NMR (376 MHz, DMSO) δ -54.8. ESI$^+$ (LC/MS) calc'd 558.1959, found 558.1974. *Note: the $^{13}$C and $^{19}$F NMR resonances of the F$_3$CCO$_2^-$ anion are not included in this list.*

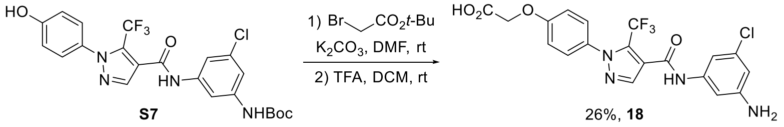

**2-(4-(4-((3-Amino-5-chlorophenyl)carbamoyl)-5-(trifluoromethyl)-1*H*-pyrazol-1-yl)phenoxy)acetic acid (18).**

Step 1. *tert*-Butyl (3-chloro-5-(1-(4-hydroxyphenyl)-5-(trifluoromethyl)-1*H*-pyrazole-4-carboxamido)phenyl)carbamate (49.7 mg, 0.100 mmol) was dissolved in dry DMF (0.4 ml) in an oven-dried 1 dram vial under Ar. Potassium carbonate (55 mg, 0.40 mmol, 4.0 equiv) was added, and the mixture was stirred at ambient temperature for 20 min. *tert*-Butyl bromoacetate (18 µl, 23 mg, 0.12 mmol, 1.2 equiv) was added with stirring, and the mixture was stirred at ambient temperature overnight. The mixture was diluted with ethyl acetate and water (ca. 2 ml each), and the layers were mixed and separated. The aqueous layer was extracted twice more with ethyl acetate (ca. 2 ml each), and the organic layers were filtered through Na$_2$SO$_4$. Volatiles were removed at reduced pressure, and the crude product was purified by flash column chromatography (hexanes/ethyl acetate gradient, Rf 0.85 in 50% EA/Hex). The resulting sample (57.3 mg) contained mono- and di-alkylated products in a 2:1 ratio.

Step 2. The sample obtained from step 1 was dissolved in DCM (1 ml) in a 20 ml scintillation vial under Ar. TFA (1 ml) was added with stirring, and the mixture was stirred at ambient temperature for 1 h. Volatiles were evaporated under a stream of Ar, and the crude product was purified by high pressure liquid chromatography (C18, water/acetonitrile gradient with 0.1% formic acid). Acetonitrile was removed at reduced pressure, and the resulting aqueous mixture was frozen and lyophilized to give the title compound as a white solid (11.9 mg, 0.0262 mmol, 26% yield).

$^1$H NMR (400 MHz, DMSO) δ 13.12 (br, 1H), 10.33 (s, 1H), 8.22 (s, 1H), 7.44 (d, *J* = 8.9 Hz, 2H), 7.09 (d, *J* = 8.9 Hz, 2H), 6.95 (t, *J* = 1.9 Hz, 1H), 6.89 (t, *J* = 1.9 Hz, 1H), 6.35 (t, *J* = 2.0 Hz, 1H), 5.50 (br, 2H), 4.78 (s, 2H). $^{19}$F NMR (376 MHz, DMSO) δ -55.0. ESI$^+$ (LC/MS) calc'd 455.0729, found 455.0724.

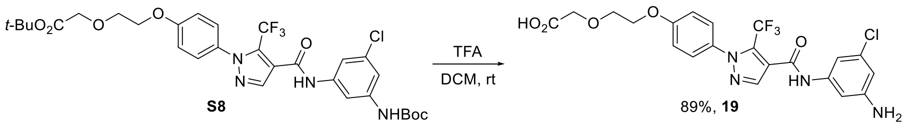

**2-(2-(4-(4-((3-Amino-5-chlorophenyl)carbamoyl)-5-(trifluoromethyl)-1*H*-pyrazol-1-yl)phenoxy)ethoxy)acetic acid (19).** *tert*-Butyl 2-(2-(4-(4-((3-((tert-butoxycarbonyl)amino)-5-chlorophenyl)carbamoyl)-5-(trifluoromethyl)-1*H*-pyrazol-1-yl)phenoxy)ethoxy)acetate (105 mg, 0.160 mmol) was dissolved in DCM (1 ml) in a 20 ml scintillation vial flushed with Ar, and the solution was cooled to 0 °C in an ice bath. TFA (1 ml) was added with stirring at 0 °C. After 15 min, the cooling bath was removed, and the solution was stirred at ambient temperature for 2 h. Volatiles were evaporated under a stream of Ar. The crude product was purified by high pressure liquid chromatography (C18, water/acetonitrile gradient with 0.1% formic acid). Acetonitrile was removed at reduced pressure, and the resulting aqueous mixture was frozen and lyophilized to give the title compound as a white solid (71.3 mg, 0.143 mmol, 89% yield).

$^{1}$H NMR (400 MHz, DMSO) δ 12.68 (br, 1H), 10.33 (s, 1H), 8.22 (s, 1H), 7.43 (d, *J* = 8.9 Hz, 2H), 7.12 (d, *J* = 9.0 Hz, 2H), 6.95 (t, *J* = 1.9 Hz, 1H), 6.89 (t, *J* = 1.9 Hz, 1H), 6.35 (t, *J* = 2.0 Hz, 1H), 5.50 (br, 2H), 4.24 – 4.16 (m, 2H), 4.10 (s, 2H), 3.88 – 3.82 (m, 2H). $^{19}$F NMR (376 MHz, DMSO) δ -55.0. ESI$^+$ (LC/MS) calc'd 499.0991, found 499.0994.

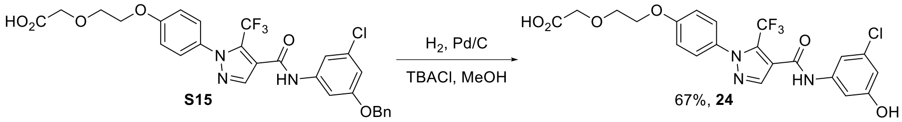

**2-(2-(4-(4-((3-Chloro-5-hydroxyphenyl)carbamoyl)-5-(trifluoromethyl)-1*H*-pyrazol-1-yl)phenoxy)ethoxy)acetic acid (24).** 2-(2-(4-(4-((3-(Benzyloxy)-5-chlorophenyl)carbamoyl)-5-(trifluoromethyl)-1*H*-pyrazol-1-yl)phenoxy)ethoxy)acetic acid (28.0 mg, 0.0475 mmol) and tetrabutylammonium chloride (15 mg, 0.054 mmol, 1.1 equiv) were suspended in dry methanol (1 ml) in a 20 ml scintillation vial under Ar. The mixture was cooled to 0 °C in an ice bath, and palladium on carbon (10% w/w, 2.0 mg, 0.0019 mmol, 4.0 mol%) was added. The atmosphere was purged with hydrogen (balloon), the cooling bath was removed, and the mixture was stirred at ambient temperature until LC/MS analysis indicated high conversion of the benzyl ether. The atmosphere was purged with Ar, and the mixture was filtered through a plug of Celite, which was rinsed thrice with methanol (1 ml each). Volatiles were removed at reduced pressure, and the crude product was purified by high pressure liquid chromatography (C18, water/acetonitrile gradient with 0.1% formic acid). Acetonitrile was removed at reduced pressure, and the resulting aqueous mixture was frozen and lyophilized to give the title compound as a white solid (16.0 mg, 0.032 mmol, 67% yield).

$^{1}$H NMR (400 MHz, DMSO) δ 12.64 (br, 1H), 10.53 (s, 1H), 9.99 (s, 1H), 8.25 (br, 1H), 7.44 (d, *J* = 8.9 Hz, 2H), 7.26 (t, *J* = 1.9 Hz, 1H), 7.19 (t, *J* = 2.0 Hz, 1H), 7.13 (d, *J* = 9.0 Hz, 2H), 6.56 (t, *J* = 2.1 Hz, 1H), 4.24 – 4.18 (m, 2H), 4.10 (s, 2H), 3.88 – 3.82 (m, 2H). $^{19}$F NMR (376 MHz, DMSO) δ -55.0. ESI$^+$ (LC/MS) calc'd 500.0831, found 500.0826.

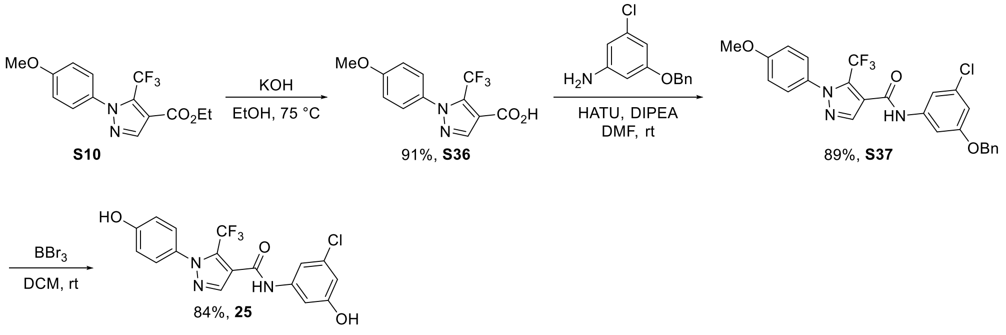

**1-(4-Methoxyphenyl)-5-(trifluoromethyl)-1*H*-pyrazole-4-carboxylic acid (S36).** Ethyl 1-(4-methoxyphenyl)-5-(trifluoromethyl)-1*H*-pyrazole-4-carboxylate (4.47 g, 14.2 mmol) was dissolved in absolute ethanol (35 ml). Potassium hydroxide (1.20 g, 21.4 mmol, 1.5 equiv) was added with stirring, and the reaction mixture was heated at a gentle reflux for 4 h. The mixture was allowed to cool to ambient temperature, and volatiles were removed at reduced pressure. The crude conjugate base was dissolved in water (ca. 100 ml) and transferred to a separatory

funnel. The solution was acidified by adding concentrated $HCl_{(aq)}$ (ca. 5 ml) and extracted thrice with ethyl acetate (ca. 80 ml each). Each organic layer was washed in sequence with brine (ca. 80 ml). The organic layers were dried over $Na_2SO_4$ and filtered. Volatiles were removed at reduced pressure. The crude acid was recrystallized by dissolving in boiling toluene (ca. 80 ml) then cooling to 4 °C. The solid was collected by filtration, washed with cold toluene and hexanes (ca. 20 ml each), and dried under vacuum. The filtrate was concentrated, and a second crop was obtained by a similar procedure with lower volumes (ca. 30 ml toluene). The two crops were combined to give the title compound as a white solid (3.72 g, 13.0 mmol, 91% yield).
$^1$H NMR (400 MHz, DMSO) δ 13.29 (s, 1H), 8.20 (s, 1H), 7.45 (d, *J* = 8.9 Hz, 2H), 7.09 (d, *J* = 8.9 Hz, 2H), 3.84 (s, 3H). $^{19}$F NMR (376 MHz, DMSO) δ -54.6. ESI$^+$ (LC/MS) calc'd 287.0638, found 287.0643.

***N*-(3-(Benzyloxy)-5-chlorophenyl)-1-(4-methoxyphenyl)-5-(trifluoromethyl)-1*H*-pyrazole-4-carboxamide (S37).** 1-(4-Methoxyphenyl)-5-(trifluoromethyl)-1*H*-pyrazole-4-carboxylic acid (57.2 mg, 0.200 mmol), 3-benzyloxy-5-chloroaniline (56.0 mg, 0.240 mmol, 1.20 mmol), and HATU (114 mg, 0.300 mmol, 1.5 equiv) were added to an oven-dried 1 dram vial under Ar, and the mixture was cooled to 0 °C in an ice bath. DMF (0.8 ml) and DIPEA(0.11 ml, 0.63 mmol, 3.2 equiv) were added in sequence with stirring. The mixture was stirred for 10 min, then the cooling bath was removed, and the mixture was stirred at ambient temperature overnight. The mixture was diluted with ethyl acetate and saturated $NH_4Cl_{(aq)}$ (ca. 1.5 ml each), and the layers were mixed and separated. The organic layer was washed further with brine (ca. 1.5 ml). The aqueous layers were extracted twice more in the same sequence with ethyl acetate (ca. 1.5 ml each). The organic layers were filtered through $Na_2SO_4$, and volatiles were removed at reduced pressure. The crude product was purified by flash column chromatography (hexanes/ethyl acetate gradient, Rf 0.67 in 50% EA/Hex) to give the title compound as an off-white solid (89.7 mg, 0.179 mmol, 89% yield).
$^1$H NMR (400 MHz, DMSO) δ 10.63 (s, 1H), 8.26 (s, 1H), 7.50 – 7.31 (m, 9H), 7.12 (d, *J* = 9.0 Hz, 2H), 6.90 (t, *J* = 2.1 Hz, 1H), 5.13 (s, 2H), 3.85 (s, 3H). $^{19}$F NMR (376 MHz, DMSO) δ -55.0. ESI$^+$ (LC/MS) calc'd 502.1140, found 502.1126.

***N*-(3-Chloro-5-hydroxyphenyl)-1-(4-hydroxyphenyl)-5-(trifluoromethyl)-1*H*-pyrazole-4-carboxamide (25).** *N*-(3-(Benzyloxy)-5-chlorophenyl)-1-(4-methoxyphenyl)-5-(trifluoromethyl)-1*H*-pyrazole-4-carboxamide (88.3 mg, 0.176 mmol) was dissolved in dry DCM in a 20 ml scintillation vial under Ar. A solution of boron tribromide (10% v/v, 0.68 ml, 0.71 mmol, 4.0 equiv) was added slowly with stirring, and the mixture was stirred at ambient temperature overnight (16 h). The reaction was quenched by slowly adding absolute ethanol (0.7 ml) and stirring at ambient temperature for 15 min. Volatiles were evaporated under a stream of Ar. The crude product was purified by flash column chromatography (hexanes/ethyl acetate gradient) to give the title compound as a white solid (57.4 mg, 0.144 mmol, 82% yield). A small portion of this sample (18 mg) was further purified by high pressure liquid chromatography (C18, water/acetonitrile gradient with 0.1% formic acid) for use in biochemical experiments.
$^1$H NMR (400 MHz, DMSO) δ 10.50 (s, 1H), 10.08 (s, 1H), 9.97 (s, 1H), 8.21 (s, 1H), 7.30 (d, *J* = 8.8 Hz, 2H), 7.25 (t, *J* = 1.9 Hz, 1H), 7.19 (t, *J* = 2.0 Hz, 1H), 6.91 (d, *J* = 8.8 Hz, 2H), 6.56 (t, *J* = 2.1 Hz, 1H). $^{19}$F NMR (376 MHz, DMSO) δ -55.1. ESI$^+$ (LC/MS) calc'd 398.0514, found 398.0482.

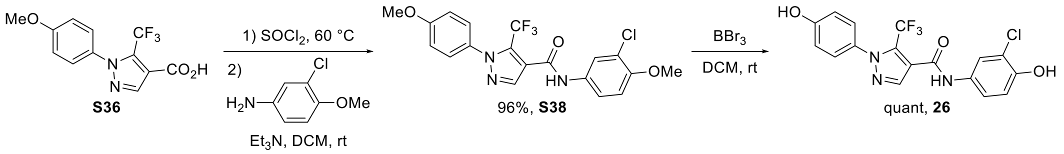

***N*-(3-Chloro-4-methoxy-phenyl)-1-(4-methoxyphenyl)-5-(trifluoromethyl)-1*H*-pyrazole-4-carboxamide (S38).** 1-(4-Methoxyphenyl)-5-(trifluoromethyl)-1H-pyrazole-4-carboxylic acid (57.2 mg, 0.200 mmol) and thionyl chloride (0.20 ml, 0.33 g, 2.8 mmol, 14 equiv) were added to an oven-dried 1-dram vial under Ar. The mixture was heated with stirring at 60 °C for 30 min, then cooled to ambient temperature. Volatiles were evaporated under a stream of Ar. The residue was redissolved in dry DCM (0.5 ml), then volatiles were again evaporated by the same procedure. The oily yellow residue was redissolved in dry DCM (0.5 ml), then 3-chloro-4-methoxy-aniline (47.3 mg, 0.300 mmol. 1.5 equiv) and triethylamine (56 µl, 40 mg, 0.40 mmol, 2.0 equiv) were added with stirring. An additional portion of dry DCM (0.5 ml) was added to enable stirring of the resulting thick slurry. The mixture was

stirred at ambient temperature overnight, then loaded directly onto a plug of silica and purified by flash column chromatography to give the title compound as a white solid (81.7 mg, 0.192 mmol, 96% yield).
$^1$H NMR (400 MHz, $CDCl_3$) δ 7.98 (s, 1H), 7.66 (s, 1H), 7.48 (d, *J* = 8.8 Hz, 1H), 7.41 (s, 1H), 7.36 (d, *J* = 8.8 Hz, 2H), 7.00 (d, *J* = 9.0 Hz, 2H), 6.93 (d, *J* = 8.9 Hz, 1H), 3.91 (s, 3H), 3.88 (s, 3H). $^{19}$F NMR (376 MHz, DMSO) δ -55.2. ESI$^+$ (LC/MS) calc'd 426.0827, found 426.0855.

***N*-(3-Chloro-4-hydroxyphenyl)-1-(4-hydroxyphenyl)-5-(trifluoromethyl)-1*H*-pyrazole-4-carboxamide (26).** *N*-(3-Chloro-4-methoxy-phenyl)-1-(4-methoxyphenyl)-5-(trifluoromethyl)-1*H*-pyrazole-4-carboxamide (42.6 mg, 0.100 mmol) was dissolved in dry DCM (2 ml) in a 20 ml scintillation vial flushed with Ar. The solution was cooled to 0 °C in an ice bath, and BBr3 (0.1 ml, 1 mmol, 10 equiv) was added with stirring. The cooling bath was removed, and the mixture was stirred at ambient temperature for 2h. At this point, TLC (50% EA/Hex) indicated that the starting material had been partially converted to three products with lower Rf values. An additional portion of $BBr_3$ (0.2 ml, 2 mmol, 20 equiv) was added, and the mixture was stirred at ambient temperature overnight. At this point, TLC indicated that the starting material had been fully converted and with high selectivity for a single product. The mixture was cooled to 0 °C in an ice bath, and the reaction was quenched by adding absolute ethanol with stirring (ca. 1 ml). Volatiles were evaporated under a stream of Ar, and the residue was suspended in water (ca. 5 ml). This mixture was extracted twice with ethyl acetate (ca. 2 ml each), and the organic layers were filtered through $Na_2SO_4$. Volatiles were removed at reduced pressure, and the crude product with purified by flash column chromatography (2 to 15% MeOH in DCM, Rf 0.43 in 10% MeOH/DCM) to give the title compound as a white solid (39.6 mg, 0.100 mmol, 99.5% yield).
$^1$H NMR (600 MHz, DMSO) δ 10.38 (s, 1H), 10.09 (s, 1H), 10.03 (s, 1H), 8.19 (s, 1H), 7.77 (d, *J* = 2.5 Hz, 1H), 7.39 (dd, *J* = 8.8, 2.5 Hz, 1H), 7.30 (d, *J* = 8.7 Hz, 2H), 6.95 (d, *J* = 8.8 Hz, 1H), 6.91 (d, *J* = 8.8 Hz, 2H). $^{19}$F NMR (376 MHz, DMSO) δ -55.0. ESI$^+$ (LC/MS) calc'd 398.0514, found 398.0510.

**Chemical abbreviations**
DCM – dichloromethane
THF – tetrahydrofuran
DMF – *N*,*N*-dimethylformamide
DMSO – dimethyl sulfoxide
Tol - toluene
Hex – hexanes (mixture of isomers)
EA – ethyl acetate
TBACl – tetra-*n*-butylammonium chloride
TBAF – tetra-*n*-butylammonium fluoride
TFA – trifluoroacetic acid
DIPEA – di-*iso*-propylethylamine
HATU – 1-[bis(dimethylamino)methylene]-1*H*-1,2,3-triazolo[4,5-*b*]pyridinium 3-oxide hexafluorophosphate
$Pd_2dba_3$ – tris(dibenzylacetone)dipalladium(0)
BINAP – (*rac*)-2,2′-bis(diphenylphosphino)-1,1′-binaphthyl
$[Ir(cod)OMe]_2$ – (1,5-cyclooctadiene)(methoxy)iridium(I) dimer
dtbpy – di-*tert*-butyl-2,2'-bipyridyl
Boc – *tert*-butyloxycarbonyl
TBDMS – *tert*-butyldimethylsilyl
Ts – *para*-toluenesulfonyl

*References*

1. Ostrem, J. M.; Peters, U.; Sos, M. L.; Wells, J. A.; Shokat, K. M., K-Ras(G12C) inhibitors allosterically control GTP affinity and effector interactions. *Nature* **2013,** *503* (7477), 548-551.
2. Ahmadian, M. R.; Zor, T.; Vogt, D.; Kabsch, W.; Selinger, Z.; Wittinghofer, A.; Scheffzek, K., Guanosine triphosphatase stimulation of oncogenic Ras mutants. *Proceedings of the National Academy of Sciences* **1999,** *96* (12), 7065-7070.
3. John, J.; Sohmen, R.; Feuerstein, J.; Linke, R.; Wittinghofer, A.; Goody, R. S., Kinetics of interaction of nucleotides with nucleotide-free H-ras p21. *Biochemistry* **1990,** *29* (25), 6058-6065.
4. Maurer, T.; Garrenton, L. S.; Oh, A.; Pitts, K.; Anderson, D. J.; Skelton, N. J.; Fauber, B. P.; Pan, B.; Malek, S.; Stokoe, D.; Ludlam, M. J. C.; Bowman, K. K.; Wu, J.; Giannetti, A. M.; Starovasnik, M. A.; Mellman, I.; Jackson, P. K.; Rudolph, J.; Wang, W.; Fang, G., Small-molecule ligands bind to a distinct pocket in Ras and inhibit SOS-mediated nucleotide exchange activity. *Proceedings of the National Academy of Sciences* **2012,** *109* (14), 5299-5304.
5. Skinner, S. P.; Fogh, R. H.; Boucher, W.; Ragan, T. J.; Mureddu, L. G.; Vuister, G. W., CcpNmr AnalysisAssign: a flexible platform for integrated NMR analysis. *Journal of Biomolecular NMR* **2016,** *66* (2), 111-124.
6. Pálfy, G.; Vida, I.; Perczel, A., 1H, 15N backbone assignment and comparative analysis of the wild type and G12C, G12D, G12V mutants of K-Ras bound to GDP at physiological pH. *Biomolecular NMR Assignments* **2020,** *14* (1), 1-7.
7. Dalvit, C.; Fagerness, P. E.; Hadden, D. T. A.; Sarver, R. W.; Stockman, B. J., Fluorine-NMR Experiments for High-Throughput Screening: Theoretical Aspects, Practical Considerations, and Range of Applicability. *Journal of the American Chemical Society* **2003,** *125* (25), 7696-7703.
8. Lu, Z.; Chen, Y.-H.; Smith, C.; Li, H.; Thompson, C. F.; Sweis, R.; Sinclair, P.; Kallashi, F.; Hunt, J.; Adamson, S. E.; Dong, G.; Ondeyka, D. L.; Qian, X.; Sun, W.; Vachal, P.; Zhao, K. Cyclic amine substituted oxazolidinone cetp inhibitor. WO2012058187, 2012.

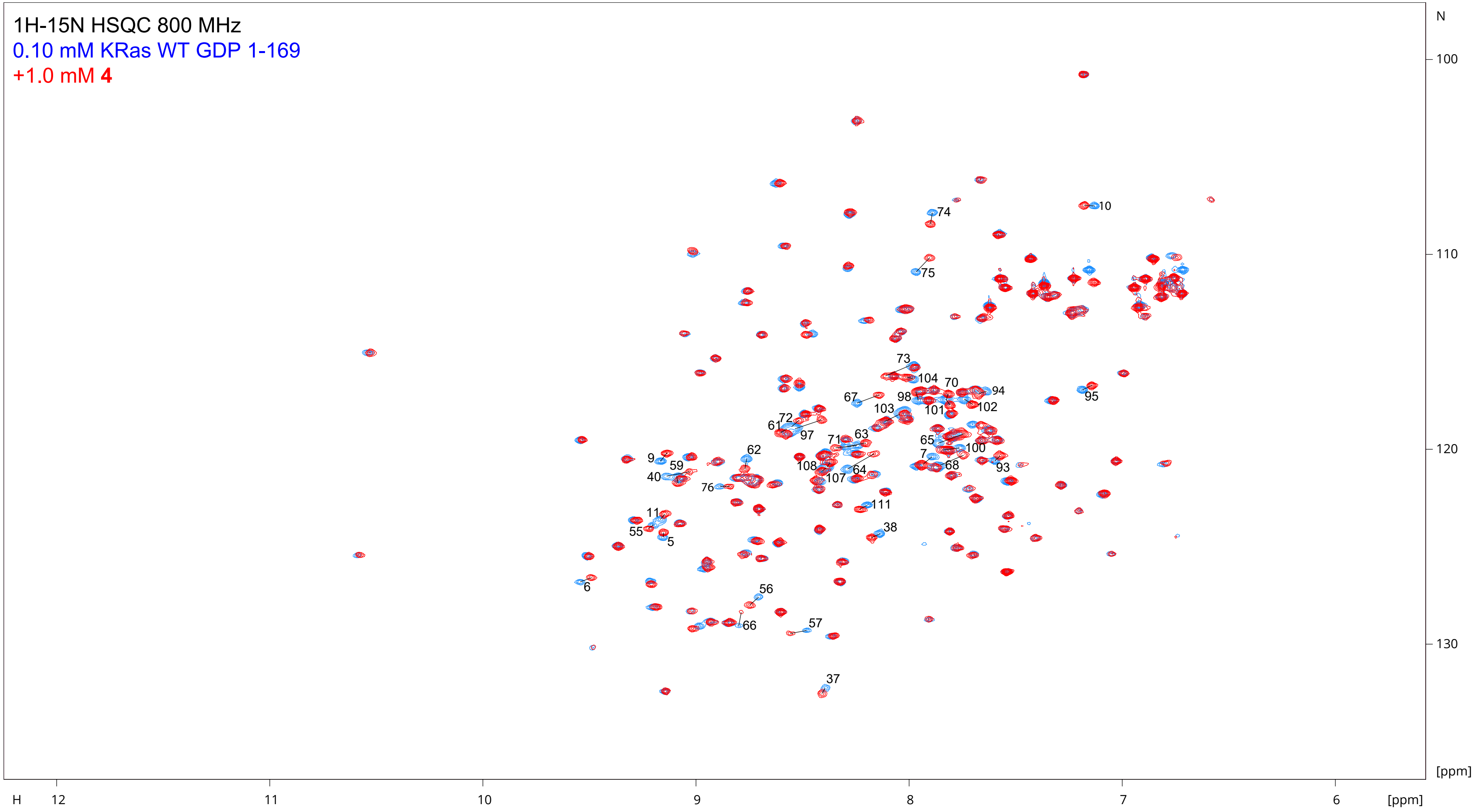

The following pairs of peaks overlap in the spectrum of unbound protein, and their CSPs should be interpreted with caution:
40/59, 68/22, 70/101, 72/97, 103/112, and 107/108

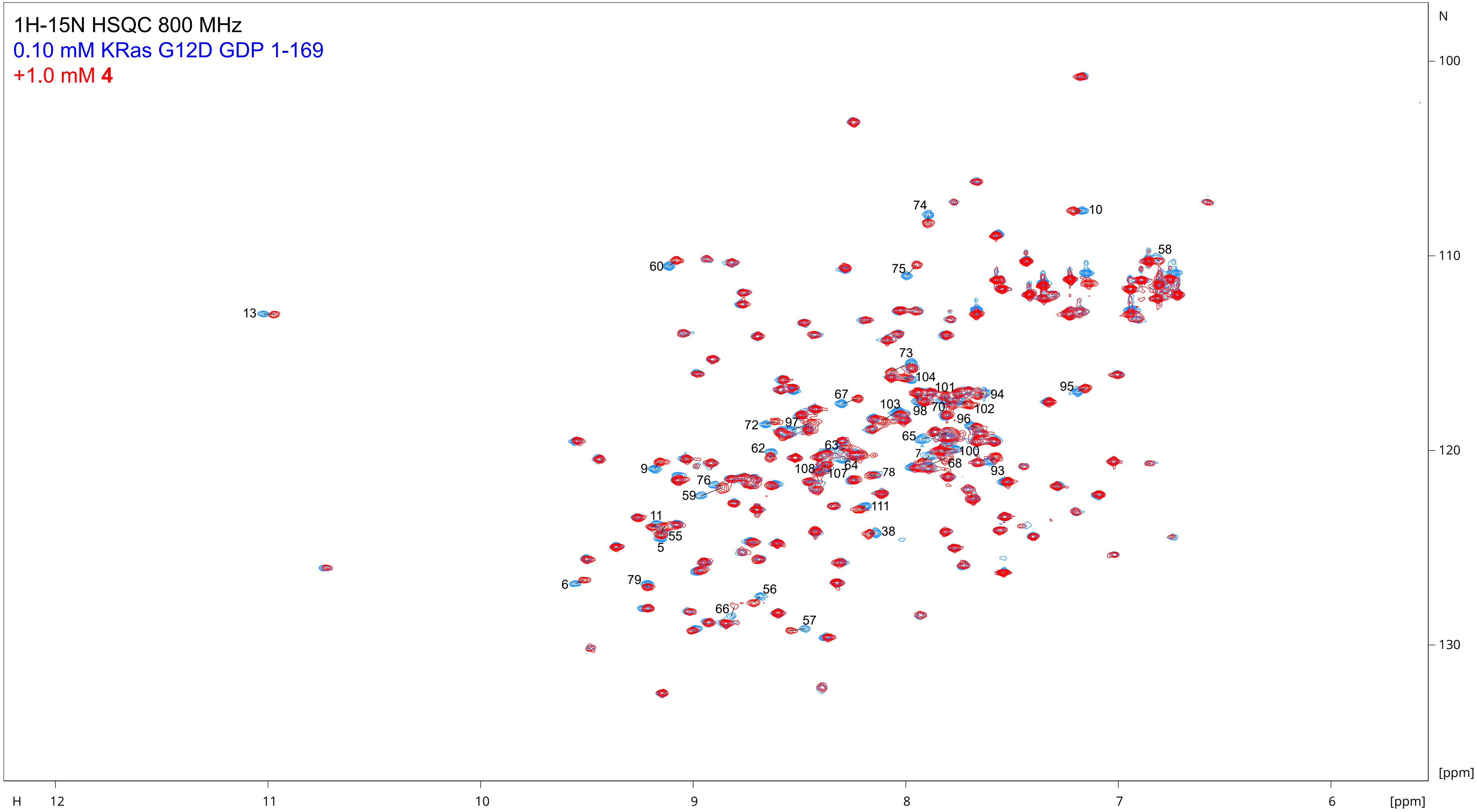

The following pairs of peaks overlap in the spectrum of unbound protein, and their CSPs should be interpreted with caution: 68/22, 70/101, 97/164, 98/136, 103/112, and 107/108

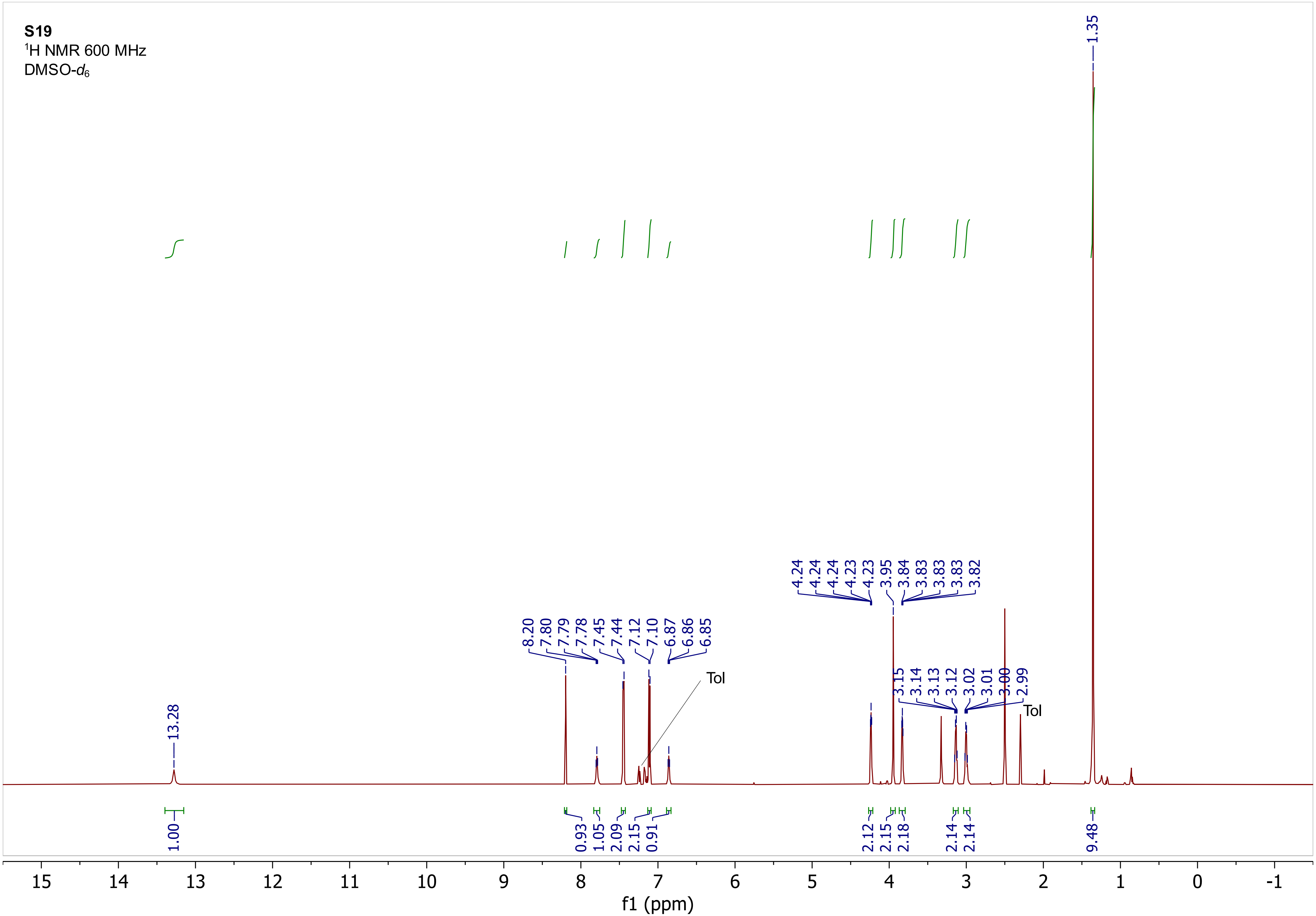

S19
$^{1}$H NMR 600 MHz
DMSO-$d_6$
13.28
8.20
7.80
7.79
7.78
7.45
7.44
7.12
7.10
6.87
6.86
6.85
Tol
4.24
4.24
4.24
4.23
4.23
3.95
3.84
3.83
3.83
3.83
3.82
3.15
3.14
3.13
3.12
3.02
3.01
3.00
2.99
Tol
1.35
1.00
0.93
1.05
2.09
2.15
0.91
2.12
2.15
2.18
2.14
2.14
9.48
15
14
13
12
11
10
9
8
7
6
5
4
3
2
1
0
-1
f1 (ppm)

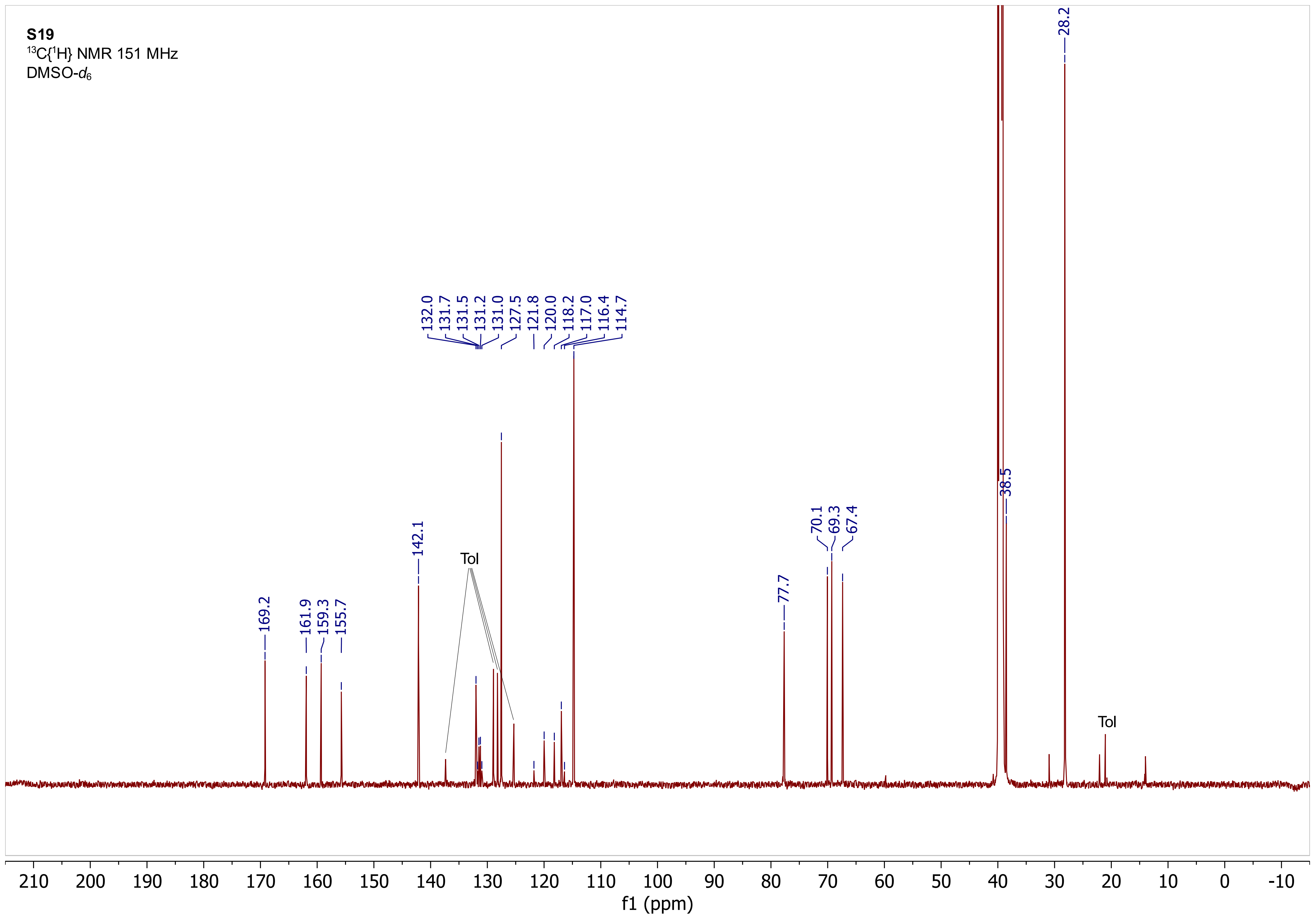
S19
$^{13}C\{^1H\}$ NMR 151 MHz
DMSO-$d_6$
169.2
161.9
159.3
155.7
142.1
132.0
131.7
131.5
131.2
131.0
127.5
121.8
120.0
118.2
117.0
116.4
114.7
Tol
77.7
70.1
69.3
67.4
38.5
28.2
Tol
f1 (ppm)

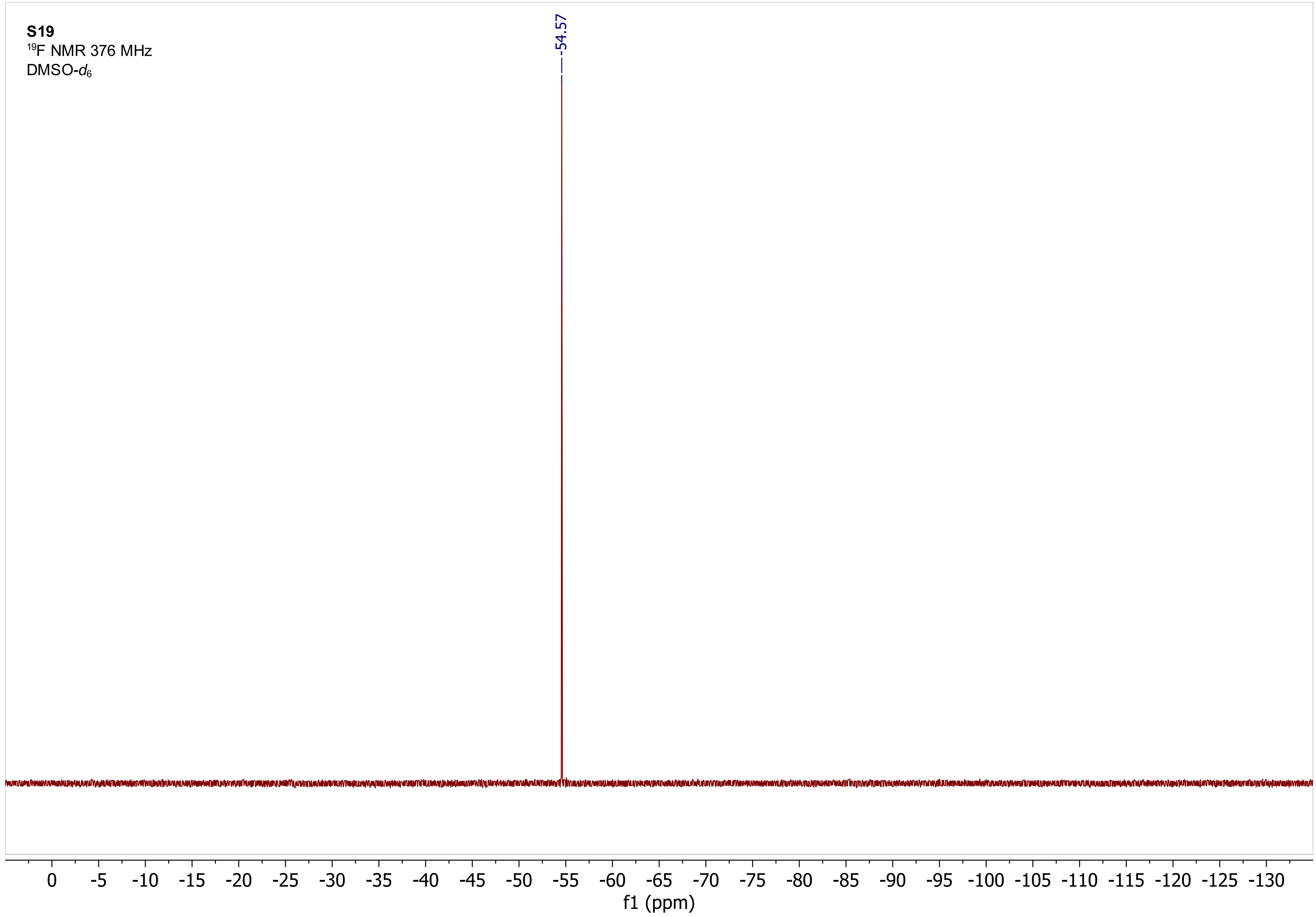

S19
19F NMR 376 MHz
DMSO-d6
-54.57
0 -5 -10 -15 -20 -25 -30 -35 -40 -45 -50 -55 -60 -65 -70 -75 -80 -85 -90 -95 -100 -105 -110 -115 -120 -125 -130
f1 (ppm)

**S19**
$^{1}$H COSY 600 MHz
DMSO-$d_6$

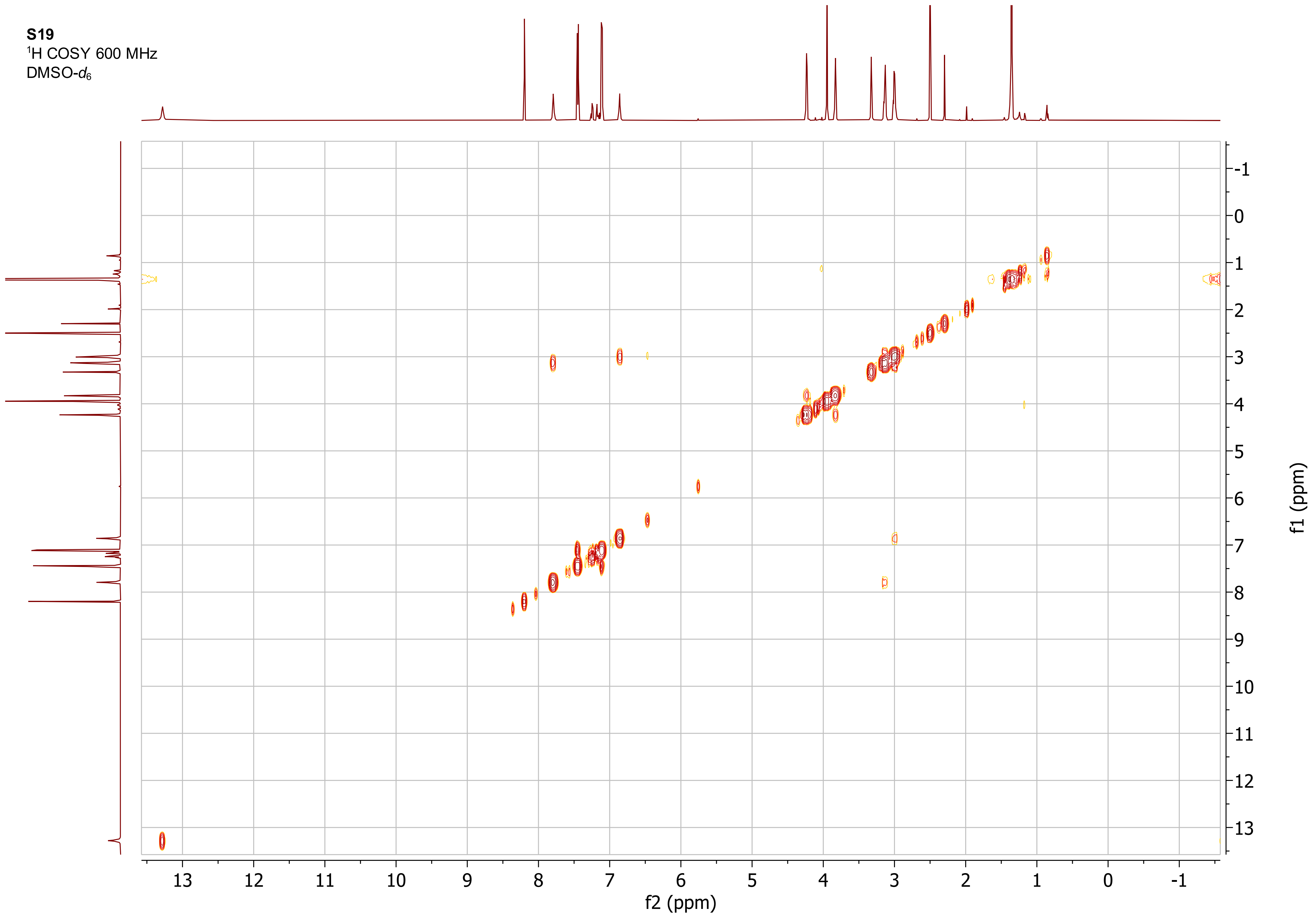

**S19**
$^{1}H$-$^{13}C$ ed-HSQC 600 MHz
DMSO-$d_6$

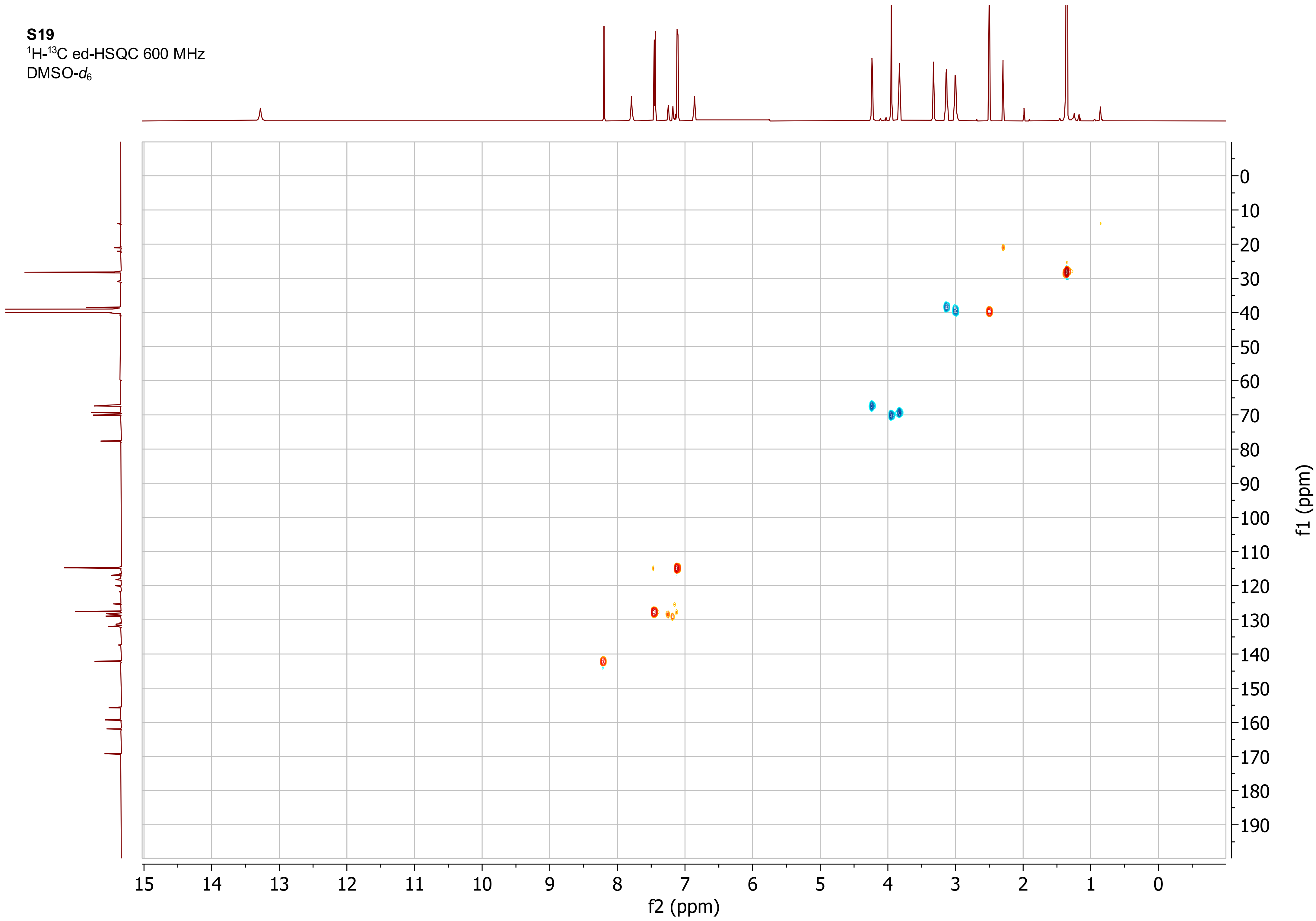

**S22**
$^{1}$H NMR 400 MHz
DMSO-$d_6$

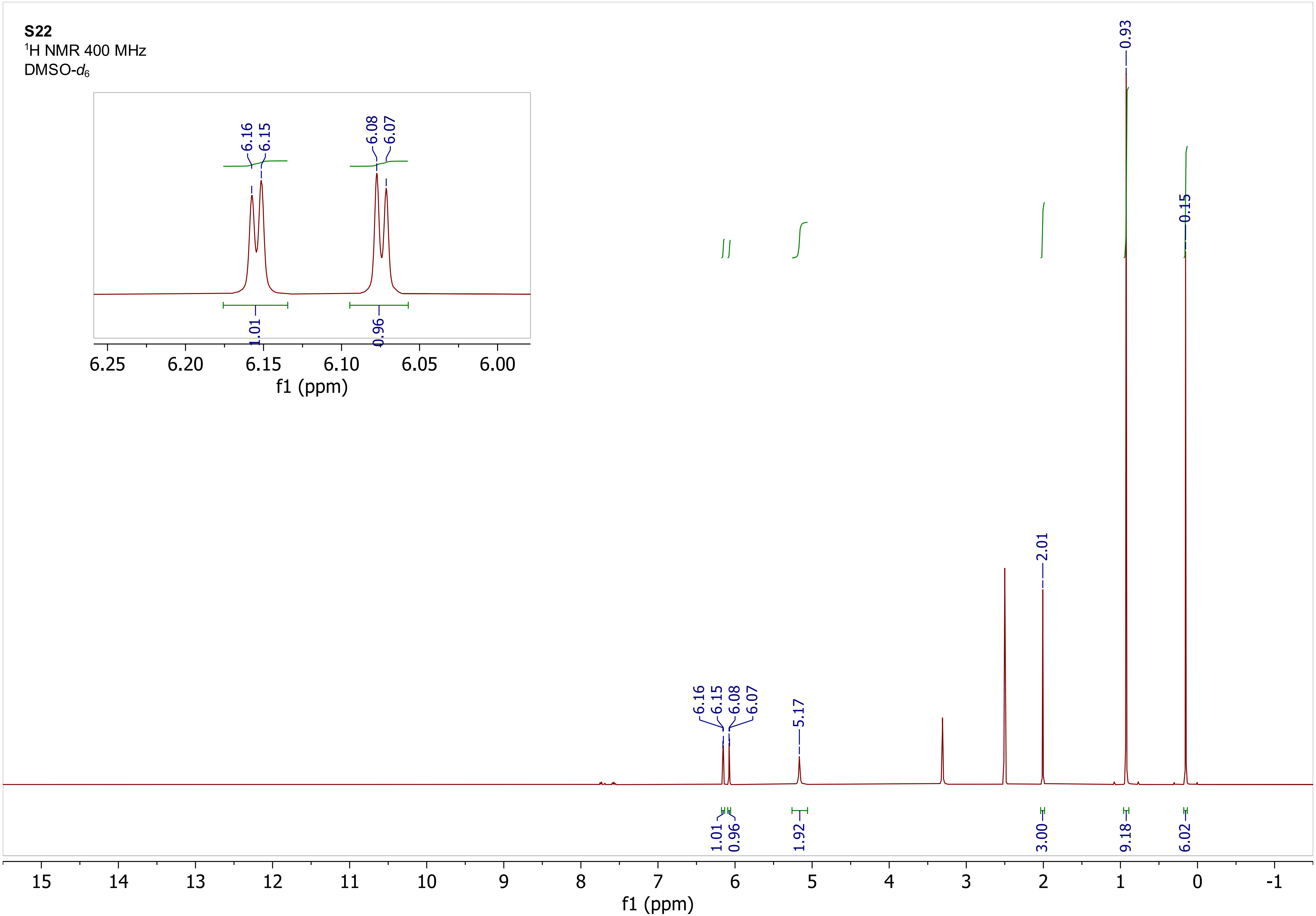

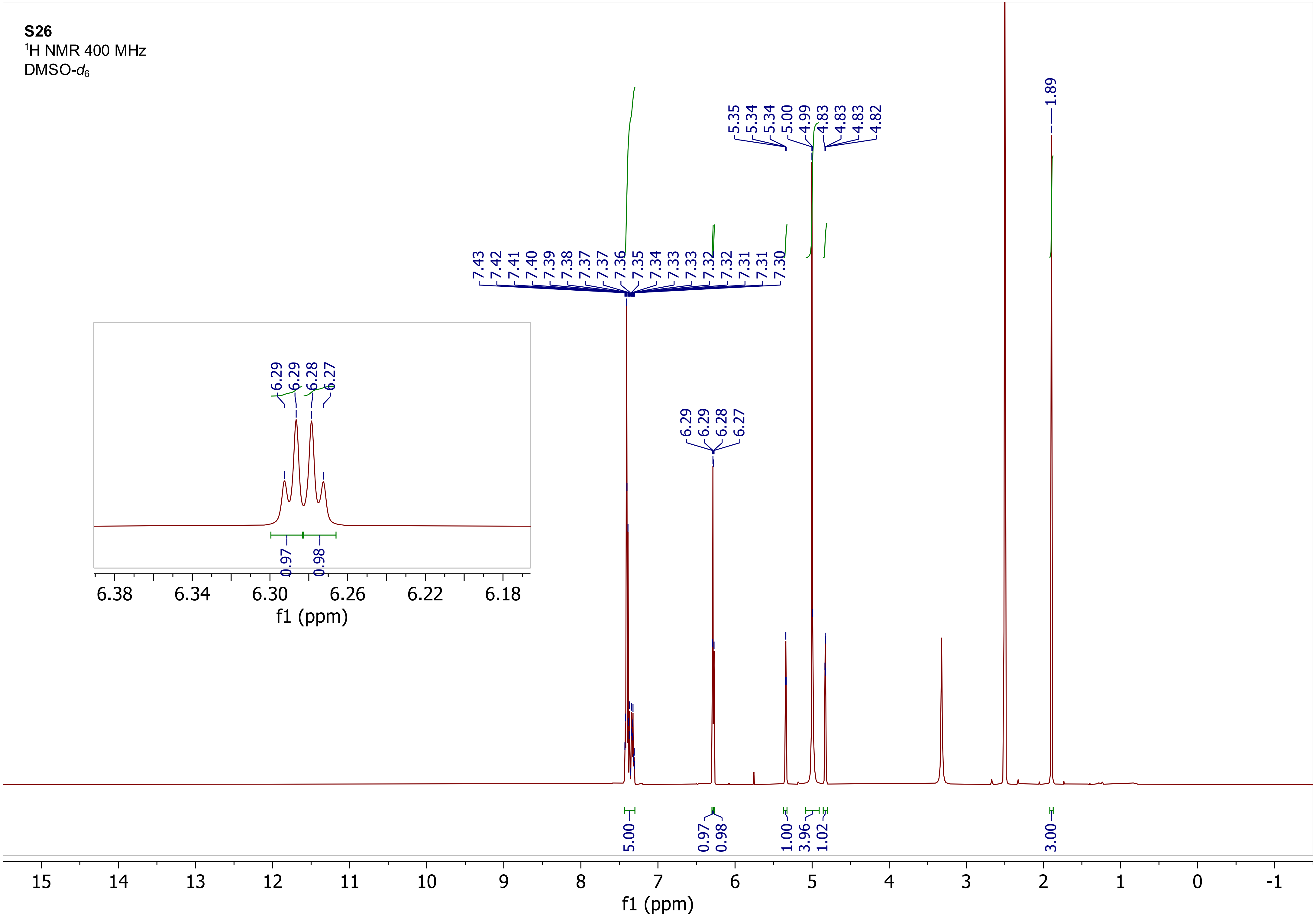

S26
$^1$H NMR 400 MHz
DMSO-$d_6$
7.43 7.42 7.41 7.40 7.39 7.38 7.37 7.37 7.36 7.35 7.34 7.33 7.33 7.32 7.32 7.31 7.31 7.30
6.29 6.29 6.28 6.27
5.35 5.34 5.34 5.00 4.99 4.83 4.83 4.83 4.82
1.89
5.00
0.97
0.98
1.00
3.96
1.02
3.00
0.97
0.98
6.38
6.34
6.30
6.26
6.22
6.18
f1 (ppm)
15
14
13
12
11
10
9
8
7
6
5
4
3
2
1
0
-1
f1 (ppm)

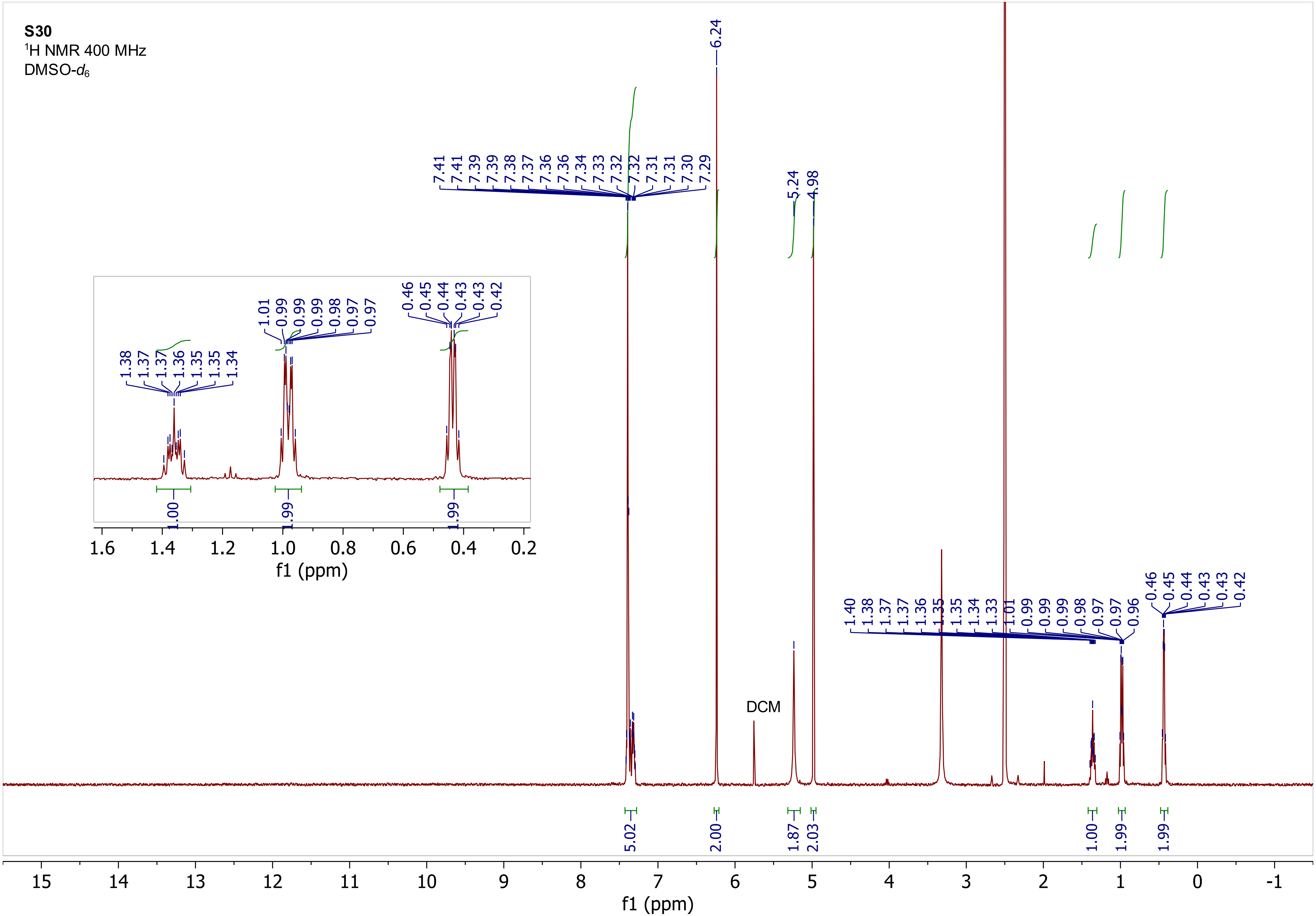

S30
$^{1}$H NMR 400 MHz
DMSO-$d_6$
DCM
f1 (ppm)

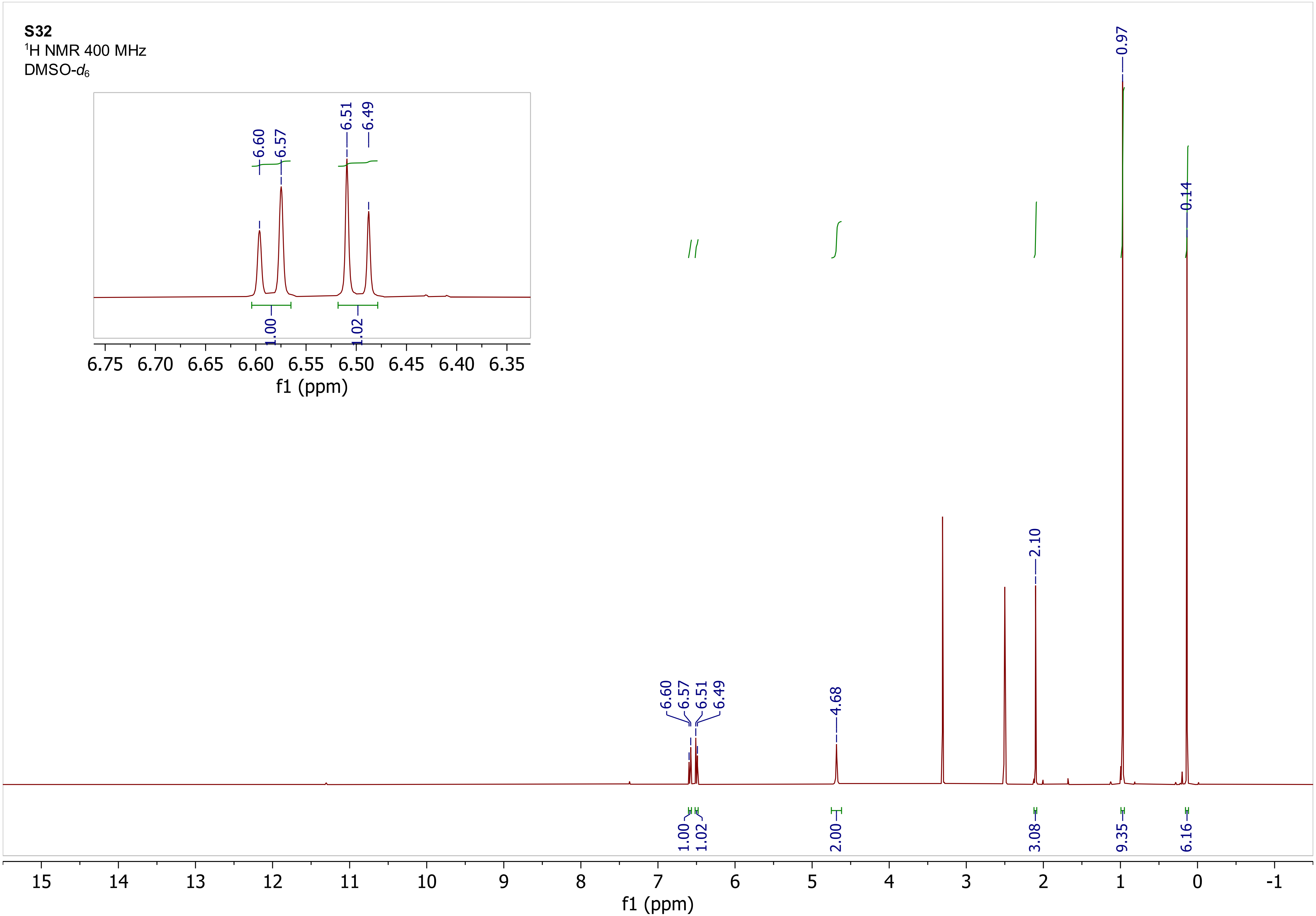

S32
$^1$H NMR 400 MHz
DMSO-$d_6$
6.60
6.57
6.51
6.49
1.00
1.02
6.75 6.70 6.65 6.60 6.55 6.50 6.45 6.40 6.35
f1 (ppm)
4.68
2.10
0.97
0.14
2.00
3.08
9.35
6.16
15 14 13 12 11 10 9 8 7 6 5 4 3 2 1 0 -1
f1 (ppm)

**S34**
$^{1}$H NMR 400 MHz
DMSO-$d_6$

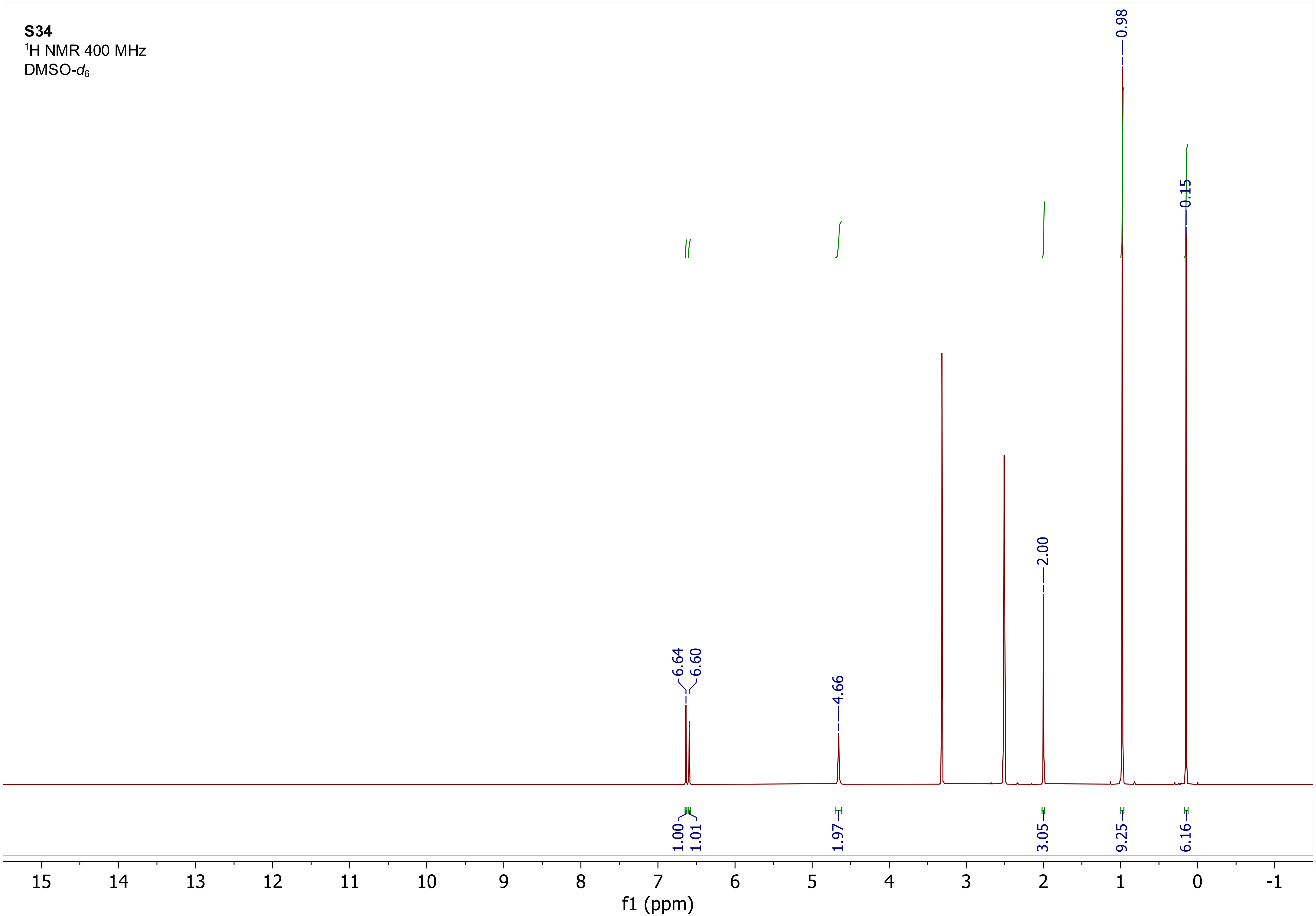

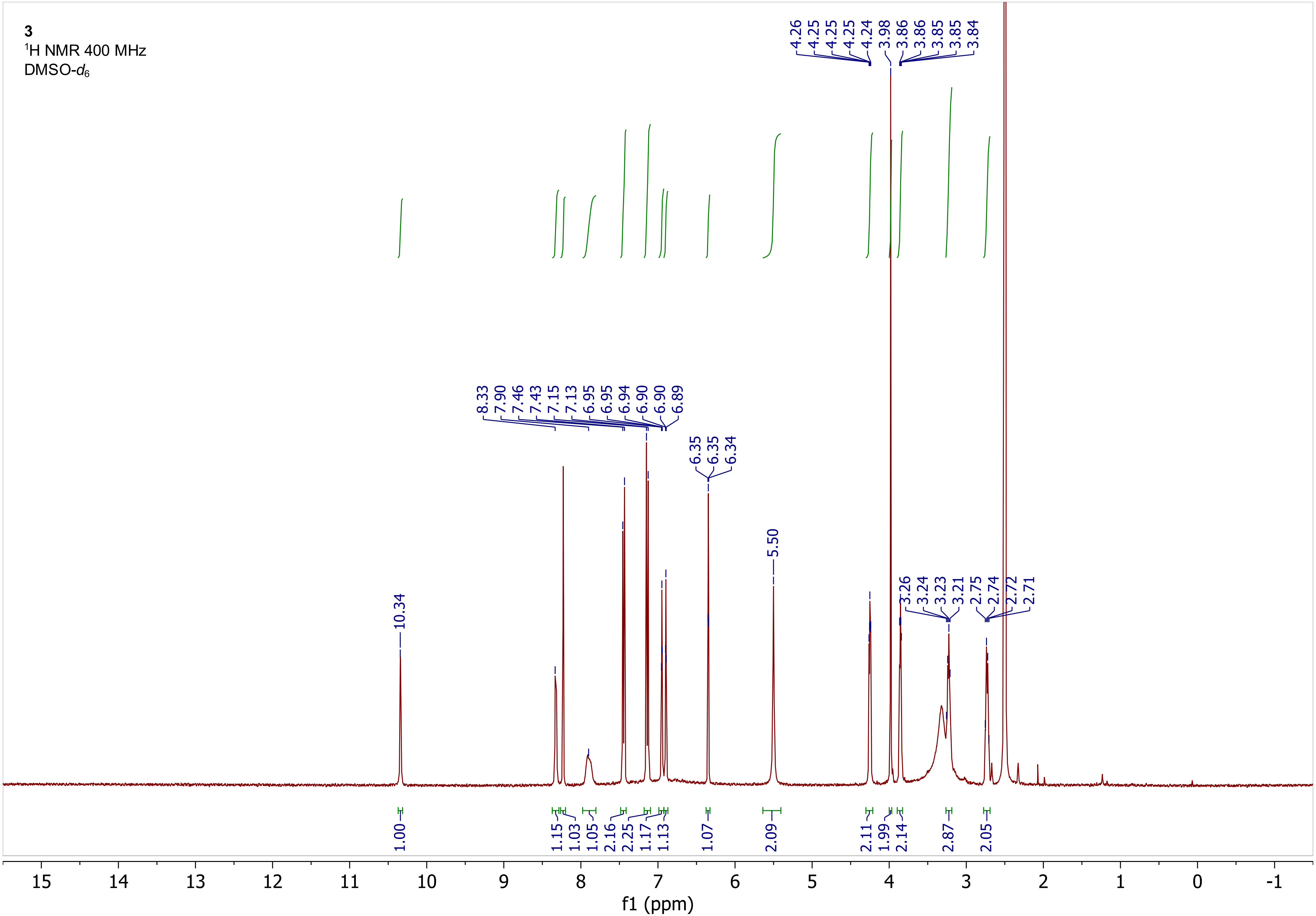
3
$^1$H NMR 400 MHz
DMSO-$d_6$
4.26 4.25 4.25 4.25 4.24 3.98 3.86 3.86 3.85 3.85 3.84
8.33 7.90 7.46 7.43 7.15 7.13 6.95 6.95 6.94 6.90 6.90 6.89
6.35 6.35 6.34
5.50
10.34
3.26 3.24 3.23 3.21 2.75 2.74 2.72 2.71
1.00 1.15 1.03 1.05 2.16 2.25 1.17 1.13 1.07 2.09 2.11 1.99 2.14 2.87 2.05
15 14 13 12 11 10 9 8 7 6 5 4 3 2 1 0 -1
f1 (ppm)

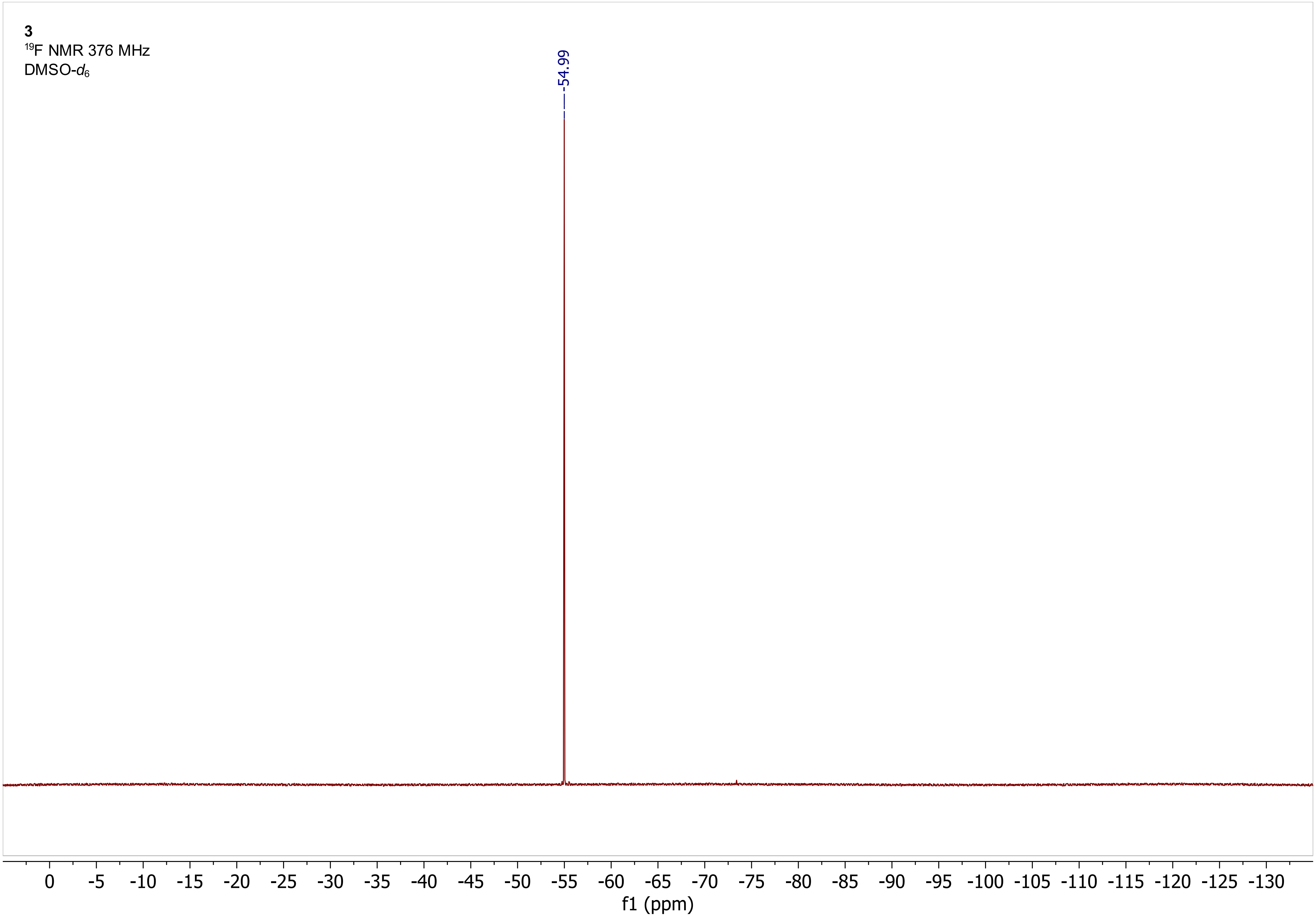
3
$^{19}$F NMR 376 MHz
DMSO-$d_6$
-54.99
0 -5 -10 -15 -20 -25 -30 -35 -40 -45 -50 -55 -60 -65 -70 -75 -80 -85 -90 -95 -100 -105 -110 -115 -120 -125 -130
f1 (ppm)

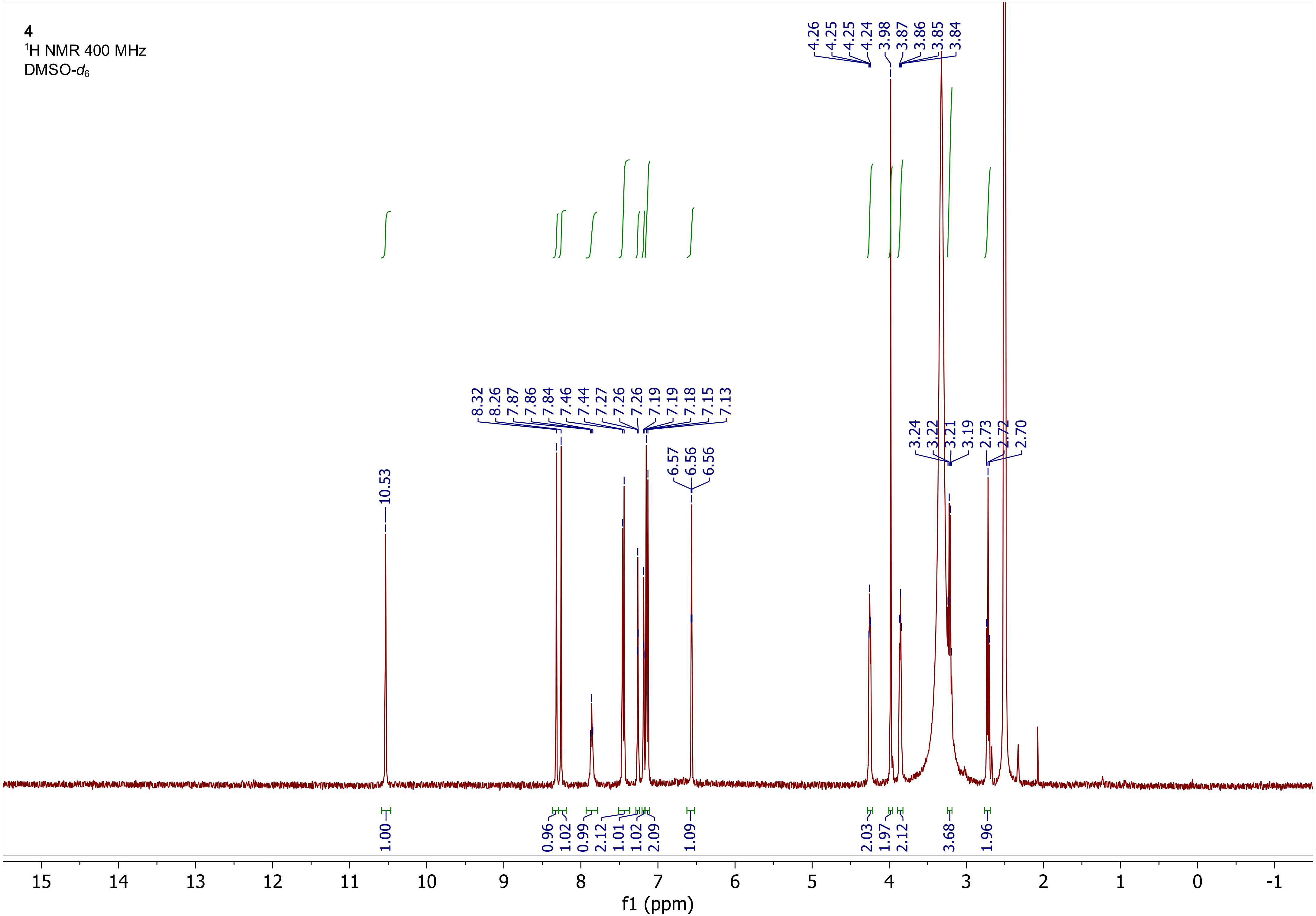
4
1H NMR 400 MHz
DMSO-d6
10.53
8.32 8.26 7.87 7.86 7.84 7.46 7.44 7.27 7.26 7.26 7.19 7.19 7.18 7.15 7.13
6.57 6.56 6.56
4.26 4.25 4.25 4.24 3.98 3.87 3.86 3.85 3.84
3.24 3.22 3.21 3.19 2.73 2.72 2.70
1.00 0.96 1.02 0.99 2.12 1.01 1.02 2.09 1.09 2.03 1.97 2.12 3.68 1.96
15 14 13 12 11 10 9 8 7 6 5 4 3 2 1 0 -1
f1 (ppm)

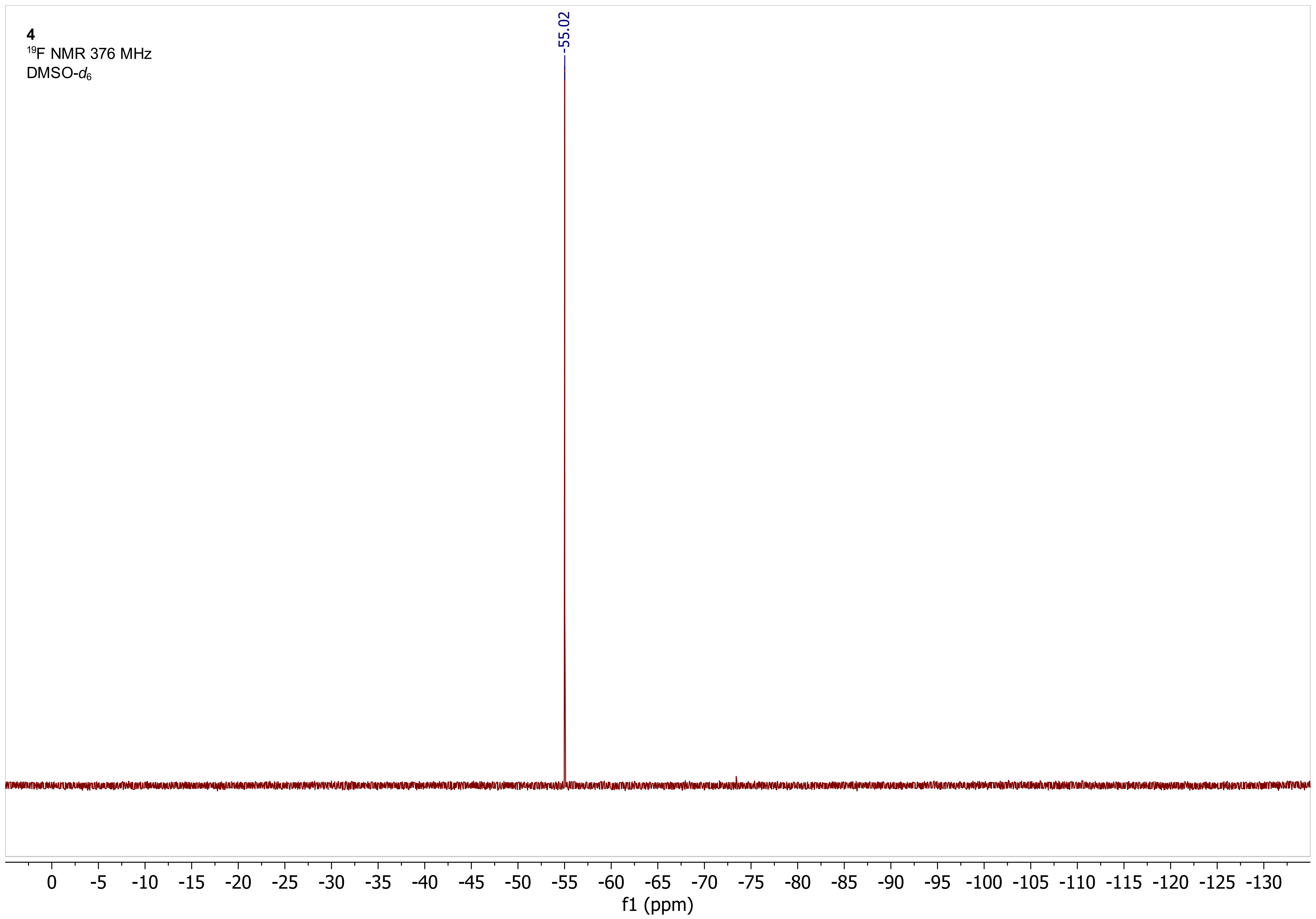

4
19F NMR 376 MHz
DMSO-d6
-55.02
0 -5 -10 -15 -20 -25 -30 -35 -40 -45 -50 -55 -60 -65 -70 -75 -80 -85 -90 -95 -100 -105 -110 -115 -120 -125 -130
f1 (ppm)

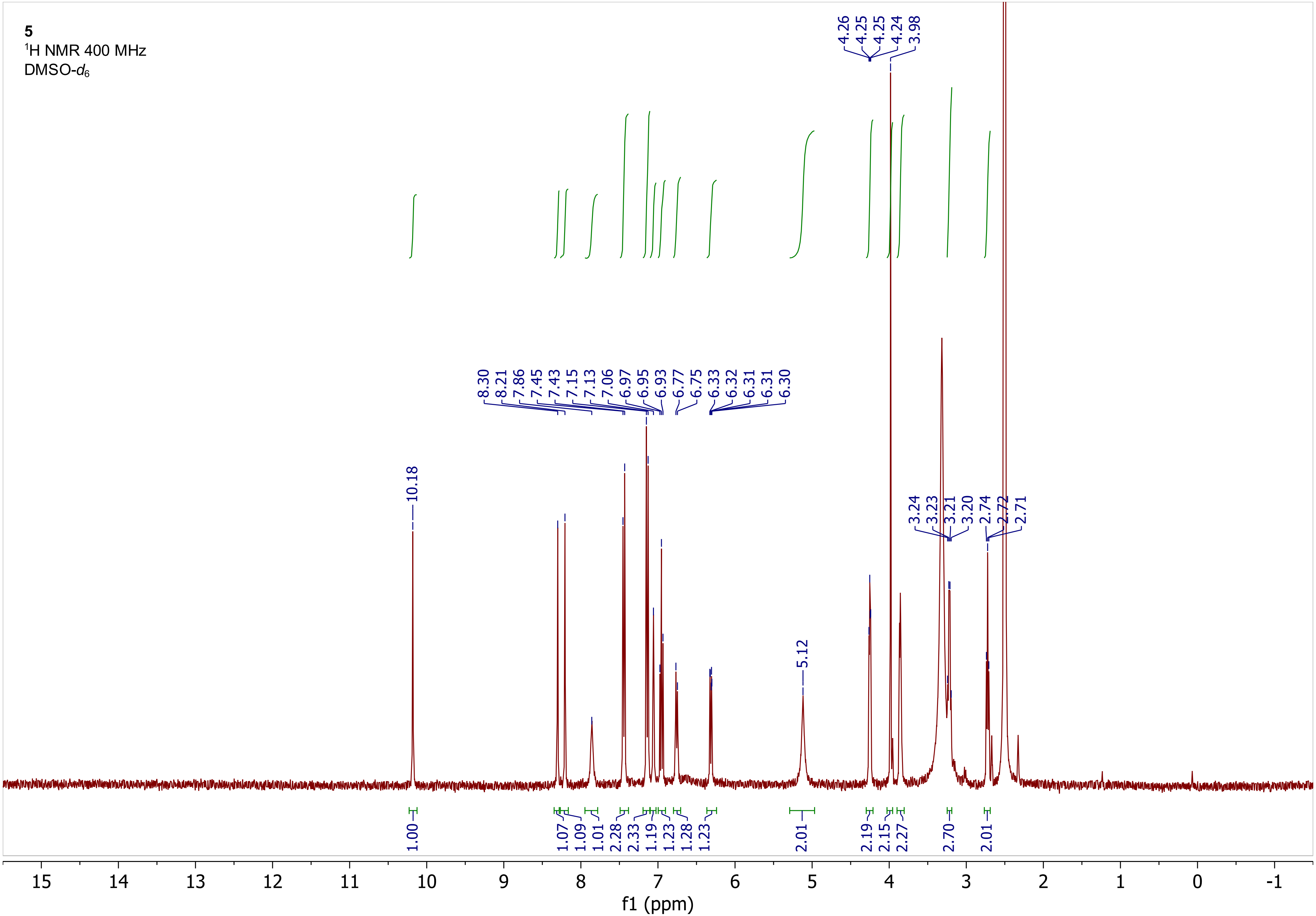
5
1H NMR 400 MHz
DMSO-d6
f1 (ppm)

**5**
$^{19}F$ NMR 376 MHz
DMSO-$d_6$

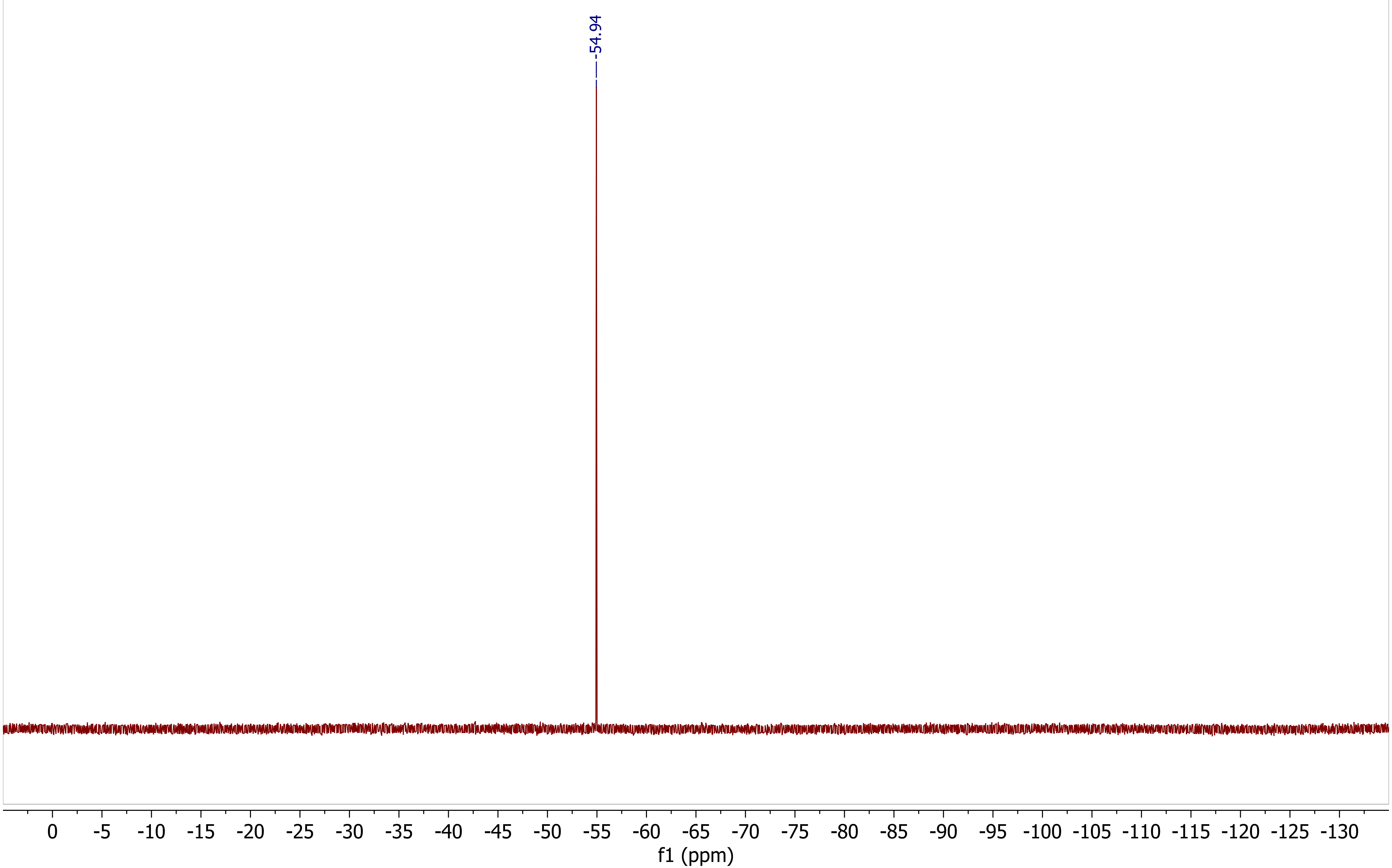

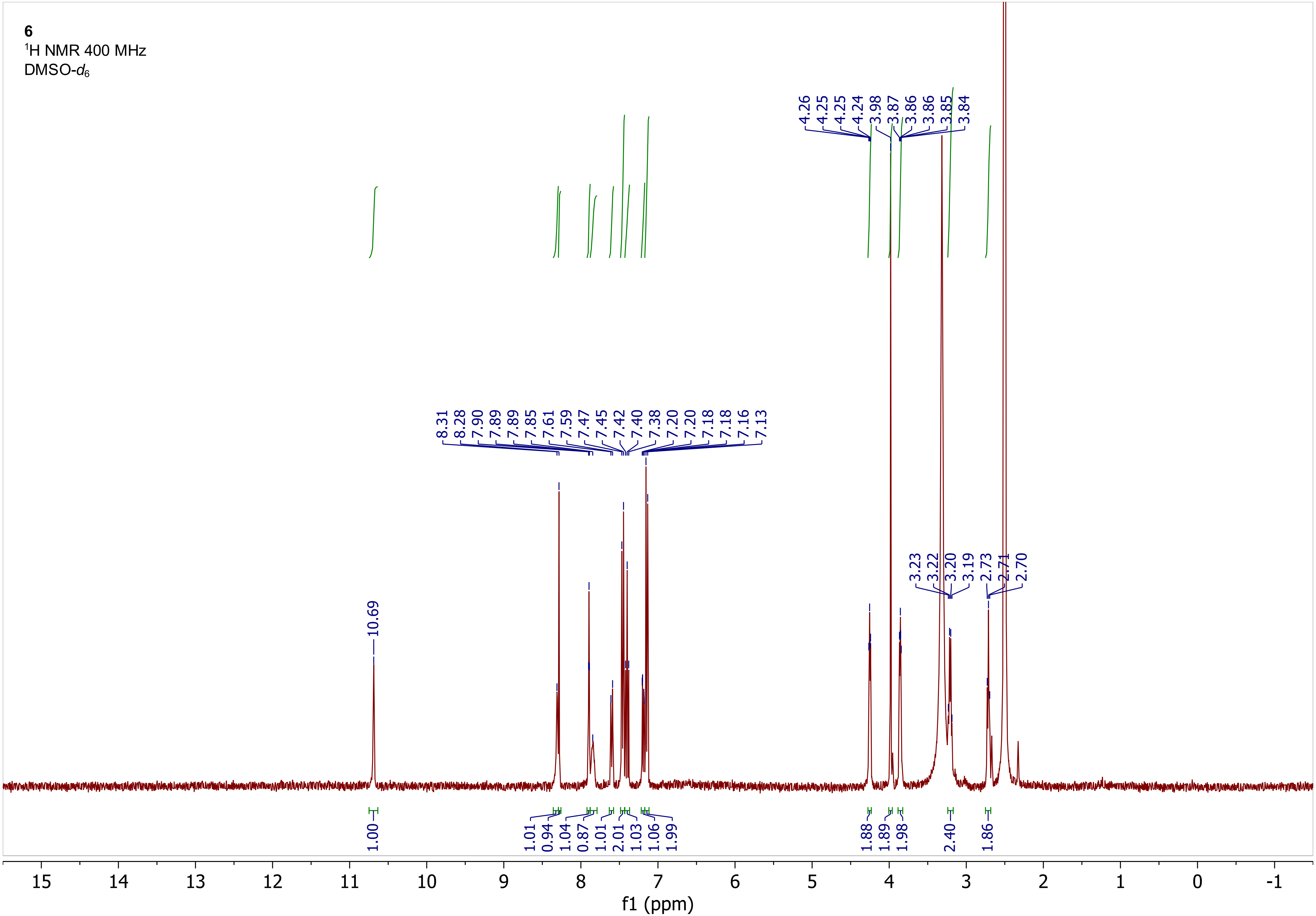
6
$^1$H NMR 400 MHz
DMSO-$d_6$
10.69
8.31 8.28 7.90 7.89 7.89 7.85 7.61 7.59 7.47 7.45 7.42 7.40 7.38 7.20 7.20 7.18 7.18 7.16 7.13
4.26 4.25 4.25 4.24 3.98 3.87 3.86 3.86 3.85 3.84
3.23 3.22 3.20 3.19 2.73 2.71 2.70
1.00 1.01 0.94 1.04 0.87 1.01 2.01 1.03 1.06 1.99 1.88 1.89 1.98 2.40 1.86
15 14 13 12 11 10 9 8 7 6 5 4 3 2 1 0 -1
f1 (ppm)

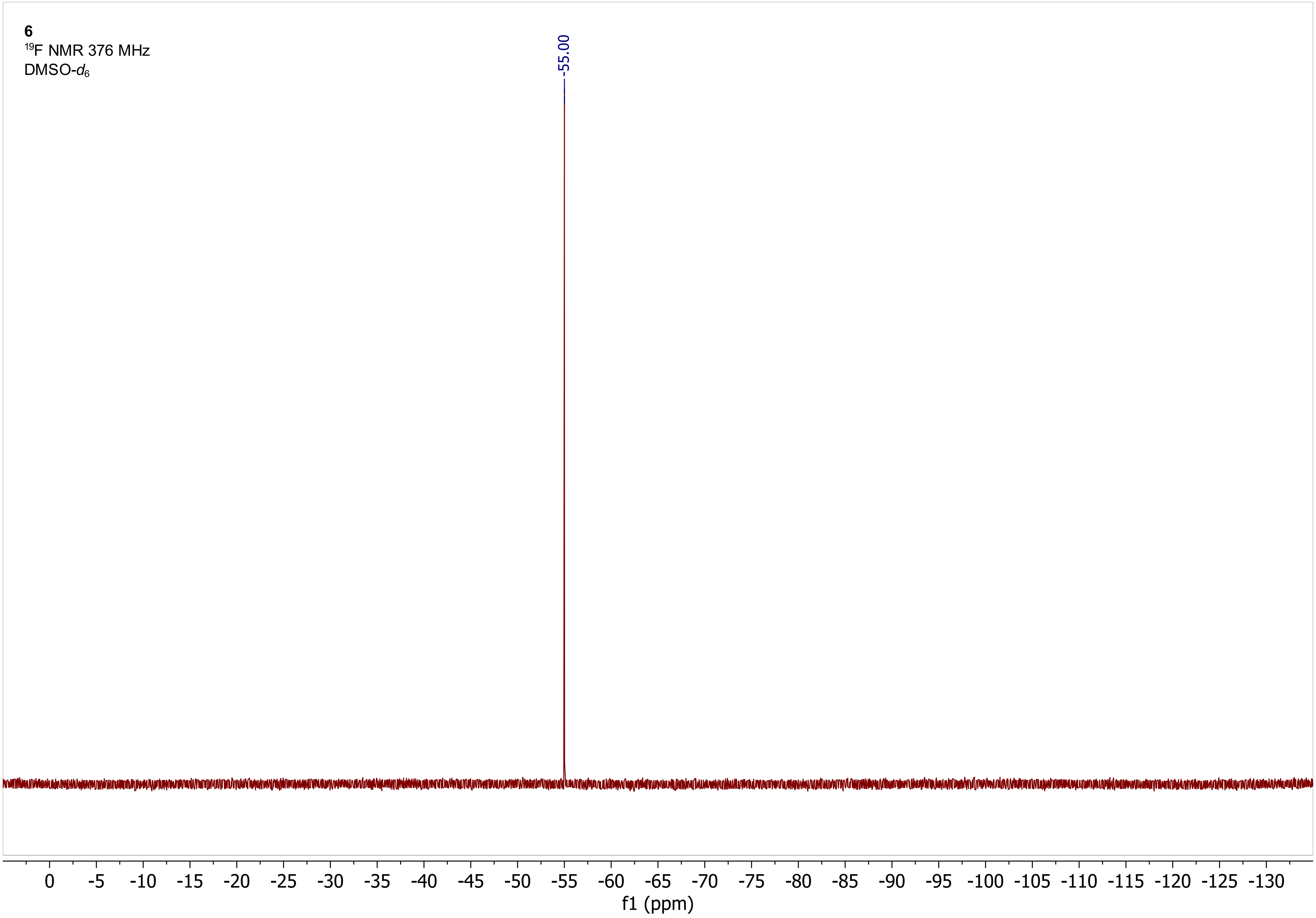
6
$^{19}$F NMR 376 MHz
DMSO-$d_6$
-55.00
0 -5 -10 -15 -20 -25 -30 -35 -40 -45 -50 -55 -60 -65 -70 -75 -80 -85 -90 -95 -100 -105 -110 -115 -120 -125 -130
f1 (ppm)

**7**

$^{1}$H NMR 400 MHz

DMSO-$d_6$

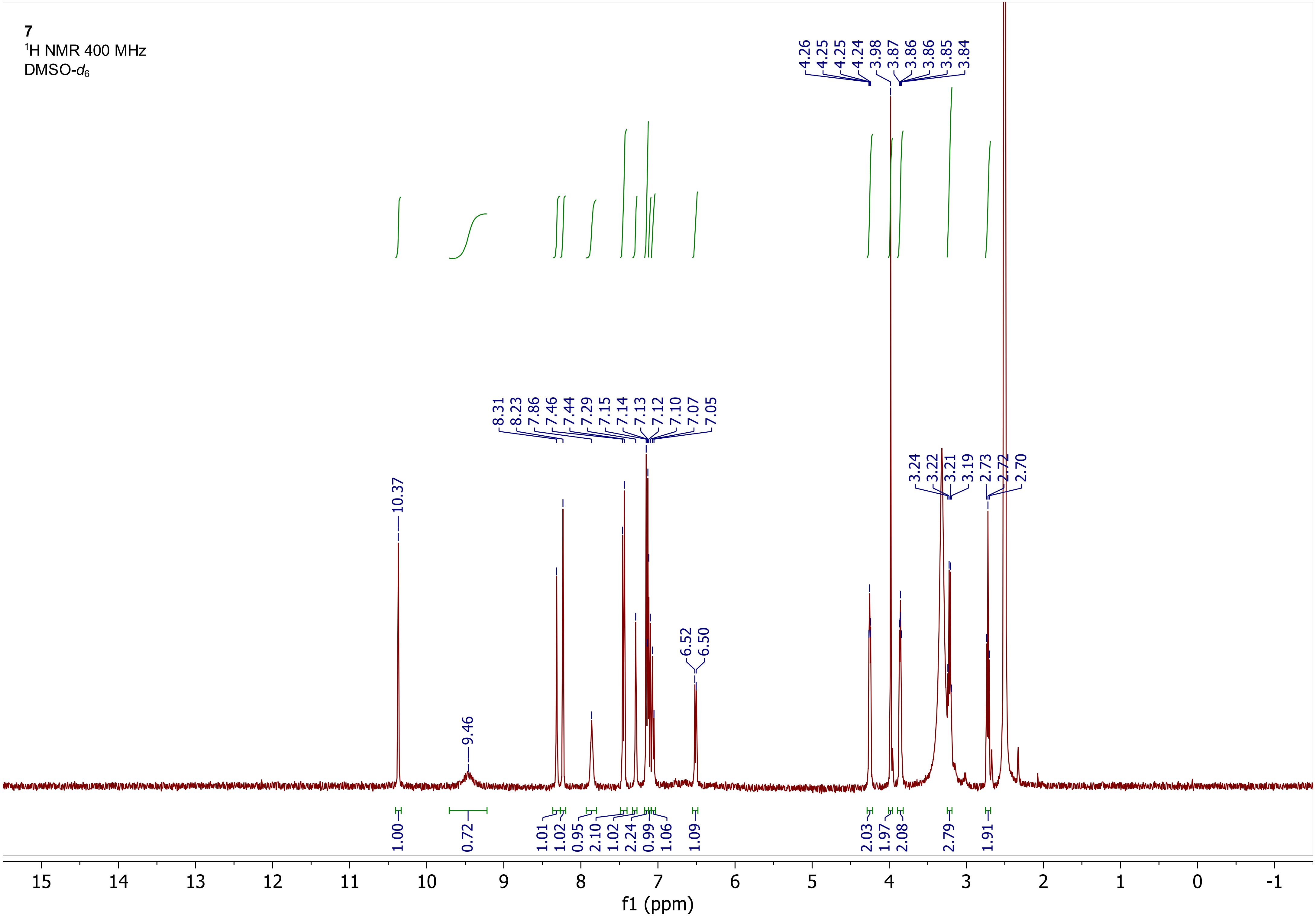

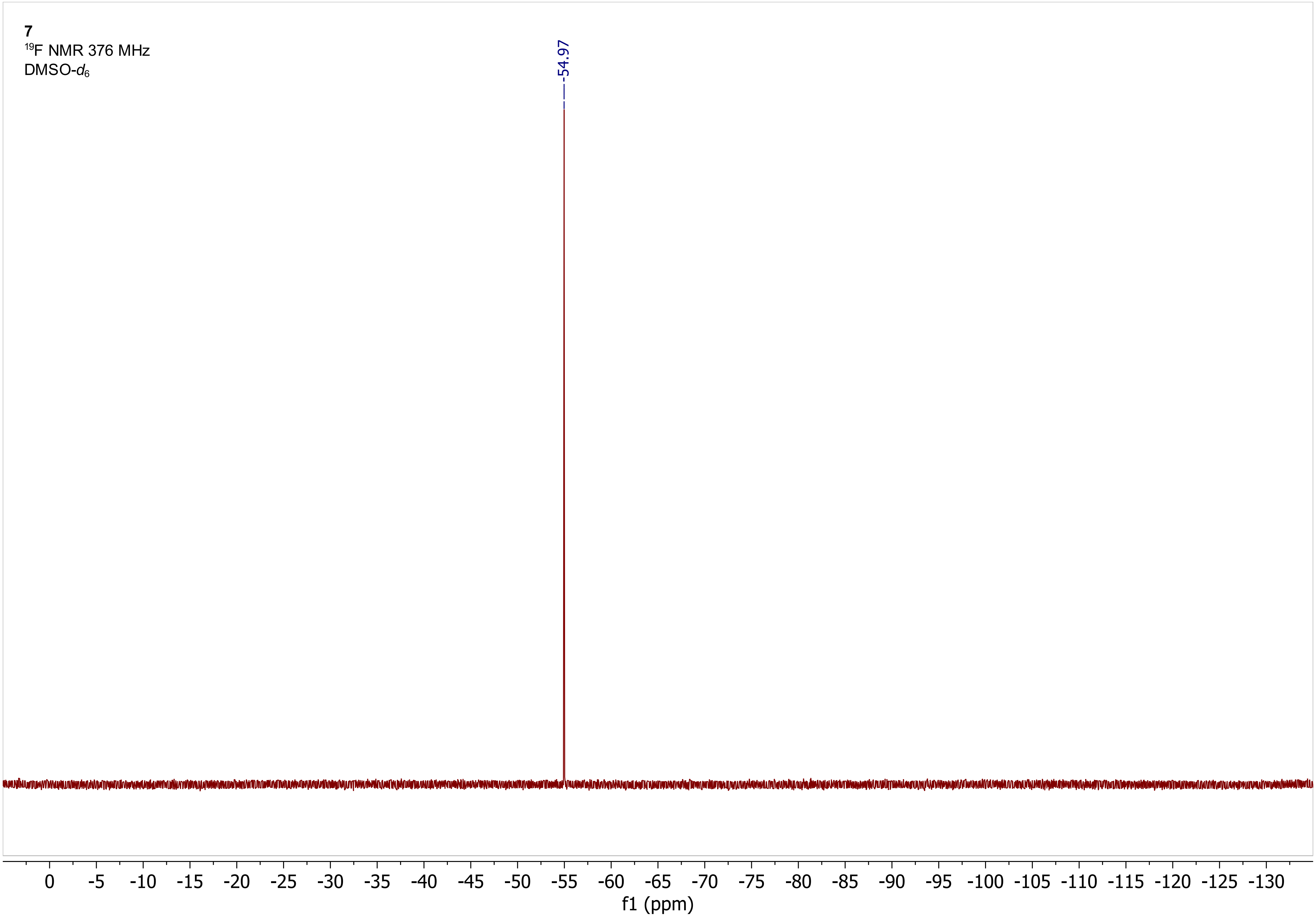
7
19F NMR 376 MHz
DMSO-d6
-54.97
0
-5
-10
-15
-20
-25
-30
-35
-40
-45
-50
-55
-60
-65
-70
-75
-80
-85
-90
-95
-100
-105
-110
-115
-120
-125
-130
f1 (ppm)

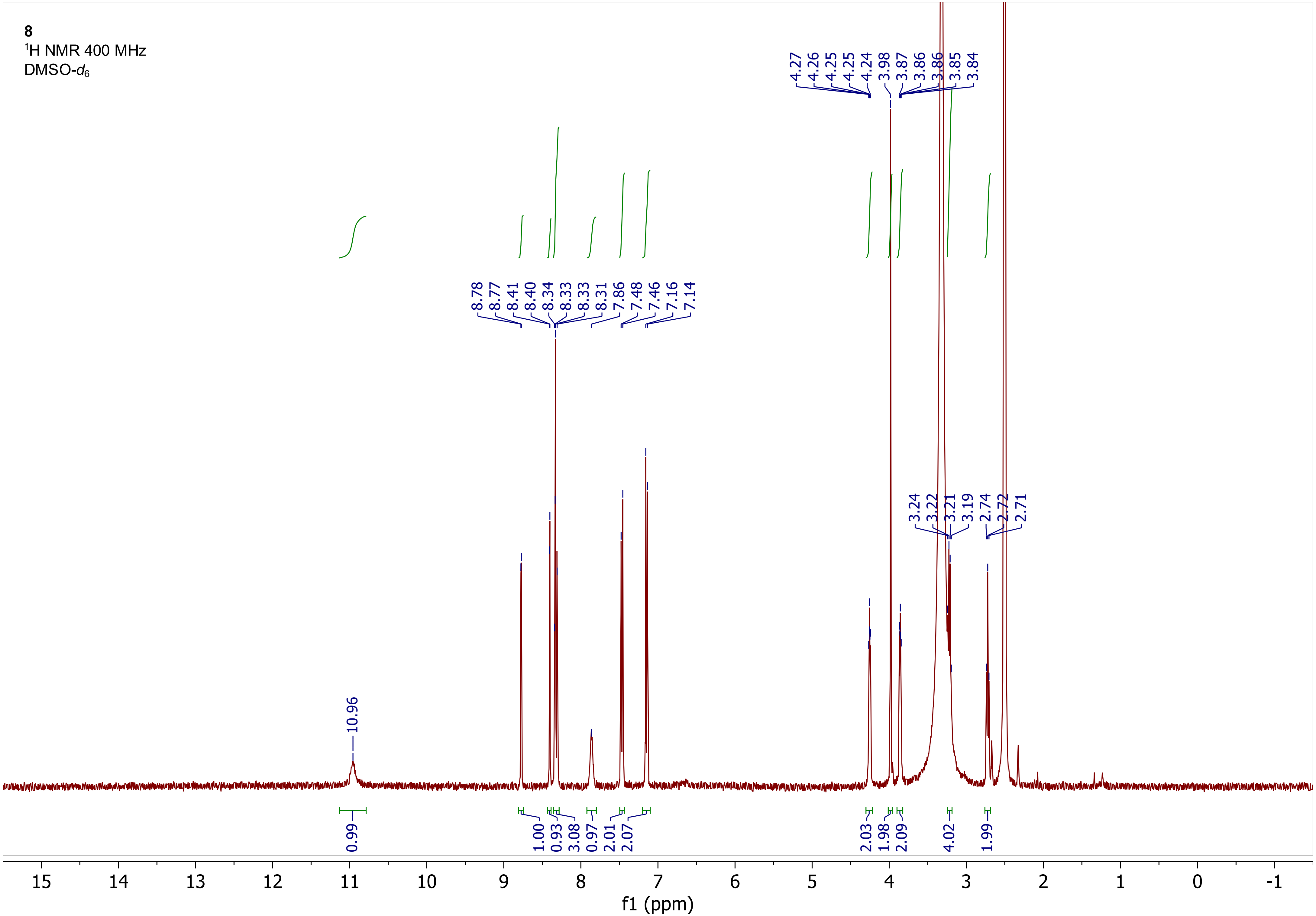

8
1H NMR 400 MHz
DMSO-d6
4.27
4.26
4.25
4.25
4.24
3.98
3.87
3.86
3.86
3.85
3.84
8.78
8.77
8.41
8.40
8.34
8.33
8.33
8.31
7.86
7.48
7.46
7.16
7.14
3.24
3.22
3.21
3.19
2.74
2.72
2.71
10.96
0.99
1.00
0.93
3.08
0.97
2.01
2.07
2.03
1.98
2.09
4.02
1.99
15
14
13
12
11
10
9
8
7
6
5
4
3
2
1
0
-1
f1 (ppm)

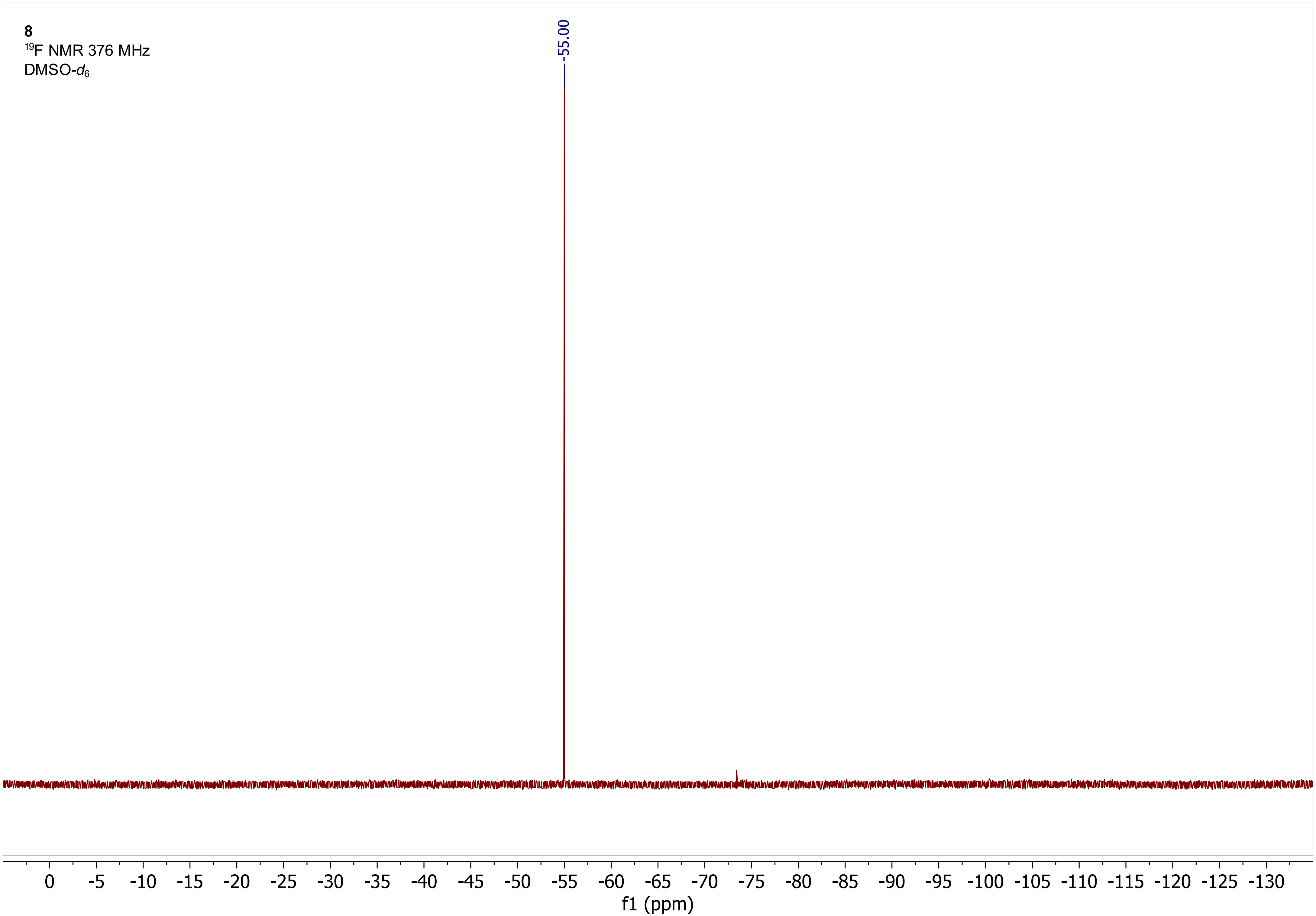
8
$^{19}$F NMR 376 MHz
DMSO-$d_6$
-55.00
0 -5 -10 -15 -20 -25 -30 -35 -40 -45 -50 -55 -60 -65 -70 -75 -80 -85 -90 -95 -100 -105 -110 -115 -120 -125 -130
f1 (ppm)

**9**
$^1$H NMR 600 MHz
DMSO-$d_6$

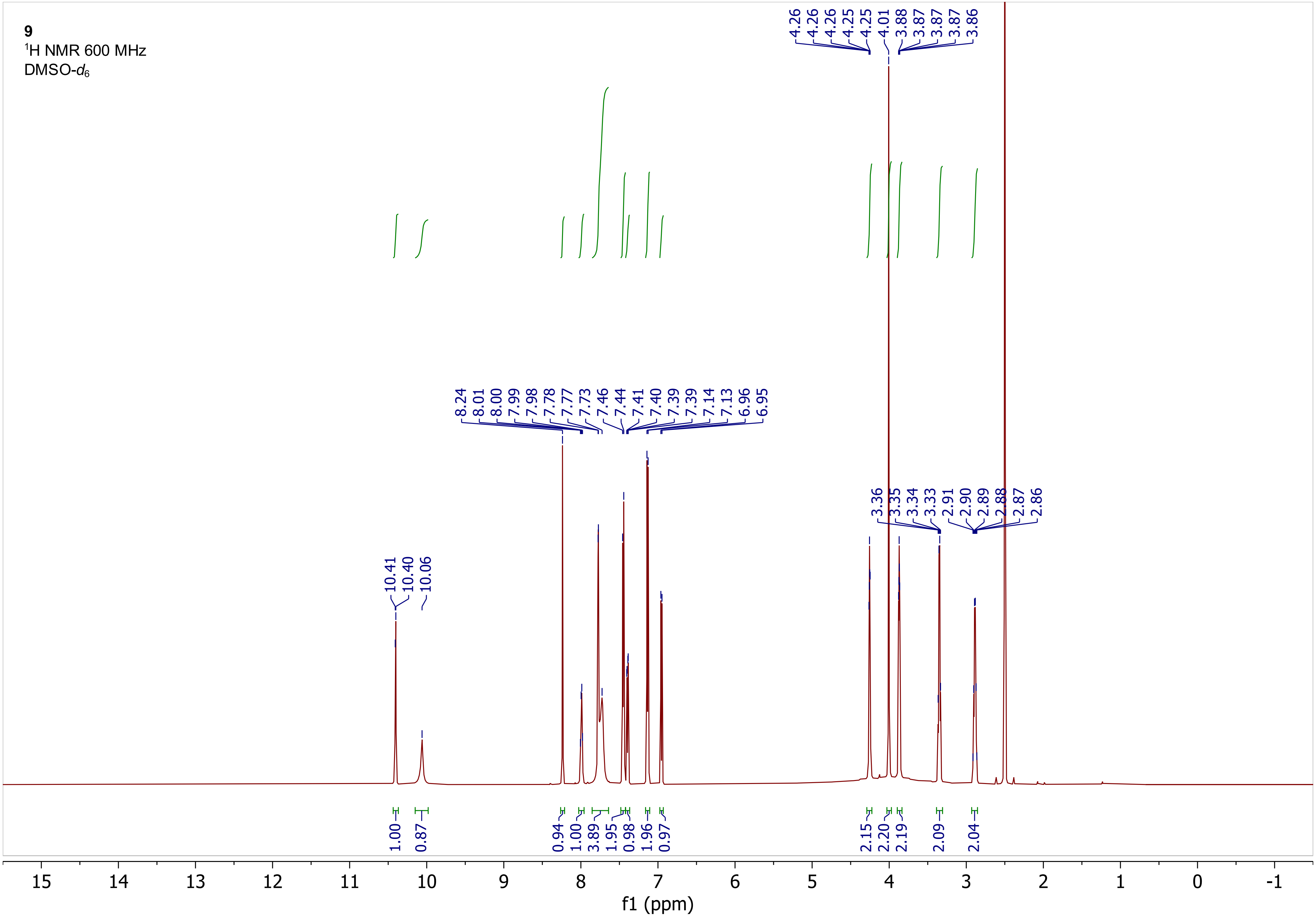

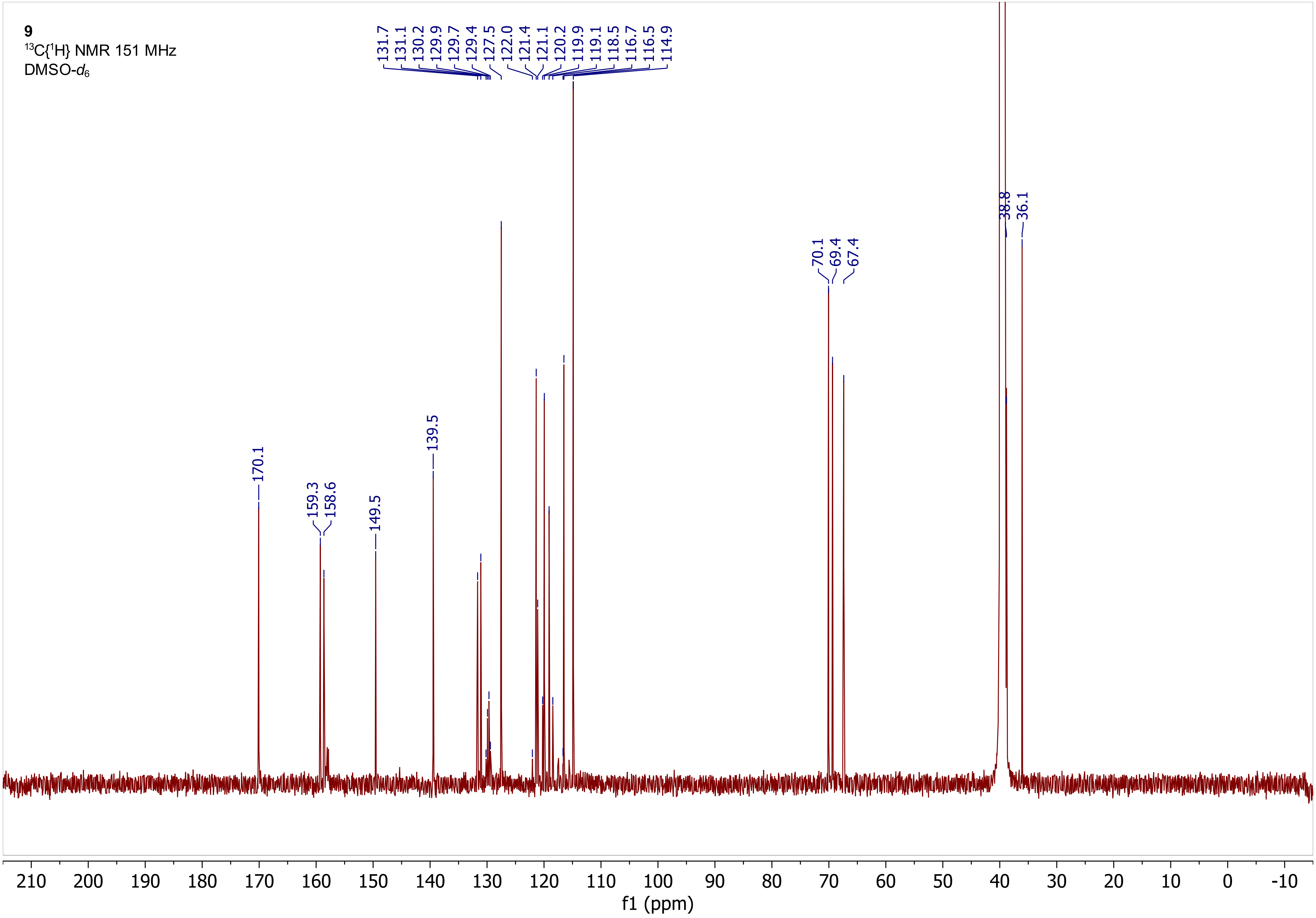

9
$^{13}C\{^1H\}$ NMR 151 MHz
DMSO-$d_6$
131.7
131.1
130.2
129.9
129.7
129.4
127.5
122.0
121.4
121.1
120.2
119.9
119.1
118.5
116.7
116.5
114.9
170.1
159.3
158.6
149.5
139.5
70.1
69.4
67.4
38.8
36.1
210
200
190
180
170
160
150
140
130
120
110
100
90
80
70
60
50
40
30
20
10
0
-10
f1 (ppm)

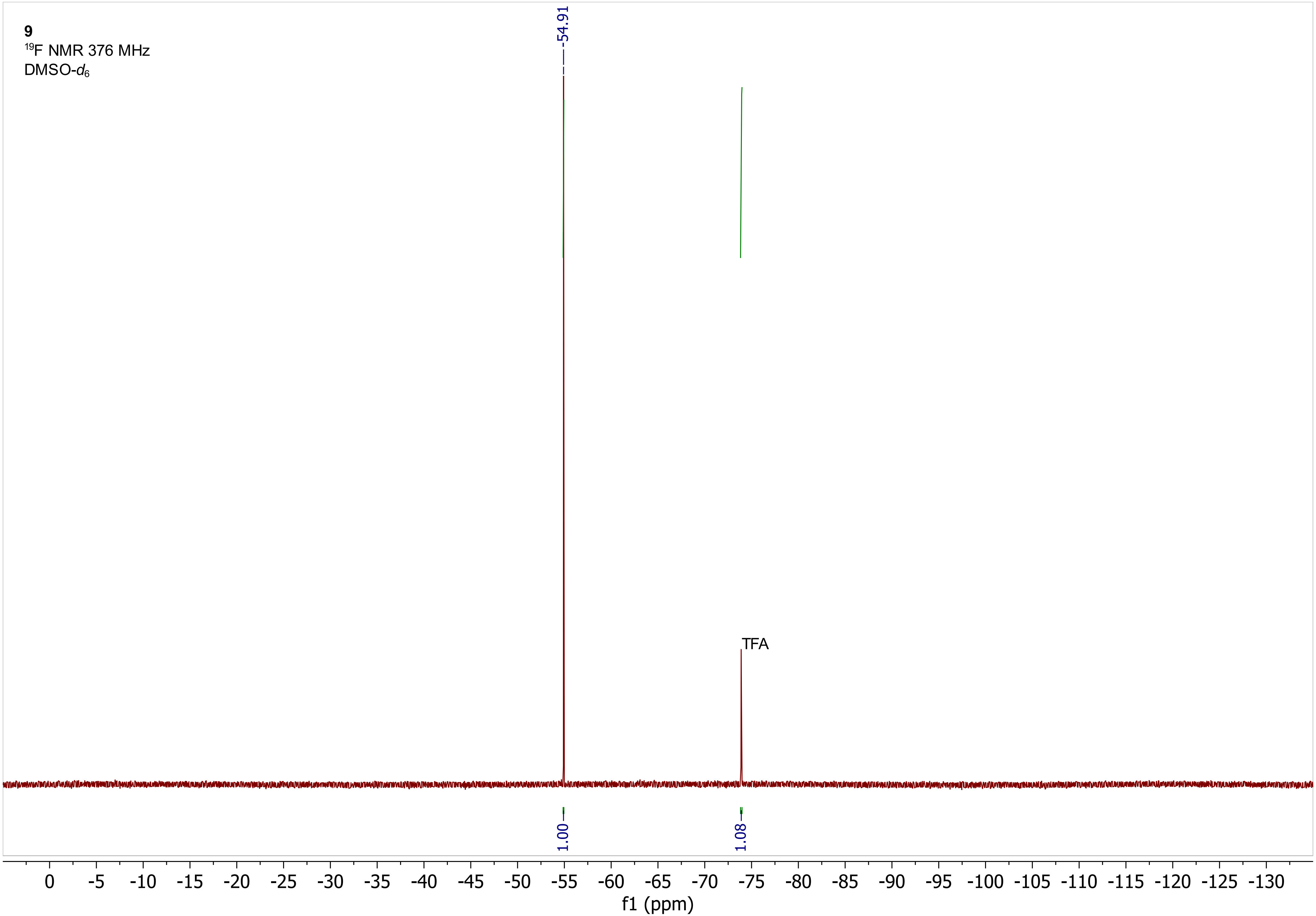

9
$^{19}$F NMR 376 MHz
DMSO-$d_6$
-54.91
TFA
1.00
1.08
0
-5
-10
-15
-20
-25
-30
-35
-40
-45
-50
-55
-60
-65
-70
-75
-80
-85
-90
-95
-100
-105
-110
-115
-120
-125
-130
f1 (ppm)

**9**
$^{1}$H COSY 600 MHz
DMSO-$d_6$

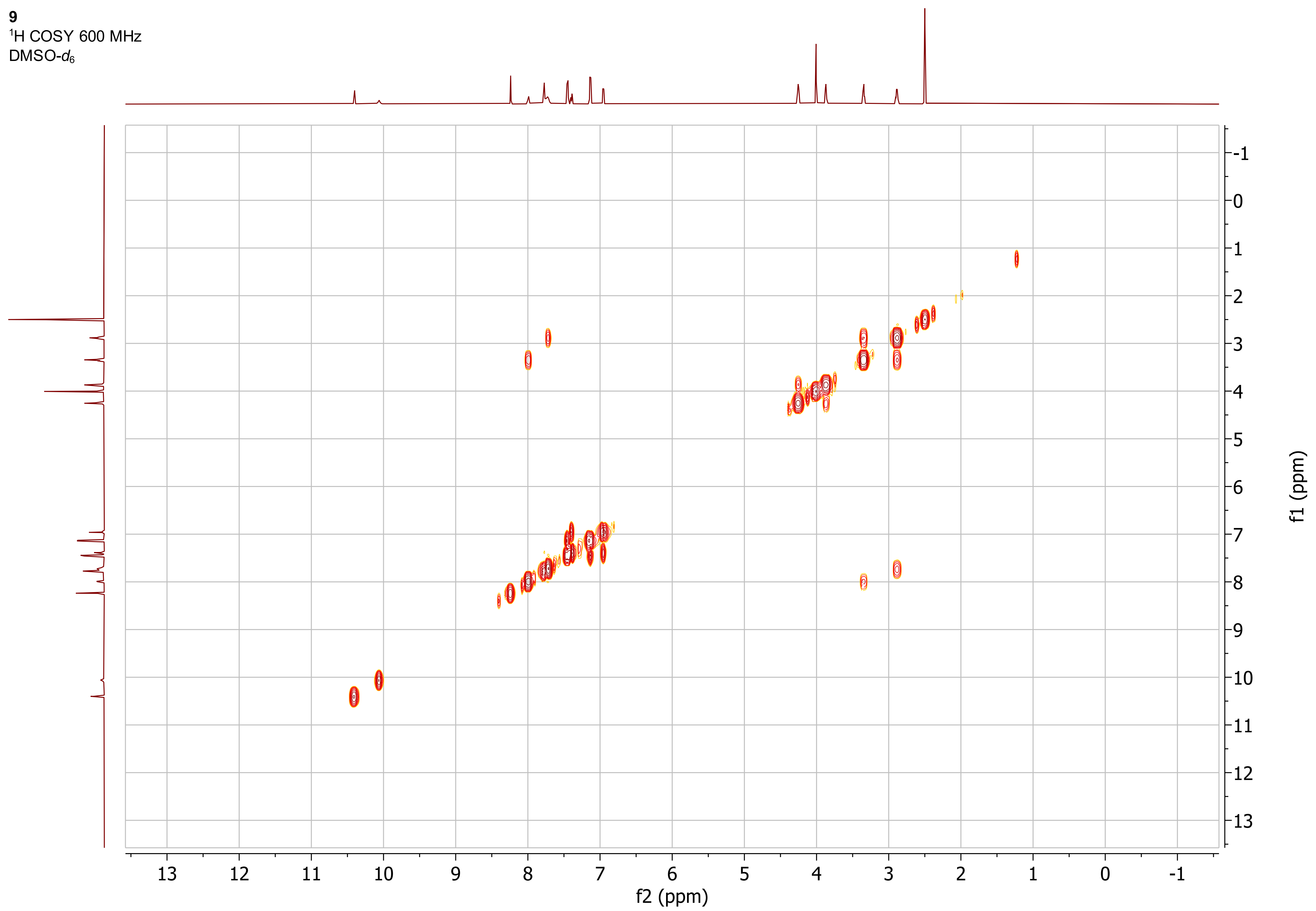

**9**
$^1H$-$^{13}C$ ed-HSQC 600 MHz
DMSO-$d_6$

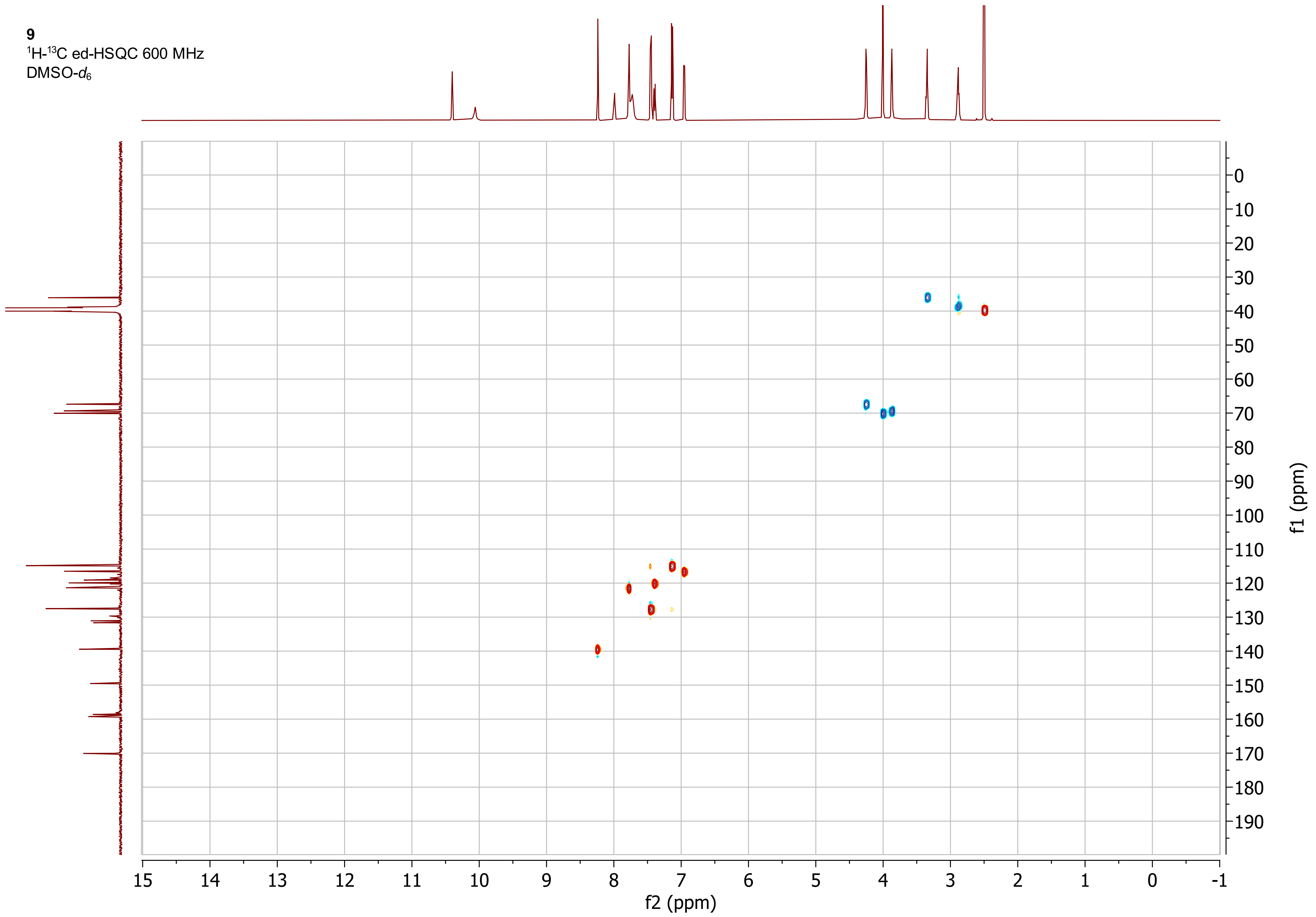

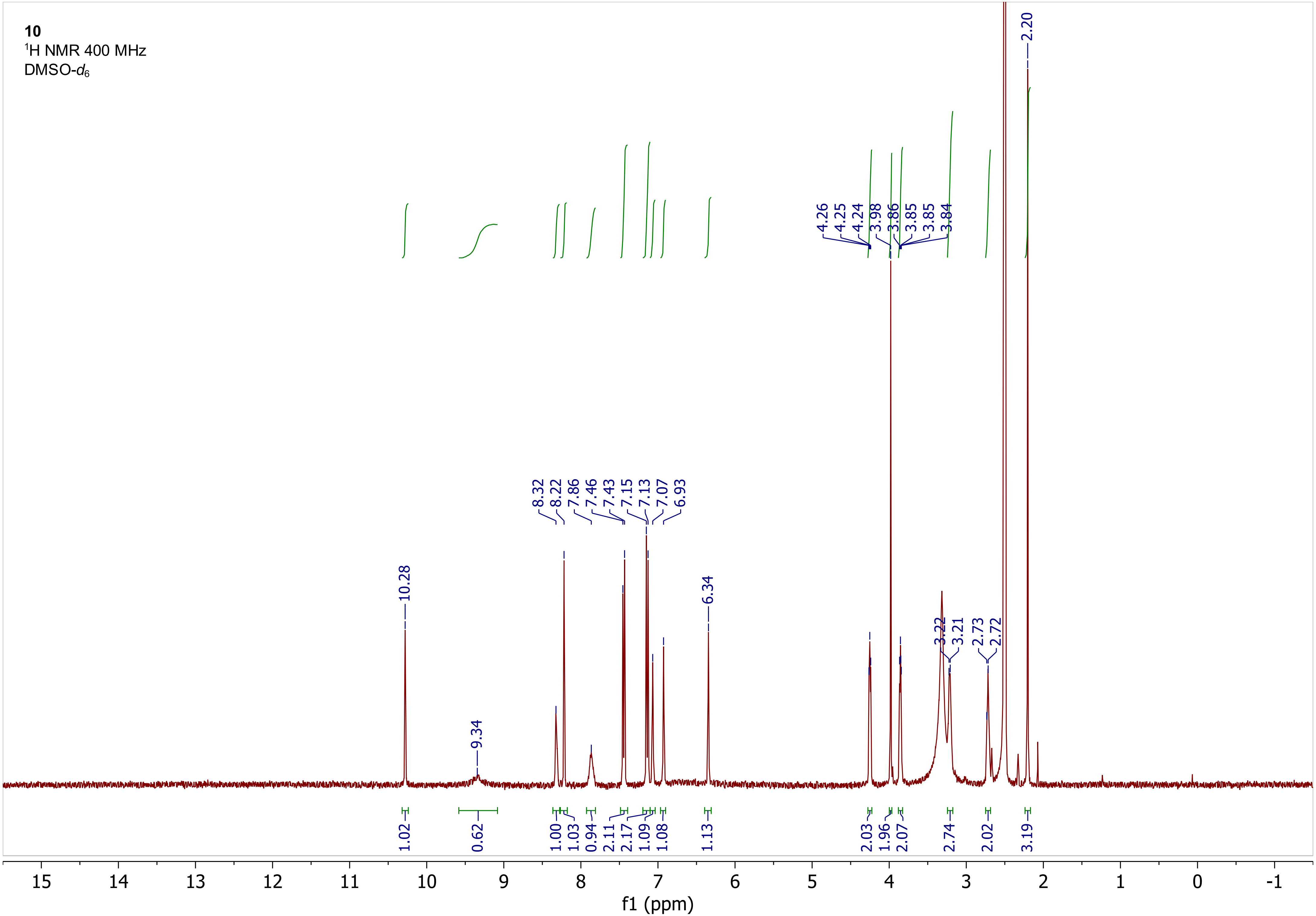
10
1H NMR 400 MHz
DMSO-d6
10.28
9.34
8.32
8.22
7.86
7.46
7.43
7.15
7.13
7.07
6.93
6.34
4.26
4.25
4.24
3.98
3.86
3.85
3.85
3.84
3.22
3.21
2.73
2.72
2.20
1.02
0.62
1.00
1.03
0.94
2.11
2.17
1.09
1.08
1.13
2.03
1.96
2.07
2.74
2.02
3.19
15
14
13
12
11
10
9
8
7
6
5
4
3
2
1
0
-1
f1 (ppm)

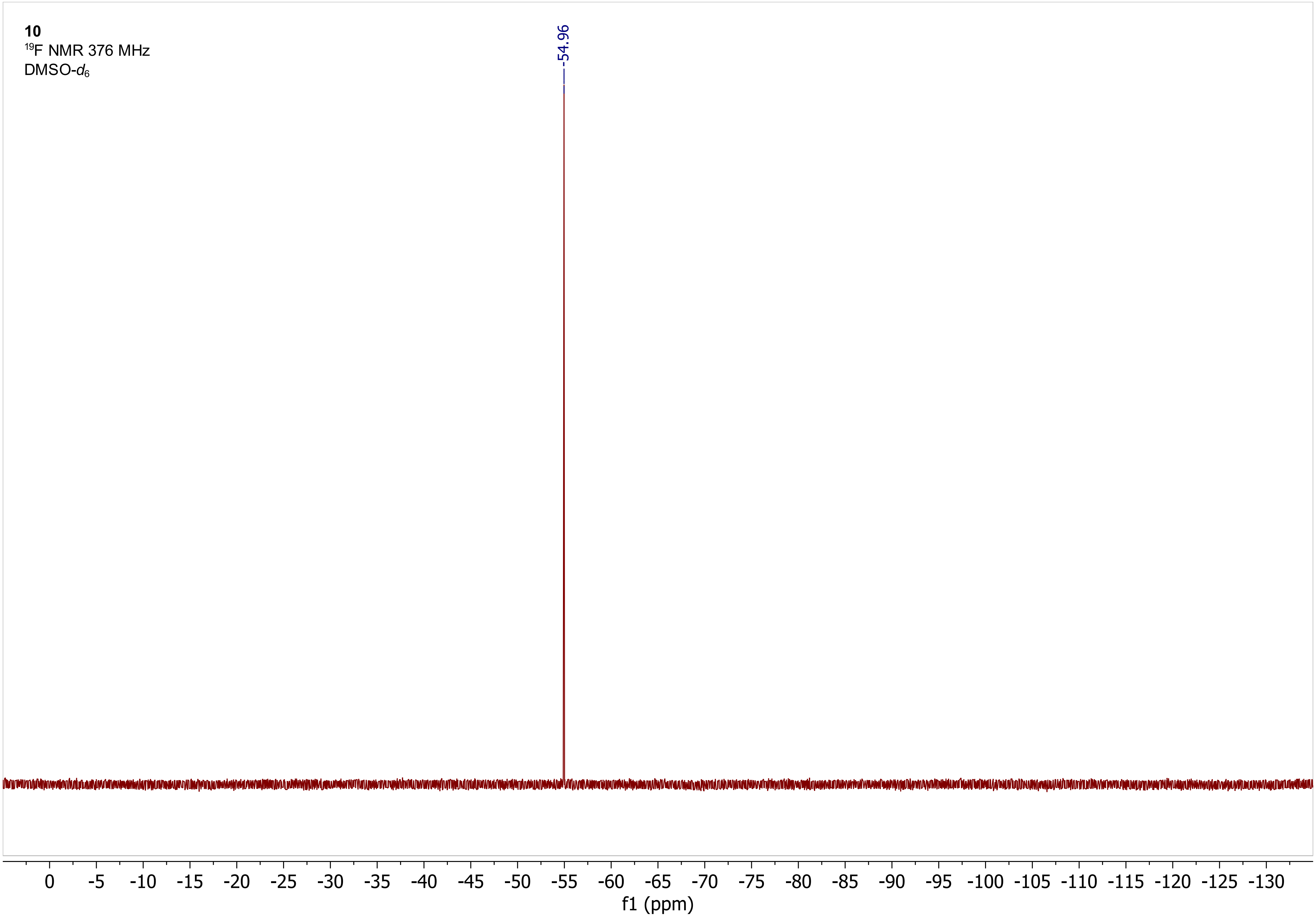
10
19F NMR 376 MHz
DMSO-d6
-54.96
0 -5 -10 -15 -20 -25 -30 -35 -40 -45 -50 -55 -60 -65 -70 -75 -80 -85 -90 -95 -100 -105 -110 -115 -120 -125 -130
f1 (ppm)

**11**

$^1$H NMR 400 MHz

DMSO-$d_6$

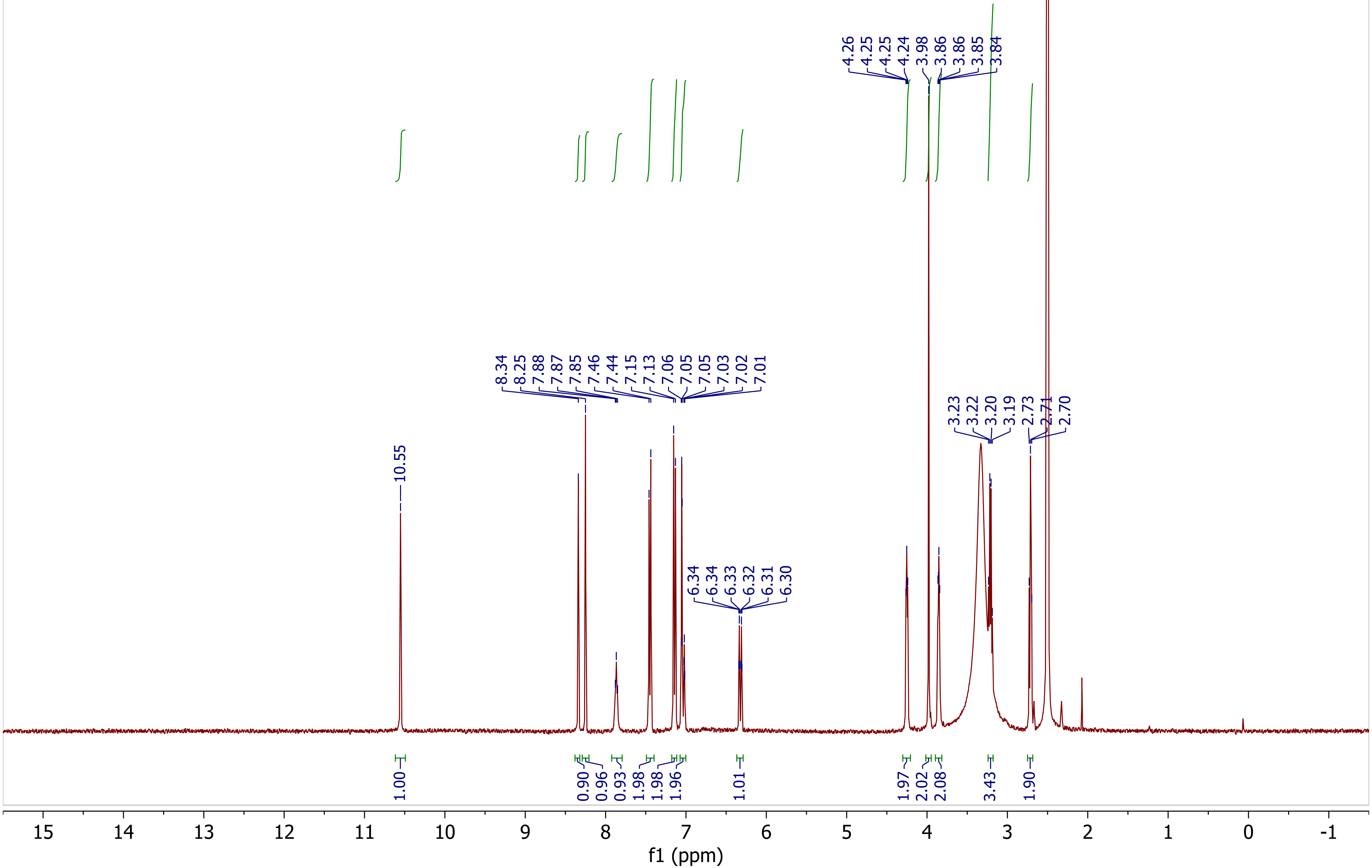

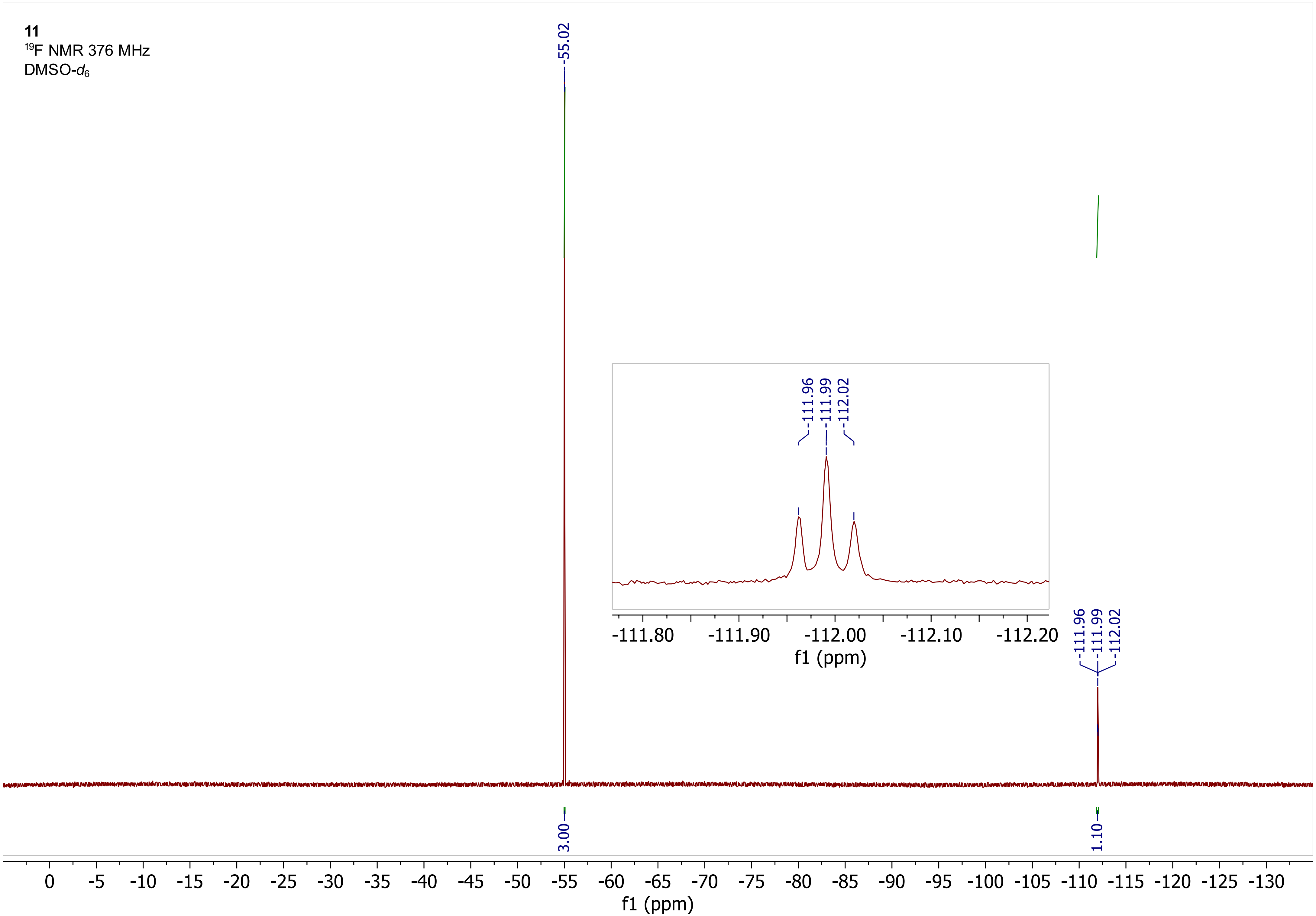

11
19F NMR 376 MHz
DMSO-d6
-55.02
-111.96
-111.99
-112.02
3.00
1.10
-111.80
-111.90
-112.00
-112.10
-112.20
f1 (ppm)
0
-5
-10
-15
-20
-25
-30
-35
-40
-45
-50
-55
-60
-65
-70
-75
-80
-85
-90
-95
-100
-105
-110
-115
-120
-125
-130
f1 (ppm)

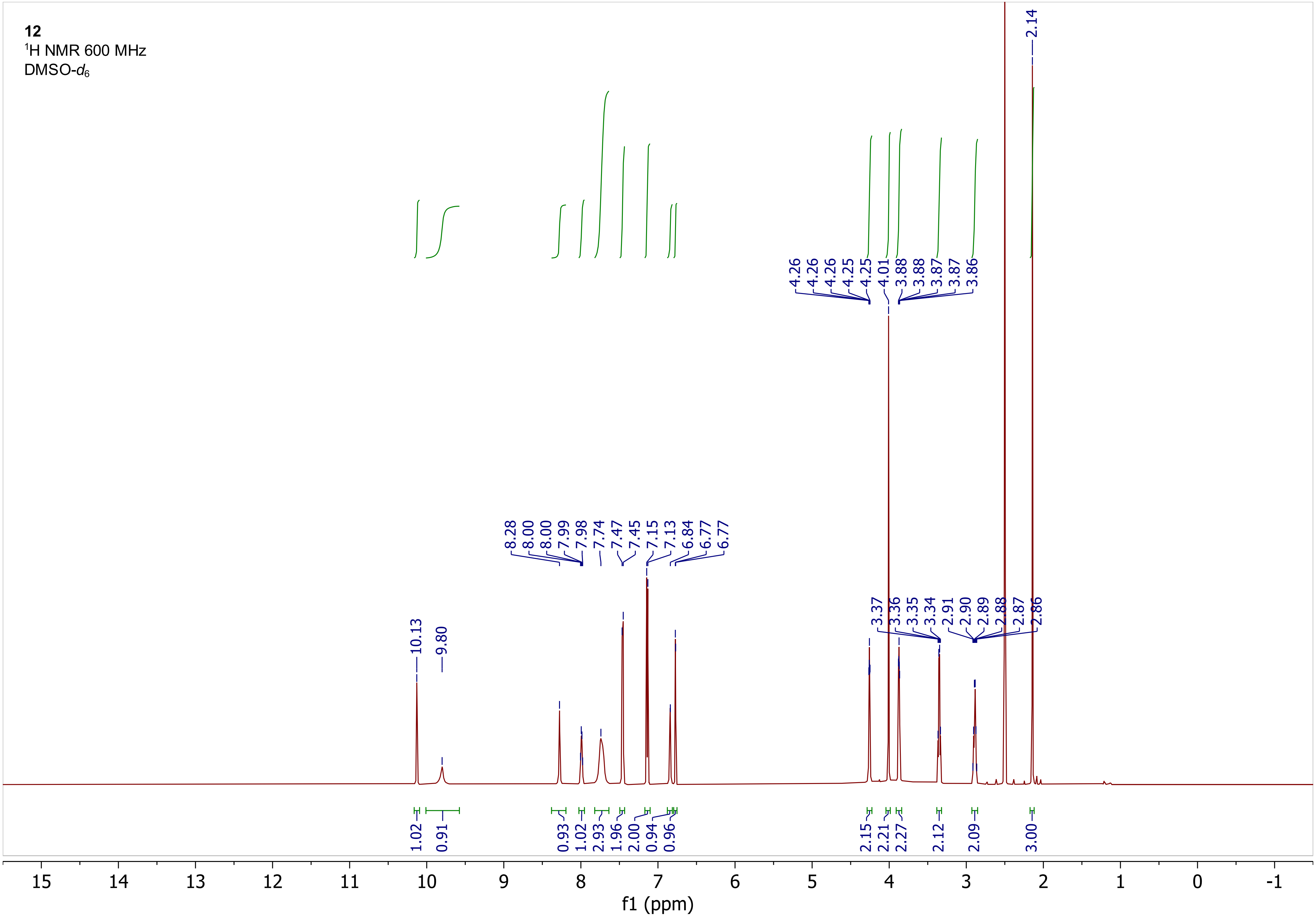

12
1H NMR 600 MHz
DMSO-d6
10.13
9.80
8.28
8.00
8.00
7.99
7.98
7.74
7.47
7.45
7.15
7.13
6.84
6.77
6.77
4.26
4.26
4.26
4.25
4.25
4.01
3.88
3.88
3.87
3.87
3.86
3.37
3.36
3.35
3.34
2.91
2.90
2.89
2.88
2.87
2.86
2.14
1.02
0.91
0.93
1.02
2.93
1.96
2.00
0.94
0.96
2.15
2.21
2.27
2.12
2.09
3.00
15
14
13
12
11
10
9
8
7
6
5
4
3
2
1
0
-1
f1 (ppm)

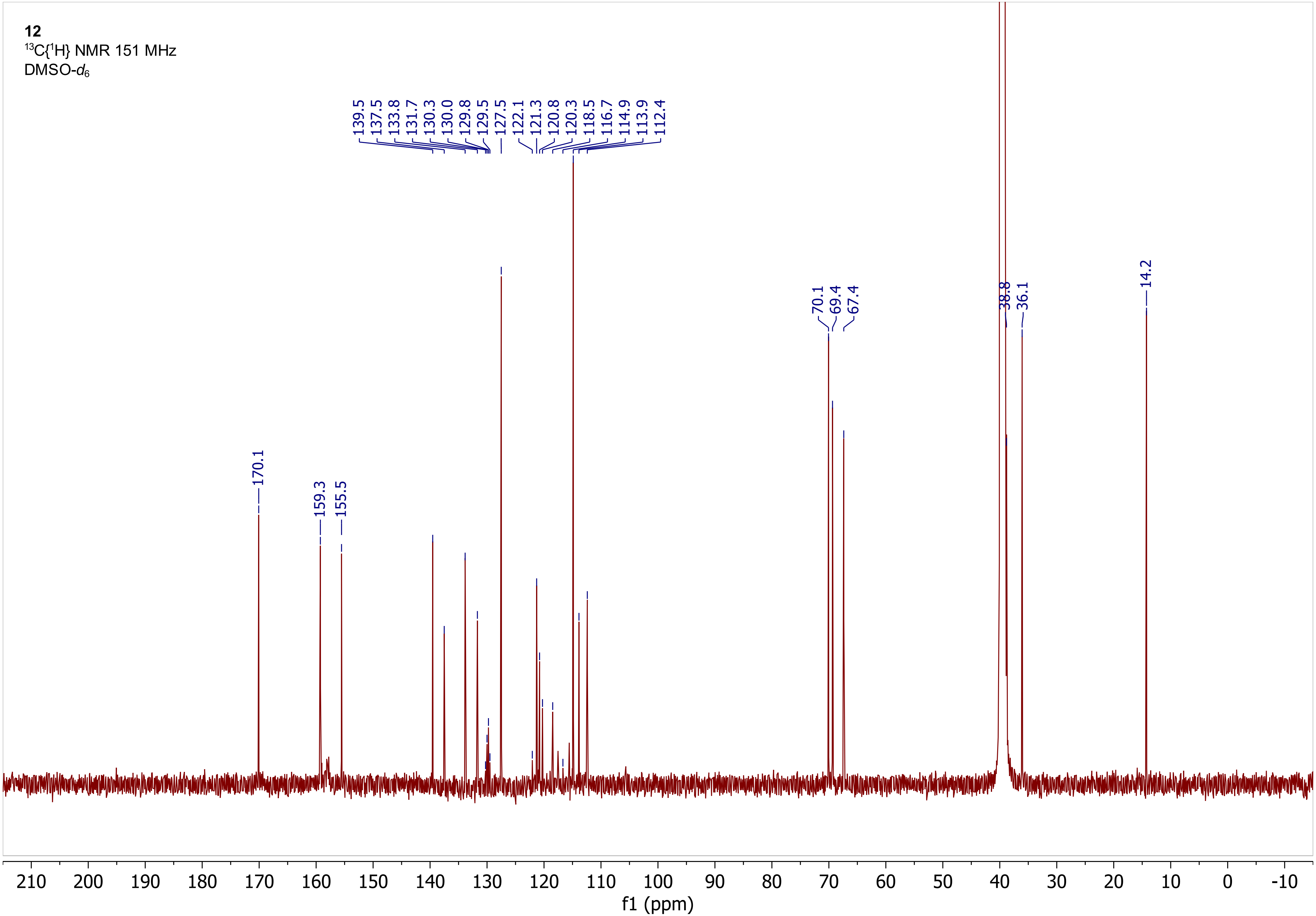
12
$^{13}C\{^1H\}$ NMR 151 MHz
DMSO-$d_6$
139.5
137.5
133.8
131.7
130.3
130.0
129.8
129.5
127.5
122.1
121.3
120.8
120.3
118.5
116.7
114.9
113.9
112.4
170.1
159.3
155.5
70.1
69.4
67.4
38.8
36.1
14.2
210
200
190
180
170
160
150
140
130
120
110
100
90
80
70
60
50
40
30
20
10
0
-10
f1 (ppm)

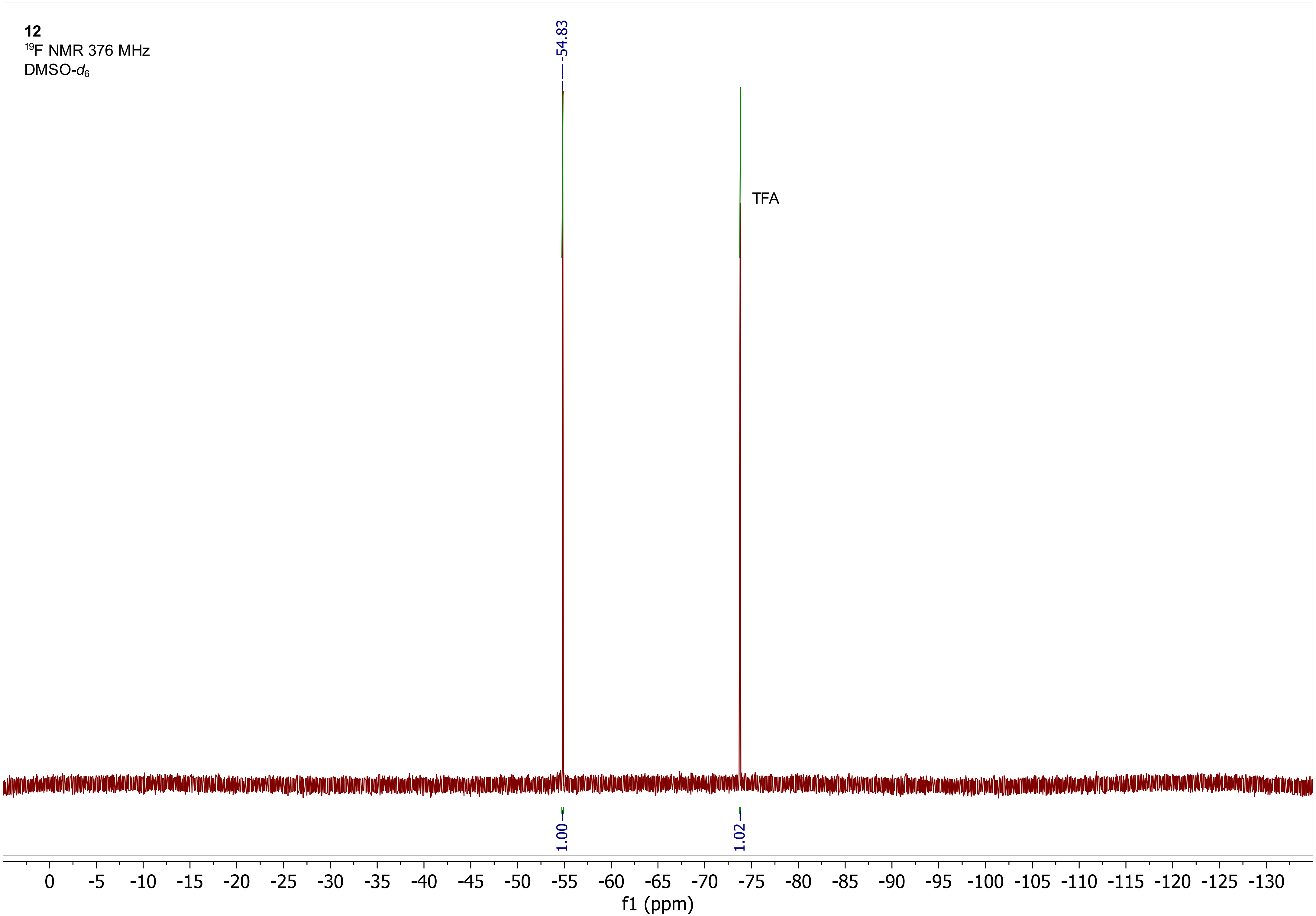
12
19F NMR 376 MHz
DMSO-d6
-54.83
TFA
1.00
1.02
0 -5 -10 -15 -20 -25 -30 -35 -40 -45 -50 -55 -60 -65 -70 -75 -80 -85 -90 -95 -100 -105 -110 -115 -120 -125 -130
f1 (ppm)

**12**
$^1$H COSY 600 MHz
DMSO-$d_6$

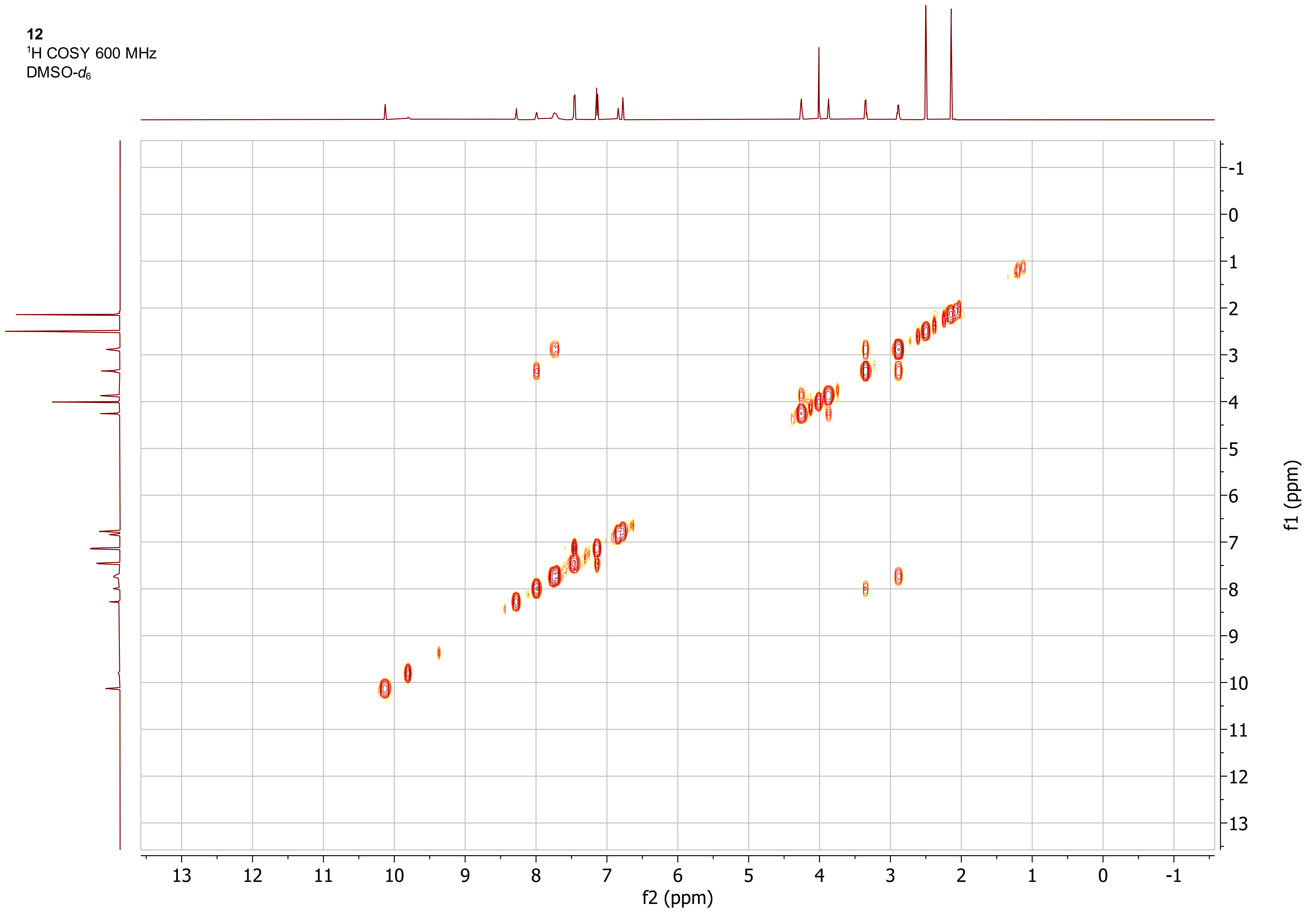

**12**
$^{1}H$-$^{13}C$ ed-HSQC 600 MHz
DMSO-$d_6$

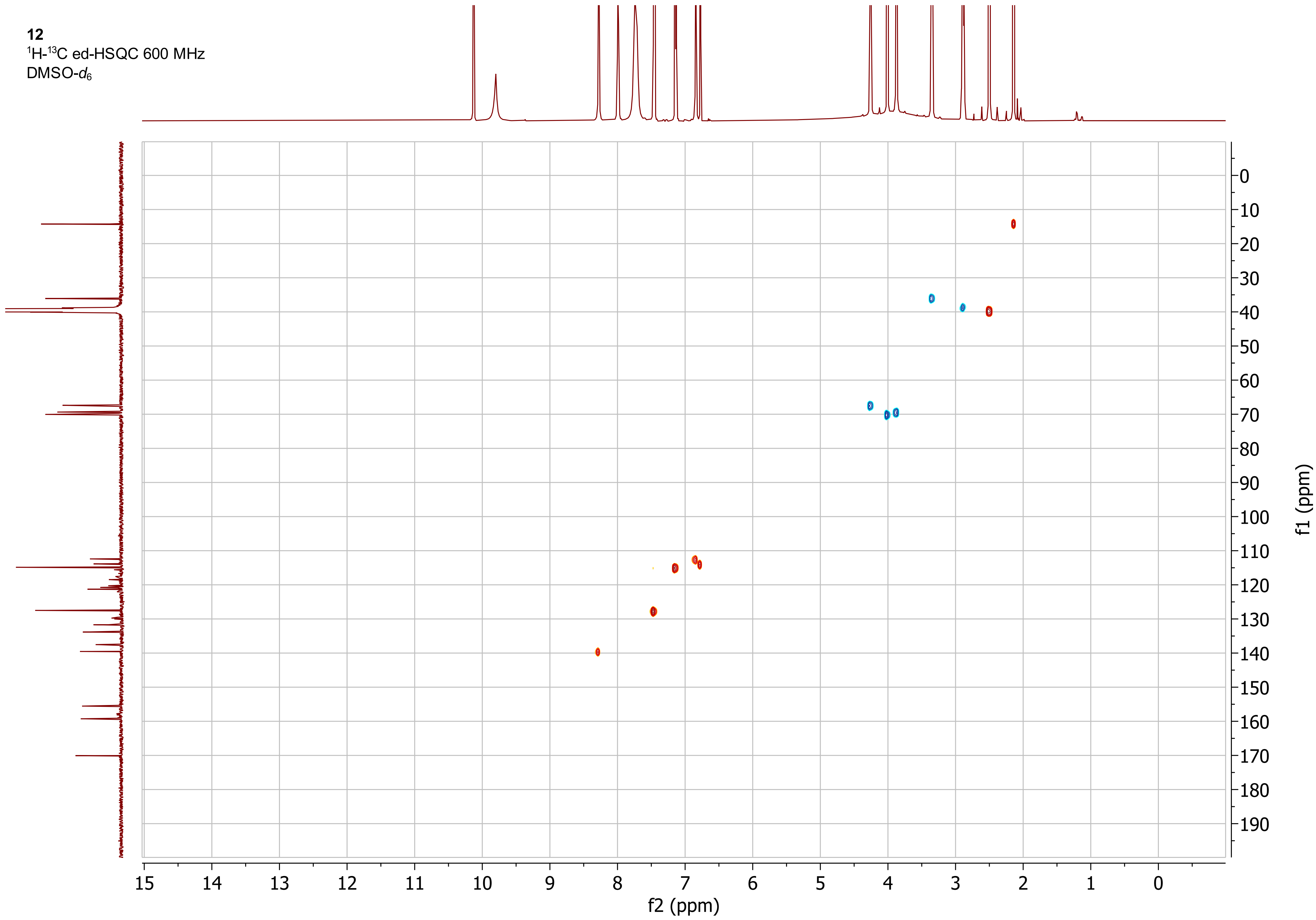

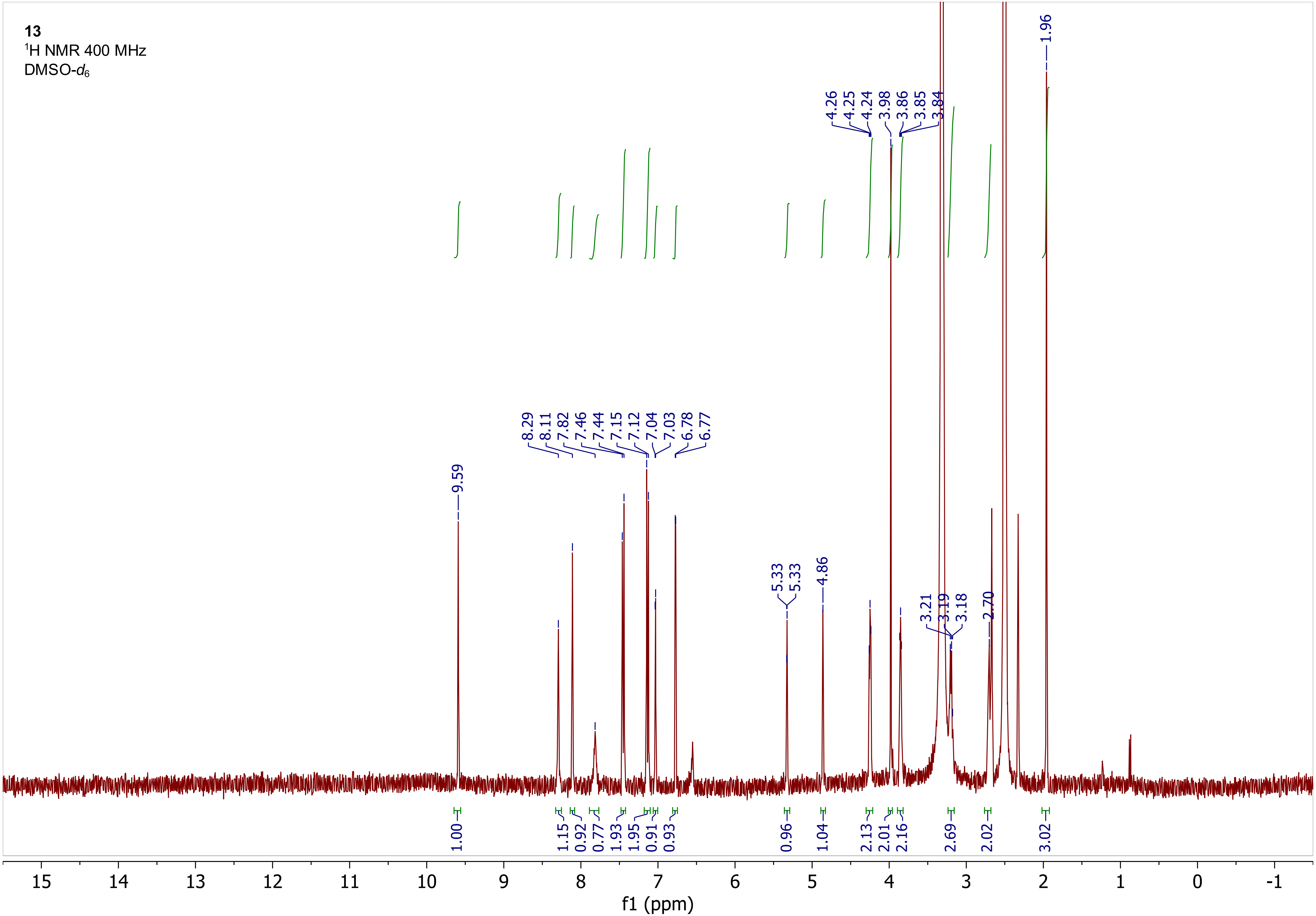
13
1H NMR 400 MHz
DMSO-d6
9.59
8.29 8.11 7.82 7.46 7.44 7.15 7.12 7.04 7.03 6.78 6.77
5.33 5.33
4.86
4.26 4.25 4.24 3.98 3.86 3.85 3.84
3.21 3.19 3.18
2.70
1.96
1.00 1.15 0.92 0.77 1.93 1.95 0.91 0.93 0.96 1.04 2.13 2.01 2.16 2.69 2.02 3.02
15 14 13 12 11 10 9 8 7 6 5 4 3 2 1 0 -1
f1 (ppm)

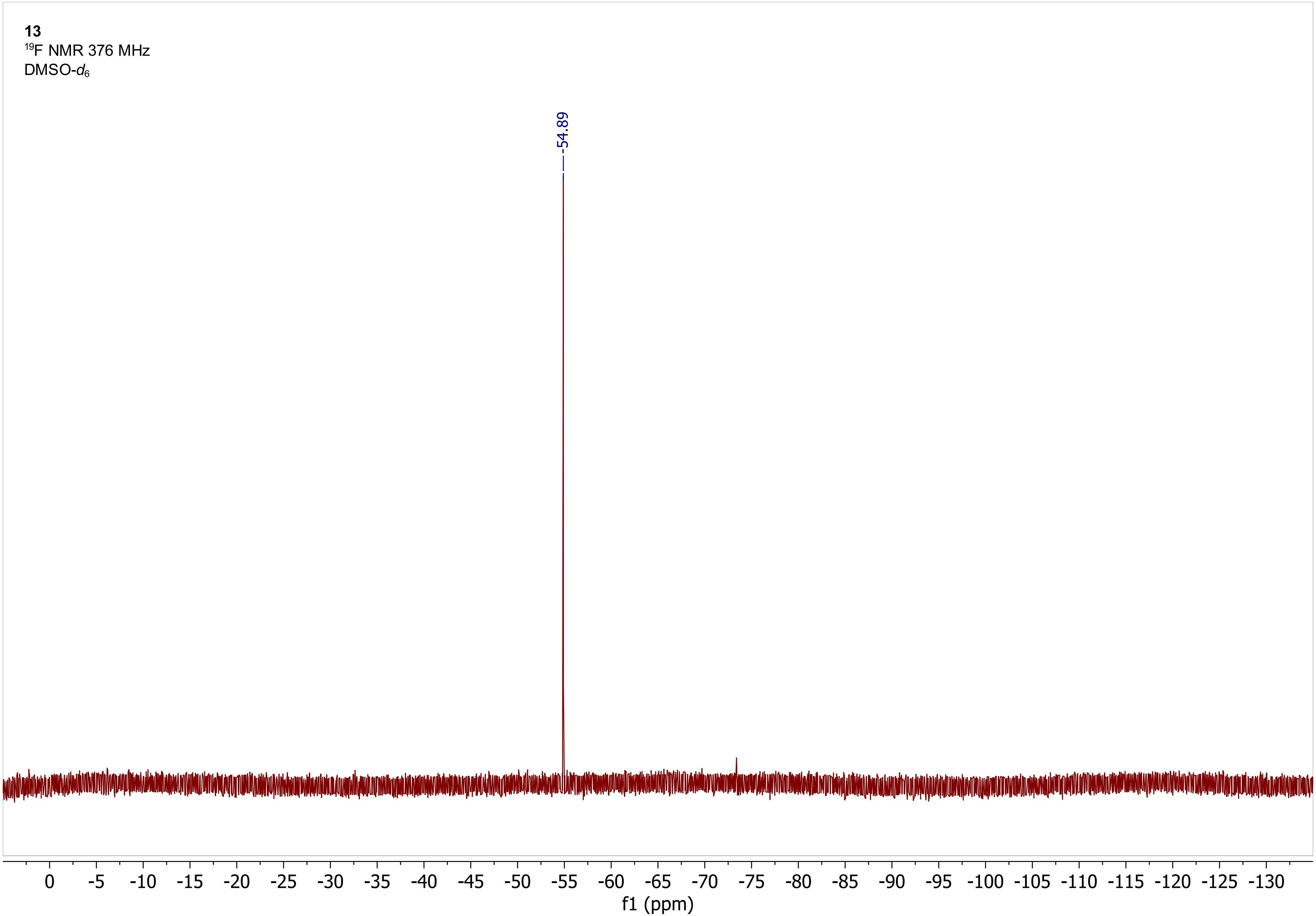
13
$^{19}$F NMR 376 MHz
DMSO-$d_6$
-54.89
0 -5 -10 -15 -20 -25 -30 -35 -40 -45 -50 -55 -60 -65 -70 -75 -80 -85 -90 -95 -100 -105 -110 -115 -120 -125 -130
f1 (ppm)

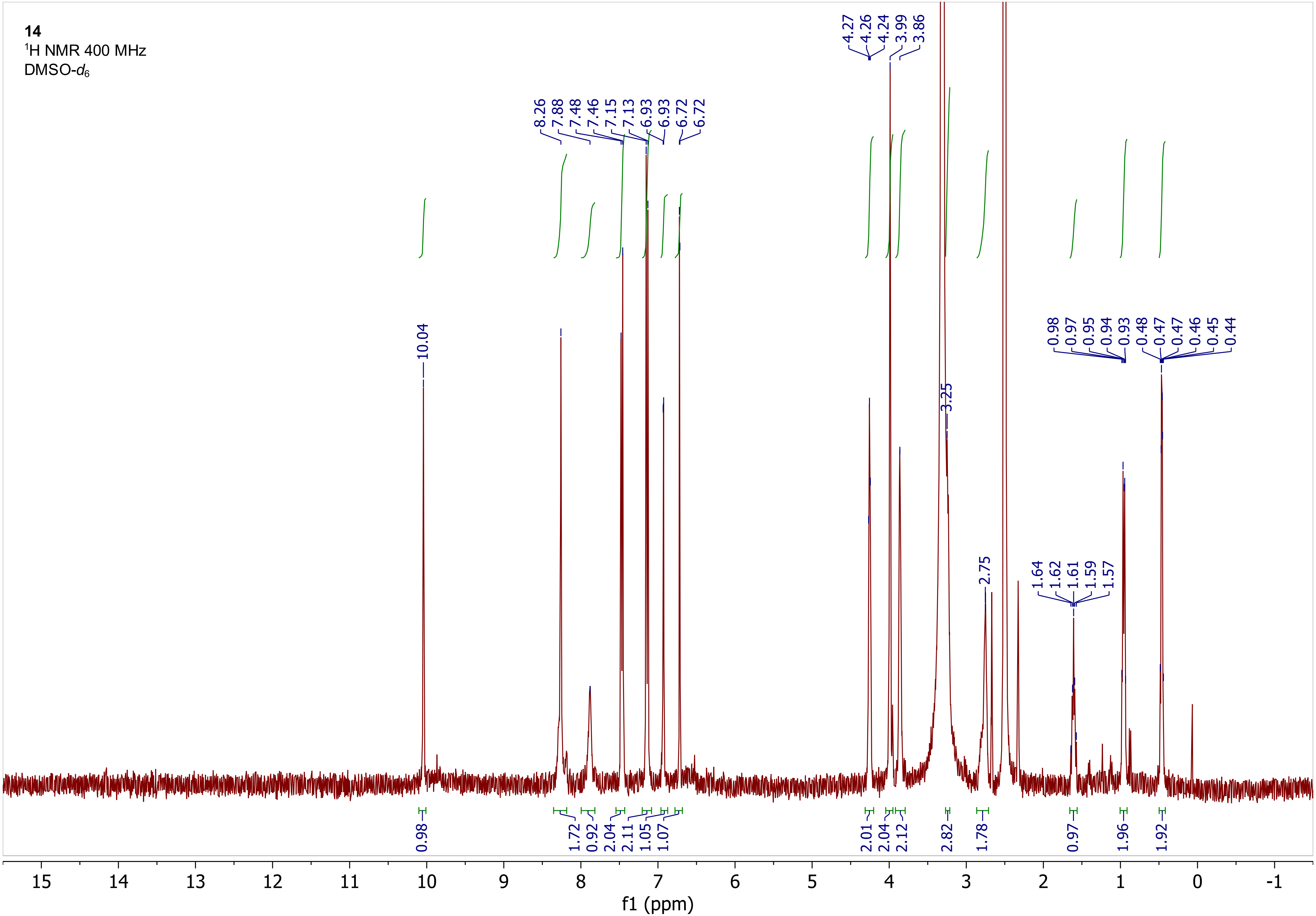
14
$^1$H NMR 400 MHz
DMSO-$d_6$
f1 (ppm)
10.04
8.26
7.88
7.48
7.46
7.15
7.13
6.93
6.93
6.72
6.72
4.27
4.26
4.24
3.99
3.86
3.25
2.75
1.64
1.62
1.61
1.59
1.57
0.98
0.97
0.95
0.94
0.93
0.48
0.47
0.47
0.46
0.45
0.44
0.98
1.72
0.92
2.04
2.11
1.05
1.07
2.01
2.04
2.12
2.82
1.78
0.97
1.96
1.92

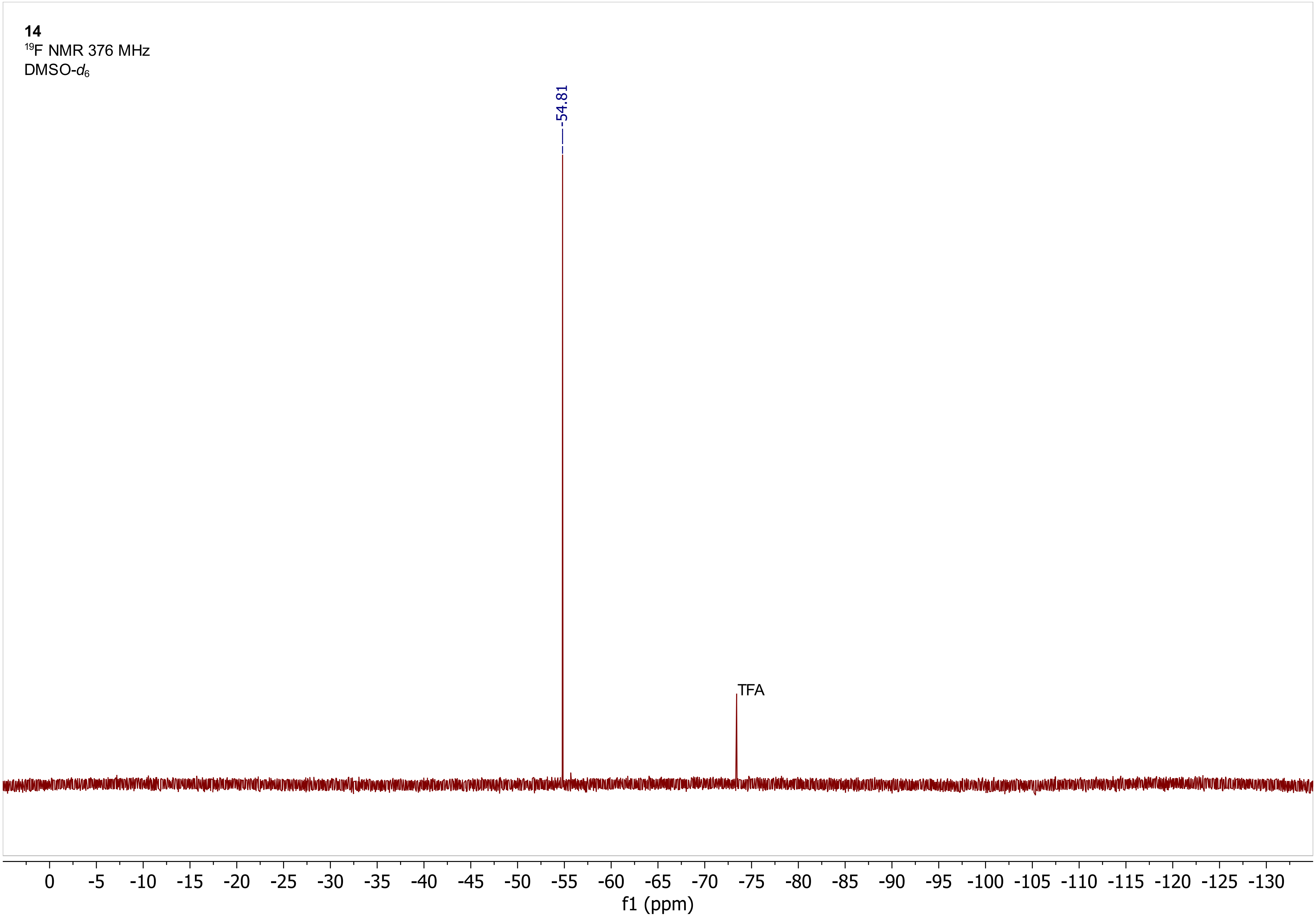
14
$^{19}F$ NMR 376 MHz
DMSO-$d_6$
-54.81
TFA
0 -5 -10 -15 -20 -25 -30 -35 -40 -45 -50 -55 -60 -65 -70 -75 -80 -85 -90 -95 -100 -105 -110 -115 -120 -125 -130
f1 (ppm)

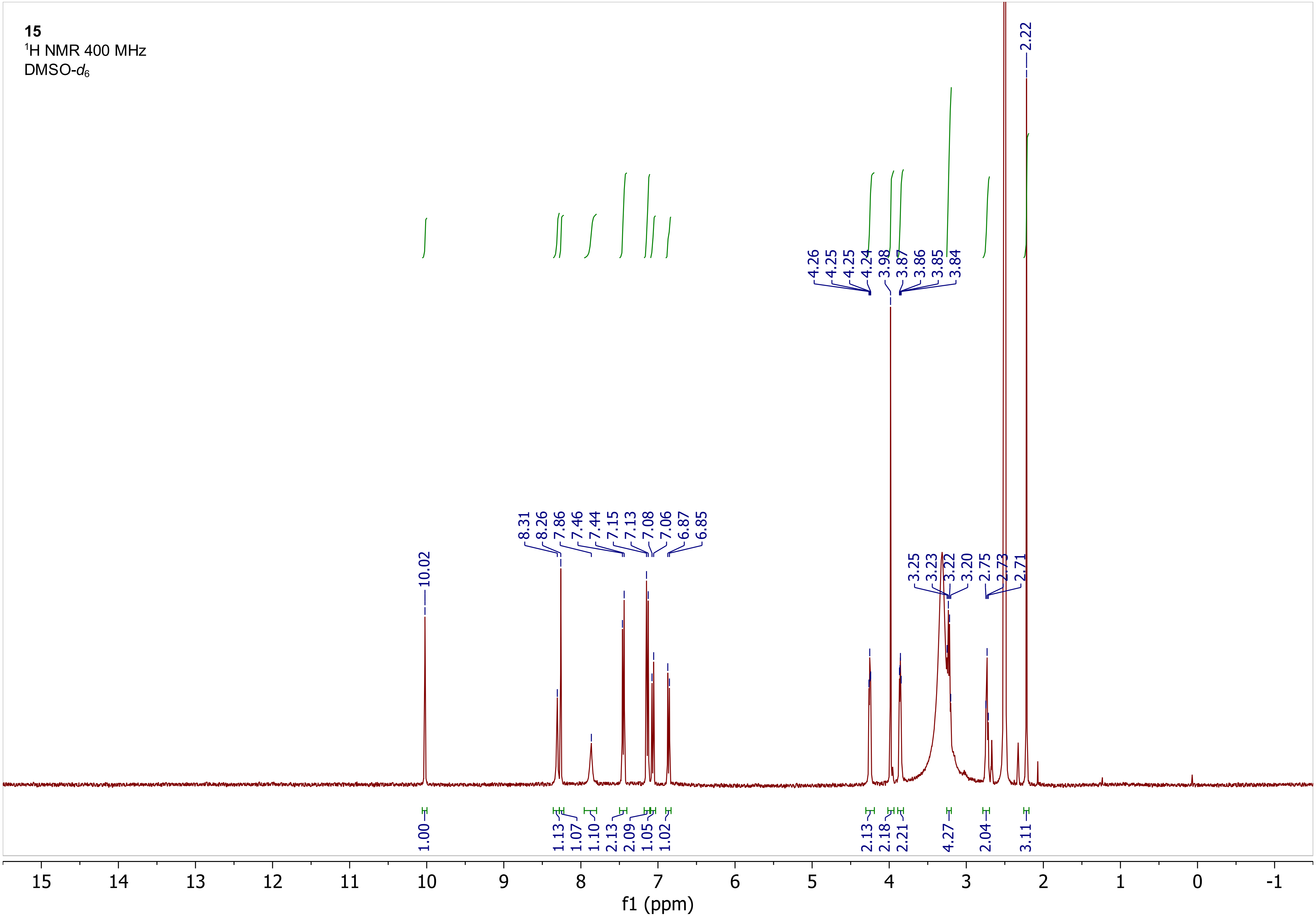

15
1H NMR 400 MHz
DMSO-d6
10.02
8.31
8.26
7.86
7.46
7.44
7.15
7.13
7.08
7.06
6.87
6.85
4.26
4.25
4.25
4.24
3.98
3.87
3.86
3.85
3.84
3.25
3.23
3.22
3.20
2.75
2.73
2.71
2.22
1.00
1.13
1.07
1.10
2.13
2.09
1.05
1.02
2.13
2.18
2.21
4.27
2.04
3.11
15
14
13
12
11
10
9
8
7
6
5
4
3
2
1
0
-1
f1 (ppm)

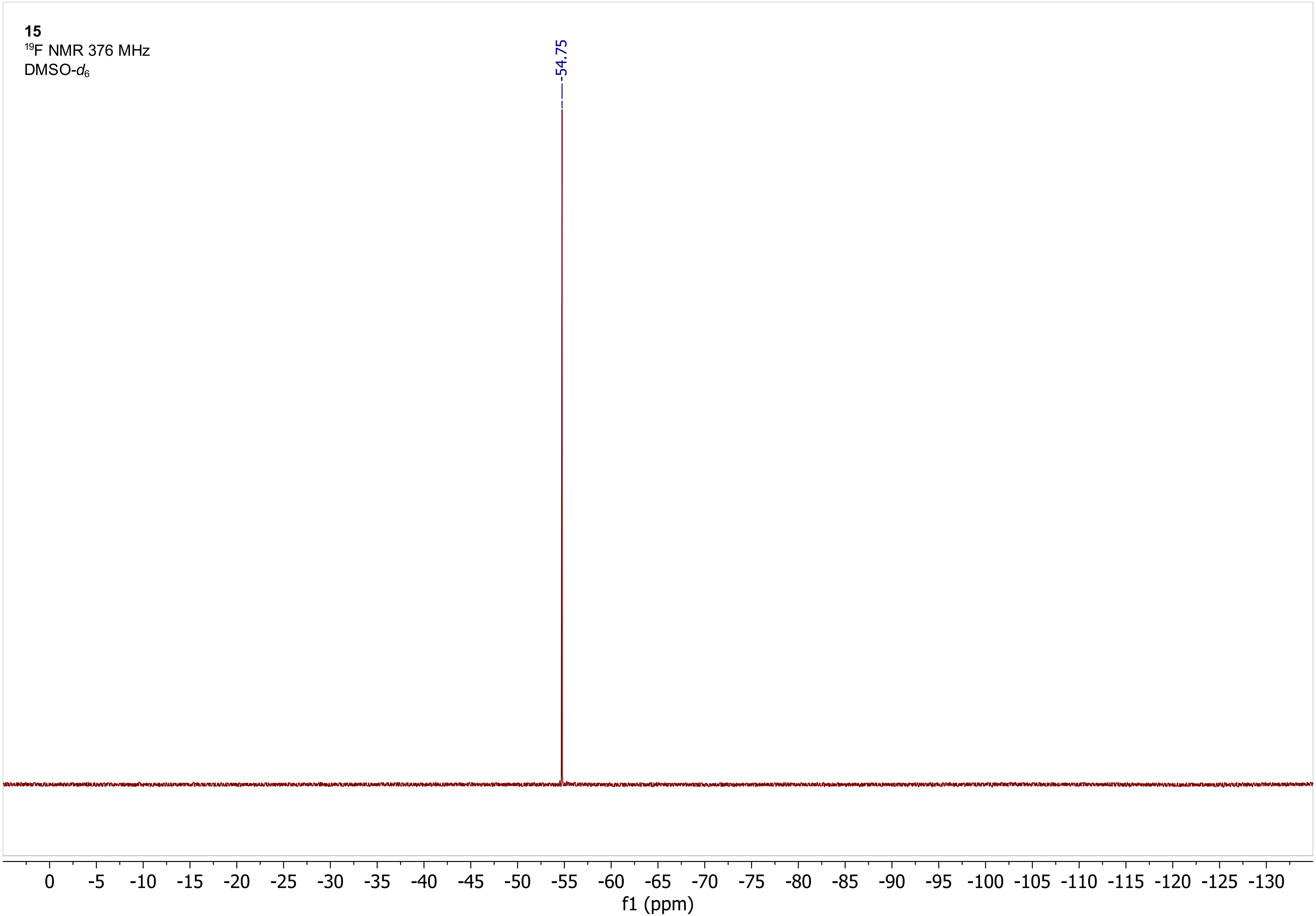
15
19F NMR 376 MHz
DMSO-d6
-54.75
0 -5 -10 -15 -20 -25 -30 -35 -40 -45 -50 -55 -60 -65 -70 -75 -80 -85 -90 -95 -100 -105 -110 -115 -120 -125 -130
f1 (ppm)

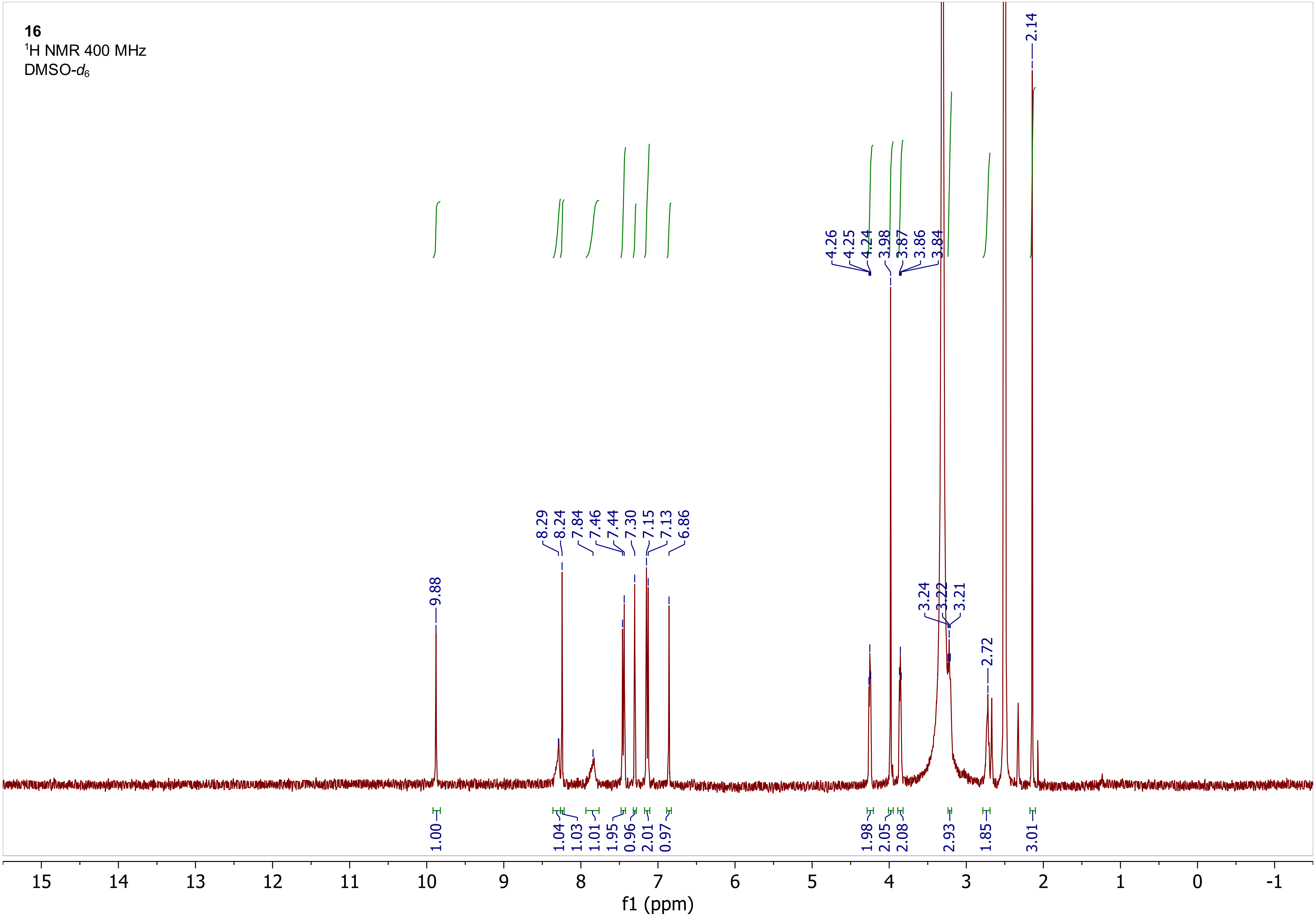

16
¹H NMR 400 MHz
DMSO-d6
9.88
8.29
8.24
7.84
7.46
7.44
7.30
7.15
7.13
6.86
4.26
4.25
4.24
3.98
3.87
3.86
3.84
3.24
3.22
3.21
2.72
2.14
1.00
1.04
1.03
1.01
1.95
0.96
2.01
0.97
1.98
2.05
2.08
2.93
1.85
3.01
15 14 13 12 11 10 9 8 7 6 5 4 3 2 1 0 -1
f1 (ppm)

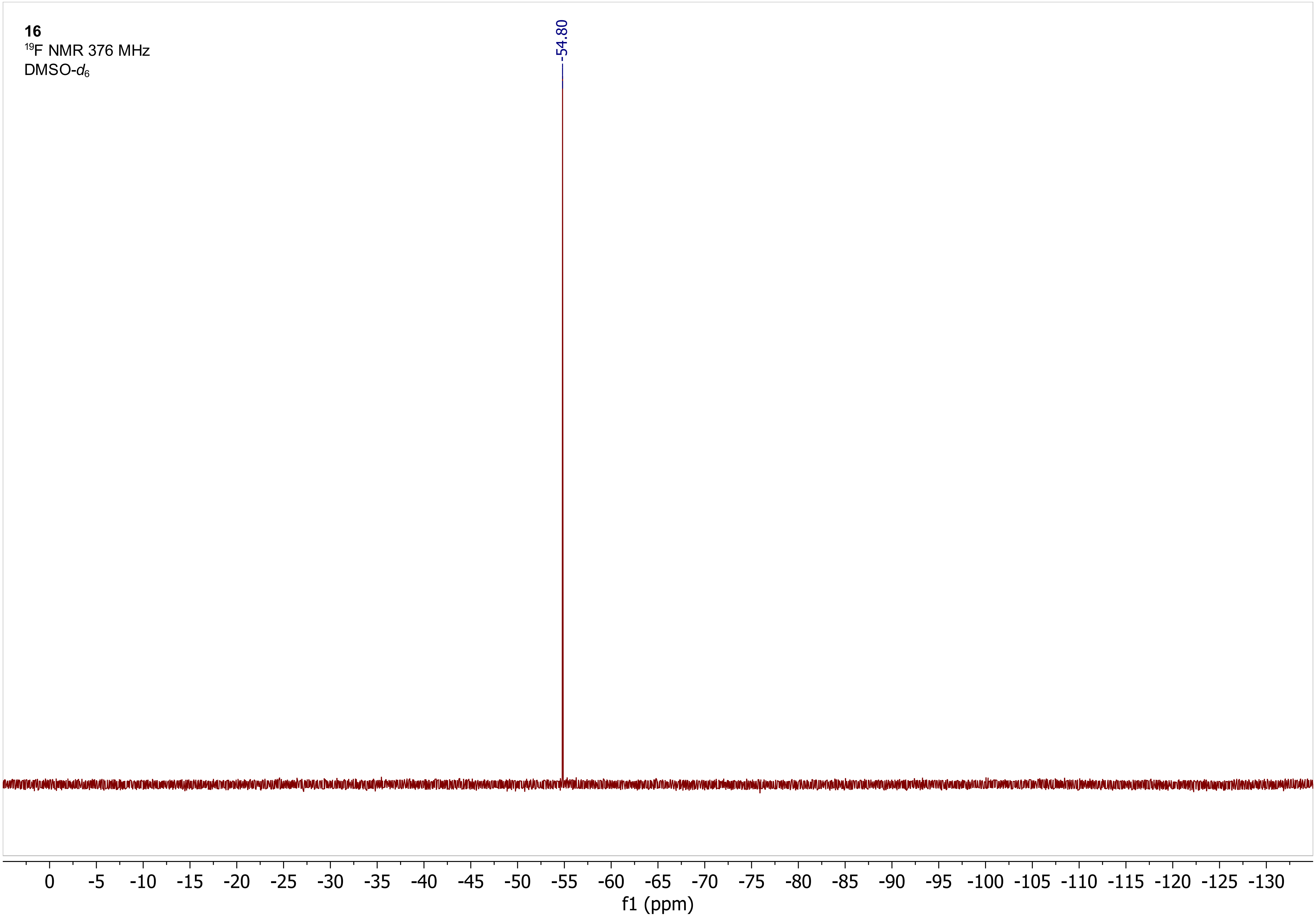

16
$^{19}$F NMR 376 MHz
DMSO-$d_6$
-54.80
0 -5 -10 -15 -20 -25 -30 -35 -40 -45 -50 -55 -60 -65 -70 -75 -80 -85 -90 -95 -100 -105 -110 -115 -120 -125 -130
f1 (ppm)

**17**
$^{1}$H NMR 600 MHz
DMSO-$d_6$

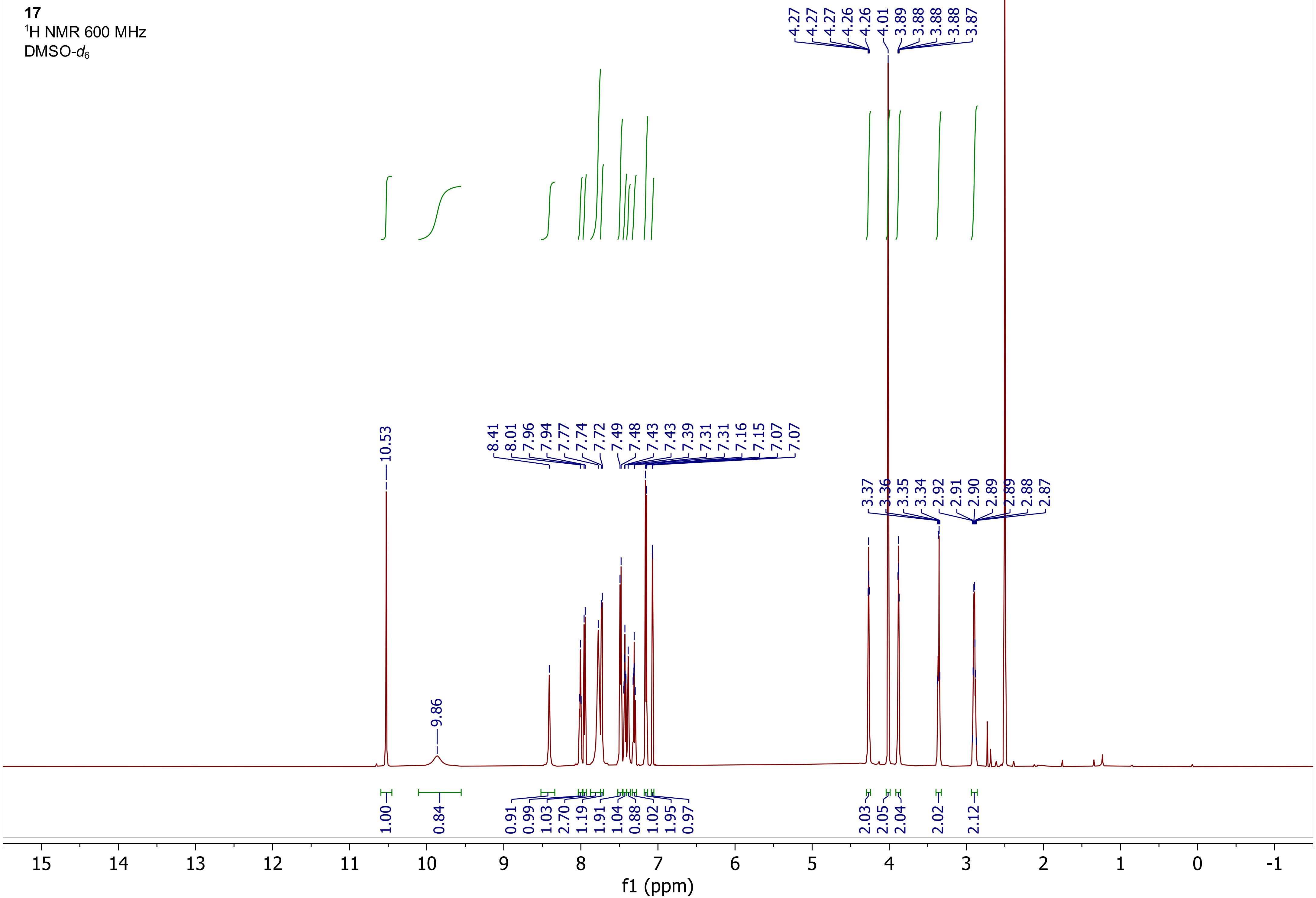

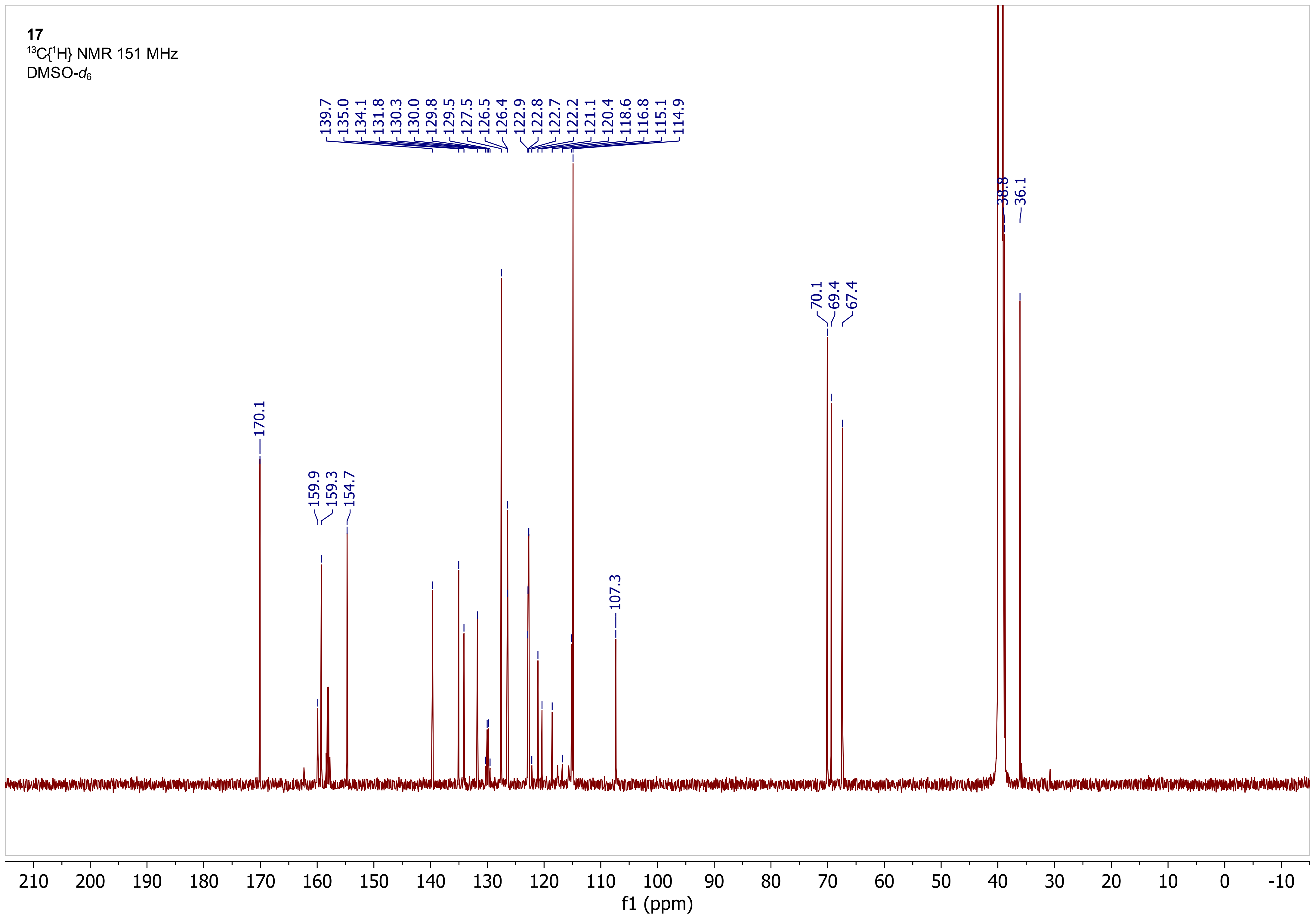
17
13C{1H} NMR 151 MHz
DMSO-d6
139.7
135.0
134.1
131.8
130.3
130.0
129.8
129.5
127.5
126.5
126.4
122.9
122.8
122.7
122.2
121.1
120.4
118.6
116.8
115.1
114.9
170.1
159.9
159.3
154.7
107.3
70.1
69.4
67.4
38.8
36.1
210
200
190
180
170
160
150
140
130
120
110
100
90
80
70
60
50
40
30
20
10
0
-10
f1 (ppm)

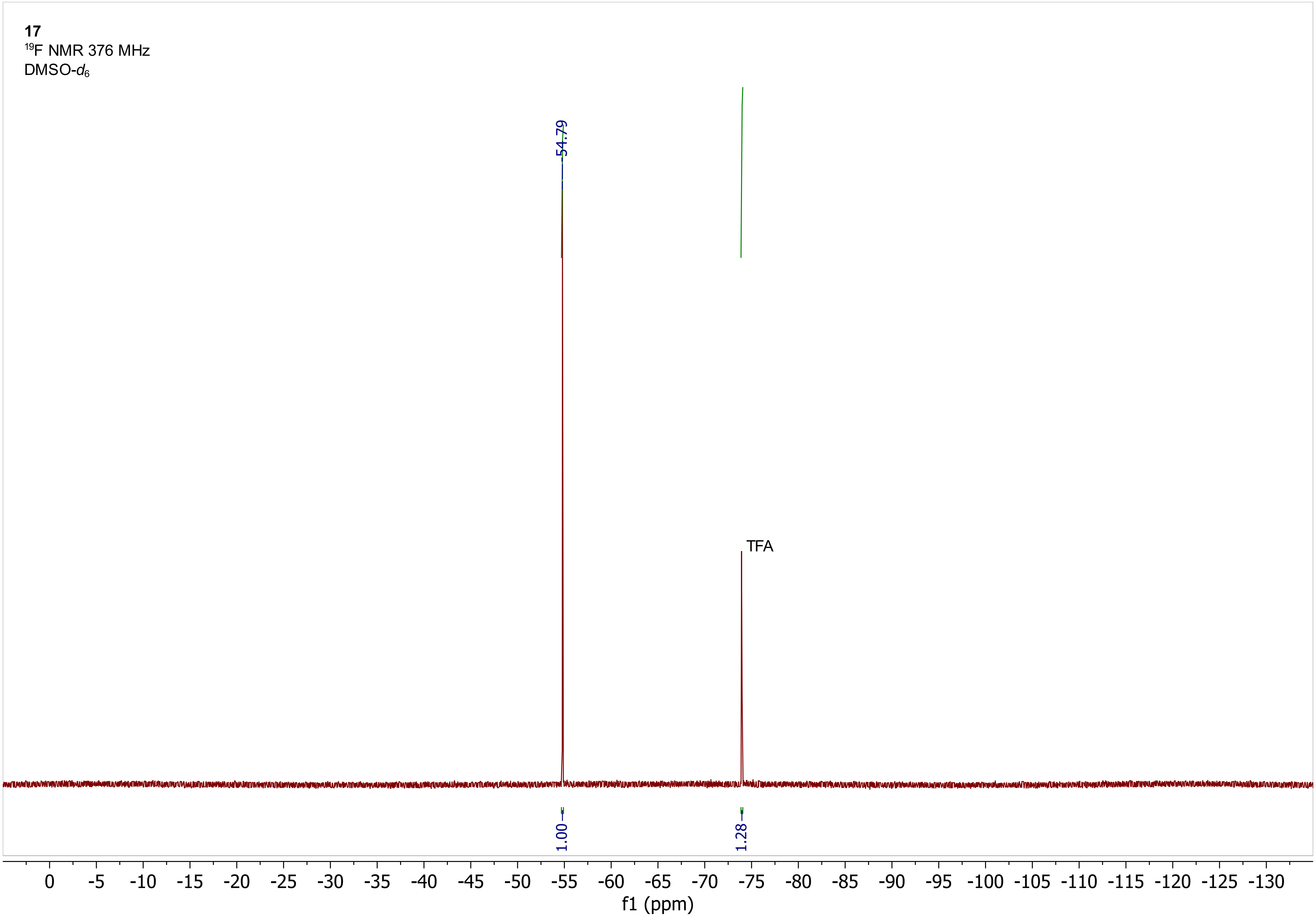
17
19F NMR 376 MHz
DMSO-d6
-54.79
TFA
1.00
1.28
0
-5
-10
-15
-20
-25
-30
-35
-40
-45
-50
-55
-60
-65
-70
-75
-80
-85
-90
-95
-100
-105
-110
-115
-120
-125
-130
f1 (ppm)

**17**
$^1$H COSY 600 MHz
DMSO-$d_6$

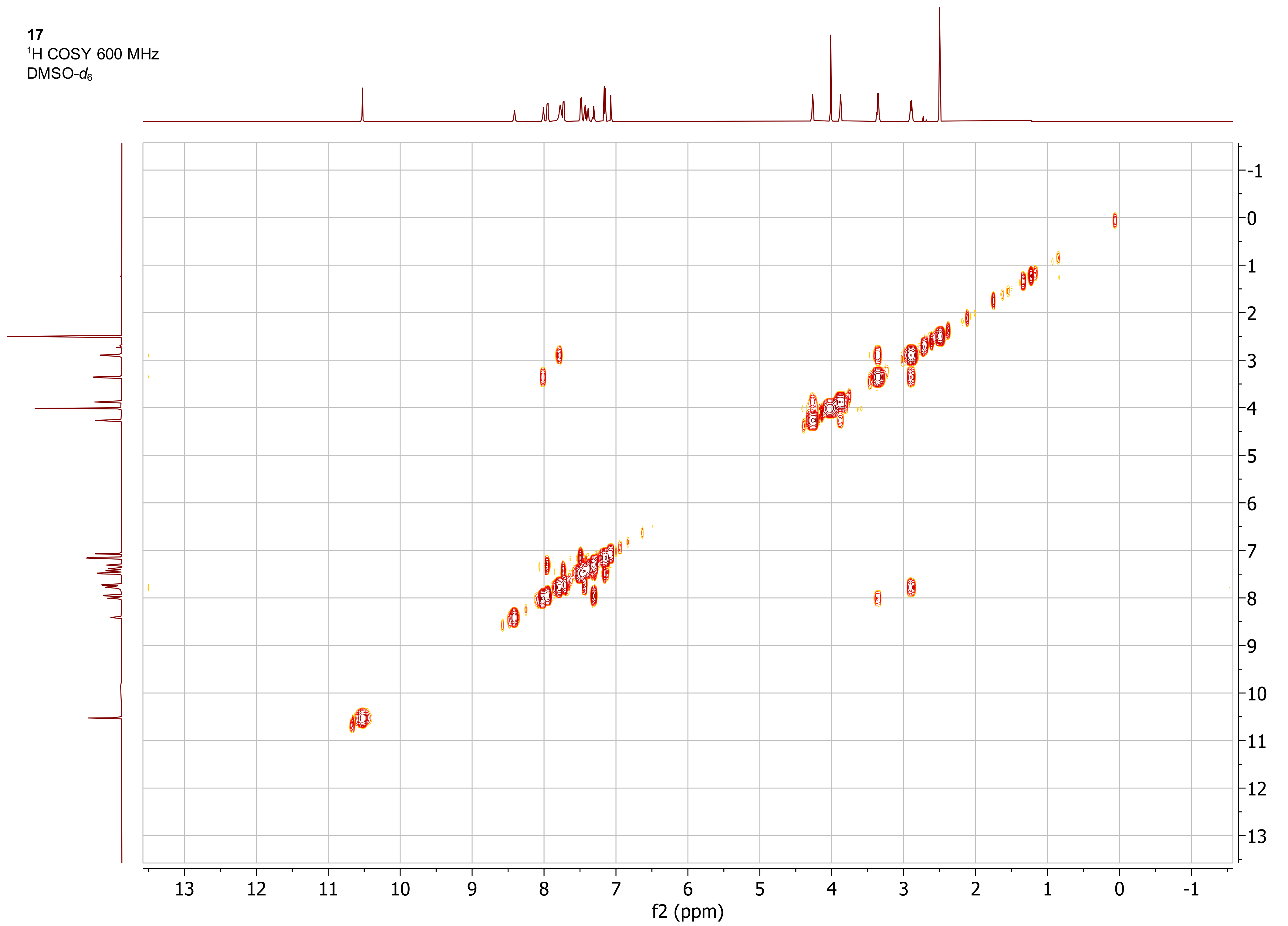

**17**
$^1H$-$^{13}C$ ed-HSQC 600 MHz
DMSO-$d_6$

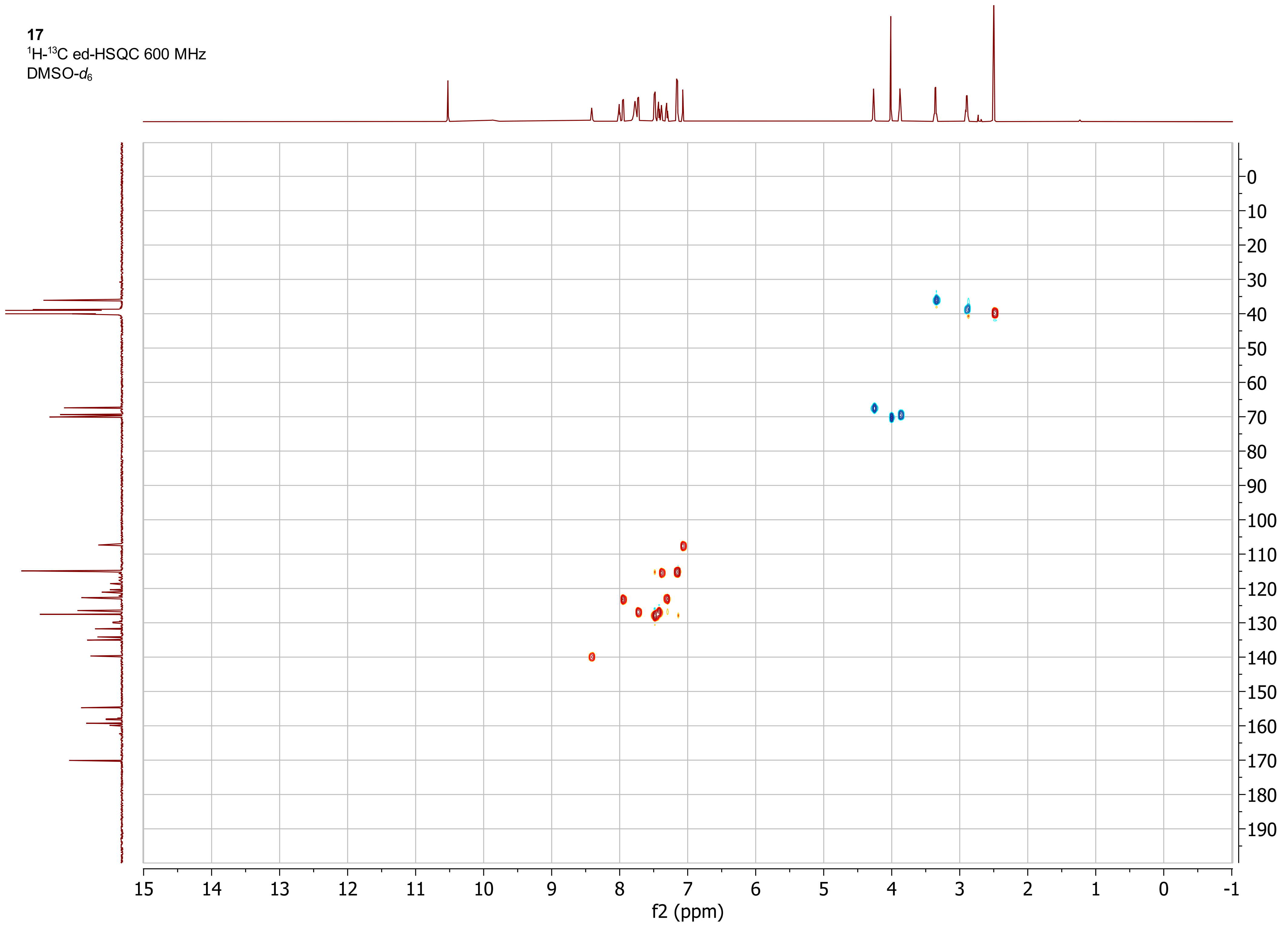

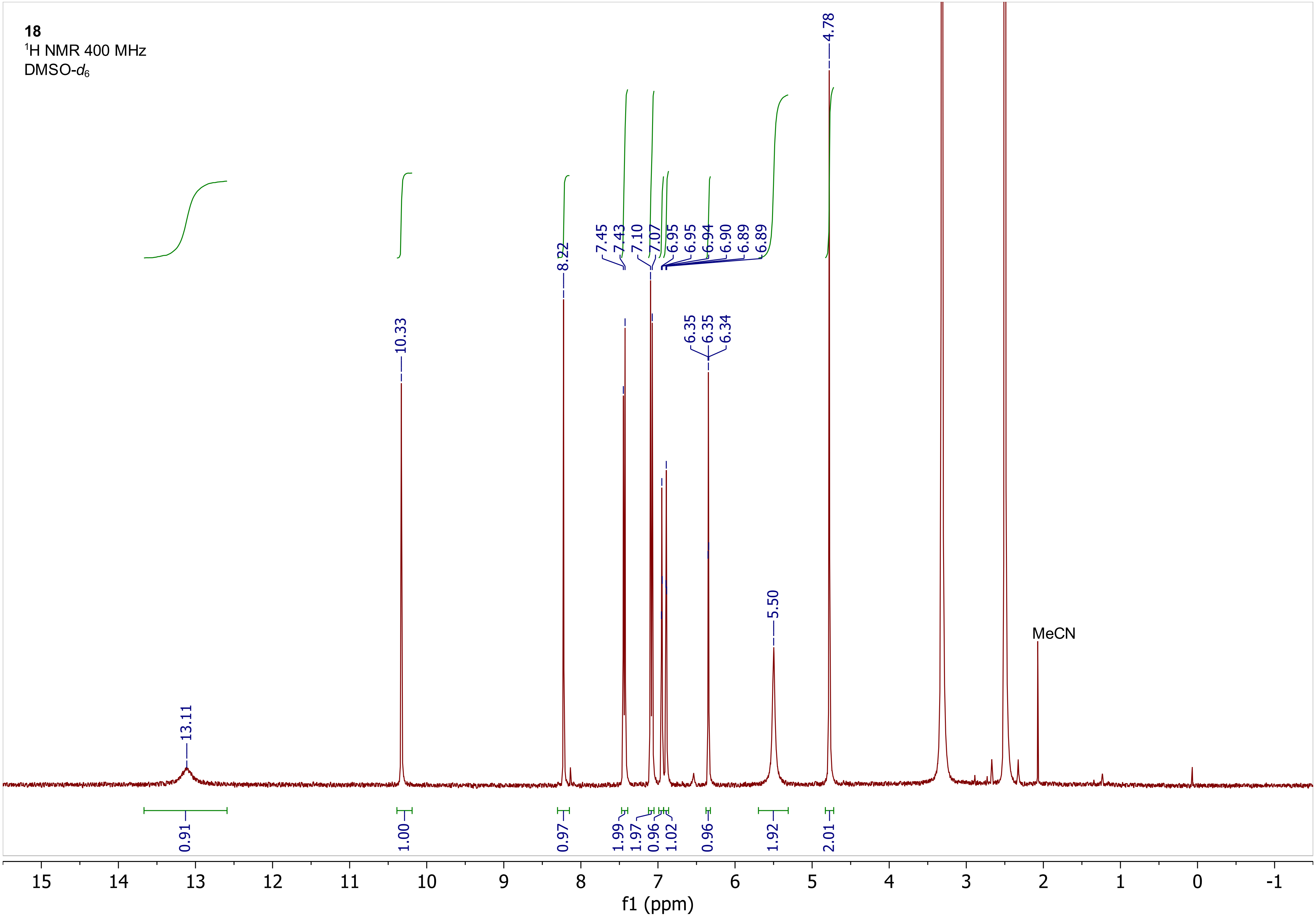

18
1H NMR 400 MHz
DMSO-d6
13.11
10.33
8.22
7.45
7.43
7.10
7.07
6.95
6.95
6.94
6.90
6.89
6.89
6.35
6.35
6.34
5.50
4.78
MeCN
0.91
1.00
0.97
1.99
1.97
0.96
1.02
0.96
1.92
2.01
15
14
13
12
11
10
9
8
7
6
5
4
3
2
1
0
-1
f1 (ppm)

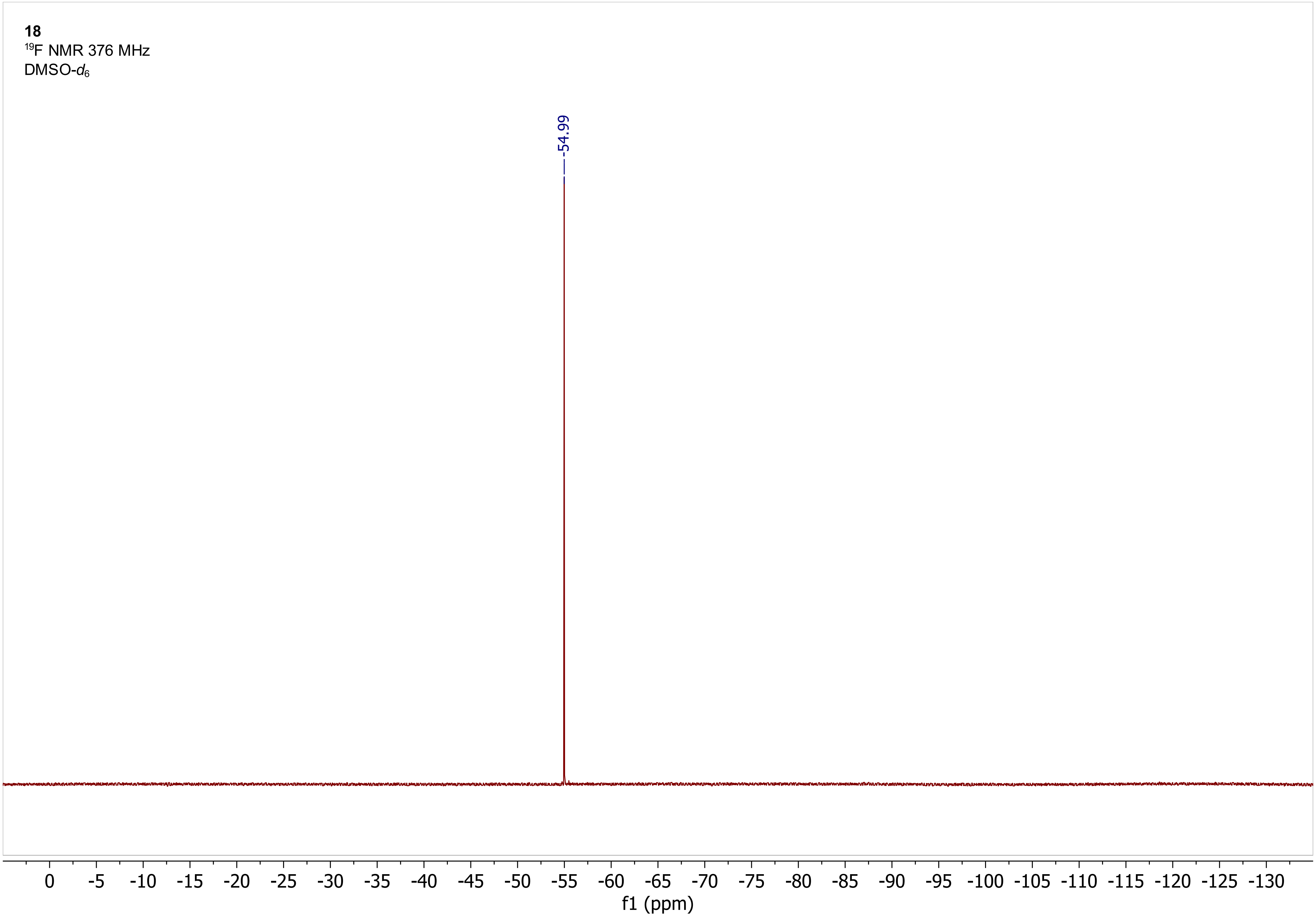
18
$^{19}F$ NMR 376 MHz
DMSO-$d_6$
-54.99
0 -5 -10 -15 -20 -25 -30 -35 -40 -45 -50 -55 -60 -65 -70 -75 -80 -85 -90 -95 -100 -105 -110 -115 -120 -125 -130
f1 (ppm)

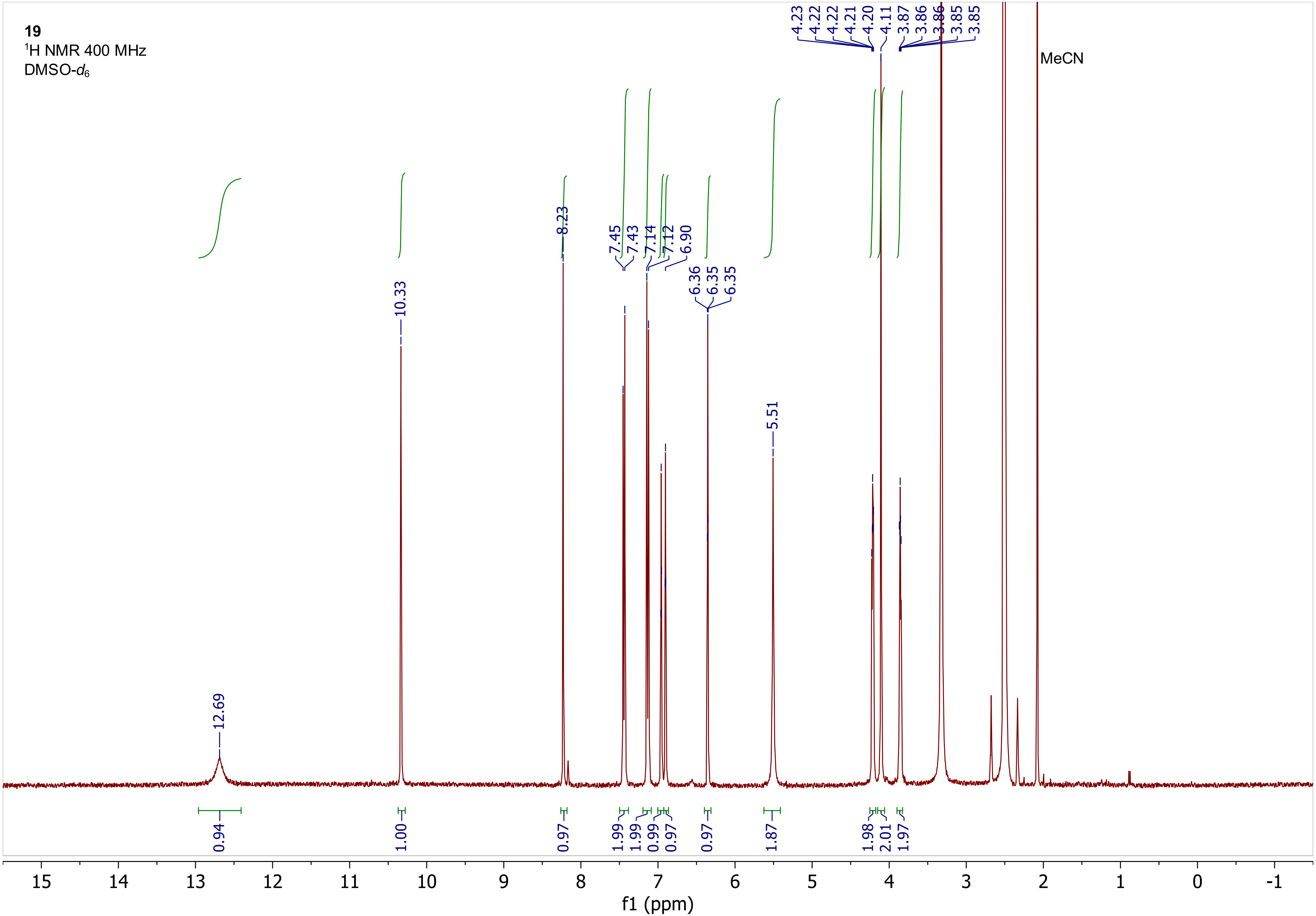
19
¹H NMR 400 MHz
DMSO-d6
MeCN
12.69
10.33
8.23
7.45
7.43
7.14
7.12
6.90
6.36
6.35
6.35
5.51
4.23
4.22
4.22
4.21
4.20
4.11
3.87
3.86
3.86
3.85
3.85
0.94
1.00
0.97
1.99
1.99
0.99
0.97
0.97
1.87
1.98
2.01
1.97
15
14
13
12
11
10
9
8
7
6
5
4
3
2
1
0
-1
f1 (ppm)

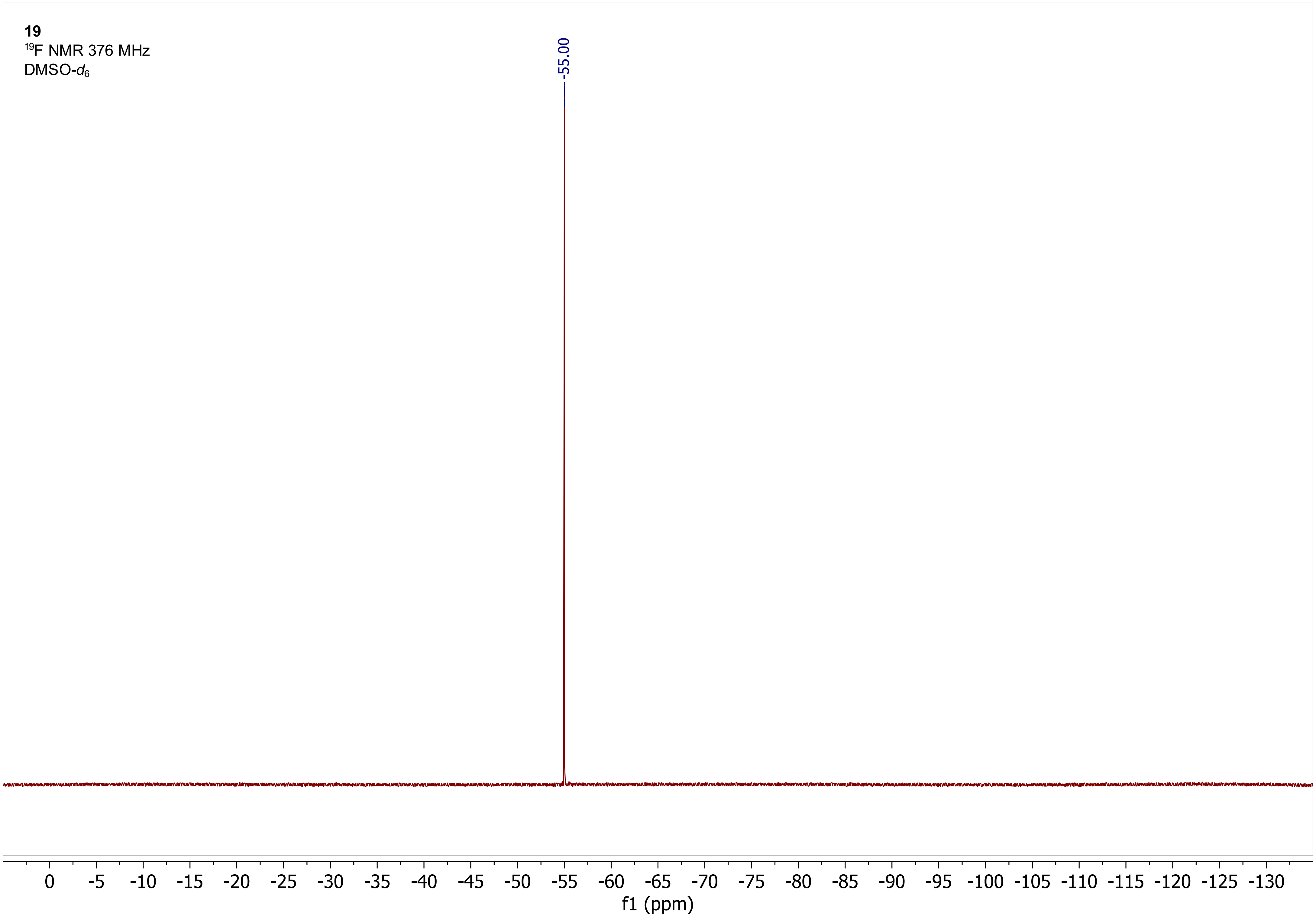
19
19F NMR 376 MHz
DMSO-d6
-55.00
0
-5
-10
-15
-20
-25
-30
-35
-40
-45
-50
-55
-60
-65
-70
-75
-80
-85
-90
-95
-100
-105
-110
-115
-120
-125
-130
f1 (ppm)

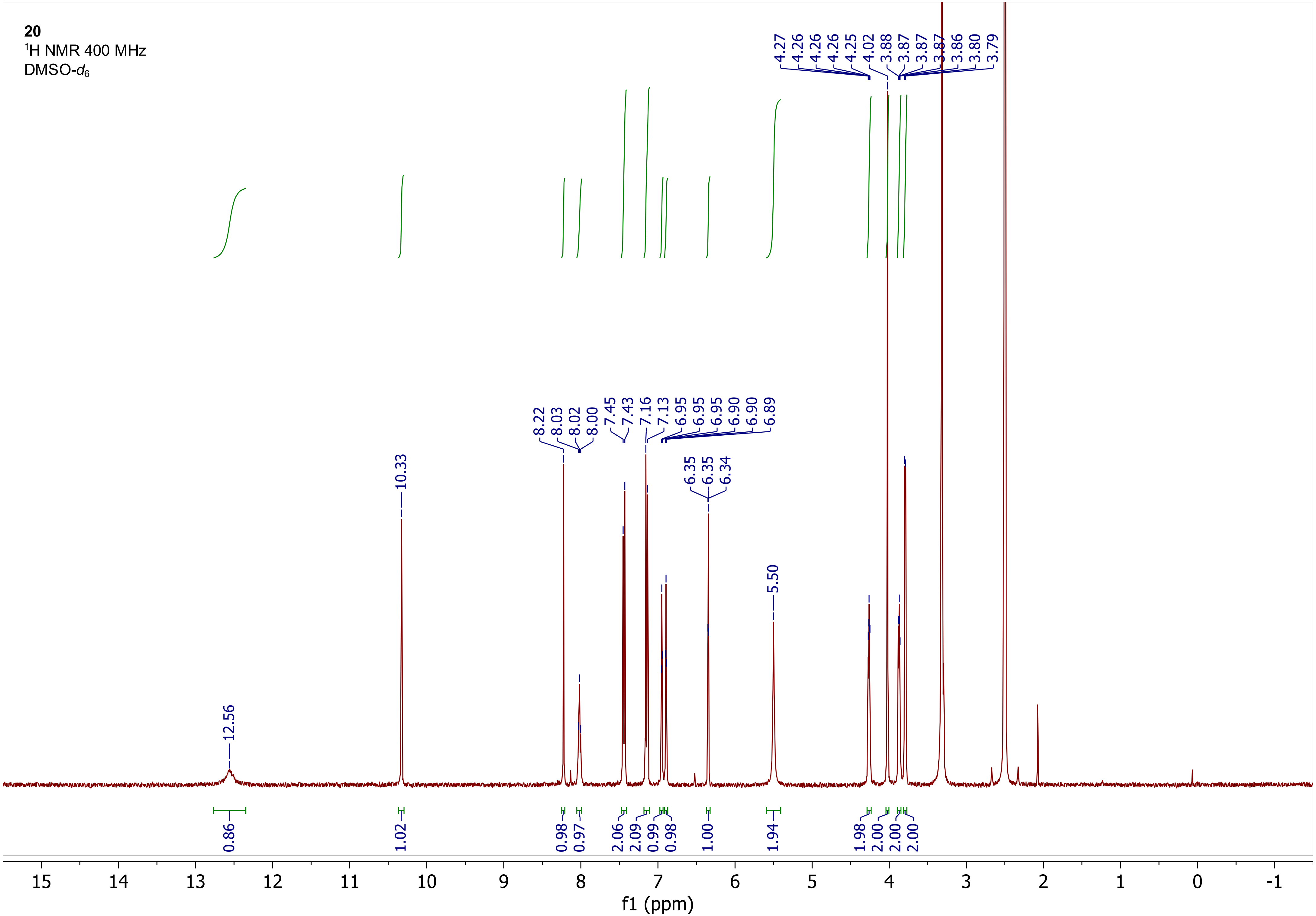

20
1H NMR 400 MHz
DMSO-d6
4.27 4.26 4.26 4.26 4.25 4.02 3.88 3.87 3.87 3.87 3.86 3.80 3.79
8.22 8.03 8.02 8.00 7.45 7.43 7.16 7.13 6.95 6.95 6.95 6.90 6.90 6.89
10.33
6.35 6.35 6.34
5.50
12.56
0.86 1.02 0.98 0.97 2.06 2.09 0.99 0.98 1.00 1.94 1.98 2.00 2.00 2.00
15 14 13 12 11 10 9 8 7 6 5 4 3 2 1 0 -1
f1 (ppm)

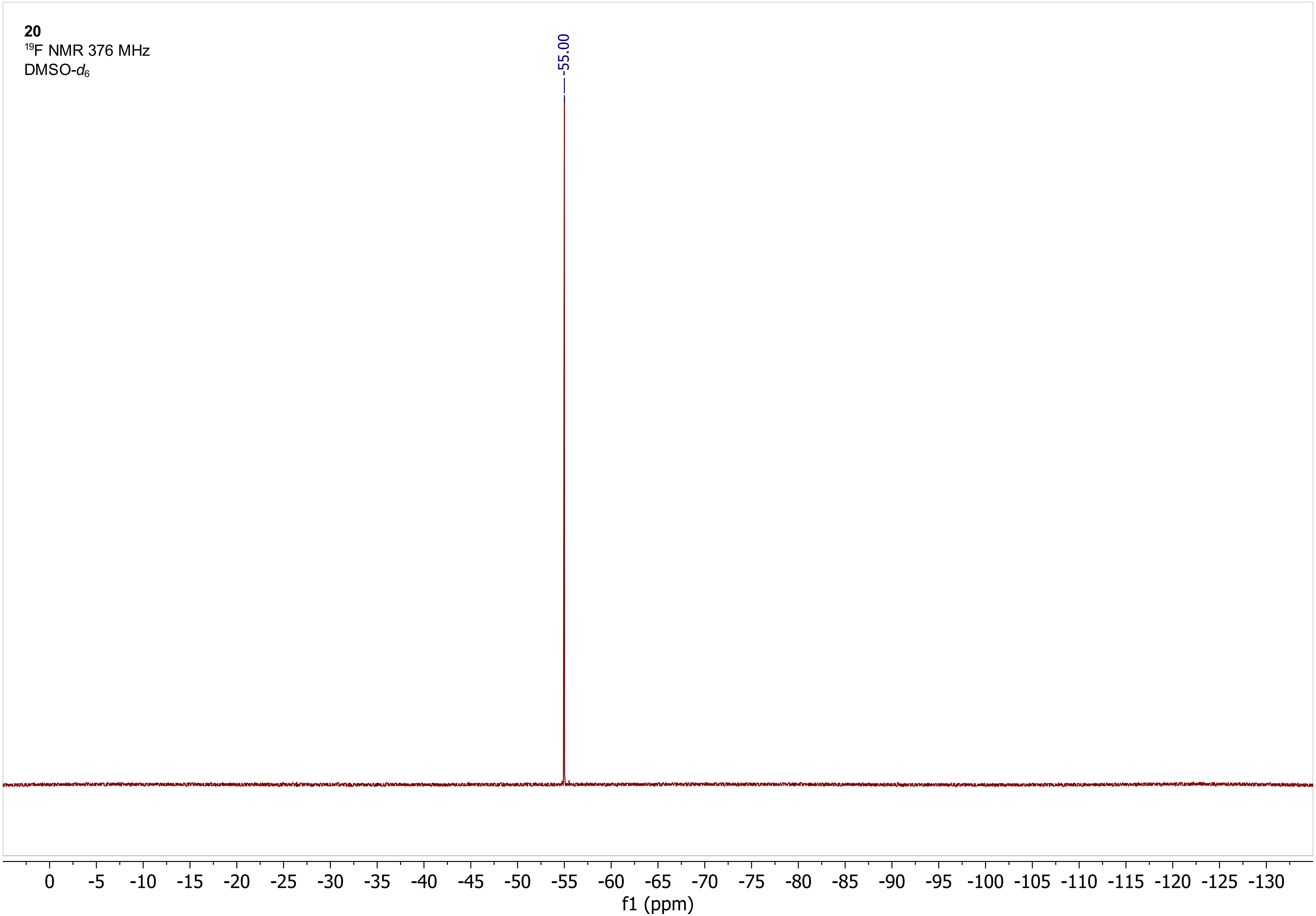

20
19F NMR 376 MHz
DMSO-d6
-55.00
0
-5
-10
-15
-20
-25
-30
-35
-40
-45
-50
-55
-60
-65
-70
-75
-80
-85
-90
-95
-100
-105
-110
-115
-120
-125
-130
f1 (ppm)

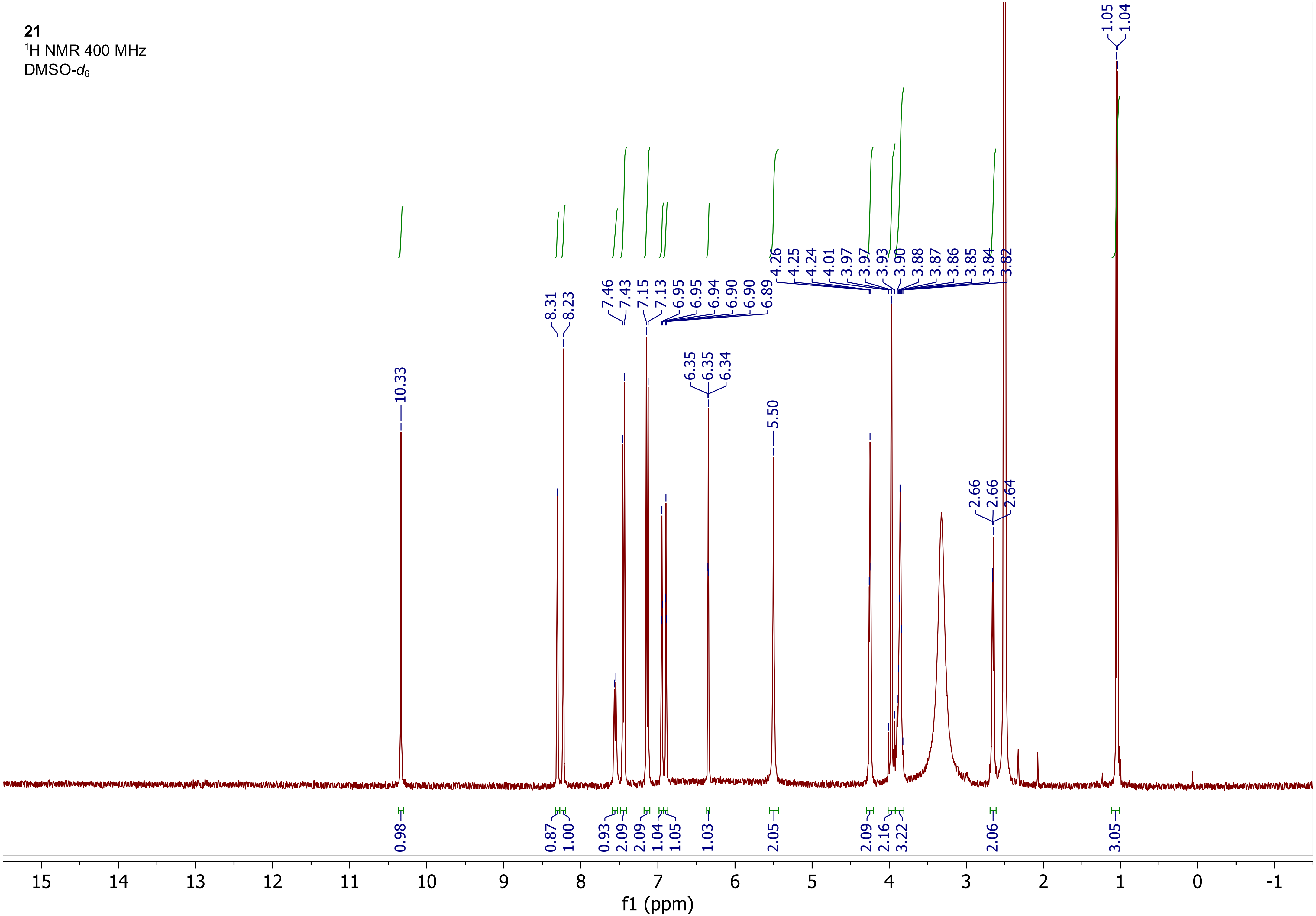
21
1H NMR 400 MHz
DMSO-d6
10.33
8.31
8.23
7.46
7.43
7.15
7.13
6.95
6.95
6.94
6.90
6.90
6.89
6.35
6.35
6.34
5.50
4.26
4.25
4.24
4.01
3.97
3.97
3.93
3.90
3.88
3.87
3.86
3.85
3.84
3.82
2.66
2.66
2.64
1.05
1.04
0.98
0.87
1.00
0.93
2.09
2.09
1.04
1.05
1.03
2.05
2.09
2.16
3.22
2.06
3.05
f1 (ppm)

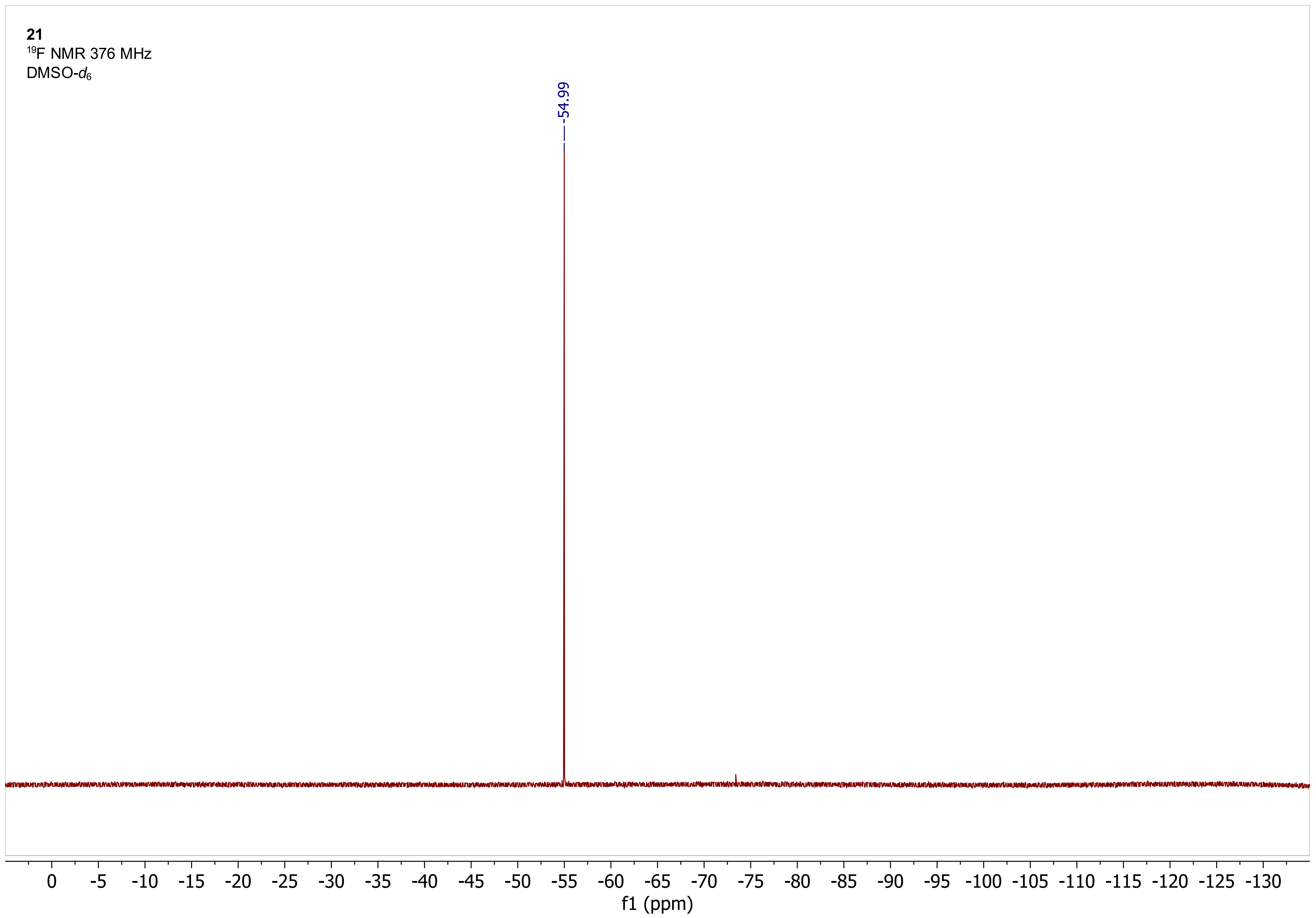
21
19F NMR 376 MHz
DMSO-d6
-54.99
0
-5
-10
-15
-20
-25
-30
-35
-40
-45
-50
-55
-60
-65
-70
-75
-80
-85
-90
-95
-100
-105
-110
-115
-120
-125
-130
f1 (ppm)

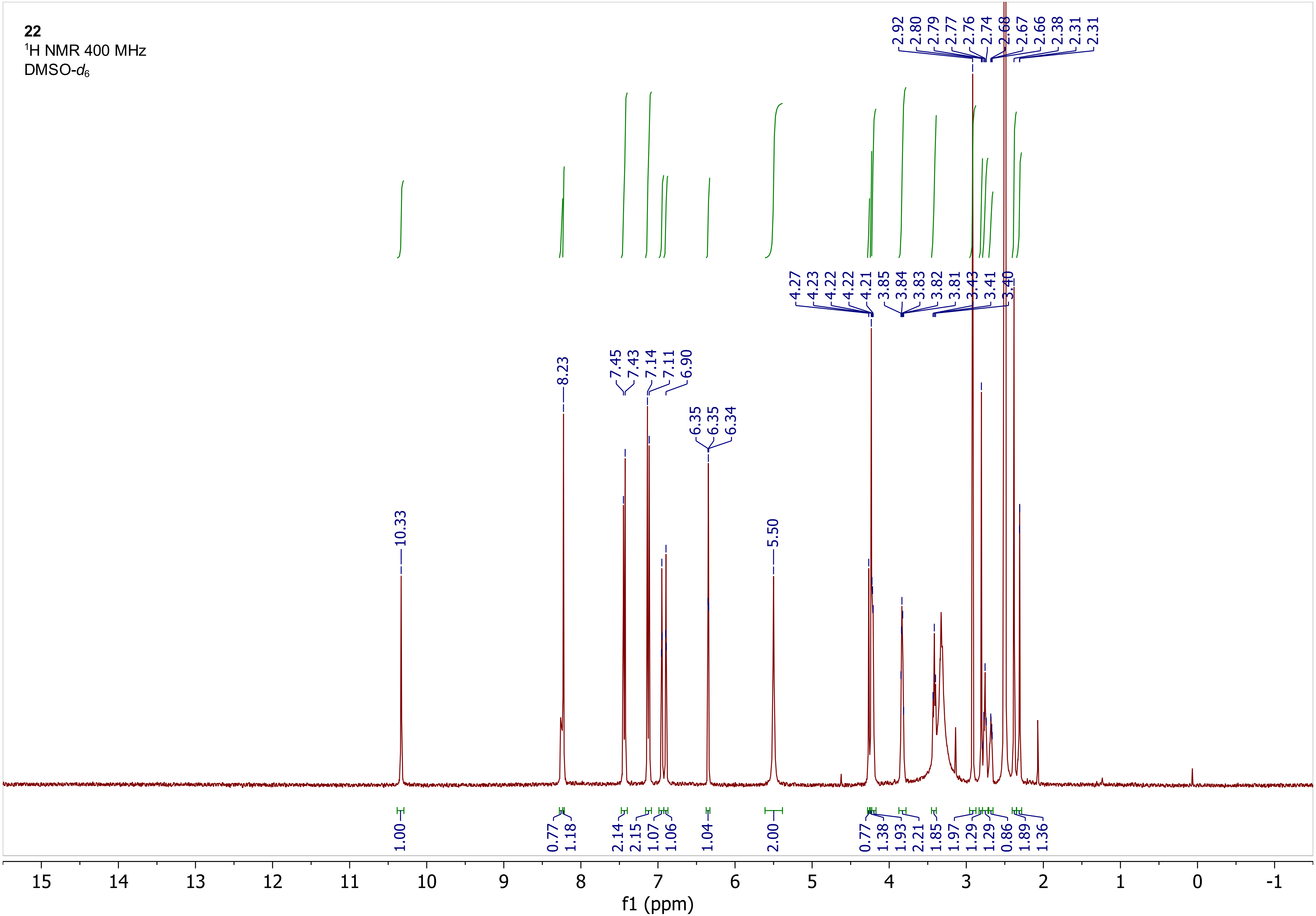
22
1H NMR 400 MHz
DMSO-d6
2.92 2.80 2.79 2.77 2.76 2.74 2.68 2.67 2.66 2.38 2.31 2.31
4.27 4.23 4.22 4.22 4.21 3.85 3.84 3.83 3.82 3.81 3.43 3.41 3.40
8.23
7.45 7.43 7.14 7.11 6.90
6.35 6.35 6.34
10.33
5.50
1.00 0.77 1.18 2.14 2.15 1.07 1.06 1.04 2.00 0.77 1.38 1.93 2.21 1.85 1.97 1.29 1.29 0.86 1.89 1.36
15 14 13 12 11 10 9 8 7 6 5 4 3 2 1 0 -1
f1 (ppm)

**22**
$^{19}$F NMR 376 MHz
DMSO-$d_6$

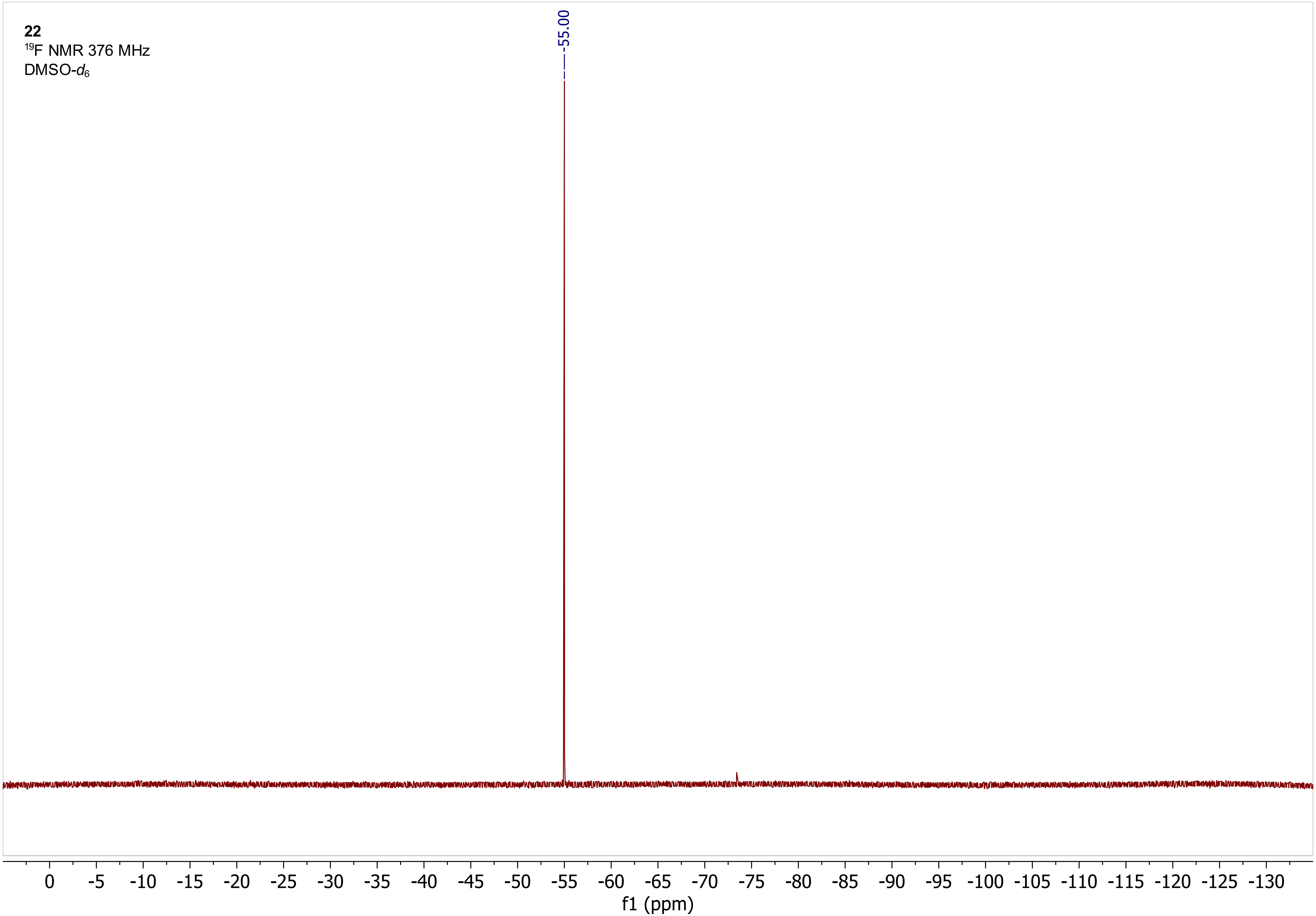

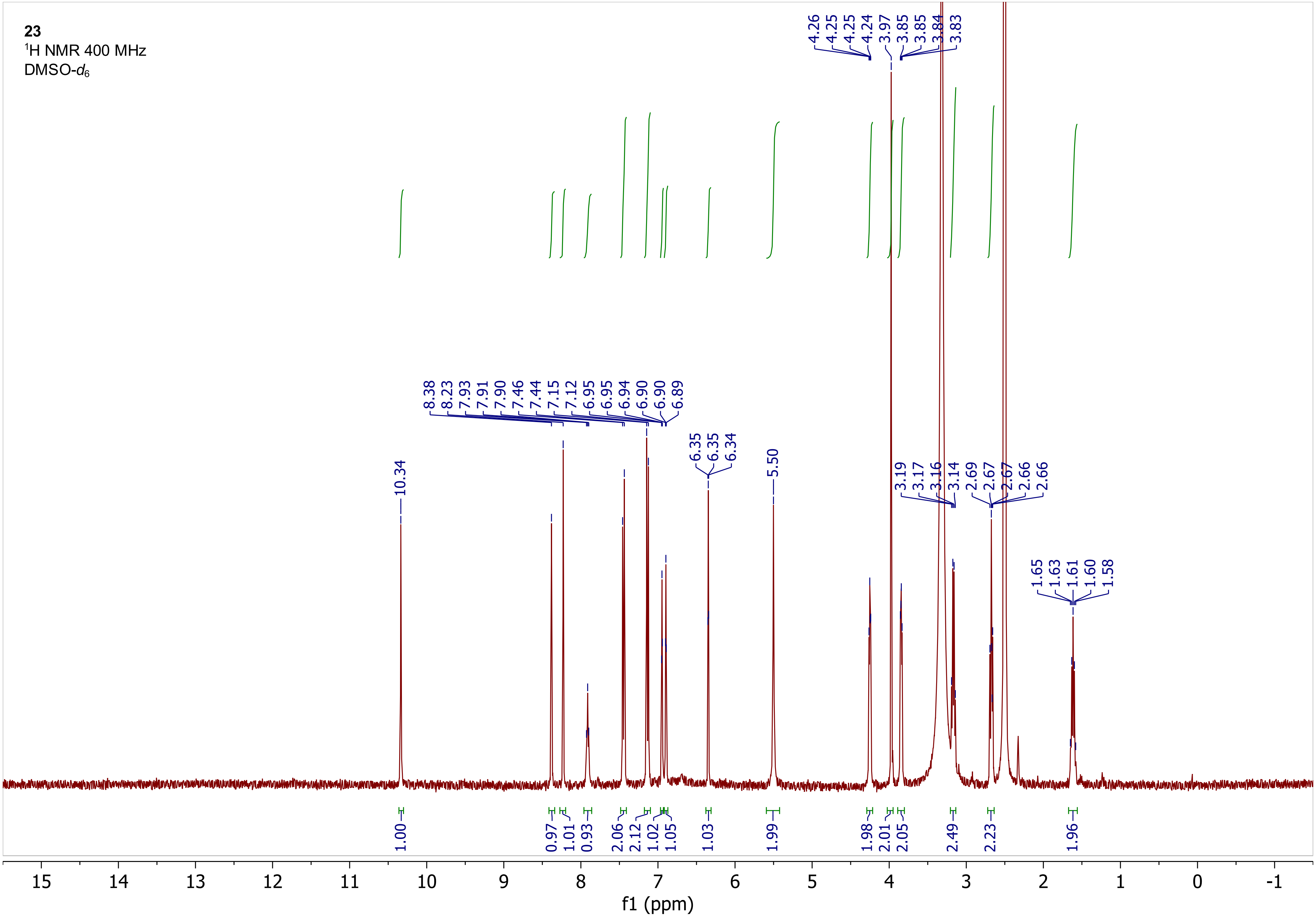

23
$^{1}$H NMR 400 MHz
DMSO-$d_6$
4.26 4.25 4.25 4.24 3.97 3.85 3.85 3.84 3.83
8.38 8.23 7.93 7.91 7.90 7.46 7.44 7.15 7.12 6.95 6.95 6.94 6.90 6.90 6.89
6.35 6.35 6.34
5.50
10.34
3.19 3.17 3.16 3.14
2.69 2.67 2.67 2.66 2.66
1.65 1.63 1.61 1.60 1.58
1.00 0.97 1.01 0.93 2.06 2.12 1.02 1.05 1.03 1.99 1.98 2.01 2.05 2.49 2.23 1.96
f1 (ppm)

**23**
$^{19}F$ NMR 376 MHz
DMSO-$d_6$

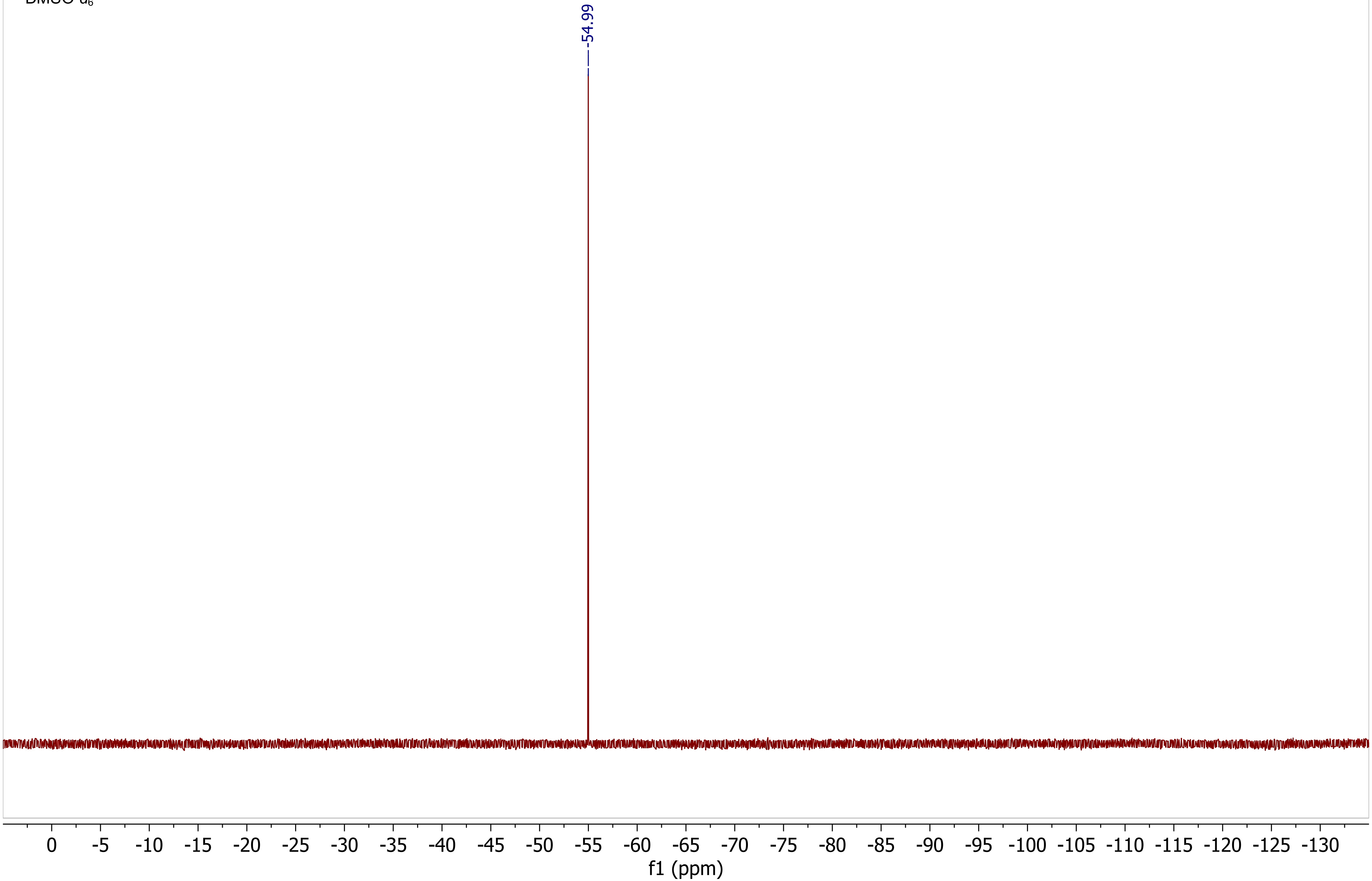

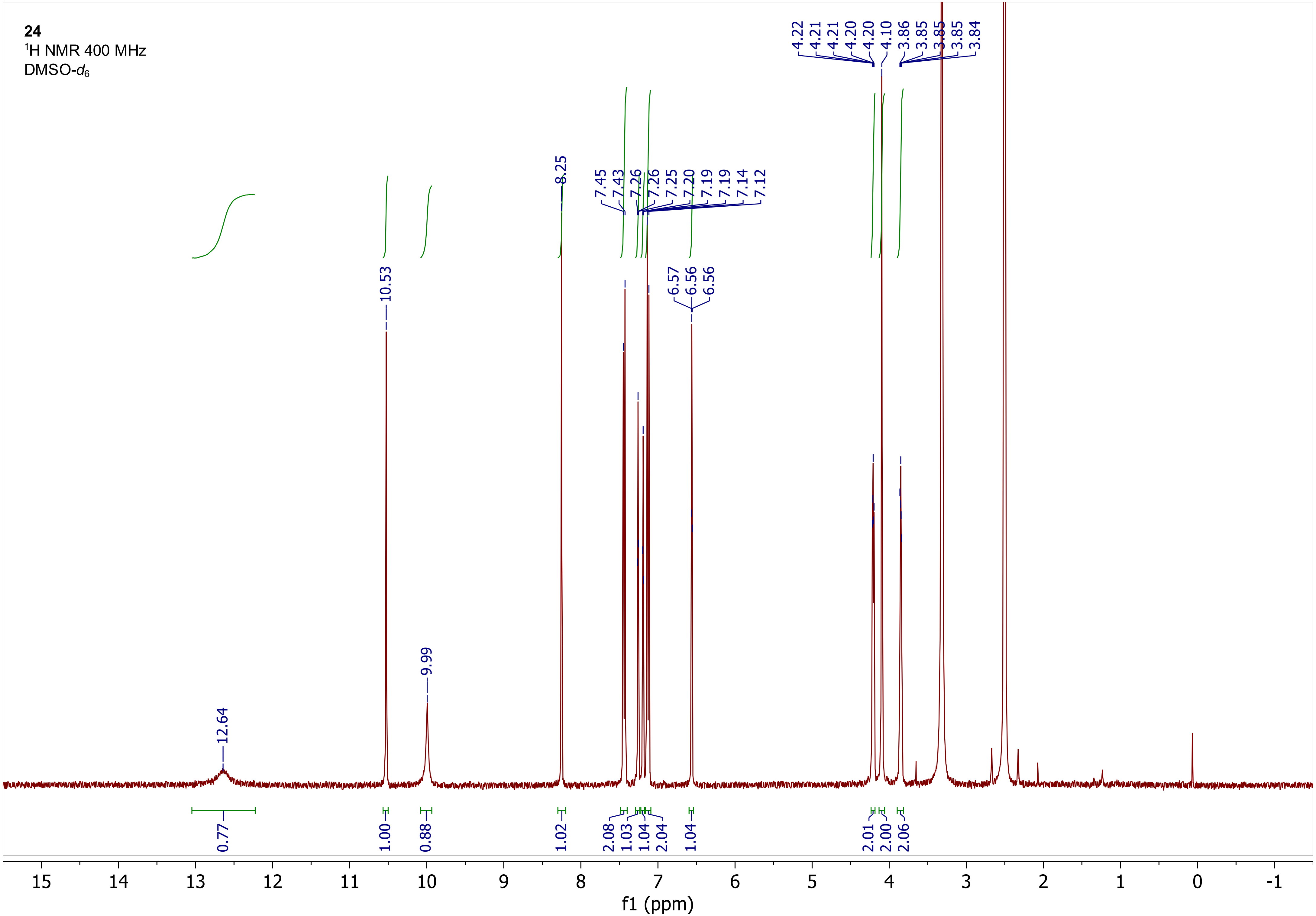

**24**
$^1$H NMR 400 MHz
DMSO-$d_6$
4.22 4.21 4.21 4.20 4.20 4.10 3.86 3.85 3.85 3.85 3.84
8.25
7.45 7.43 7.26 7.26 7.25 7.20 7.19 7.19 7.14 7.12
10.53
6.57 6.56 6.56
9.99
12.64
0.77 1.00 0.88 1.02 2.08 1.03 1.04 2.04 1.04 2.01 2.00 2.06
15 14 13 12 11 10 9 8 7 6 5 4 3 2 1 0 -1
f1 (ppm)

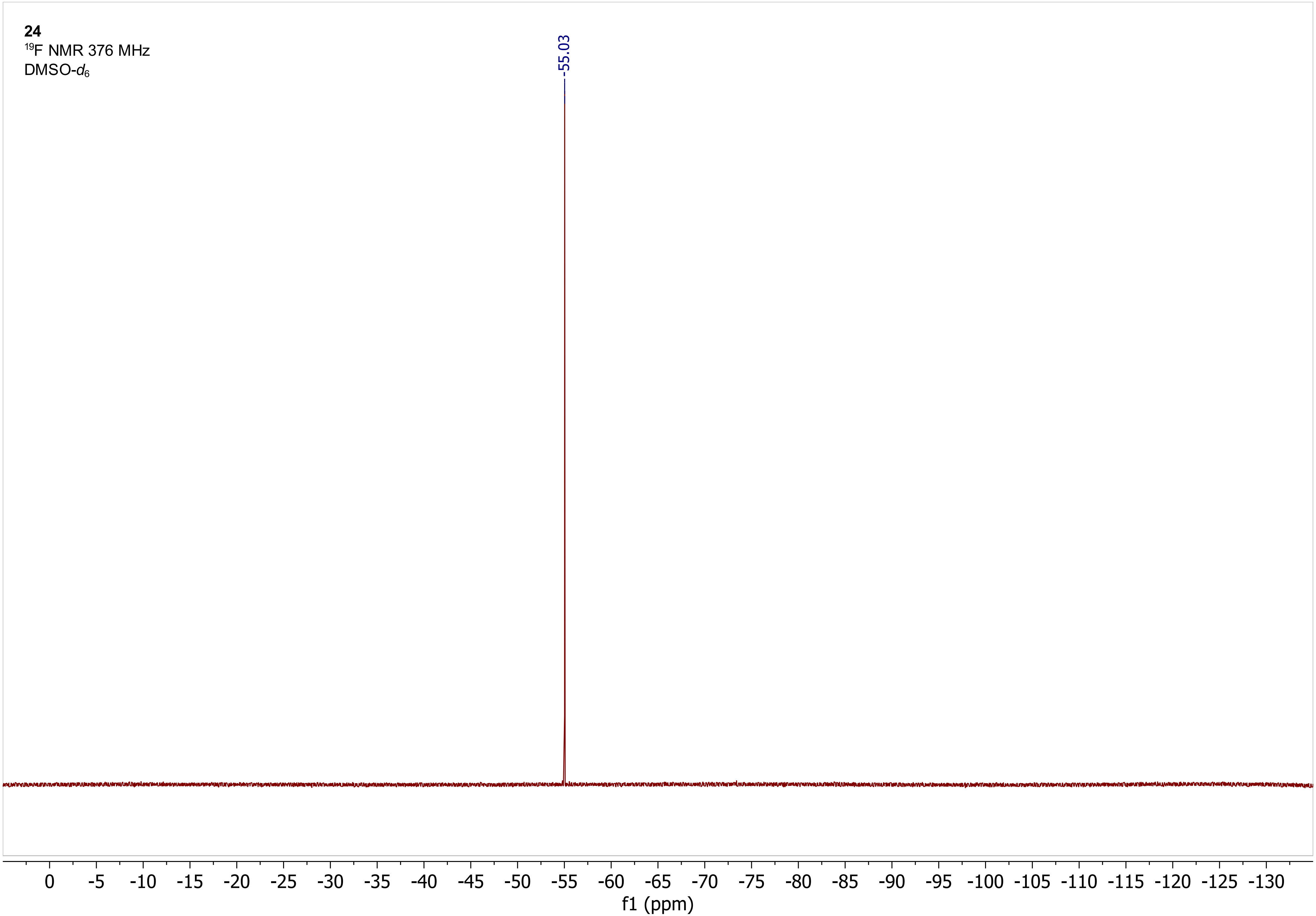
24
$^{19}$F NMR 376 MHz
DMSO-$d_6$
-55.03
0 -5 -10 -15 -20 -25 -30 -35 -40 -45 -50 -55 -60 -65 -70 -75 -80 -85 -90 -95 -100 -105 -110 -115 -120 -125 -130
f1 (ppm)

**25**
$^{1}$H NMR 400 MHz
DMSO-$d_6$

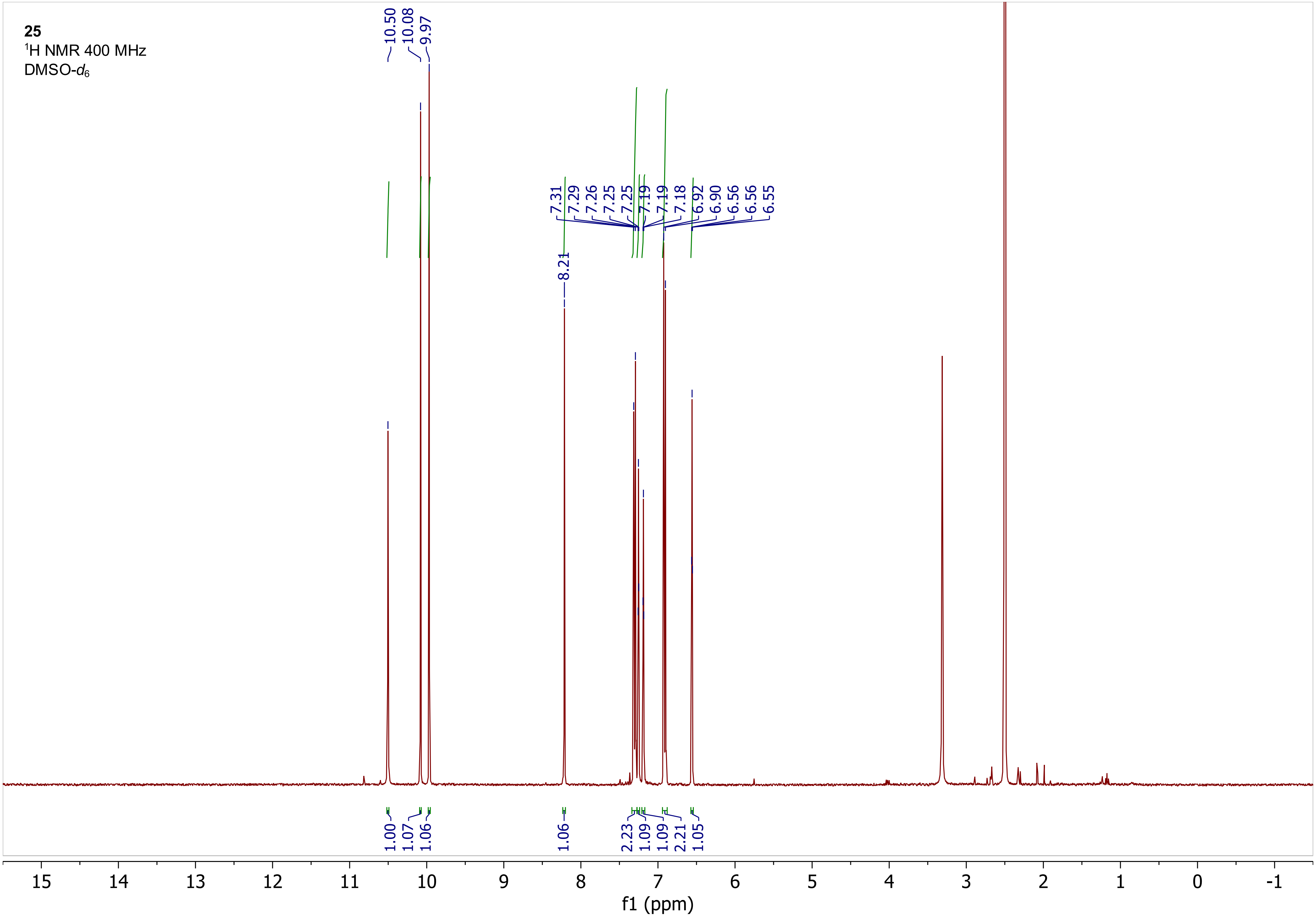

**25**
$^{19}F$ NMR 376 MHz
DMSO-$d_6$

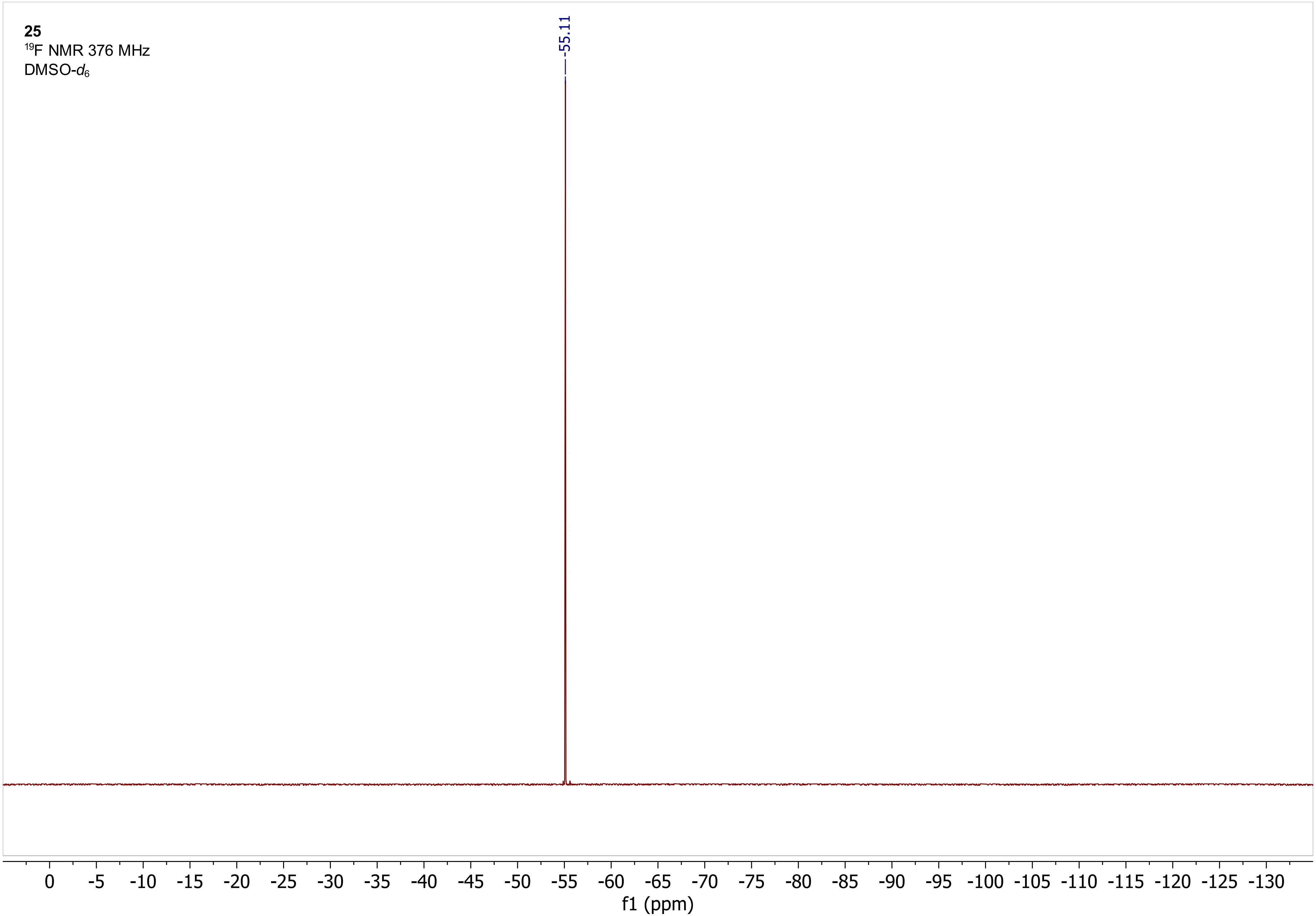

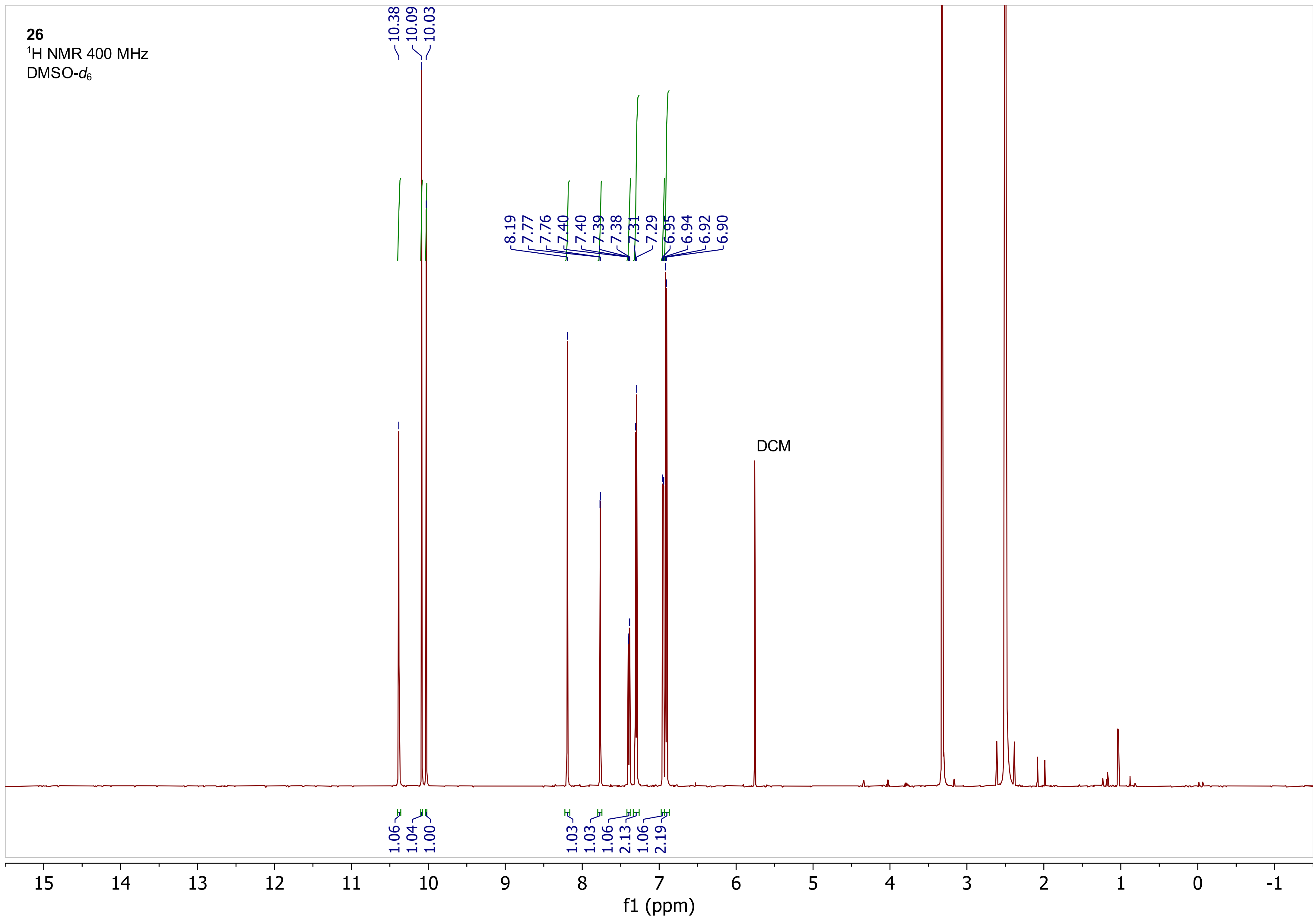

26
¹H NMR 400 MHz
DMSO-d6
10.38
10.09
10.03
8.19
7.77
7.76
7.40
7.40
7.39
7.38
7.31
7.29
6.95
6.94
6.92
6.90
DCM
1.06
1.04
1.00
1.03
1.03
1.06
2.13
1.06
2.19
15
14
13
12
11
10
9
8
7
6
5
4
3
2
1
0
-1
f1 (ppm)

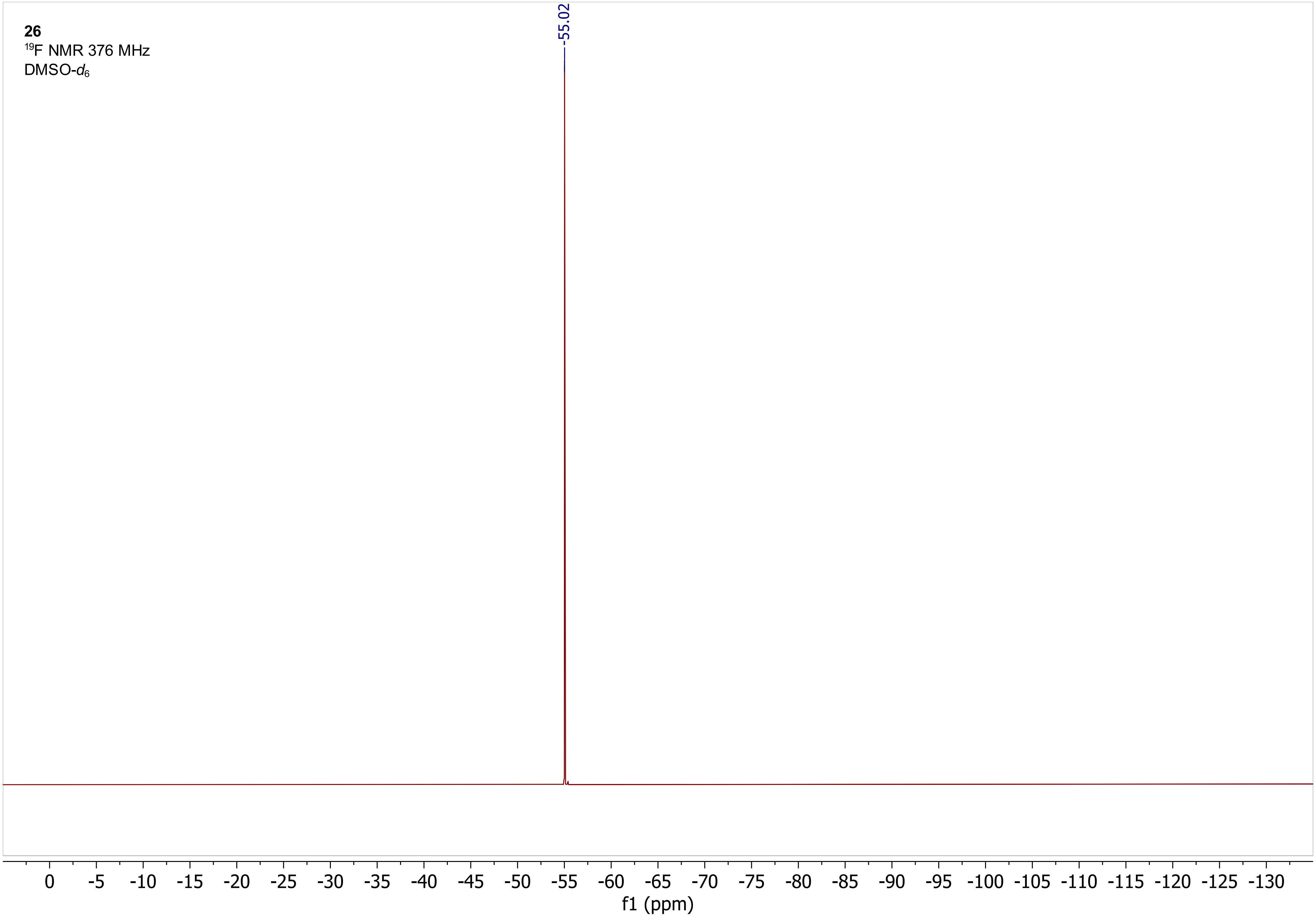
26
19F NMR 376 MHz
DMSO-d6
-55.02
0 -5 -10 -15 -20 -25 -30 -35 -40 -45 -50 -55 -60 -65 -70 -75 -80 -85 -90 -95 -100 -105 -110 -115 -120 -125 -130
f1 (ppm)
